# Atomic-Resolution Prediction of Degrader-mediated Ternary Complex Structures by Combining Molecular Simulations with Hydrogen Deuterium Exchange

**DOI:** 10.1101/2021.09.26.461830

**Authors:** Tom Dixon, Derek MacPherson, Barmak Mostofian, Taras Dauzhenka, Samuel Lotz, Dwight McGee, Sharon Shechter, Utsab R. Shrestha, Rafal Wiewiora, Zachary A. McDargh, Fen Pei, Rajat Pal, João V. Ribeiro, Tanner Wilkerson, Vipin Sachdeva, Ning Gao, Shourya Jain, Samuel Sparks, Yunxing Li, Alexander Vinitsky, Xin Zhang, Asghar M. Razavi, István Kolossváry, Jason Imbriglio, Artem Evdokimov, Louise Bergeron, Wenchang Zhou, Jagat Adhikari, Benjamin Ruprecht, Alex Dickson, Huafeng Xu, Woody Sherman, Jesus A. Izaguirre

## Abstract

Targeted protein degradation (TPD) has emerged as a powerful approach in drug discovery for removing (rather than inhibiting) proteins implicated in diseases. A key step in this approach is the formation of an induced proximity complex, where a degrader molecule recruits an E3 ligase to the protein of interest (POI), facilitating the transfer of ubiquitin to the POI and initiating the proteasomal degradation process. Here, we address three critical aspects of the TPD process: 1) formation of the ternary complex induced by a degrader molecule, 2) conformational heterogeneity of the ternary complex, and 3) assessment of ubiquitination propensity via the full Cullin Ring Ligase (CRL) macromolecular assembly. The novel approach presented here combines experimental biophysical data—in this case hydrogen-deuterium exchange mass spectrometry (HDX-MS, which measures the solvent exposure of protein residues)—with all-atom explicit solvent molecular dynamics (MD) simulations aided by enhanced sampling techniques to predict structural ensembles of ternary complexes at atomic resolution. We present results demonstrating the efficiency, accuracy, and reliability of our approach to predict ternary structure ensembles using the bromodomain of SMARCA2 (SMARCA2*^BD^*) with the E3 ligase VHL as the system of interest. The simulations reproduce X-ray crystal structures – including prospective simulations validated on a new structure that we determined in this work (PDB ID: 7S4E) – with root mean square deviations (RMSD) of 1.1 to 1.6 Å. The simulations also reveal a structural ensemble of low-energy conformations of the ternary complex within a broad energy basin. To further characterize the structural ensemble, we used snapshots from the aforementioned simulations as seeds for Hamiltonian replica exchange molecular dynamics (HREMD) simulations, and then perform 7.1 milliseconds of aggregate simulation time using Folding@home. The resulting free energy surface identifies the crystal structure conformation within a broad low-energy basin and the dynamic ensemble is consistent with solution-phase biophysical experimental data (HDX-MS and small-angle x-ray scattering, SAXS). Finally, we graft structures from the ternary complexes onto the full CRL and perform enhanced sampling simulations, where we find that differences in degradation efficiency can be explained by the proximity distribution of lysine residues on the POI relative to the E2-loaded ubiquitin. Several of the top predicted ubiquitinated lysine residues are validated prospectively through a ubiquitin mapping proteomics experiment.

## 1 Introduction

Heterobifunctional degraders are a class of molecules that induce proximity between a target protein of interest (POI) and a E3 ubiquitin ligase, which can lead to ubiquitination of the POI and its subsequent proteosomal degradation through a complex machinery of proteins.^1^ Degrader molecules provide the opportunity of a novel therapeutic modality as compared with traditional small molecule inhibitors – single molecules induce catalytic turnover of the POI and potentially offer an avenue for modulation of targets traditionally labeled as “undruggable” by classical therapeutic strategies.^2–4^ Heterobifunctional degraders consists of two separate protein binding moieties (the “warhead” and the “E3-ligand”) joined by a “linker”. The warhead binds to the POI (and we note that the degrader molecules studied here all have a non-covalently binding warhead) and the E3-ligand binds to an E3 ubiquitin ligase such as Cereblon (CRBN),^5–7^ cIAP,^8^ KEAP1,^9^ von Hippel-Lindau protein (VHL),^10–12^ or, potentially, to any of the more than 600 known E3 ubiquitin ligases.^13^ The ternary complex induced by the E3-ligand-linker-warhead degrader molecule is critical for bridging the interactions between the POI and a ubiquitin ligase (which can be the native *or* a non-native degradation partner of the POI). An important consideration when assessing putative degrader molecules is the cooperativity of the ternary complex, i.e., the difference between the binding affinity of the ternary complex and the binary components, which can influence degradation efficiency. The cooperativity is thought to result from interactions across the induced interface of the POI-ligase pair.^14,15^

The formation of the POI-degrader-ligase ternary complex is central to the targeted protein degradation (TPD) process, but how the formation of the ternary structure impacts protein degradation is still poorly understood, especially given the dynamic nature of the complex.^16–19^ X-ray crystallography of the ternary complex^20^ provides a high resolution structure of a single conformational state, but a growing body of evidence suggests that the dynamic nature of the ternary structure may not be accurately represented by this lowest energy crystallization snapshot. For instance, a study of several heterobifunctional degraders found that different degraders displayed different degrees of efficiency, although the corresponding ternary complex structures are nearly identical, thus raising questions about the static structural representations of the ternary complex and degradation efficiency. Studies targeting the degradation of Burton Tyrosine Kinase (BTK) by CRBN or cIAP found that high degradation efficiencies can also be achieved through degrader molecules that induce a non-cooperative ternary complex, demonstrating a disconnect between binding affinity and degradation efficiency.^21,22^ It appears that for degraders that bind with relative weak affinity (1 uM) to either the target or the ligase, cooperativity is crucial to optimize degradation. On the other hand, for degraders with very high binding affinity (low nM) to the target or the ligase, cooperativity is less crucial.

This and other findings^23–25^ suggest that degradation efficiency is more complex than can be understood through the thermodynamics of binding or the analysis of static structures. As such, determining the dynamic ensemble of the ternary complex may reveal mechanistic insights to facilitate the design of more effective degrader molecules.^20,26–29^ Previous work to computationally predict ternary structures has primarily consisted of protein-protein docking protocols with rigid protein structures, possibly followed by refinement of the initial structures with molecular dynamics (MD) simulations to assess the stability of the predicted models.^28–34^ However, these docking protocols fail to predict experimentally determined structures with high fidelity and they neglect the aforementioned dynamic nature of the ternary structure, highlighting the challenge associated with the generation of ternary structure models.

Recently, Eron et al. demonstrated how ternary complex structures of BRD4 do not represent the biologically relevant conformer of the ternary complex induced with CRBN, as demonstrated using HDX-MS. Molecular modeling revealed the dynamic nature and alternative conformations, which helped explain the dramatically increased cooperativity, ternary complex formation and degradation of their molecule CFT-1297 compared to the literature standard, dBET6.^35^ The authors use experimental data to improve protein-protein docking predictions, but they admit that the high flexibility of degrader-induced ternary complexes impedes a complete description of the bound conformations using their approach.

The goal of our work here is to understand the structural and dynamic basis of targeted protein degradation and ultimately design molecules for synthesis. We specifically focus on three different VHL-recruiting degraders of SMARCA2, for which crystal structures exist. PROTAC 1 (PDB ID: 6HAY) and PROTAC 2 (PDB ID: 6HAX) have been solved previously and ACBI1 (PDB ID: 7S4E) was solved and deposited as part of this work. The cooperativities and degradation efficiencies for each of these molecules is summarized in Table 1. We carry out MD simulations in combination with hydrogen-deuterium exchange mass-spectrometry (HDX-MS), shedding light on the dynamics of the ternary complexes beyond what is provided by static crystal structures. Specifically, we use “protection data” derived from HDX-MS as collective variables in weighted-ensemble MD simulations that predict ternary complex conformations, enhancing both the speed and accuracy of the computational predictions. We also show the usefulness of HDX-MS data as constraints for protein-protein docking when higher throughput and lower resolution models are sought, such as when screening many degrader molecules. Furthermore, we introduce methods that includes long-timescale MD simulations augmented with small-angle X-ray scattering (SAXS) data and Markov state modeling to determine the conformational free energy landscapes of the ternary complexes, which is the foundation for quantifying the populations of different conformational states. Finally, as an example of downstream use of these models, we assemble the entire cullin-RING ligase (CRL) to explore structural and dynamic factors that may be associated with ubiquitination. Mass spectrometry-based proteomics experiments validate the predicted ubiquitination of several lysines of SMARCA2 induced by ACBI1, supporting the use of the CRL model as a criterion for explaining degradation.

**Table 1:** Binding affinity (*K_d_*), efficiencies (IC50, DC50), and cooperativity (*α*) of PROTAC 1, PROTAC 2, and ACBI1 degraders. Ternary IC50 and binary (SMARCA2) DC50 values are reported; the cooperativity is the ratio of binary over ternary IC50. Table adapted from Farnaby et al.^36^

|  | **K<sub>d</sub>, VHL(nM)** | **K<sub>d</sub>, SMARCA2(nM)** | **IC50(nM)** | **DC50(nM)** | **Dmax(%)** | **$\alpha$** |
| --- | --- | --- | --- | --- | --- | --- |
| **PROTAC 1** | 98 ± 26 | 4500 ± 480 | 205 ± 15 | 300 | 65 | 12 |
| **PROTAC 2** | 100 ± 10 | 770 ± 51 | 45 ± 9 | 70 | 90 | 18 |
| **ACBI1** | 250 ± 64 | 1800 ± 980 | 26 ± 3 | 6 | ≈100 | 30 |

This work offers unique insights into the dynamic nature of the ternary structure ensemble and that of the full CRL macromolecular assembly that could explain ubiquitination and downstream protein degradation. Our results can be used to guide the design of novel degrader molecules that induce a productive ternary complex ensemble. In particular, having a small set of high-population ternary complex structures can provide an avenue for structure-based degrader discovery, particularly focused on the design of linkers that improve drug-like properties of the degrader molecule while maintaining or improving the aspects of the ternary structure ensemble that lead to ubiquitination. We make the simulation and experimental results available to the research community, including source codes, the release of a new X-ray crystal structure of ACBI1 connecting the bromodomain of SMARCA2 to VHL that has been deposited into the Protein Data Bank (PDB ID: 7S4E), and the release of the HDX proteomics and ubiquitin mapping proteomics. Data are available via ProteomeXchange with identifiers PXD033849 and PXD033763.

## 2 Results

### 2.1 Degraders with different efficiency induce similar ternary complex structures in X-ray crystallography

The ternary complexes of the bromodomain of SMARCA2 isoform 2 (iso2-SMARCA2*^BD^*) and the VHL/ElonginC/ElonginB (VCB) complex induced by different heterobifunctional degraders have been studied extensively.^28,37^ In particular, PROTAC 1, PROTAC 2, and ACBI1 are three degrader molecules that induce a ternary SMARCA2*^BD^*:VCB complex with quite different degradation efficiencies (see Table 1). Whereas crystal structures of the ternary complexes induced by PROTAC 1 (PDB ID: 6HAY) and PROTAC 2 (PDB ID: 6HAX) exist, none has been reported to date for ACBI1, the most potent degrader among them. Thus, we determined the structure of SMARCA2*^BD^*:VHL liganded by ACBI1 via X-ray crystallography. The structure was obtained by hanging drop vapor diffusion (see Methods 4.2)^28^ and solved by molecular replacement to 2.25 Å in the highest resolution shell (Supplemental Table 1), using the PROTAC 2 crystal structure (PDB ID:6HAX) as the search model (Fig. 1a).

**Fig. 1:**
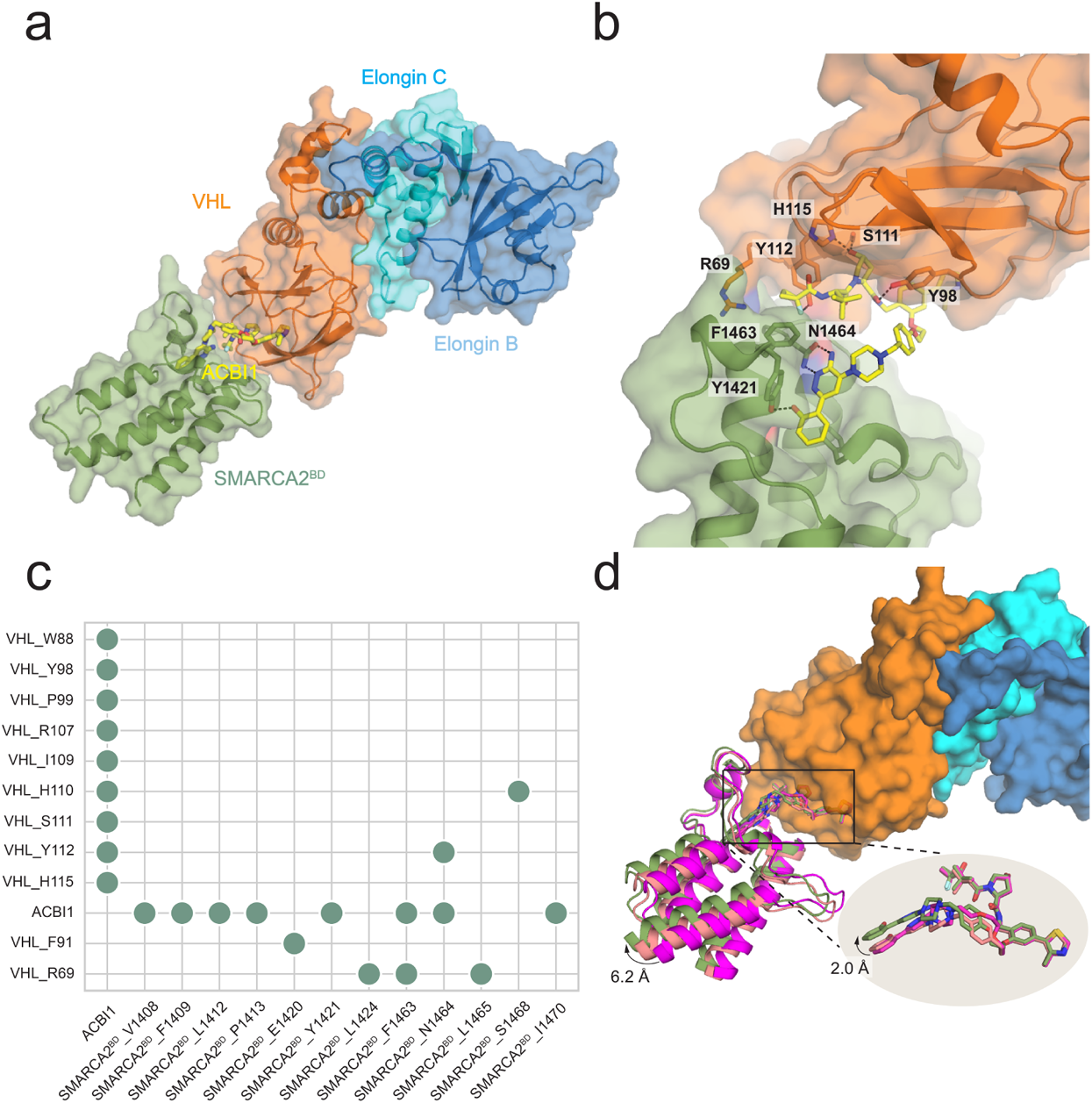
Ternary complex of SMARCA2*^BD^* and VHL/Elongin C/Elongin B (VCB) induced by ACBI1 shows structural similarities with PROTAC 1 and PROTAC 2: a) Overall perspective of iso2-SMARCA2*^BD^* and the VCB complex induced by degrader molecule ACBI1 (shown as yellow stick representation). b) ACBI1-induced interface contacts between SMARCA2*^BD^* and VHL. Annotated residues are among those that make the highest number of contacts (see panel c). c) A contact map for the interface of the crystal structure (obtained by the Arpeggio software^38^). Contacts are indicated when ≥ 10 atomic contacts (i.e., distance ≤ 4.5 Å) are present. d) Superposition of the crystal structures of PROTAC 1 (PDB ID: 6HAY, purple), PROTAC 2 (6HAX, salmon), and ACBI1 (7SE4, green) by aligning VHL (orange surface representation) shows varied conformations of the warheads of the three degraders (up to 1.7 Å), resulting in alterations of SMARCA2*^BD^* within the ternary complex.

ACBI1 bridges the induced interface, forming contacts with both proteins. Importantly, the degrader induces favorable contacts across the non-native interface, such as VHL:R69 and SMARCA2*^BD^*:F1463 (Fig. 1b,c). SMARCA2*^BD^*:N1464 maintains critical bivalent contacts to the aminopyridazine group of ACBI1, positioning the terminal phenol group for pi-stacking interactions with residues F1409 and Y1421 (Fig. 1b,c). On the ligase side of the interface, the interactions between Y98 and ACBI1 are consistent with those between the same residue and PROTAC 1 or PROTAC 2 (Fig 1b,c).^28^

Despite differences in the linker compositions, the protein-protein interface induced by ACBI1 is structurally similar to that induced by PROTACs 1 or 2^28^ (see Fig. 1d). A slight 1.7 Å twist of ACBI1 compared to the other two degraders, which can be ascribed to their minor differences (e.g. the ACBI1 linker has one additional ether group compared to the PROTAC 2 linker), results in a subtle “swing” of the protein in the crystal structure (Fig. 1d). However, the protein-protein interface remains the same (Supplemental Fig. 1), and the structural differences do not align with the markedly different degradation efficiency obtained^36^ suggesting that the (dynamic) ensemble of ternary complex structures may be fairly different among them and responsible for the degradation differential. Consistent with other studies,^22,39^ this implies that “crystallographic snapshots” are not suitable to provide a holistic view of the ensemble of all possible ternary complex structures in solution, but merely represent a subset of the relevant conformations favored by crystallization.^40^

### 2.2 Hydrogen Deuterium Exchange Reveals Extended Protein-Protein Interfaces

In order to assess the impact of different degrader molecules on the dynamic nature of the SMARCA2*^BD^*:VHL interactions, we performed hydrogen-deuterium exchange (HDX) experiments on the respective APO, binary and ternary (complex) species, thus characterizing the induced protein-protein interface in solution.^35^ This approach is a promising alternative to previous attempts at characterizing degrader ternary complexes that employed multiple crystal structures,^39^ NMR,^22^ and SAXS coupled with various forms of modeling. Additionally, there exists a wealth of knowledge for the integration of HDX-MS coupled with computational modeling.^41,42^ Importantly, changes in the rate of deuterium incorporation are dependent on factors like pH, temperature, solvent occlusion and molecular interactions like hydrogen bonding.^43^ We control the temperature and pH using robotics systems that enable precise temporal control over D_2_O exposure, probing the effects of (binary and ternary) complex formation on hydrogen bonding and solvent exposure. To ascertain the changes in solvent protection in the binary or ternary complex, the uptake of the APO or binary species is subtracted from that of the corresponding binary or ternary states (referred to as BinaryΔAPO and TernaryΔBinary), respectively. The results are summarized in difference plots that highlight the statistically significant (95% or 98% confidence interval) changes in deuterium uptake (see Supplemental Fig. 35a-d for the SMARCA2*^BD^*:VCB complex induced by ACBI1).

Fig. 2a reveals that large regions of SMARCA2*^BD^* become protected upon ternary complex formation induced by ACBI1 (see TernaryΔBinary difference plot). These stretches of protected residues, e.g. amino acids 1409-1422 and 1456-1470, overlap with the warhead binding site based on the ternary complex structure published in this work (7S4E) and those published previously (6HAY, 6HAX), which confirms the similarity of the ternary complex interface among the three degrader molecules discussed above. Additionally, there are also stretches of protected amino acids, 1394-1407, that are too distant from the established binding interface to result from complex formation (Fig. 2a and f). Interestingly, the BinaryΔAPO difference plot suggests that, under our experimental conditions, the warhead concentration is close to the dissociation constant KD = 10*µ*M,^28^ as there is minimal difference between the exchange of SMARCA2*^BD^* in presence and in absence of the warhead due to the mixed population of free SMARCA2*^BD^* in solution outweighing the signature generated from the bound state (Fig. 2a and c).

**Fig. 2:**
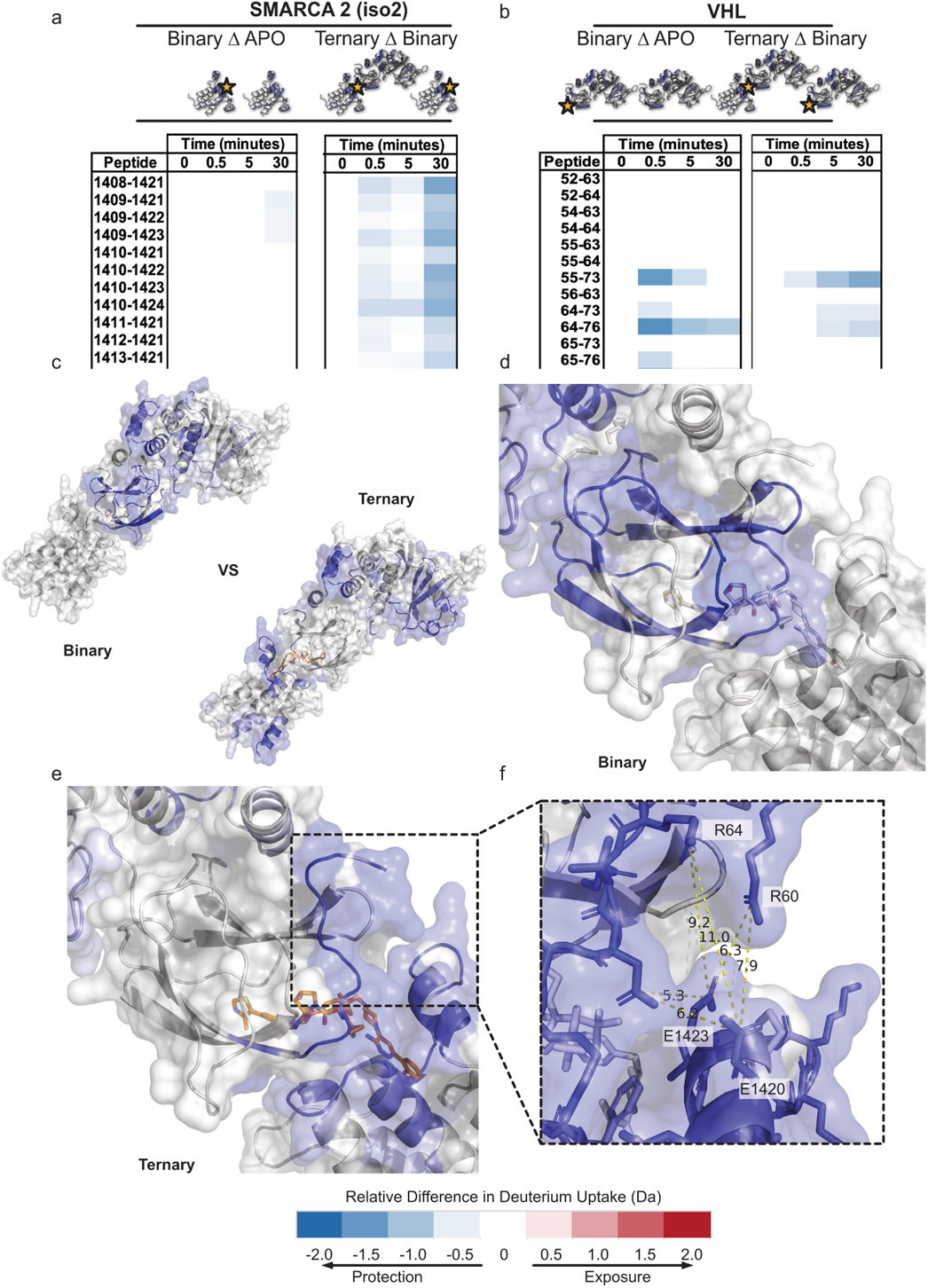
HDX-MS extends the ACBI1-induced SMARCA2*^BD^*:VHL compared to crystallographic data. a) SMARCA2*^BD^* HDX difference plots covering residues 1400-1423. Binary as compared to the APO, and ternary as compared to the binary states reveal increased protection induced by the presence of ACBI1 and VCB complex. b) Binary compared to APO and ternary compared to binary states of the VHL subunit highlighting extended exchange patterns due to the presence of the ternary complex. c) Exchange patterns induced by the binary and ternary forms of the complex superimposed on the crystal structure (PDB ID: 7S4E). d) Binary-specific induced HD exchange near the ligand binding site of VHL and SMARCA2*^BD^*. e) Ternary-specific induced HD exchange near the ligand binding site of VHL and SMARCA2*^BD^*. f) Proposed solution-state extended protein interface that may take advantage of salt-bridge interactions to increase cooperativity of the protein-protein complex.

Large regions of VHL are protected in the presence of the E3-ligand as indicated by the BinaryΔAPO difference plot (Fig. 2b and e). The most protected residues in the binary state are centered around amino acids 87-116, which include all 9 residues in the E3-ligand binding site of VHL. In the presence of SMARCA2*^BD^* (see Fig. 2b, TernaryΔBinary difference plot), much of the allosteric network due to E3-ligand binding can be subtracted away leaving only the most significantly protected residues induced by ternary complex formation (Fig. 2b and d). In particular, residues 60-72, which house the critical interaction of R69 show significant protection due to ternary complex formation (Fig. 2b and d). Moreover, we observe protection of residues 166-176 and residues 187-201 on VHL (see Supplemental Fig. 35b and f) as well as some regions on Elongin B and C that show protection upon ternary complex formation (see Supplemental Fig. 35c and d). Although these sites are distal from the binding interface, they spatially align with one another when grafted onto the structure (Fig. 2c) potentially uncovering a critical network of allosteric changes^44^ induced by ACBI1 that may play a role in downstream positioning of SMARCA2 to the E2 enzymes in the CRL complex.

The difference between HDX-MS binary and ternary SMARCA2*^BD^* experiments reveals that the interactions at the protein-protein interface help stabilize the ternary complex. Many of the charged interface residues, that are solvent-exposed and outside the range of traditional hydrogen bonding or salt-bridge interactions (*>* 6.3 Å) in the corresponding X-ray crystal structure (e.g. K1416, E1420, E1423 on SMARCA2*^BD^* and R60, R64 on VHL) are determined to be protected based on the HDX-MS results (Fig. 2e). In fact, the protected, charged interface residues of SMARCA2*^BD^* lie outside the direct ligand binding pocket in the crystal structure of the ternary complex. Interestingly, R60 through R64 on VHL are protected in the ternary complex for a longer duration than in the binary complex alone. This enhanced protection across the interface suggests that conformational rearrangements are responsible for protein-protein interactions. Our simulations presented below (Section 2.6) support this hypothesis, finding contacts between several of these charged interface residues. Taken together, these results underscore the importance of cooperativity driving the formation of the ternary complex for ligases with poor binding affinity to the POI.

Interestingly, we find that iso1-SMARCA2*^BD^*:ACBI1:VCB shows a slightly different protection pattern from iso2-SMARCA2*^BD^*:ACBI1:VCB, mainly in that residues G104 through L116 of VHL show significant protection in the former compared to the latter ternary complex. In our crystal structure of the iso2-SMARCA2*^BD^*:ACBI1:VCB system, these protected residues are close to the site where the additional 17 residues of iso1-SMARCA2*^BD^* appear, suggesting that the protected residues in VHL may be interacting with these residues that are not present in iso2-SMARCA2*^BD^*. Consistent with this hypothesis, residues I1414-N1417 of the iso1-SMARCA2*^BD^* extension show some protection in the ternary complex.

Studying the solution-state dynamics of degrader ternary complexes uncovers key details that are missed by crystallographic “snapshots” alone. As many of the crystallographic contacts are nearly identical between the different degrader molecules, many key interactions may be underrepresented in the crystal structure. Utilizing HDX-MS information, or other data derived from solution-state experiments, as restraints in modeling and simulation opens a pathway from a single accepted protein structure to a vast ensemble of conformations. Production of accurate ternary complex ensembles enables alternative routes for the design, optimization, and mechanism-of-action studies of heterobifunctional degraders.

### 2.3 HDX data enhance weighted ensemble simulations of ternary complex formation

We simulate the formation of iso2-SMARCA2*^BD^*:VHL degrader ternary complexes using weighted ensemble (WE) simulations, where a set of weighted trajectories are evolved in parallel along pre-defined collective variables, providing a means to compute non-equilibrium properties and predict likely binding pathways.^45,46^ This path-sampling strategy can sample rare events by orders of magnitude more efficiently than conventional MD simulations and it has been employed before for tasks such as protein-protein^47^ and protein-ligand binding.^48^ It is noteworthy, however, that our simulations are not informed by any structural data about the ternary complex interface from X-ray crystallography experiments.

Starting from a dissociated configuration, in which the degrader molecule is bound to VHL, yet both are clearly apart from SMARCA2*^BD^* (initial separation distance ~ 20 Å), the formation of ternary aggregates is simulated yielding complexes with interface structures well comparable to those obtained experimentally or in the low free energy basins to which experimental structures belong. As HDX experiments show, and our simulations of ternary complexes below confirm, the ternary complex exists as a dynamic ensemble of multiple conformations, of which the X-ray structure is a snapshot. Thus, we assess the quality of bound complexes by the minimum interface-RMSD (I-RMSD)^49^ of each simulated aggregate with respect to a set of structurally diverse reference ternary structures (see Supplemental Fig. 4 for SMARCA2*^BD^*:PROTAC 2:VHL). This set of distinct structures is obtained from long-timescale (*>* 1 *µs*) brute-force MD simulations, thus allowing a comparison to a variety of possible ternary complexes and not merely to a single experimental reference structure. We provide detailed descriptions of the methodology and the evaluation of all simulations performed in the Supplemental Information (SI). For these simulations, we use a collective variable defined by the number of atomic contacts and the warhead-RMSD (w-RMSD) with respect to the crystal structure of the binary target-warhead complex (see Methods 4.7).

Protein-protein encounter complexes, i.e., the formation of protein contacts, are usually observed within 500 ns of aggregate simulation time. An ensemble of about 500 bound ternary complexes with a minimum I-RMSD *<* 2 Å can usually be obtained after ~ 2 *µs*, which takes ~ 12 days using a single A40 GPU per simulation, but it is highly parallelizable to more GPUs.

Remarkably, when introducing as a collective variable the number of contacts formed by the protected residues, as determined by the HDX-MS experiments described above, the prediction accuracy of ternary complex formation is significantly improved compared to simulations in which any protein-protein contacts were considered (see Supplemental Figs. 2, 3). Supplemental Movie 1 shows the continuous trajectory of one such ternary complex binding event, where the addition of protected-residue contacts enhances the ternary complex binding.

**Fig. 3:**
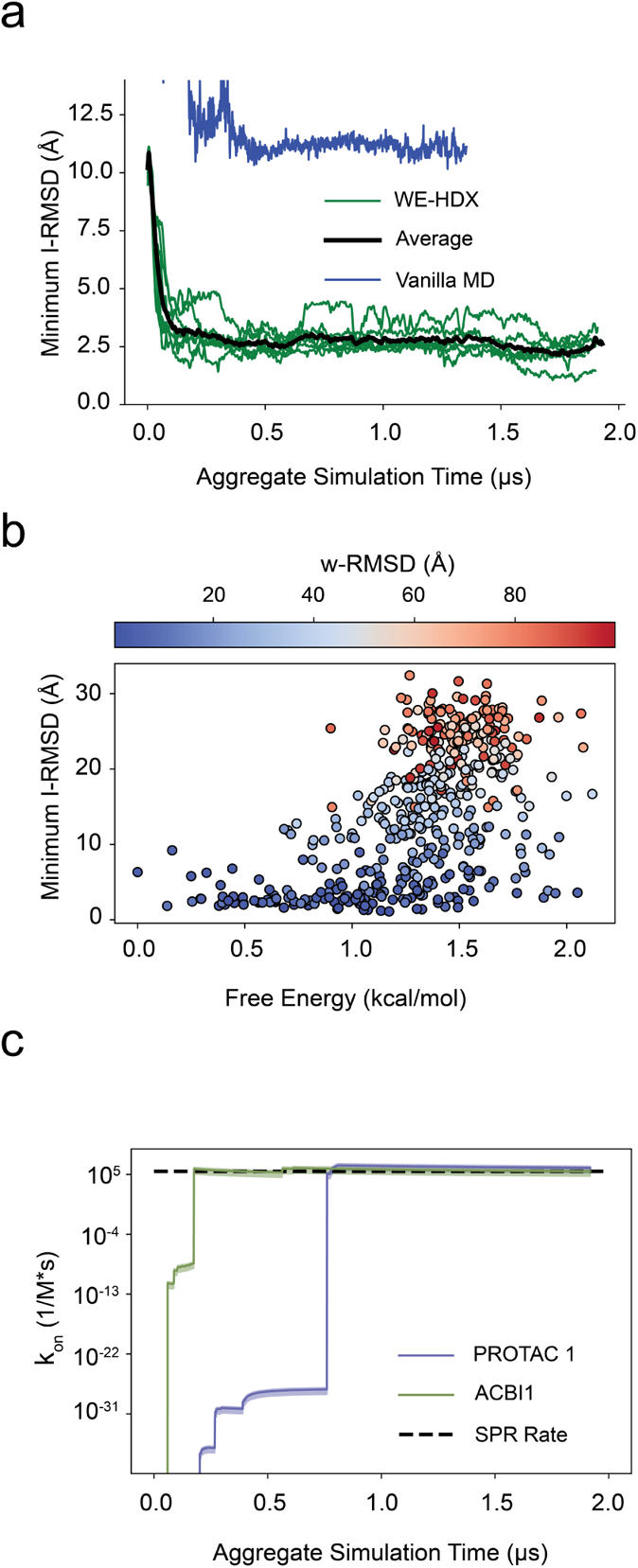
Assessing ternary complex formation. a) The minimum I-RMSD over time during the WE-HDX simulations of the PROTAC 2 system. Each green line indicates one replica and the black line is the average between all runs. The blue line indicates the minimum I-RMSD for a vanilla molecular dynamics simulation. b) A scatter plot of the free energy vs the minimum I-RMSD of each of the 500 clusters from the PROTAC 2 simulations. The circles are colored by w-RMSD. c) The predicted binding rates for the PROTAC 1 system (purple) and the ACBI1 system (green). The black line is the experimental binding rate determined via SPR.

We note that the use of HDX-MS data in our approach is rather qualitative, as the simulations are solely informed by the existence of specific interaction sites and not by the degree of those interactions. HDX rate constants are not estimated during the simulations as often performed in quantitative approaches that combine HDX-MS experiments with simulation.^50^ Rather, our method falls in the category of simulations *guided* by HDX-MS data, in which qualitative correlations between simulation and experiment are attempted to be established^42^ (see discussion in the SI). We present a particularly interesting example of synergy between molecular simulations and HDX-MS experiments, in which the path-sampling algorithm is furnished with a fairly simple parameter derived from the experimental measurements, i.e., the contact numbers between distinct sites. We call this integrated approach WE-HDX. Despite its simplicity, WE-HDX seems particularly appropriate for the formation of ternary complexes that have distinct contacts across their binding interface.

To systematically study the formation of SMARCA2*^BD^*:VHL ternary complexes with all three degraders, we run seven independent WE-HDX simulations with PROTAC 2 for an aggregate simulation time of 12.5 *µs* and three such simulations totaling ~ 6 *µs* for both PROTAC 1 and ACBI1. The difference in simulations corresponds to the greater flexibility of the PROTAC 2 ternary complexes, compared to the other two degraders. Ensembles of bound ternary complexes were formed with minimum I-RMSDs of 0.5 Å for ACBI1, 0.7 Å for PROTAC 1, and 1.1 Å for PROTAC 2, respectively.

To highlight the sampling ability of WE-HDX simulations, Fig. 3a compares the minimum I-RMSD of the SMARCA2*^BD^*:PROTAC 2:VHL simulation with that from vanilla MD simulations of the same system as a function of aggregate simulation time. While the minimum I-RMSD converges to 2.5 Å in the WE-HDX simulations within 0.5 *µs* of aggregate simulation time (Fig. 3 A), that for the vanilla MD remains as high as 10 Å after 1.4 *µs* of simulation.

The very high prediction accuracy of the WE-HDX simulations is illustrated for the SMARCA2*^BD^*:PROTAC 2:VHL system in Fig. 4. One example of a predicted structure is visualized in Fig. 4a,b. The contact maps presented in Fig. 4c compare the ternary interface of the experimental crystal structure to that of the minimum I-RMSD structure produced by the WE-HDX simulations. Each point reflects the degree of interaction, revealing an interaction pattern from the WE-HDX simulations that is comparable to that from experiment. The near-perfect alignment (minimum I-RMSD = 1.1 Å) of one sampled conformation with the crystal structure shown in Fig. 4d further emphasizes that the interactions of degrader ternary complexes observed experimentally can be recaptured by WE-HDX.

**Fig. 4:**
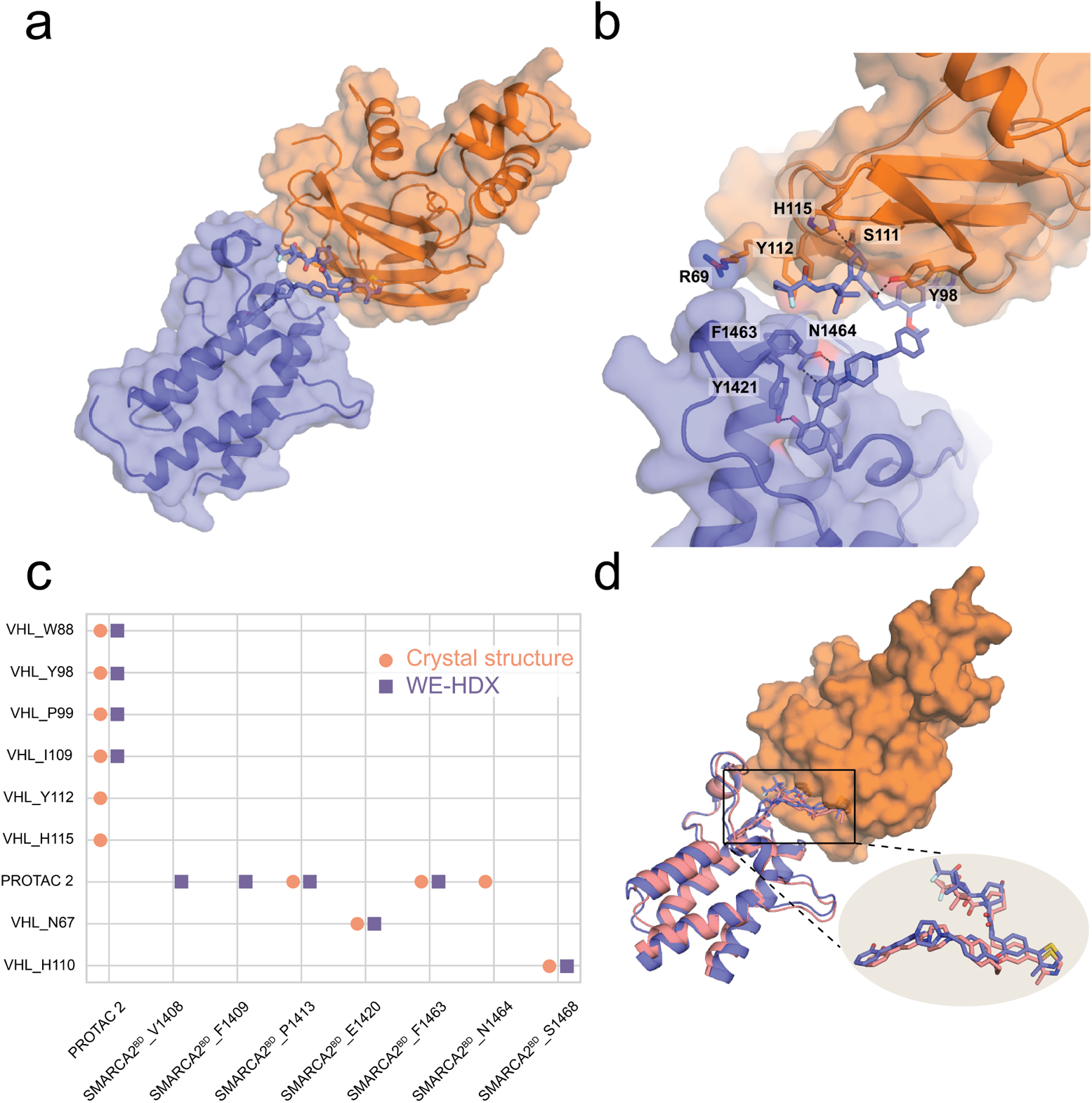
Illustration of one representative prediction of SMARCA2*^BD^*:PROTAC 2:VHL produced by WE-HDX simulations and its comparison to the crystal structure (PDB ID: 6HAX). a) A simulated ternary structure with minimum I-RMSD = 1.1 Å. SMARCA2*^BD^* (purple) and VHL (orange) are shown in cartoon and transparent surface representations and PROTAC 2 is shown in in stick representation. b) Structural details of the binding interface. Annotated residues are among those that make the highest number of contacts (see panel c). c) A contact map of the interfaces from the crystal (salmon) and the simulated structure (purple). d) Structural alignment of the simulated (purple) with the crystal structure (salmon) with a detailed PROTAC 2 comparison.

Six out of seven of the SMARCA2*^BD^*:PROTAC 2:VHL simulations observed binding events for a total of 3278 unique observations. In order to assess the degree of heterogeneity within this ensemble, we clustered the WE results into 500 macrostates with a k-means algorithm using the *Cα − Cα* distances between the ligase and target protected residues. As expected, all states with a low minimum I-RMSD have low values of w-RMSD too (Fig. 3b). States with high free energies, i.e., above 1.5 kcal/mol, have large minimum I-RMSDs, ranging from 1.5 to 30 Å. However, the minimum I-RMSD distribution among the 20 low free energy states below 0.5 kcal/mol is significantly tighter, ranging from 1.1 to 9.2 Å with an average value of 3.7 Å and 12 out of 20 states even having a minimum I-RMSD below 3 Å.

We predict ternary complex binding rate constants for the three different degraders directly from WE-HDX simulations using the probability flux into a bound state (minimum I-RMSD *<* 2 Å). While the predicted rates for PROTAC 1 and ACBI1 are on the same order of magnitude as in experiments (Fig. 3c), we predict a significantly slower binding rate for PROTAC 2, which is not yet determined experimentally (see Table 2). However, for all three rates there are large uncertainties, as has previously been observed in WE rate calculations.^51,52^ Better statistics can be achieved by longer simulation times or the use of recently proposed algorithms that converge these rates more efficiently,^53,54^ which is beyond the scope of this work.

**Table 2:** Comparison of ternary complex binding rate constants between simulation and experiment for the PROTAC 1, PROTAC 2, and ACBI1 systems. The experimental rate for PROTAC 2 has not been determined yet.

| degrader | Predicted Rate ($M^{-1}s^{-1}$) | Experimental Rate ($M^{-1}s^{-1}$) |
| --- | --- | --- |
| PROTAC 1 | $10 * 10^5 \pm 8 * 10^5$ | $2.9 * 10^5$ |
| PROTAC 2 | $2.2 * 10^2 \pm 1.7 * 10^2$ | N/A |
| ACBI1 | $3 * 10^5 \pm 2 * 10^5$ | $2.4 * 10^5$ |

In most of the analysis above, we have used the minimum I-RMSD with respect to a set of reference structures, as described, to assess the quality of structures obtained from WE-HDX simulations. Alternatively, the C*_α_*-RMSD of the entire ternary complex has been used before as a parameter to gauge their prediction accuracy.^47^ Supplemental Fig. 3b shows that the interface-RMSD, and, in particular, the threshold at 2 Å is indeed an appropriate metric for the identification of ternary complexes, as all such complexes formed in our WE-HDX simulations of the system with PROTAC 2 have a minimum I-RMSD *<* 2 Å for a C*_α_*-RMSD *≤ ∼*5 Å, which is clearly below the threshold used in other studies (e.g. C*_α_*-RMSD *≤* 10 Å used by Drummond et al.^34^).

As in most design projects X-ray structures may not be readily available, it is important to determine the usefulness of predictive features that do not depend on ternary complex X-ray structures. To this end, we filtered the ensemble of simulated SMARCA2*^BD^*:PROTAC 2:VHL structures for bound complexes with warhead-RMSD *<* 2 Å and *>* 30 contacts between protected residues (see Fig. 5a). Among these, the bulk of the density was limited to minimum I-RMSD values between 1 and 4 Å, with 90% below 3 Å and 43% even below 2 Å (see Fig. 5b), indicating that observables such as the warhead-RMSD and the number of contacts between protected residues can be used to characterize bound ternary complexes.

**Fig. 5:**
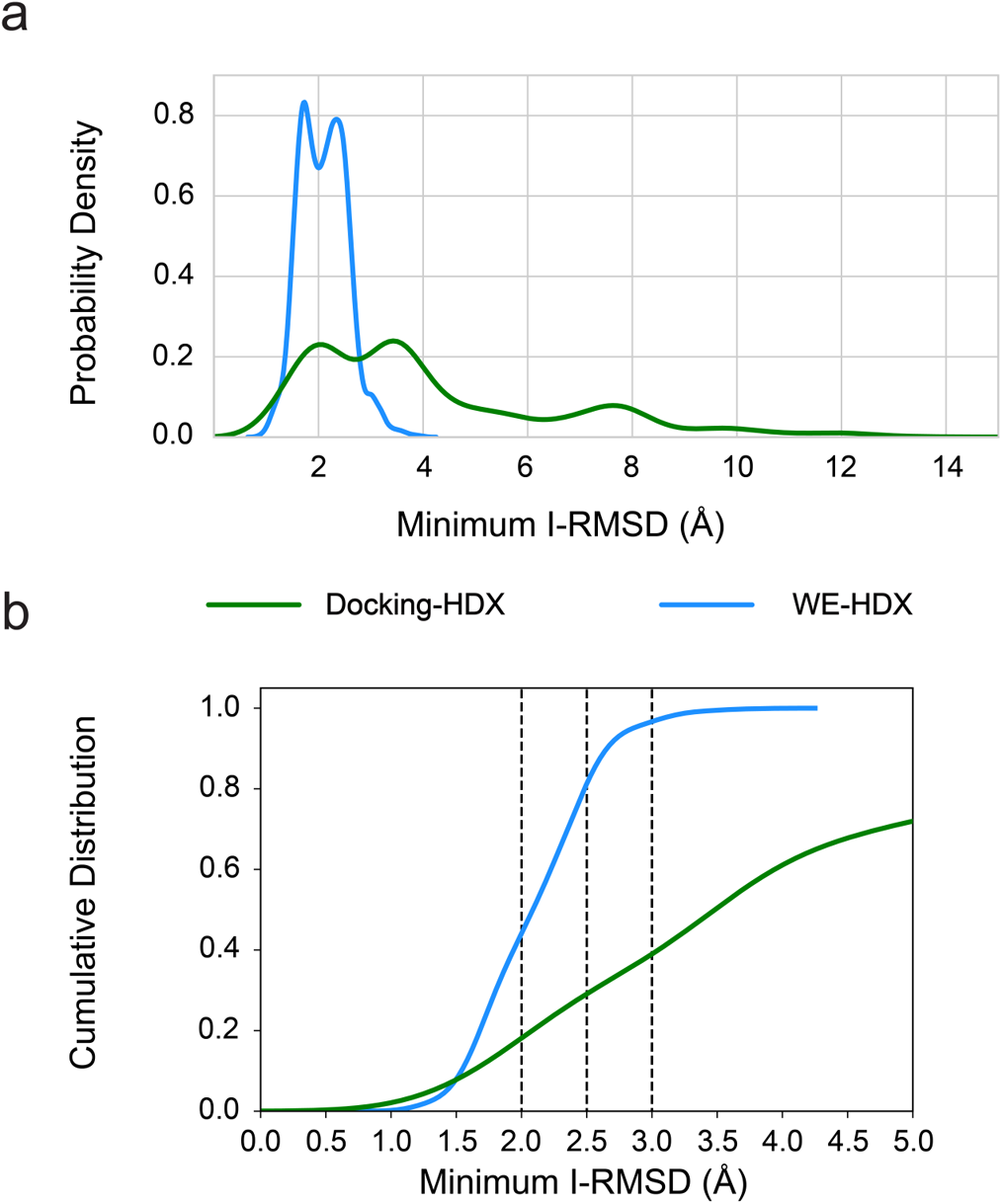
Comparing the bound ensembles determined by docking and WE simulations with information from HDX-MS for the PROTAC 2-induced ternary complex. Simulated structures with a warhead-RMSD ≤ 2 Å and *>* 30 contacts between the SMARCA2*^BD^* and VHL interface are considered bound, whereas the docked bound structures are determined as the top-100 from Rosetta-scoring. a) Probability densities of minimum I-RMSD values for the bound ensembles from WE-HDX and Docking-HDX. b) Cumulative distributions of minimum-I-RMSD values for the probability densities shown in panel a, illustrating that larger ensembles of bound ternary complexes can be obtained from WE-HDX compared to Docking-HDX. The dashed vertical lines indicate three specific thresholds of minimum IRMSD (2 Å, 2.5 Å, and 3 Å), below which a complex can be considered bound.

Knowledge of a large number of degrader-induced ternary complexes is essential to understanding the structural and dynamic features that lead to targeted protein degradation. As the WE-HDX results reveal, the level of detail associated with such simulations allows an entire ensemble of ternary complexes, including many conformations with a pronounced protein interface, to be generated *ab initio*, i.e., even from a fairly dissociated state and with no additional information on the protein-protein binding pose. This is a significant achievement with regard to the design of effective degrader molecules, for which ternary complex structures are not obtained experimentally.

### 2.4 HDX-MS improves prediction of ternary complexes using docking

Several docking procedures to predict ternary complexes of degrader molecules have been described. Most of them have stages for generating protein-protein complexes in the absence of the degrader, linker, alignment of linker or whole degrader to the protein-protein complexes, and some sort of scoring (^30–32,34^). We used an approach comparable to that published by Bai et al.^31^

In contrast to recent work,^35^ our docking method uses HDX-MS data to impose additional distance restraints at the sampling stage (instead of post-sampling scoring). Also, differently from the distance restraints derived from chemical cross-linking experiments,^55^ our approach is based on the statistics of the length of the linker in a degrader molecule. Application of the HDX-MS data for re-ranking of the docking predictions, as described by Eron et al.,^35^ may lead to a more quantitative assessment of structures. Discussion of the interplay of HDX-MS-derived restraints and HDX-MS-based re-rankings in docking is beyond the scope of the present work.

We show that incorporating experimentally retrieved distance restraints into the docking protocol significantly improves its ability to predict ternary complexes of high quality (see detailed comparisons in Supplemental Figs 5 and 6). In particular, it is striking how strongly the incorporation of HDX-MS data can boost the accuracy of the docking protocol among the highest-ranked docking poses.

**Fig. 6:**
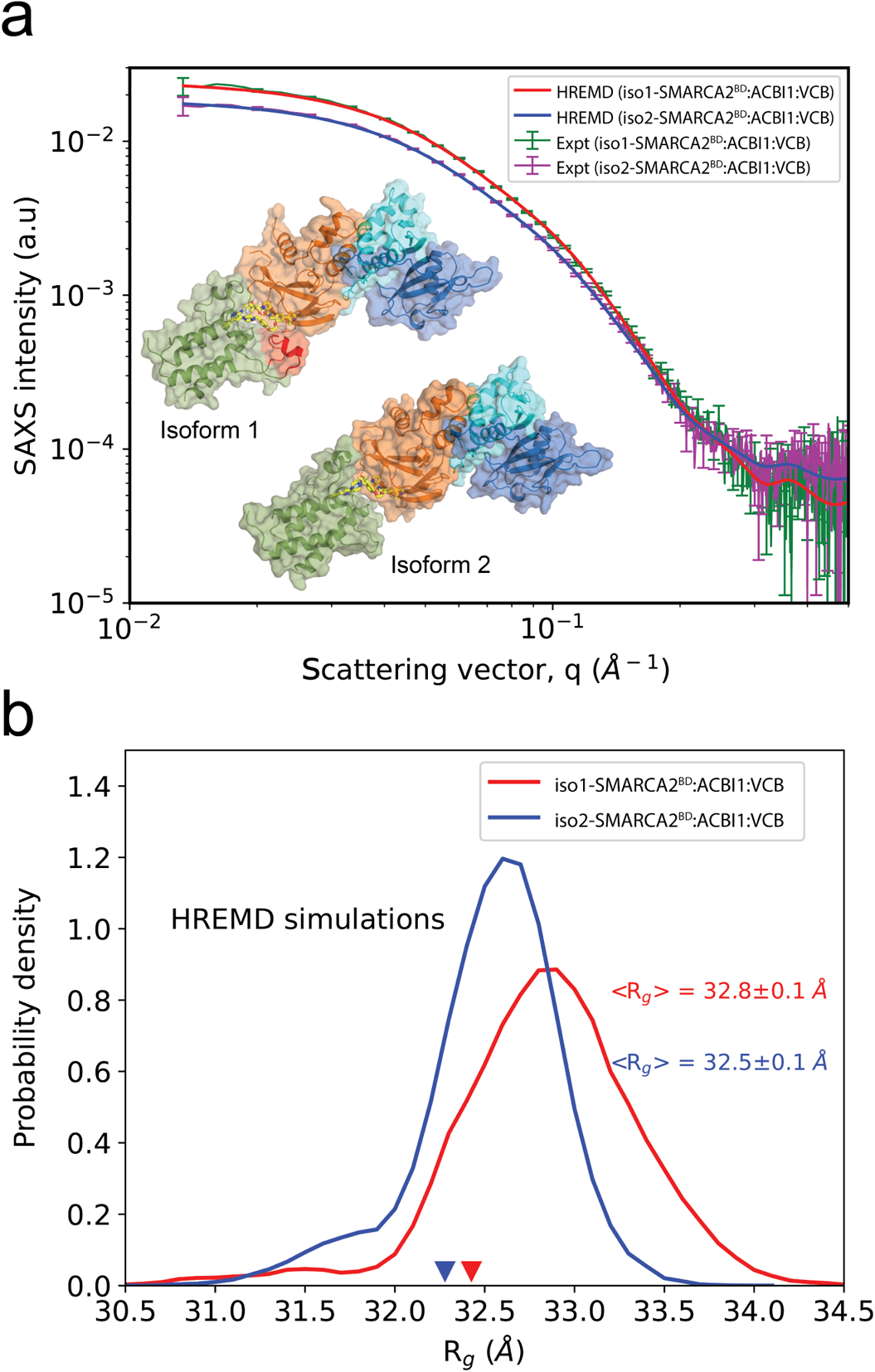
SAXS profiles and structural ensembles of iso1-/iso2-SMARCA2*^BD^*:ACBI1:VCB complexes. a) Comparison of theoretical and experimental SAXS profiles, SAXS intensity vs. *q*. b) The histograms of R*_g_* of iso1-SMARCA2*^BD^*:ACBI1:VCB (red) and iso2-SMARCA2*^BD^*ACBI1:VCB (blue) complexes calculated from HREMD simulations. The inverted red and blue triangles are the R*_g_* values of starting structures of iso1-/iso2-SMARCA2*^BD^*:ACBI1:VCB from homology model and crystallography respectively.

Although WE-HDX simulations consistently outperform the HDX-enhanced docking routine (see Fig. 5), docking, in combination with HDX-MS (Docking-HDX), is a useful tool for the quick filtering of a large number of degrader designs considering the significantly less computational cost of this approach (25 CPU hours for the generation of one ensemble compared to ~ 12 A40 GPU days for the WE-HDX method).

### 2.5 HREMD simulations and SAXS experiments reveal highly flexible ternary complex ensembles

The HDX-MS measurements revealed substantial flexibility, which is consistent with the structural diversity obtained from WE-HDX simulations and from the docking protocol of the SMARCA2*^BD^*:VHL ternary degrader-protein complexes studied here. To further enhance the exploration of their conformational heterogeneity, we perform atomistic Hamiltonian replica-exchange MD (HREMD) simulations based on the X-ray structures. HREMD is a parallel tempering simulation method that efficiently samples large conformational changes of proteins in aqueous solution and, therefore, is a promising strategy to study the protein-protein interactions and the flexibility of degraders in ternary complexes (see Methods 4.9). In particular, we simulate ternary complexes of both isoforms of SMARCA2*^BD^* connected only to the VHL subunit or, in order to be consistent with our experiments, to the larger VCB complex by PROTAC 1, PROTAC 2, or ACBI1 (see Supplemental Table 4 for a list of all HREMD simulations performed). The structure of iso1-SMARCA2*^BD^*, which is not experimentally resolved, is obtained by homology modeling with the iso2-SMARCA2*^BD^* structure used as template (see Methods 4.5). HREMD simulations with iso1-SMARCA2*^BD^* were performed to test whether they could explain the ternary complex protection differential observed between that isoform and Isoform 2. To ensure the HREMD-generated ensembles are accurate and reliable, we validate the simulations by directly comparing against the size exclusion chromatography coupled to small-angle X-ray scattering (SEC-SAXS) data, Fig. 6a.

The excellent agreement (*χ*^2^ = 1.55 and *χ*^2^ = 1.23 for iso1- and iso2-SMARCA2*^BD^*:ACBI1:VCB respectively, where *χ*^2^ is defined in Eq. 11) between SAXS profiles obtained from experiment and such calculated from simulations shows that the HREMD simulations capture the long timescale conformational ensembles to experimental accuracy. Furthermore, the ensemble-averaged *R_g_* of the two complexes from simulation are in excellent agreement to *R_g_* values obtained by Guinier approximation (Eq. 1) to experimental SAXS data (Supplemental Fig. 15), *R_g_* = 33.4*±*0.4 Å and 32.3*±*0.3 Å for iso1- and iso2-SMARCA2*^BD^*:ACBI1:VCB, respectively. The histograms of *R_g_* (calculated from atomic coordinates using Eq. 2) suggest that ternary complexes are flexible in solution leading to a change in overall conformation compared to their corresponding simulation starting structures, i.e., a homology model of iso1-SMARCA2*^BD^*:ACBI1:VCB and the crystal structure of iso2-SMARCA2*^BD^*:ACBI1:VCB (see Fig. 6b. These results illustrate the need for enhanced sampling methods, such as HREMD, to rigorously probe the conformational changes of the inherently flexible ternary degrader complexes.

To demonstrate the value of the HREMD simulations in aiding in the prediction of degrader efficacy, we analyze the thermodynamics of ternary complex formation by estimating a conformational free energy penalty for the binding of a fully-dissolved PROTAC 1, PROTAC 2, or ACBI1 to SMARCA2*^BD^* and VHL in a ternary complex. To this end, we simulate the individual degraders in solution (Methods 4.10), in addition to the ternary complex simulations presented above, and compare, as an observable proxy, the average linker end-to-end distance (normalized by the number of backbone atoms in the linker) of each degrader when fully dissolved to the corresponding value obtained when bound in a ternary complex. We observe that, in both environments, PROTAC 2 and ACBI1 adopt a significantly more expanded linker conformation compared to PROTAC 1 (see Supplemental Fig. 20), which has a lower SMARCA2-degradation efficiency than the other two degraders (see Table 1). This suggests, in accord with previous empirical findings,^34^ that degraders with extended linkers in solution more easily induce SMARCA2*^BD^*:VHL ternary complexes (Supplemental Fig. 20). We consider other uses of the ternary complex ensembles found with HREMD in the next section.

### 2.6 Structural determinants of degrader ternary complexes are revealed by long-timescale simulations

We quantify the free energy landscapes of several of the ternary complexes sampled in the HREMD simulations, namely iso2-SMARCA2*^BD^*:ACBI1:VHL, iso2-SMARCA2*^BD^*:PROTAC 1:VHL, iso2-SMARCA2*^BD^*:PROTAC 2:VHL, and iso1-SMARCA2*^BD^*:ACBI1:VCB. We begin this analysis by performing principal component analysis (PCA) decomposition of the distances between interface residues to identify high-variance collective variables (see Methods 4.11). The probability distribution of these high-variance features allows us to determine a more easily interpretable free energy landscape from our simulation data. We find that the landscape of each ternary complex contains several local minima differing by only a few kcal/mol (Fig. 8a and Supplemental Fig. 21).

Using *k*-means clustering in the PCA feature space, we then identify distinct clusters of conformations. Cluster centers roughly correspond to local minima in the free energy landscape (see Fig. 8a and Supplemental Fig. 21). These clusters of simulated conformations are consistent with our HDX-MS protection data: Fig. 7 shows that interface residues that were found to be protected in HDX-MS experiments are observed to interact in either the most populated or second most populated cluster. Notably, this analysis shows that in representative structures (namely the second most populated cluster centers) of iso1-SMARCA2*^BD^*:ACBI1:VCB, the helix formed by the 17-residue extension of iso1-SMARCA2*^BD^* interacts with a beta sheet of the VHL subunit, Fig. 7b, in accordance with our HDX-MS experiments that found this beta sheet to be protected in presence of iso1-SMARCA2*^BD^*, but not in the presence of iso2-SMARCA2*^BD^* (Fig. 7a). Similarly, representative structures from highly populated clusters of iso2-SMARCA2*^BD^*:ACBI1:VHL and iso2-SMARCA2*^BD^*:PROTAC 2:VHL show contacts between residues that were observed to be protected in HDX-MS experiments (see blue-colored regions in Figs. 7a,c), whereas such from the most populated cluster of iso2-SMARCA2*^BD^*:PROTAC 1:VHL Fig. 7e do not show these contacts. Representative structures from the most populated cluster of iso2-SMARCA2*^BD^*:degrader:VHL with all three degraders are displayed in Supplemental Fig. 22.

**Fig. 7:**
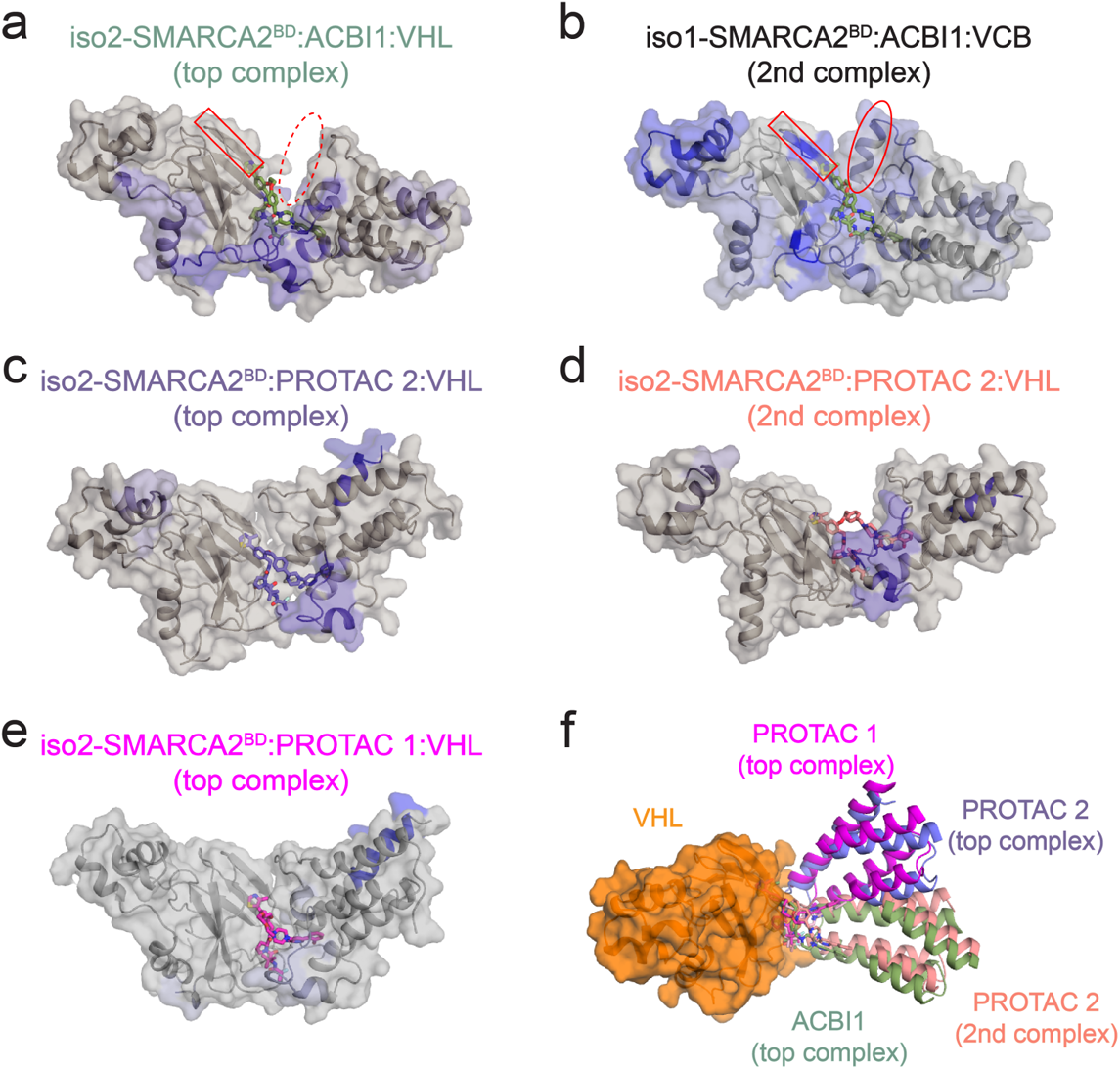
Most populated structures of SMARCA2*^BD^* bound to VHL with different degrader molecules, identified by dimension reduction and clustering of HREMD simulation data. a)-e) The blue-colored regions of SMARCA2*^BD^* and VHL represent HDX-MS protection in the presence of the corresponding degrader molecule relative to SMARCA2*^BD^*:VHL or SMARCA2*^BD^*:VCB complexes in the absence of the degrader. Representative structures from the second most populated clusters (2nd complex) of iso1-SMARCA2*^BD^*:ACBI1:VCB (panel b) and iso2-SMARCA2*^BD^*:PROTAC 2:VHL (panel d) support our HDX-MS results. In panels a and b, note that the beta sheet highlighted by a red rectangle does not show HDX-MS protection in iso2-SMARCA2*^BD^*:ACBI1:VHL, and does not contact VHL in simulations of that system. This region does show HDX-MS protection in iso1-SMARCA2*^BD^*:ACBI1:VCB, and we find in simulations that it forms contacts with an alpha helix that is only present in iso1-SMARCA2*^BD^* (indicated by a red oval). Note that Elongin B and Elongin C are included the simulations associated with in panel b, but omitted here for clarity. f) The ternary complexes from panels a, c, d, and e are compared after aligning VHL (orange surface representation) to illustrate the conformational heterogeneity among highly populated structures of ternary complexes with different degraders.

**Fig. 8:**
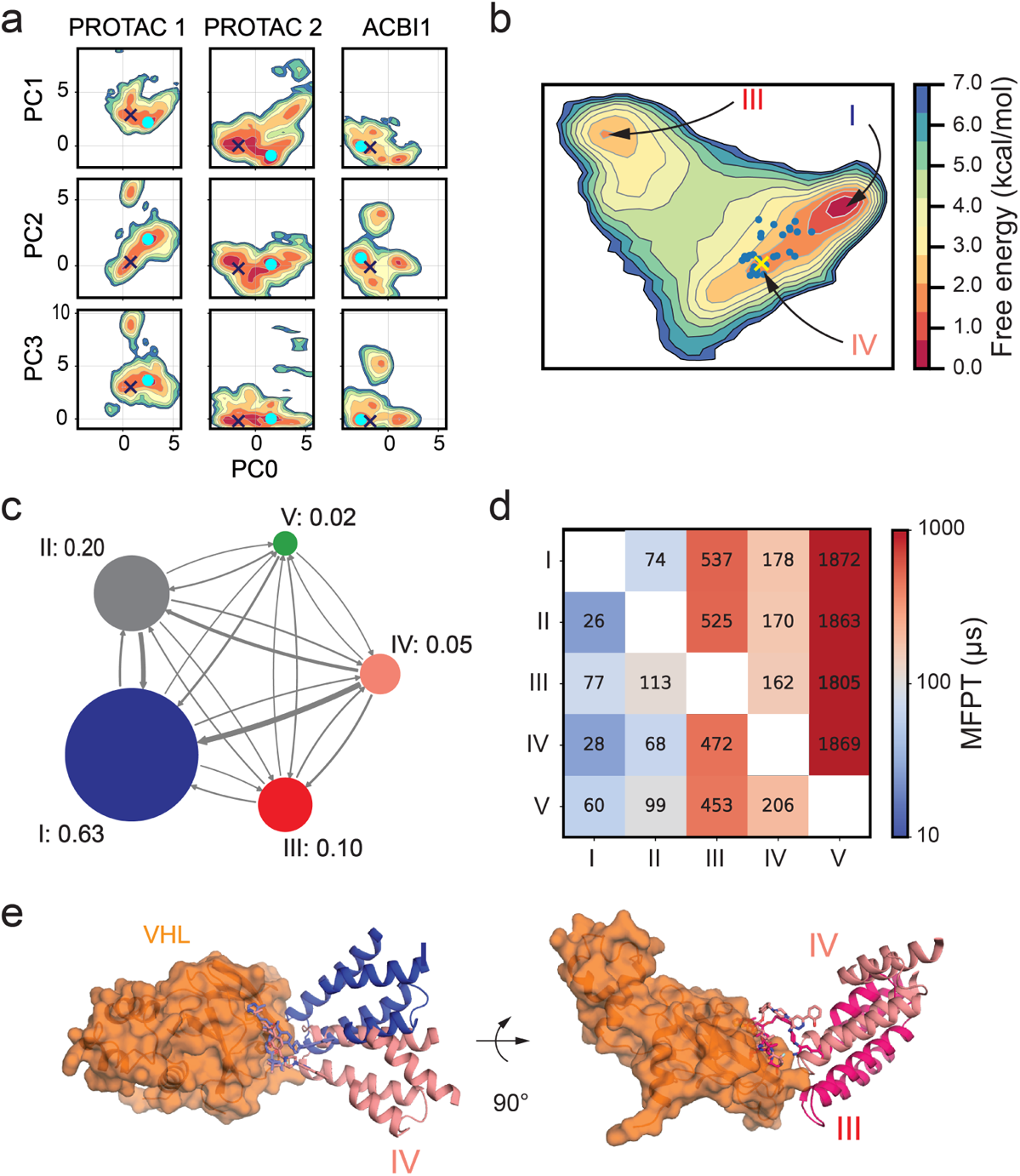
a) Conformational free energy landscapes of the iso2-SMARCA2*^BD^*:PROTAC 1:VHL, iso2-SMARCA2*^BD^*:PROTAC 2:VHL, and iso2-SMARCA2*^BD^*:ACBI1:VHL systems in the PCA space defined by our analysis of HREMD simulations. The crystal structure of each system is shown as a dark blue X, while the center of the largest *k*-means cluster is shown as a cyan point. Energy scale bar shown in panel b. b) Conformational free energy landscape as a function of the first two tICA features of iso2-SMARCA2*^BD^*:PROTAC 2:VHL ternary complex inferred from a Markov state model (MSM) determined using long time scale Folding@Home simulations. The ensemble of bound states from WE-HDX simulations is shown as blue points; the crystal structure (PDB ID: 6HAX) is shown as a yellow X. In this projection, states II and V are close to state I. c) Network diagram of the coarse-grained MSM calculated using a lag time of 50 ns, with the stationary probabilities associated with each state indicated. d) Mean first-passage times (MFPTs) to transition between MSM states. Numbers indicate predicted MFPTs in *µ*s. e) Comparison of the crystal structure (salmon) with the lowest free energy state (blue) and a metastable state (red) predicted by the MSM. Arrows indicate a change of orientation.

Our analysis shows that both iso2-SMARCA2*^BD^*:ACBI1:VHL and iso2-SMARCA2*^BD^*:PROTAC 1:VHL assume quite stable conformations: in both cases, the majority of snapshots fall into the largest cluster of conformations, Supplemental Fig. 22. The ground state (lowest free energy) structures are also quite similar to the corresponding crystal structures (*Cα*-RMSD 1.7 *±* 0.3 Å for iso2-SMARCA2*^BD^*:PROTAC 1:VHL and 0.8 *±* 0.1 Å for iso2-SMARCA2*^BD^*:ACBI1:VHL). However, iso2-SMARCA2*^BD^*:PROTAC 2:VHL shows a much more dynamic landscape, and samples conformations similar to both the ground state of iso2-SMARCA2*^BD^*:ACBI1:VHL and iso2-SMARCA2*^BD^*:PROTAC 1:VHL. This result, based on the enhanced sampling of ternary complexes, allows us to rationalize the differential in degradation efficiencies observed among the three degraders (see Table 1). We suggest that PROTAC 1 may fail to mediate the degradation of SMARCA2 because the (stable) conformation adopted by the ternary complex cannot be productively ubiquitinated. ACBI1, on the other hand, induces a productive conformation of the ternary complex, facilitating ubiquitination. Hence, PROTAC 2 would then fall between the two, as the corresponding ternary complexes sample both the productive conformation induced by ACBI1 and the non-productive PROTAC 1like conformation, Fig. 8a.

To characterize the free energy landscape of iso2-SMARCA2*^BD^*:PROTAC 2:VHL more comprehensively, we select 98 representative structures from the corresponding HREMD simulation as initial configurations for simulations on Folding@home (F@H), one of the largest distributed computing networks. Each initial condition was cloned 100 times and run for ~ 650 ns, for a total of ~ 6 ms of simulation time. These independent MD trajectories provide the basis for fitting a Markov state model (MSM),^56^ which provides a full thermodynamic and kinetic description of the system and allows for the prediction of experimental observables of interest.^57^ We use time-lagged independent component analysis (tICA)^58^ to determine the collective variables with the slowest dynamics. The distance between points in the tICA feature space corresponds roughly to a kinetic distance.^59^

The MSM uses the observed dynamics of the simulations to predict a stationary probability distribution on tICA space that is, in general, different from the empirical distribution of our simulation data. The result is shown in Fig. 8b. This model is coarsegrained to obtain a five-state MSM, of which the following three states are of particular interest: the ground state I with a stationary probability of 0.63, a metastable state III with 0.10 probability, and state IV, to which the experimental crystal structure can be assigned and which has a stationary probability of 0.05 (Fig. 8c,d).

Importantly, the MSM predicts that the ternary complex crystal structure with PROTAC 2 is 1.5 kcal/mol higher in free energy than the global free energy minimum and that they differ by an I-RMSD of 3.6 Å (Fig. 8b,e), thus lending credence to our approach of extensive conformational sampling to identify previously undetermined structures. The model further predicts a relative free energy of 2.2 kcal/mol for the metastable state with an I-RMSD of 4.4 Å relative to the crystal structure (Fig. 8b,e). Interestingly, the SMARCA2*^BD^*:PROTAC 2:VHL ternary complex structures simulated by the WE-HDX strategy described above can be well identified on this free energy landscape too (blue points on the projection in Fig. 8b), demonstrating how the simulation of ternary complexes formation yields valid conformations.

The classification into five macro-states can be attributed to structural differences at the ternary complex interface. For instance, the global minimum state is stabilized by a number of protein-protein contacts and, furthermore, contacts between PROTAC 2 and R1403, N1464, and I1470 of SMARCA2*^BD^*, that are missing in the metastable state (Supplemental Fig. 23). On the other hand, contacts between VHL and PROTAC 2 are largely unchanged between the metastable and global minimum states, likely due to the tight interaction between VHL and the degrader. The area of the binding interface is substantially increased in both the metastable and global minimum states relative to the crystal structure: the global minimum state has a buried surface area of 2962 Å^2^, compared to 2800 Å^2^ for the metastable state and only 2369 Å^2^ for the crystal structure. We note that these differences observed at the interfaces of distinct ternary complexes further support the adequacy of the minimum I-RMSD metric we used above to measure the prediction accuracy.

We also performed F@H simulations of iso2-SMARCA2*^BD^*:PROTAC 1:VHL (900 *µs* of aggregate simulation time across 1000 trajectories from 99 initial structures coming from HREMD) and iso2-SMARCA2*^BD^*:ACBI1:VHL (500 *µs* of aggregate simulation time across 2000 trajectories from 100 initial structures coming from HREMD). These simulations were used to fit MSMs for these systems using the same procedure above (Supplemental Figs. 24 and 25). The resultant MSMs predict that the crystal structure of the iso2-SMARCA2*^BD^*:PROTAC 1:VHL system is 2.2 kcal/mol higher than its global free energy minimum, while the crystal structure of the iso2-SMARCA2*^BD^*:ACBI1:VHL system is only 0.7 kcal/mol higher in energy than its ground state. Coarse-graining the PROTAC 1 model yields a two-state MSM, while a three-state MSM is obtained for the ACBI1 system. In both cases, the crystal structure falls into the most probable macro-state. Interestingly, in the predicted ground state of the ternary complex with PROTAC 1, SMARCA2*^BD^* is oriented relative to VHL (Supplemental Fig. 25) in a similar fashion as in the predicted ground state with PROTAC 2 (Supplemental Fig. 24d), while in the ground state of the ACBI1 system, the position of SMARCA2*^BD^* relative to VHL (Supplemental Fig. 25) is more similar to that in the crystal structure, which, as described above, is comparable among all three ternary complexes. This illustrates that notable conformational changes can be induced by different degrader molecules.

Interestingly, the simulations of ternary complexes of iso2-SMARCA2*^BD^* and VHL mediated by the 3 degraders confirm important interactions between charged residues at the SMARCA2*^BD^*:VHL interface that were suggested by the HDX-MS experiments presented above. In particular, R60 on VHL, which is experimentally found to be protected for a longer duration in the ternary complex, preferentially forms contacts with E1420 on the SMARCA2*^BD^* interface (see Supplemental Fig. 33) for ACBI1 and PROTAC 2 but not for PROTAC 1. ACBI1 also induces contacts between K1416 of SMARCA2*^BD^* and N90/D92 of VHL, which are notably reduced in the presence of PROTAC 1 and PROTAC 2. This consistent observation in both experiment and simulation may contribute to the stronger cooperativity observed for ACBI1 compared to PROTAC 1 and PROTAC 2.

The millisecond-long simulations presented here are, to the best of our knowledge, the most extensive sampling of ternary degrader complexes to date, permitting examination of their free energy landscapes in unprecedented detail. Remarkably, these simulations capture key structural determinants observed experimentally, such as HDX-MS residue protection and ternary complex stability and, furthermore, reveal structural differences between the energetically most favorable states of SMARCA2*^BD^*:VHL induced by different degraders that may contribute to cooperativity.

### 2.7 Large-scale simulations of the Cullin-RING ligase with VHL and SMARCA2***^BD^*** yield accurate predictions of ubiquitination

In addition to simulating the ternary complex formation and associated dynamics, a more complete understanding of the ubiquitination process should involve the full Cullin-RING E3 ubiquitin ligase (CRL). To this end, we probe the different ternary degrader complexes in the context of the full CRL macromolecular assembly by examining the separation of different solvent-exposed POI lysine residues from the ubiquitination zone of the CRL^60^ (Fig. 9a), specifically focusing on the probability of POI lysine residue density within this zone.

**Fig. 9:**
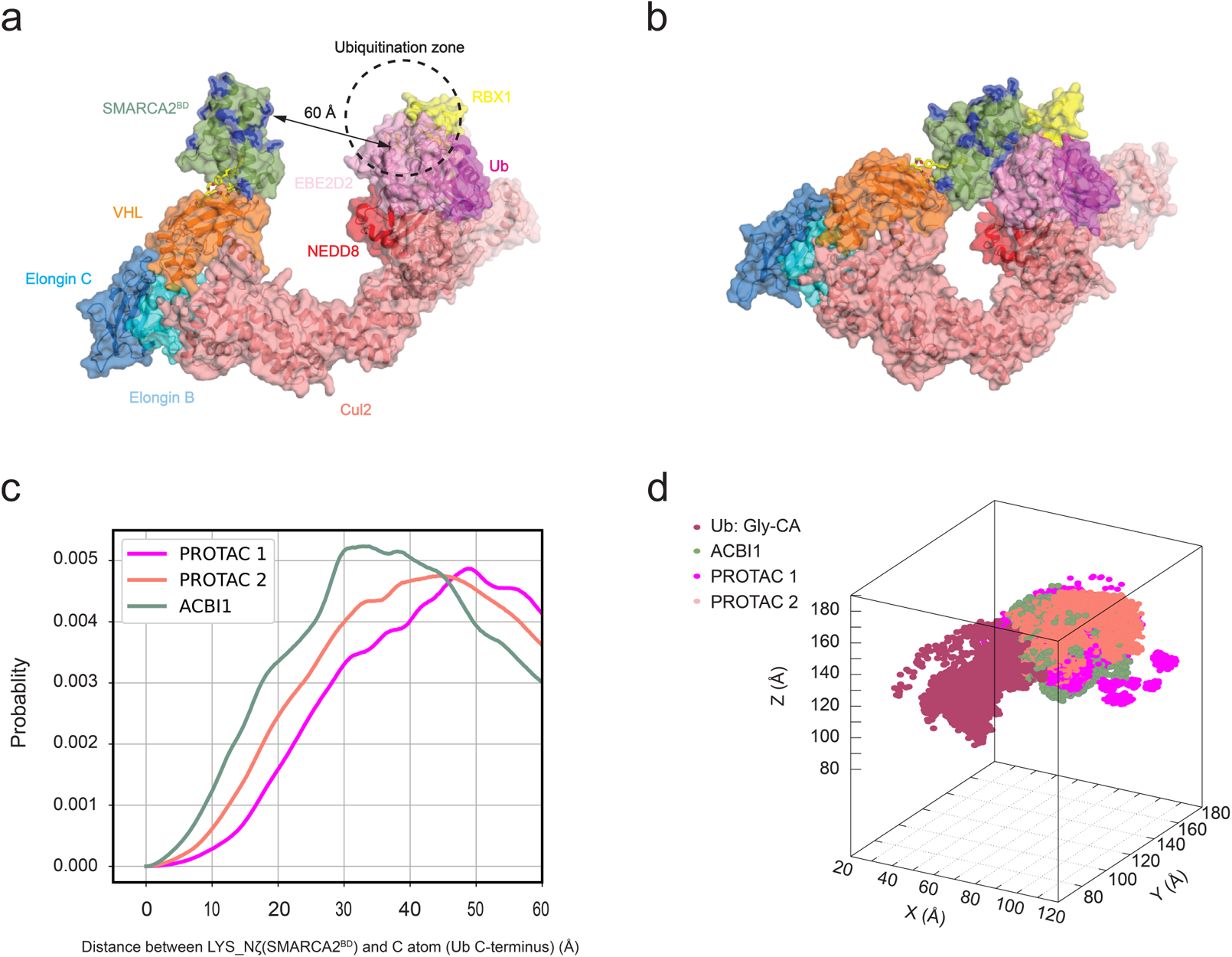
Degrader-dependent SMARCA2*^BD^* lysine densities in the CRL-VHL ubiquitination zone. a) Active form of CRL-VHL with bound SMARCA2*^BD^* and E2-ubiquitin in the open CRL conformation. b) Same as panel a with a closed conformation of CRL generated by meta-eABF simulations. c) Probability of distances of lysine residues (side-chain nitrogen atom) from SMARCA2*^BD^* to the C-terminal glycine C atom of ubiquitin for the three different degraders PROTAC 1, PROTAC 2, and ACBI1. d) Density of lysine residues in 3D space near the ubiquitination zone of CRL-VHL.

The hypothesis is that the ubiquitination rate depends on the probability of finding a lysine residue in the ubiquitination zone. As such, this analysis can provide insight into the degradation potency of degrader molecules. First, we build an entire E2-E3 complex for CRL-VHL in its activated form using a recently obtained structure of the active form of the closely related CRL-*β*TrCP as reference^61^ (see Methods 4.13). Second, we use the meta-eABF simulation approach (see Methods 4.14) to sample CRL open-closed conformations in the presence of SMARCA2*^BD^*(Fig. 9b). These conformations are then used as reference states to superimpose structures from HREMD simulations of ternary SMARCA2*^BD^*:VHL complexes on the active state of the CRL-VHL, allowing us to obtain lysine densities from SMARCA2*^BD^* in the ubiquitination zone of the CRL-VHL.

Comparing the lysine densities of the three degraders (Fig. 9c), we observe that ACBI1 places the most lysine density in the ubiquitination zone of CRL-VHL, followed by PROTAC 2 and PROTAC 1. This order of lysine density in the ubiquitination zone agrees with the experimentally observed degradation data between ACBI1 and PROTAC 1,^36^ and also places PROTAC 2 between these two, thus establishing a procedure to qualitatively predict the ubiquitination likelihood of the target protein.

To experimentally validate degrader-induced changes in global protein and ubiquitination levels, we treated Hela cells with 300 nM of ACBI1 for 1h, followed by global mass spectrometry-based proteome and ubiquitinomics analysis. In total, we quantified 13,300 ubiquitination sites on 5300 proteins (Supplemental Excel Sheet). As expected, our results confirm ACBI1-induced degradation of the SMARCA2 protein (Fig.10a). The loss of SMARCA2 protein abundance was rescued by co-treatment with 1uM proteasomal inhibitor MG132 that impedes the targeted degradation. In addition, global ubiquitination profiling identified several SMARCA2 lysine sites, some of which show a statistically significant increase in ubiquitination levels after the ACBI1 treatment, compared with the vehicle control DMSO (Fig. 10b, Table 3, Supplemental Excel Sheet). Ubiquitinated lysine residues were detected both on (e.g., K1398 and K1416) and outside the bromodomain (e.g., K1101, K1197/K1207, K1323), with the most significantly ubiquitinated residue (K1416) located on the SMARCA2*^BD^* (Fig. 10c). Not all of the lysine residues from bromodomain can be detected in this experiment due to the repeated occurrence of lysine and arginine residues within short intervals, hence the cleaved peptide is too small to be detected by the mass spectrometer. However, among those detected, the general trend is in agreement with the above-mentioned prediction that ACBI1 tends to position lysine residues closer to ubiquitin (Supplemental Fig. 34). Our results are in agreement with recent data from Arvinas and Genentech showing that a potent and selective SMARCA2 degrader is most significantly inducing ubiquitination of a lysine residue on the bromodomain of SMARCA2, although other lysine residues are ubiquitinated to lesser degrees.^62^ These results further validate the hypothesis that degraders like ACBI1 directly influence ubiquitination of lysine residues in the ubiquitination zone of CRL-VHL by modulating their global proximity to ubiquitin.

**Fig. 10:**
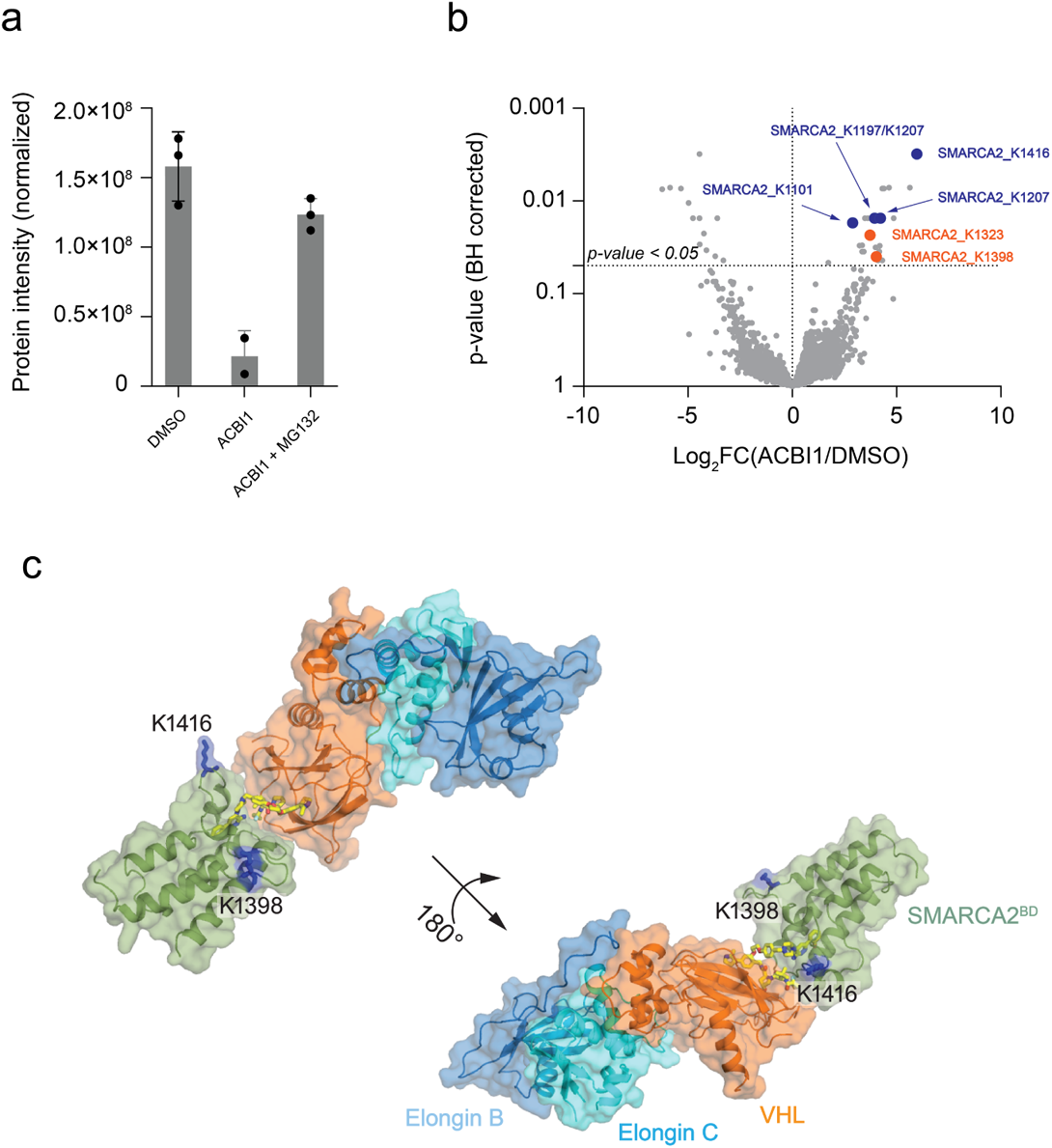
Changes in ubiquitination levels on the proteome of HeLa cells upon treatment with ACBI1 at 300 nM for 1 h. a) Change in SMARCA2 protein abundance upon treatment with DMSO, ACBI1, and ACBI1 + MG132. The ACBI1 treatment significantly decreases the SMARCA2 protein abundance compared to the DMSO alone and, upon co-treatment with the proteasomal inhibitor (MG132), the abundance is rescued to almost levels similar to the DMSO alone. b) Distribution of changes in ubiquitination levels plotted as Log_2_ fold change in ACBI1 versus DMSO control against Benjamini-Hochberg corrected P value for each ubiquitinated sites from triplicate measurements. The SMARCA2 sites with significant changes in ubiquitination levels (p-value *<* 0.05 and Log_2_ FC(ACBI1/DMSO) ≥ 1) are marked. The sites unique to SMARCA2 are marked as solid orange circles and SMARCA2/4 shared sites are shown as solid blue circles. c) Location of the two SMARCA2*^BD^* lysine residues K1398 and K1416 (shown in blue stick representation) on the SMARCA2*^BD^*:ACBI1:VCB crystal structure.

**Table 3:**
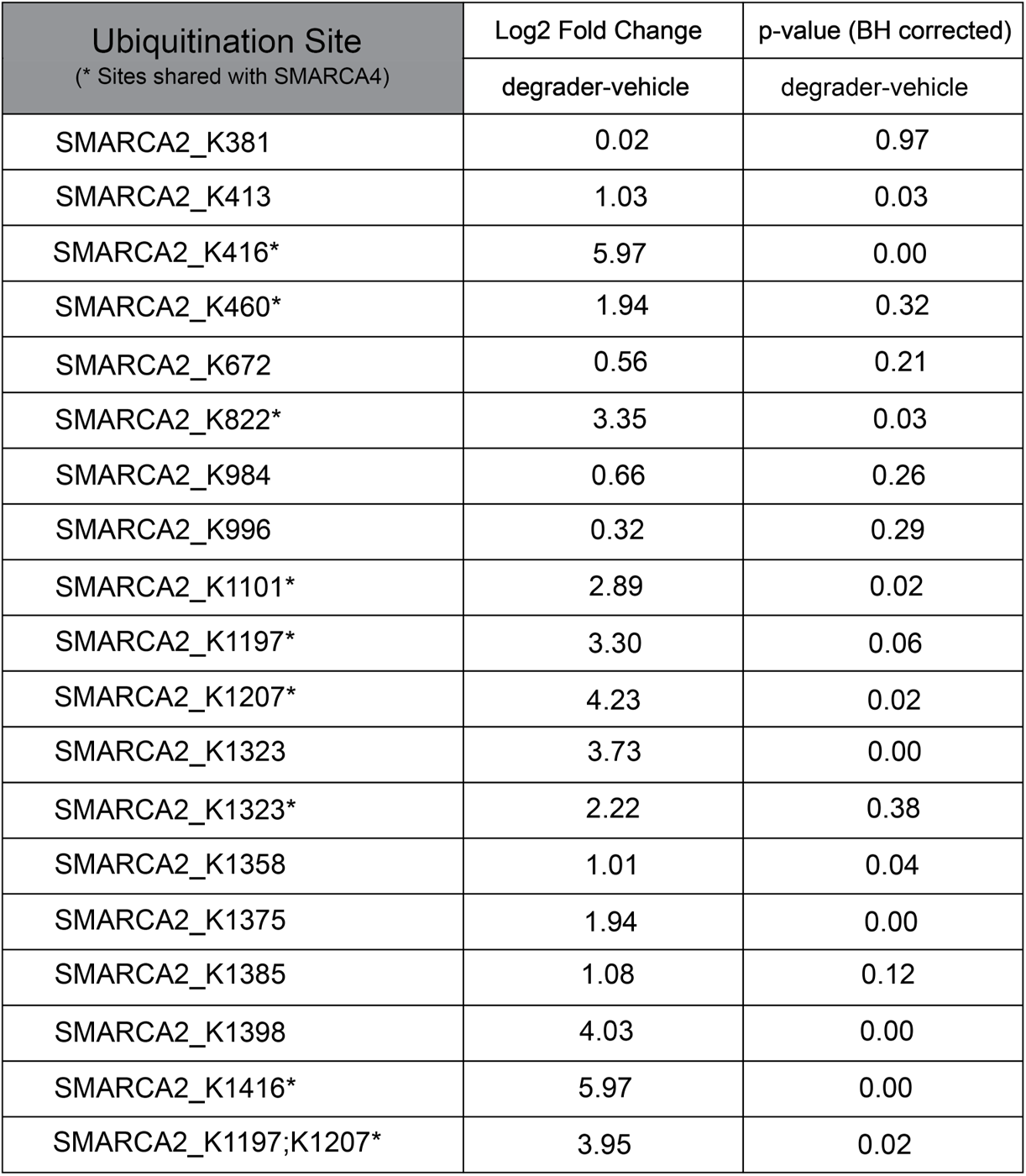
Lysine residues identified as ubiquitinated upon ACBI1 treatment. The change in abundance of ACBI1 treated ubiquitination levels compared to the DMSO treated sample are shown with associated Benjamini-Hochberg FDR corrected p-values. The residues(sites) marked with an asterisk(*) are shared sites with SMARCA4 protein.

## 3 Discussion

The formation of a ternary complex is a critical step in targeted protein degradation. However, accurately predicting the structural ensemble of the ternary complex is challenging due to the size of the multi-protein system, the inherent conformational flexibility associated with forming non-native protein-protein interactions, the relevant timescales for biological motions, and the limited data associated with the solution-phase dynamics of ternary structures. The ability to accurately predict the formation of degrader-induced ternary complexes and the corresponding structural ensembles would provide a better understanding of TPD and enable more precise optimization of degrader molecules (e.g. linker length, composition, and attachment points).

Here, we studied three different degrader molecules in complex with SMARCA2*^BD^* and VHL that have similar thermodynamic binding profiles and protein-protein interactions observed in the crystal structures but different degradation efficiencies. The crystal structure determined in this work of ACBI1 complexed with SMARCA2*^BD^* and VHL (PDB ID: 7S4E) reveals a similar conformation to previously published and close degrader analogs: PROTAC 1 (PDB ID: 6HAY) and PROTAC 2 (PDB ID: 6HAX). The similar binding thermodynamics and crystal structure complexes, yet different degradation efficiencies, motivated our work to explore the dynamic nature of the ternary structure, which might be the source of the differing degradation efficiencies (although other factors such as permeability may also play a role). The approach we describe here combines MD simulations with solution-phase biophysical experiments to produce dynamic ternary structure predictions that could be helpful in elucidating the characteristics that impact binding cooperativity and degradation efficiency.

We apply enhanced Hamiltonian replica exchange molecular dynamics (HREMD) simulations, validated by experimental SAXS data, to derive heterogeneous ensembles of ternary complex conformations that constitute the basis for millisecond-long MD simulations on Folding@home. Detailed free energy landscapes predict that the experimental crystal structures are approximately 1-2 kcal/mol higher in free energy than the lowest energy (most favorable) conformations, confirming that they are snapshots in low free energy basins, but not the global minima of those basins. Simulation global minima reveal notable differences in the orientation between SMARCA2*^BD^* and VHL induced by the three degraders.

To put the simulation results in a larger context, we examine the likelihood of ubiquitination for specific SMARCA2*^BD^*:VHL degrader ternary complexes by deploying the entire Cullin-RING E3 ubiquitin ligase (CRL). The orientation of SMARCA2*^BD^* with respect to the CRL changes dramatically in these global minima from simulation compared to the crystal structures: in particular, we find that ACBI1 positions lysines of SMARCA2*^BD^* closer on average to ubiquitin in the E2 ligase than does PROTAC 1, with PROTAC 2 shifting between an ACBI1-like position and a PROTAC 1-position, suggesting that ACBI1 has the highest propensity to facilitate ubiquitination of SMARCA2*^BD^*. We employed proteomics and ubiquitinomics experiments to determine ubiquitinated lysine residues for SMARCA2 in Hela cells. The results confirm the hypothesis that ACBI1 positions several lysine residues in closer proximity of E2-ubiquitin, enhancing ubiquitination probability, and hence, degradation efficiency. For example, we predict that K1416 in SMARCA2*^BD^* is most likely to be ubiquitinated, which is also the case in the ubiquitinomics experiment.

HDX-MS experiments revealed charged interface residues that are protected and yet are not in contact in the crystal structures. Our long-timescale ternary complex simulations revealed that several of these residues form contacts in the ternary complex ensembles. Some of these are common to all three degraders, whereas some are absent from PROTAC 1 (e.g., VHL:R60 and SMARCA2*^BD^*:E1420). These contacts may underlie the differences in cooperativity, placement in the ubiquitination zone and ultimately degradation.

We developed a novel protocol that incorporates information about protected residues as contact collective variables in weighted ensemble simulations that seek to form interactions among the protected residues and to bind the warhead portion of the degrader to SMARCA2*^BD^* (WE-HDX). This method reliably produced ternary complex structures that were in low free energy basins of the ternary complex landscape, and that were similar to the conformations accessible when starting simulations from the crystal structure. This method also provides estimates of the *k_on_* for ternary complex formation. We compared WE-HDX to docking using HDX-MS protected residues as constraints. We find that HDX constraints improve the quality of ternary complexes for docking; yet WE-HDX is more accurate than docking using HDX constraints. Further usage of the HDX-MS data could be done by computing HDX-MS observables from simulation, and then reweighting the ternary complex landscape accordingly. Even though many models are proposed in the literature, we did not estimate that those models would give us accurate reweighting at this point in time, although clearly this would be a fruitful avenue for future research.

Our integrative approach provides a richer understanding of the dynamics of ternary complex ensembles, which could improve the design of degrader molecules for new systems of interest. For the three degraders studied here, the global minima from HREMD and FAH simulations showed that the orientation of SMARCA2 lysines with respect to the E2-loaded ubiquitin is a discriminating feature, particularly of ACBI1/PROTAC 2 with respect to PROTAC 1, suggesting this to be critical for a productive ternary complex. From the conformational landscape we also find that the stability of the ternary complex differs among these 3: PROTAC 1 and ACBI1 are more stable than PROTAC 2; however PROTAC 1 is in a non-productive configuration. Thus the stability of the ternary complex indueced by ACBI1 might distinguish it from PROTAC 2. Furthermore, we consider the conformational free energy penalty for the degrader to go from its conformation in solution to ternary complex, and again we find that this penalty is larger for PROTAC 1 than it is for PROTAC 2 and ACBI1. We also found protected charged residues from HDX-MS that while not in contact in the crystal structures, appear in simulations such as VHL:R69 and SMARCA2*^BD^*:E1420, giving clue to potential structural determinants of cooperativity. And we found that ACBI1 had the highest ubiquitination probability based on our CRL modeling, followed by PROTAC 2 and PROTAC 1, which was confirmed by ubiquitinomics experiments presented here. The methodologies described here rely on advanced physics-based simulations and solution-phase biophysical experiments. Since this approach is based on physical principles without the need for training data, we expect it to be transferable to other POI-ligase ternary complexes with induced proximity degrader molecules, and possibly to other induced proximity systems (e.g. phosphorylation, methylation, and acetylation). Efforts in our group are underway to expand the application to more ligands in the SMARCA2*^BD^*:VHL system and to other POI-ligase combinations. We have used the simulation methods outlined here in a prospective manner as follows: we have predicted ternary complex ensembles of potential heterobifunctional degraders using WE-HDX; used HREMD and F@H simulations out of HREMD to select the lowest free energy structures; then calculated the ubiquitination probability of these structures by modeling them in the full CRL. We have then optimized for short and rigid linkers against the ternary complex structures selected for preferential ubiquitination, also using the conformational free energy penalty for the degrader to go from solution to the ternary complex as a design objective. Based on that we have selected the heterobifunctional molecules that optimize these properties. We expect to report on this larger data set in a future publication.

We make source code, simulation results, and experimental data from this work publicly available for researchers to further advance the field of induced proximity modulation.

## 4 Methods

### 4.1 Cloning, expression and purification of SMARCA2***^BD^*** and VHL/ElonginB/C

The SMARCA2*^BD^* gene from Homo sapiens was custom-synthesized at Genscript with N-terminal GST tag (Ciulli 2019 Nature ChemBio) and thrombin protease cleavage site. The synthetic gene comprising the SMARCA2*^BD^* (UniProt accession number P51531-1; residues 1373-1511) was cloned into pET28 vector to create plasmid pL-477. The second construct of SMARCA2*^BD^* with deletion 1400-1417 (UniProt accession number P51531-2) was created as pL-478. For biotinylated SMARCA2*^BD^*, AVI-tag was gene synthesized at C-terminus of pL-478 to create pL-479. The VHL gene from Homo sapiens was custom-synthesized with N-terminal His6 tag^36^ and thrombin protease cleavage site. The synthetic gene comprising the VHL (UniProt accession number P40337; residues 54-213) was cloned into pET28 vector to create plasmid pL-476. ElonginB and ElonginC gene from Homo sapiens was custom-synthesized with AVI-tag at C-terminus of ElonginB.^27^ The synthetic genes comprising the EloB (UniProt accession number Q15370; residues 1-104) and EloC (UniProt accession number Q15369; residues 17-112) were cloned into pCDFDuet vector to create plasmid pL-474. For protein structural study, AVI-tag was deleted in pL-474 to create pL-524.

For SMARCA2*^BD^* protein expression, the plasmid was transformed into BL21(DE3) and plated on Luria-Bertani (LB) medium containing 50 *µ*g/ml kanamycin at 37 °C overnight. A single colony of BL21(DE3)/pL-477 or BL21(DE3)/pL-478 was inoculated into a 100-ml culture of LB containing 50 *µ*g/ml kanamycin and grown overnight at 37 °C. The overnight culture was diluted to OD600=0.1 in 2 × 1-liter of Terrific Broth medium containing 50 *µ*g/ml kanamycin and grown at 37 °C with aeration to midlogarithmic phase (OD600 = 1). The culture was incubated on ice for 30 minutes and transferred to 16 °C. IPTG was then added to a final concentration in each culture of 0.3 mM. After overnight induction at 16 °C, the cells were harvested by centrifugation at 5,000 xg for 15 min at 4 °C. The frozen cell paste from 2 L of cell culture was suspended in 50 ml of Buffer A consisting of 50 mM HEPES (pH 7.5), 0.5 M NaCl, 5 mM DTT, 5% (v/v) glycerol, supplemented with 1 protease inhibitor cocktail tablet (Roche Molecular Biochemical) per 50 ml buffer. Cells were disrupted by Avestin C3 at 20,000 psi twice at 4 °C, and the crude extract was centrifuged at 39,000 xg (JA-17 rotor, Beckman-Coulter) for 30 min at 4 °C. Two ml Glutathione Sepharose 4 B (Cytiva) was added into the supernatant and mixed at 4 °C for 1 hour, washed with Buffer A and eluted with 20 mM reduced glutathione (Sigma). The protein concentration was measured by Bradford assay, and GST-tag was cleaved by thrombin (1:100) at 4 °C overnight during dialysis against 1 L of Buffer B (20 mM HEPES, pH 7.5, 150 mM NaCl, 1mM DTT). The sample was concentrated to 3 ml and applied at a flow rate of 1.0 ml/min to a 120-ml Superdex 75 (HR 16/60) (Cytiva) pre-equilibrated with Buffer B. The fractions containing SMARCA2*^BD^* were pooled and concentrated by Amicon® Ultracel-3K (Millipore). The protein concentration was determined by OD280 and characterized by SDS-PAGE analysis and analytical LC-MS. The protein was stored at –80 °C.

For VHL/ElonginB/C protein expression, the plasmids were co-transformed into BL21(DE3) and plated on Luria-Bertani (LB) medium containing 50 *µ*g/ml kanamycin and 50 *µ*g/ml streptomycin at 37 °C overnight. A single colony of BL21(DE3)/pL-476/474 or BL21(DE3)/pL-476/524 was inoculated into a 100-ml culture of LB containing 50 *µ*g/ml kanamycin and 50 *µ*g/ml streptomycin and grown overnight at 37 °C. The overnight culture was diluted to OD600=0.1 in 6 × 1-liter of Terrific Broth medium containing 50 *µ*g/ml kanamycin and 50 *µ*g/ml streptomycin and grown at 37 °C with aeration to mid-logarithmic phase (OD600 = 1). The culture was incubated on ice for 30 minutes and transferred to 18 °C. IPTG was then added to a final concentration of 0.3 mM in each culture. After overnight induction at 18 °C, the cells were harvested by centrifugation at 5,000 g for 15 min at 4 °C. The frozen cell paste from 6 L of cell culture was suspended in 150 ml of Buffer C consisting of 50 mM HEPES (pH 7.5), 0.5 M NaCl, 10 mM imidazole, 1 mM TCEP, 5% (v/v) glycerol, supplemented with 1 protease inhibitor cocktail tablet (Roche Molecular Biochemical) per 50 ml buffer. Cells were disrupted by Avestin C3 at 20,000 psi twice at 4 °C, and the crude extract was centrifuged at 17000 g (JA-17 rotor, Beckman-Coulter) for 30 min at 4 °C. Ten ml Ni Sepharose 6 FastFlow (Cytiva) was added into the supernatant and mixed at 4 °C for 1 hour, washed with Buffer C containing 25 mM imidazole and eluted with 300 mM imidazole. The protein concentration was measured by Bradford assay. For protein crystallization, His-tag was cleaved by thrombin (1:100) at 4 °C overnight during dialysis against 1 L of Buffer D (20 mM HEPES, pH 7.5, 150 mM NaCl, 1 mM DTT). The sample was concentrated to 3ml and applied at a flow rate of 1.0 ml/min to a 120-ml Superdex 75 (HR 16/60) (Cytiva) pre-equilibrated with Buffer D. The fractions containing VHL/ElonginB/C were pooled and concentrated by Amicon® Ultracel-10K (Millipore). The protein concentration was determined by OD280 and characterized by SDS-PAGE analysis and analytical LC-MS. The protein was stored at –80 °C. For SPR assay, 10 mg VHL/ElonginB/C protein complex was incubated with BirA (1:20), 1 mM ATP and 0.5 mM Biotin and 10mM MgCl2 at 4 °C overnight, removed free ATP and Biotin by 120-ml Superdex 75 (HR 16/60) with the same procedure as above, and confirmed the biotinylation by LC/MS.

### 4.2 X-ray structure determination of iso2-SMARCA2***^BD^***:ACBI1:VCB Complex

Purified SMARCA2 and VCB in 50 mM HEPES, pH 7.5, 150 mM NaCl, 1 mM DTT were incubated in a 1:1:1 molar ratio with ACBI1 for 1 hour at room temperature. Incubated complex was subsequently injected on to a Superdex 10/300 GL increase (Cytiva) pre-incubated with 50 mM HEPES, pH 7.5, 150 mM NaCl, 1 mM DTT, 2% DMSO at a rate of 0.5 mL/min to separate any noncomplexed partners from the properly formed ternary complex. Eluted fractions corresponding to the full ternary complex were gathered and spun concentrated to 14.5 mg/mL using an Amicon Ultrafree 10K NMWL Membrane Concentrator (Millipore). Crystals were grown 1-3 *µ*L hanging drops by varying the ratio of protein to mother liquor from 0.5-2:0.5-2 respectively. Crystals were obtained in buffer consisting of 0.1 M HEPES, pH 7.85, 13% PEG 3350, 0.2 M sodium formate incubated at 4° C. Crystals grew within the first 24 hours but remained at 4° C for 5 days until they were harvested, cryo protected in an equivalent buffer containing 20% glycerol and snap frozen in LN2. Diffraction data was collected at NSLS2 beamline FMX (*λ*=0.97932 Å) using an Eiger × 9M detector.

Crystals were found to be in the P 21 21 21 space group with unit cell dimensions of a= 80.14, b= 116.57, c= 122.23 Å, where *α*= *β*= *γ*=90°. Crystal contained two copies of the SMARCA2:ACBI1:VCB (VHL, ElonC, ElonB) complex within the asymmetric unit cell. The structure was solved by performing molecular replacement with CCP4i243 PHASER using PDB ID 6HAX as the replacement model. MR was followed by iterative rounds of modeling (COOT44) and refinement (REFMAC545–53) by standard methods also within the CCP4i2 suite. Structures were refined to Rwork/RF ree of 23.7%/27.5%.

### 4.3 Hydrogen Deuterium Exchange Mass Spectrometry

Our HDX analyses were performed as reported previously with minor modifications.^63–65^ With the knowledge of binding constants for each of the three degraders, the assays were designed to optimize the complex formation of 80% or greater in the D2O labeling solution after the 1:13 dilution (94% ACBI1, 93% PROTAC 2, 89% PROTAC 1) to obtain maximal exchange of the ternary complexes. Maximizing complex formation in solution ensures that the ratio of liganded to free protein in solution does not complicate the downstream analysis.^66^ HDX experiments were performed using a protein stock at the initial concentration of 200 *µ*M of SMARCA2*^BD^*, VCB in the APO, binary (200 *µ*M ACBI1) and ternary (200 *µ*M PROTAC ACBI1) states in 50 mM HEPES, pH 7.4, 150 mM NaCl, 1 mM TCEP, 2% DMSO in H2O. The protein samples were injected into the nanoACQUITY system equipped with HDX technology for UPLC separation (Waters Corp.^67^) to generate mapping experiments used to assess sequence coverage. Generated maps were used for all subsequent exchange experiments. HDX was performed by diluting the initial 200 *µ*M protein stock 13-fold with D2O (Cambridge Isotopes) containing buffer (10 mM phosphate, pD 7.4, 150 mM NaCl) and incubated at 10 °C for various time points (0.5, 5, 30 min). At the designated time point, an aliquot from the exchanging experiment was sampled and diluted 1:13 into D2O quenching buffer containing (100 mM phosphate, pH 2.1, 50 mM NaCl, 3M GuHCl) at 1 °C. The process was repeated at all time points, including for non-deuterated samples in H2O-containing buffers. Quenched samples were injected into a 5-*µ*m BEH 2.1 × 30-mm Enzymate-immobilized pepsin column (Waters Corp.) at 100 *µ*l/min in 0.1% formic acid at 10 °C and then incubated for 4.5 min for on-column digestion. Peptides were collected at 0 °C on a C18 VanGuard trap column (1.7 *µ*m × 30 mm) (Waters Corp.) for desalting with 0.1% formic acid in H2O and then subsequently separated with an in-line 1.8µMHss T3 C18 2.1 × 30-mm nanoACQUITY UPLC column (Waters Corp.) for a 10-min gradient ranging from 0.1% formic acid to acetonitrile (7 min, 5–35%; 1 min, 35–85%; 2 min hold 85% acetonitrile) at 40 *µ*l/min at 0 °C. Fragments were mass-analyzed using the Synapt G2Si ESL-Q-ToF mass spectrometer (Waters Corp.). Between injections, a pepsin-wash step was performed to minimize peptide carryover. Mass and collision-induced dissociation in data-independent acquisition mode (MSE) and ProteinLynx Global Server (PLGS) version 3.0 software (Waters Corp.) were used to identify the peptides in the non-deuterated mapping experiments and analyzed in the same fashion as HDX experiments. Mapping experiments generated from PLGS were imported into the DynamX version 3.0 (Waters Corp.) with quality thresholds of MS1 signal intensity of 5000, maximum sequence length of 25 amino acids, minimum products 2.0, minimum products per amino acid of 0.3, minimum PLGS score of 6.0. Automated results were inspected manually to ensure the corresponding m/z and isotopic distributions at various charge states were assigned to the corresponding peptides in all proteins (SMARCA2*^BD^*, VHL, ElonginC, ElonginB). DynamX was utilized to generate the relative deuterium incorporation plots and HDX heat map for each peptide (see Supplemental. Fig. 37) of each protein within the complex and stable deuterium exchange (see Supplemental Figs. 38-41). The relative deuterium uptake of common peptides was determined by subtracting the weighted-average mass of the centroid of the non-deuterated control samples from the deuterated samples at each time point. All experiments were made under the same experimental conditions negating the need for back-exchange calculations but therefore are reported as relative.^43^ All HDX experiments were performed twice, on 2 separate days, and a 98 and 95% confidence limit of uncertainty was applied to calculate the mean relative deuterium uptake of each data set. Mean relative deuterium uptake thresholds were calculated as described previously.^63–65^ Differences in deuterium uptake that exceeded the error of the datasets were considered significant.

### 4.4 SEC-SAXS experiments

SAXS data were collected with an AKTAmicro (GE Healthcare) FPLC coupled to a BioXolver L SAXS system (Xenocs) that utilized an Excillum MetalJet D2+ X-ray source operating at a wavelength of 1.34 Å. We measured two protein complex samples,

i. iso1-SMARCA2*^BD^*:ACBI1:VCB, and
ii. iso2-SMARCA2*^BD^*:ACBI1:VCB.

The scattering data was detected on PILATUS3 300 K (Dectris) detector with a resulting *q* range of 0.0134 – 0.5793 Å*^−^*^1^. To ensure the resulting scattering profile is solely due to complexes with all four protein chains and a degrader, and devoid of contributions from binary or uncomplexed proteins, size exclusion chromatography is coupled to SAXS (SEC-SAXS). The elution peak 1 of the SEC profile is assigned to the ternary complexes, whereas peak 2 is attributed to binary or uncomplexed proteins, respectively (Supplemental Fig. 14). The SEC-SAXS data for each sample was collected by loading 500 *µ*L of the ternary complex formed by addition of equimolar concentrations (275 *µ*M) of SMARCA2*^BD^*, VCB and ACBI1, onto a Superdex 200 Increase 10/30 equilibrated with 20 mM HEPES pH 7.5, 150 mM NaCl and 1 mM DTT at 20 *^◦^*C. The solution scattering data was collected as a continuous 60 second data-frame measurements with a flow rate of 0.05 mL/min. The average scattering profile of all frames within the elution peak 1 was calculated and subtracted from the average buffer scattering to yield the scattering data of the protein complex. The final SAXS profile of each ternary complex (Figure 6a) was determined from the average scattering signal from the sample in the elution peak 1, where the relatively large variability in the calculated radius of gyration, *R_g_* (red solid/open circles in Supplemental Fig. 14). This indicates that complexes are dynamic or flexible. Data reduction was performed using the BioXTAS RAW 2.0.3 software.^68^ *R_g_* was estimated from experimental an SAXS curve using the Guinier approximation,

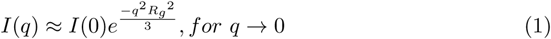

where *I*(*q*) and *I*(0) are the measured SAXS intensity and forward scattering intensity at *q*=0, respectively. *q* is the magnitude of scattering vector given by, *q* = 4*πsinθ/λ*, where 2*θ* is the scattering angle and *λ* is the wavelength of incident beam. The linear region in *ln*(*I*(*q*)) vs. *q*^2^ was fitted at low-*q* values such that *q_max_R_g_ ≤*1.3 to estimate *R_g_*, where *q_max_* is the maximum *q*-value in the Guinier fit (Supplemental Fig. 15). On the other hand, *R_g_* of the protein complex in simulation was directly calculated from atomic coordinates using following relation,

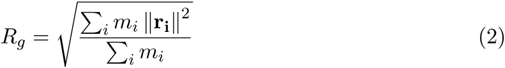

where *m_i_* is the mass of *i^th^* atom and **r_i_** is the position of *i^th^* atom with respect to the center of mass of the molecule.

### 4.5 Molecular dynamics simulations

The initial coordinates of the system were obtained from X-ray crystal structures PDB ID 6HAX, 6HAY, or 7S4E, respectively. The missing atoms were added using the LEaP module in AMBER20. The AMBER ff14SB force field^69^ was employed for the protein and the degrader force field parameters were generated using in-house programs for all MD simulations in this study. The explicit solvent was modeled using TIP3P water encapsulating the solute in a rectangular box. Counter ions were added to the system to enforce neutrality. Langevin dynamics were used to maintain the temperature at 300 K and the collision frequency was set to 2.0 ps*^−^*^1^. The SHAKE algorithm was utilized so that 2 fs time step could be achieved.

A step-wise equilibration protocol was used prior to running the production phase of the Molecular Dynamics simulations. First, a minimization was performed with a positional restraint of 5 kcal mol*^−^*^1^ Å*^−^*^2^ applied to all solute heavy atoms followed by a fully unrestrained minimization. Each minimization was composed of 500 steps of the steepest decent followed by 2000 steps of conjugate gradient. Using 5 kcal mol*^−^*^1^ Å*^−^*^2^ positional restraint on the heavy atoms of the solute, the system was linearly heated from 50 to 300 K for a duration of 500 ps (NVT ensemble) followed by a density equilibration of 750 ps (NPT ensemble). Over the course of five 250 ps simulations, the restraints on the heavy atoms of the systems were reduced from 5 to 0.1 kcal mol*^−^*^1^ Å*^−^*^2^. Then, a 500 ps simulation was run with a positional restraint of 0.1 kcal mol*^−^*^1^ Å*^−^*^2^ on the backbone atoms followed by a fully unrestrained 5 ns simulation.

Three independent regular MD simulations were performed for each of the three bound degrader complexes for up 1 *µ*s. Structures obtained from these simulations were clustered into 25 groups based on interface residue distances. One representative structure from each cluster (along with the experimentally obtained crystal structure) were used as the set of reference ternary complexes for the evaluation of bound complex predictions by WE simulations or docking.

### 4.6 Isoform 1 homology model

Since no suitable X-ray structure for iso1-SMARCA2*^BD^* is available in the PDB, we have used the YASARA (Yet Another Scientific Artificial Reality Application) homology modeling module (YASARA Biosciences GmbH) to build a high-resolution model of iso1-SMARCA2*^BD^* based on its its amino acid sequence. The sequence that was used is Uniprot P51531-1 (residues 1373-1493) has an additional 17 aa loop compared to P51531-2 (missing loop at 1400-1417). As a template for homology modeling, we used the structure from the PDB ID 6HAY. Once the model was completed, an AMBER minimization, which restrained all heavy atoms except the loop residues, was run. This ensured that the residues in the loop do not overlap and assume a stable secondary structure conformation. Minimization did not show major side-chain movements in the final minimized output which further suggested that the structure was stable

### 4.7 WE-HDX simulations

WE-HDX simulations of the formation of ternary complexes were run with both “binless” and “binned” WE variants (see Supplemental Information). These binding simulations were run with iso2-SMARCA2*^BD^* and the degrader-bound VHL subunit. The Elongin C and Elongin B subunits were omitted in these path-sampling simulations as the process of ternary complex formation is mainly determined by interactions at the SMARCA2*^BD^*:degrader:VHL interface.

Initially, the ternary complexes were unbound “manually” by separating the corresponding VHL-degrader complex from SMARCA2*^BD^* by 20 − 40°*A* (depending on the system). The (rectangular) simulation boxes of these unbound systems were then solvated with explicit water molecules and counter ions were added to neutralize their net charge. The PROTAC 1 system had 21, 191 water molecules and 10 chlorine ions. The PROTAC 2 simulations had 31, 567 water atoms and 9 chlorine ions. The ACBI1 system had 24, 093 water molecules, 9 chlorine ions. The dimensions of the simulation systems were 131*Å* × 84*Å* × 84*Å* for the PROTAC 1 system, 144*Å* × 89*Å* × 91*Å* for the PROTAC 2 system, and 123*Å* × 76*Å* × 98*Å* for the ACBI1 system.

#### 4.7.1 REVO-epsilon Weighted Ensemble method

As a bin-less WE variant, we applied the REVO algorithm. We will describe the application of the REVO algorithm as it pertains to this study, but a more detailed explanation can be found in previous works. The goal of the REVO resampling algorithm is to maximize the variation function defined as:

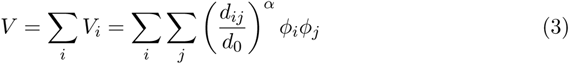

where *V_i_* is the trajectory (or walker) variation, *d_ij_* is the distance between walkers i and j determined using a specific distance metric, *d*_0_ is the characteristic distance used to make the distance term dimensionless, set to 0.148 for all simulations, the *α* is used to determine how influential the distances are to the walker variation and was set to 6 for all the simulations. The novelty terms *ϕ_i_* and *ϕ_j_* are defined as: 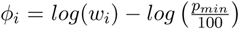. The minimum weight, *p_min_*, allowed during the simulation was 10*^−^*^50^. Cloning was attempted for the walker with the highest variance, *V_i_* when the weights of the resultant clones would be larger than *p_min_*, provided it is within distance *ɛ* of the walker with the maximal progress towards binding of the ternary complex. The two walkers selected for merging were within a distance of 2 (Å) and have a combined weight larger than the maximal weight allowed, *p_max_*, which was set to 0.1 for all REVO simulations. The merge pair also needed to minimize:

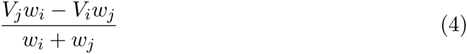

If the proposed merging and cloning operations increase the total variance of the simulation, the operations are performed and we repeat this process until the variation can no longer be increased.

Three different distance metrics were used while simulating the PROTAC 2 system: Using the warhead RMSD to the crystal structure, maximizing the contact strength (defined below) between protected residues identified by HDX data, and a linear combination of the warhead RMSD, contact strength between HDX-protected residues, and the contact strength between SMARCA2*^BD^* and the degrader. The simulations for the other systems used the last distance metric exclusively. To compute the warhead RMSD distance metric, we aligned to the binding site atoms on SMARCA2*^BD^*, defined as atoms that were within 8 Å of the warhead in the crystal structure. Then the RMSD was calculated between the warhead in each frame and the crystal structure. The distance between a set of walkers i and j is defined as: 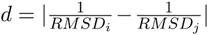. The contact strength is defined by determining the distances between residues. We calculate the minimum distance between the residues and use the following to determine the contact strength:

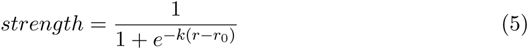

where k is the steepness of the curve, *r* is the minimum distance between any 2 residues and *r*_0_ is the distance we want a contact strength of 0.5. We used 10 for k and 5 Å for *r*_0_. The total contact strength was the sum of all residue-residue contact strengths. The distance between walkers i and j was calculated by: *d* = *|cs_i_ − cs_j_|* where cs is the contact strength of a given walker.

All REVO simulations were run using OpenMM v.7.5.0. Simulation details are as described above. The degrader-VHL interface was restrained to maintain the complex during the simulation by using a OpenMM custom centroid force defined as:

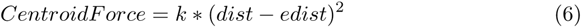

where the dist is the distance between the center of mass of the degrader and the center of mass of VHL and the edist is the distance between the center of mass of the degrader and center of mass of VHL of the crystal structure, and k is a constant set to 2 kcal/mol *∗ Å*^2^.

#### 4.7.2 Binned Weighted Ensemble method

We also applied a variant of the WE simulation, in which the pre-defined collective variable is divided into bins, using the WESTPA software.^70,71^ Each bin may contain a number (*M*) of walkers, *i*, that carry a certain weight (*w_i_*). The simulations were run for a relatively short time (*τ* = 50*ps*), after which walkers are either replicated, if their number per bin is *< M*, or they are merged, if there are *> M* walkers per bin. Similar to REVO, the sum of all *w_i_* equals 1 in any iteration, i.e., the trajectory replication and merging operations correspond to an unbiased statistical resampling of the underlying distribution.^72^ Detailed description about the WE path sampling algorithms can be found elsewhere.^45,46^

The unbound systems described above were taken as the starting configuration for each binding simulation with the GPU-accelerated version of the AMBER molecular dynamics package.^73^ To ensure the degrader remains bound to the VHL protein during these simulations, a modest (1 kcal mol*^−^*^1^ Å*^−^*^2^) flat-bottom position restraint was enforced between the center of masses of the ligand and protein binding site heavy atoms. All other MD simulation parameters were as described above *M* was set to 5 and two collective variables (CV1 and CV2) were defined to assess progress during the simulations or ternary complexes with each of the three degraders. CV1 was defined as the warhead-RMSD, or w-RMSD, of the degrader warhead with respect to the corresponding crystal structure of the bound complex. CV2 was a combination of two observables; it was either defined to be the number of native atom contacts between the warhead and the SMARCA2*^BD^* binding interface or, if the binding sites were so distant that no contacts were formed, it was defined as the distance of the binding partners, i.e., SMARCA2*^BD^* and the VHL-degrader binary complex. Contacts were counted between non-hydrogen atoms within a radius of 4.5 Å and, to ensure that CV2 is defined along one linear dimension, the contact counts were scaled by −1. This selection of CV1 and CV2 with an appropriate binning allowed the separated binding partners to assemble, during the WE simulations, into ternary complexes that are similar to the corresponding crystal structures, which were used for w-RMSD and native contact calculations.

When augmenting the WE simulations with HDX-MS data, i.e., in the WE-HDX simulations, only the protected residues of the two proteins, as informed by the corresponding experiments, were taken into consideration for the contact counts of CV2.

The ensemble of predicted bound structures was evaluated by comparing the distributions of minimum interface-RMSDs (I-RMSDs) with respect to the aforementioned set of reference ternary complexes, where the interface is defined by SMARCA2*^BD^* and VHL residues within 10 Å. Furthermore, to obtain a subset of reliable predictions, these I-RMSD distributions only contain structures with w-RMSD *<* 2 Å and *>* 30 contacts between any residues of the two proteins or, in the case of WE-HDX simulations, between the protected residues of the two proteins.

### 4.8 Ternary complex docking protocol

Following the previously reported applications of molecular docking to predictions of ternary complexes (i.e., Methods 4 and 4b from Drummond et al.^32,34^ as well as the approach from Bai et al.^31^), we assume that high fidelity structures of SMARCA2*BD*:warhead and VHL:ligand are known and available to be used in protein-protein docking. This docking of two proteins with bound degrader moieties is performed in the absence of the linker. The conformations of the linker are sampled independently with an inhouse developed protocol that uses implementation of fast quantum mechanical methods, CREST.^74–76^ Differently from the docking protocols described in,^31,32,34^ we make use of distance restraints derived either from the end-to-end distances of the sampled conformations of linker, or from the HDX-MS data. Thus, before running the protein-protein docking, we generate an ensemble of conformers for linkers and calculate the mean (*x*_0_) and standard deviation (*sd*) for the end-to-end distance. This information is then used to set the distance restraints in the RosettaDock software:^77,78^

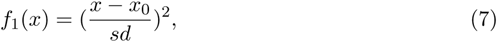

where *x* is the distance between a pair of atoms in a candidate docking pose (the pair of atoms is specified as the attachment points of the linker to warhead and ligand).

When information about the protected residues is available from HDX-MS experiments, we used them to set up a set of additional distance restraints:

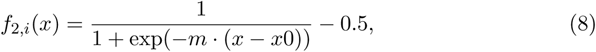

where *i* is the index of a protected residue, *x*0 is the center of the sigmoid function and *m* is its slope. As above, *x*0 value was set to be the mean end-to-end distance calculated over the ensemble of linker conformers. The value of *m* was set to be 2.0 in all the performed docking experiments. The type of RosettaDock-restraint is *SiteConstraint*, with specification of C*α* atom for each protected residue and the chain-ID of partnering protein (i.e., *x* in Eq.(8) is the distance of C*α* atom from the partnering protein). Thus, the total restraint-term used in docking takes the form:

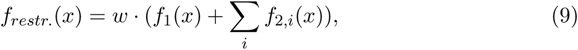

where *w* = 10 is the weight of this additional score function term.

RosettaDock implements a Monte Carlo-based multi-scale docking algorithm that samples both rigid-body orientation and side-chain conformations. The distance-based scoring terms, Eq. (9), bias sampling towards those docking poses that are compatible with specified restraints. This limits the number of output docking structures, as only those ones that pass the Metropolis criterion with the additional term of Eq. (9) will be considered.

Once the docking poses are generated with RosettaDock, all the pre-generated conformations of the linker are structurally aligned onto each of the docking predictions.^31^ Only those structures that satisfy the RMS-threshold value of *≤* 0.3 Å are saved as PDB files. All the docking predictions are re-ranked by the values of Rosetta Interface score (*I_sc_*). The produced ternary structures are examined for clashes, minimized and submitted for further investigations with Molecular Dynamics methods. Details about running the described docking protocol can be found in the Supplemental Information.

### 4.9 HREMD simulation

The simulation box of a ternary complex was solvated with explicit water and counter ions were added to neutralize the net charge of the system. We chose the Amber ff14SB force field^79^ for protein and TIP3P water model.^80^ For the degrader molecules, force field parameters were generated using in-house force field generator. The LINCS algorithm^81^ was used to constrain all bonds including hydrogen atoms. The equation of motions was numerically integrated with a time step of 2 fs using the Verlet leapfrog algorithm.^82^ The particle-mesh Ewald summation^83^ with a fourth-order interpolation and a grid spacing of 1.6 Å was employed to calculate the long-range electrostatic interactions. A cutoff of 12 Å was imposed for the short-range electrostatic and Lennard-Jones interactions. The solute and solvent were coupled separately to a temperature bath of 300 K using the Velocity-rescale thermostat^84^ with a relaxation time of 0.1 ps. The Parrinello-Rahman algorithm^85^ with a relaxation time of 2 ps and isothermal compressibility of 4.5*×*10*^−^*^5^ bar*^−^*^1^ was utilized for a pressure coupling fixed at 1 bar. We started with minimizing the energy of a system using the steepest descent algorithm. Then, the system was equilibrated at the NVT and NPT ensembles for 1 ns each. Finally, we ran the production runs in the NPT ensemble.

The details of Hamiltonian replica-exchange MD (HREMD) can be found in the Supplemental Information (Supplemental Figs. 16, 17 and Supplemental Table ??). For all HREMD simulations, we chose the effective temperatures, *T*_0_ = 300 K and *T_max_* = 425 K such that the Hamiltonian scaling parameter, *λ*_0_ = 1.00 and *λ_min_* = 0.71 for the lowest and the highest rank replicas, respectively. We estimated the number of replicas (*n*) in such a way that the average exchange probabilities (*p*) between neighboring replicas were in the range of 0.3 to 0.4. We used *n*=20 and *n*=24 for SMARCA2*^BD^*:degrader:VHL and SMARCA2*^BD^*:degrader:VCB respectively. Each simulation was run for 0.5 *µ*s/replica, and a snapshot of a complex was saved every 5 ps (total 100,001 frames per replica). Finally, we performed all the analyses on only the lowest rank replica that ran with original/unscaled Hamiltonian.

We assessed the efficiency of sampling by observing (i) the values of *p*, (ii) a good overlap of histograms of potential energy between adjacent replicas (Supplemental Fig. 16), and (iii) a mixing of exchange of coordinates across all the replicas (Supplemental Fig. 17). Furthermore, to show the convergence of HREMD simulation, we computed three metrics, radius of gyration of a ternary complex, center of mass (COM) distance between SMARCA2*^BD^* and VHL, and heavy atom contacts within 5 Å between SMARCA2*^BD^* and VHL. The distributions of these metrics are plotted with cumulative length of HREMD simulation (Supplemental Figs. 18 and 19). We noted that the distributions are similar for the last 0.3 *µ*s (0 - 0.3 *µ*s, 0 - 0.4 *µ*s and 0 - 0.5 *µ*s) of the lowest rank replica implying the convergence of the simulation.

### 4.10 MD simulation of degraders

PROTAC 1, PROTAC 2 and ACBI1 were solvated in a simulation box with 1002, 1207 and 3169 TIP3P water^80^ molecules respectively, along with counter ions to neutralize the system. All other simulation parameters were same as described in section 4.9. The production MD simulation of each degrader was run in the NPT ensemble for 1 *µ*s.

### 4.11 Conformational free energy landscape determination

In order to quantify to the conformational free energy landscape, we performed dimension reduction of our simulation trajectories using principle component analysis (PCA). First, the simulation trajectories were featurized by calculating interfacial residue contact distances. Pairs of residues were identified as part of the interface if they passed within 7 Å of each other during the simulation trajectory, where the distance between two residues was defined as the distance between their C*α* atoms. PCA was then used to identify the features that contributed most to the variance by diagonalizing the covariance matrix of the iso2-SMARCA2:PROTAC 2:VHL system; four PCA features were used in our analysis, chosen because this many features were needed to explain greater than 95% of the variance.

After projecting the simulation data onto the resultant feature space, snapshots were clustered using the *k*-means algorithm. The number of clusters *k* was chosen using the “elbow-method”, i.e., by visually identifying the point at which the marginal effect of an additional cluster was significantly reduced. In cases where no “elbow” could be unambiguously identified, *k* was chosen to be the number of local maxima of the probability distribution in the PCA feature space. The centroids determined by *k*-means approximately coincided with such local maxima, consistent with the interpretation of the centroids as local minima in the free energy landscape, see Supplemental Fig. 22.

To prepare the Folding@home simulations, HREMD data were featurized with interface distances and its dimensionality reduced with PCA as described above. The trajectory was then clustered into 98 k-means states for PROTAC 2, and 100 states for PROTAC 1 and ACBI1, whose cluster centers were selected as ‘seeds’ for Folding@home massively parallel simulations. The simulation systems and parameters were kept the same as for HREMD and loaded into OpenMM where they were energy minimized and equilibrated for 5 ns in the NPT ensemble (T = 310 K, p = 1 atm) using the openmmtools Langevin BAOAB integrator with 2 fs timestep. 100 trajectories with random starting velocities were then initialized on Folding@home for each of the seeds. The final dataset consists of 9800 trajectories, 5.7 milliseconds of aggregate simulation time, and 650 ns median trajectory length. This dataset is made publicly available at: https://console.cloud.google.com/storage/browser/paperdata.

For computational efficiency, the data was strided to 5 ns/frame, featurized with closest heavy atom interface distances, and projected into tICA space at lag time 5 ns using commute mapping. The dimensionality of the dataset was chosen to keep the number of tICs necessary to explain 95% of kinetic variance: 219 for PROTAC 1, 339 for PROTAC 2, and 197 for ACBI1. The resulting tICA space was discretized into microstates using *k*-means: we used 30 microstates for PROTAC 1, 1000 microstates for PROTAC 2, and 40 microstates for ACBI1. The Markov state models (MSM) were then estimated from the resulting discretized trajectories at lag time 50 ns. For the PROTAC 2 MSM, we used a minimum number of counts for ergodic trimming (i.e. the ‘mincount connectivity’ argument in PyEMMA) of 4, as the default setting resulted in a trapped state whose connectivity between simulation sub-ensembles starting from two different seeds was observed only due to clustering noise. The validity of the MSM was confirmed by plotting the populations from raw MD counts vs. equilibrium populations from the MSM, which is a useful test, especially when multiple seeds are used and the issue of connectivity is paramount. A hidden Markov model (HMM) was then computed to coarse-grain the transition matrix using 2 macrostates for PROTAC 1, 5 macrostates for PROTAC2, and 3 macrostates for ACBI1. Chapman-Kolmogorov tests using the transition matrices from these HMMs are shown in Supplemental Figures 31, 29, and 30. A better alternative to build macrostate models might be to construct memory kernels^86^ rather than fuzzy assignments of states as in HMMs. This may also reduce the computational resources needed to estimate free energies and transition kinetics of macrostates.

During analysis of our PROTAC 1 simulations, we found that one of our initial 100 seeded structures was kinetically separated from the others, such that reversible transitions to this state were not observed in our F@H trajectories. Transitions between this state and the ground state were therefore identified as the slowest mode by tICA. Since transitions *to* this state were never observed in our F@H simulations, we simply removed all trajectories seeded from this initial structure and omitted the first tIC from our analysis.

### 4.12 Comparison of HREMD to SAXS experiment

We validated the HREMD-generated ensembles of iso1/iso2-SMARCA2:ACBI1:VCB complexes by directly comparing to the experimental SAXS data. The theoretical SAXS profile was computed from each snapshot from the HREMD simulation trajectory using CRYSOL^87^ available in a software package ATSAS.^88^ The following CRYSOL command was used: *crysol < filename.pdb > −lm* 20 *−sm* 0.5 *−ns* 201 *−un* 1 *−eh −dro* 0.03. To expedite the writing of PDBs from HREMD trajectory and calculation of SAXS profiles, we used the multiprocessing functionality implemented in a Python package *idpflex*.^89^ The ensemble-averaged theoretical SAXS profile was determined as below,

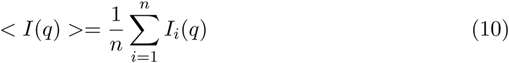

where *n* = 100,001 is the total number of frames in HREMD trajectory of each complex. The ensemble-averaged theoretical SAXS profile was compared to experiment (Fig. 6c) by minimizing chi-square (*χ*^2^) given by,

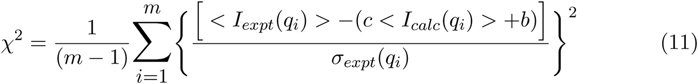

where *< I_expt_*(*q*) *>* and *< I_calc_*(*q*) *>* are the ensemble-averaged experimental and theoretical SAXS intensities respectively, *m* is the number of experimental *q* points, *c* is a scaling factor, *b* is a constant background, and *σ_expt_* is the error in *I_expt_*(*q*).

### 4.13 Cullin-RING E3 ubiquitin ligase (CRL) simulations to explore activation

To study the impact of different degraders on ubiquitination, first we constructed an active form of the Cullin-RING E3 ubiquitin ligase (CRL) with VHL and grafted it onto the ternary structures from the SMARCA2*^BD^*:degrader:VHL simulations described above. We used targeted MD simulations (TMD)^90^ to drive the activation of the CLR based on the active structure of a homologous E3 ligase, CRL-*β*TrCP (PDB ID: 6TTU).^61^ The full CRL-VHL system was built using PDB IDs 1LQB and 5N4W including VHL, ElonginB, ElonginC, Cullin2, and RBX1.^11,91^ NEDD8 was placed near residue Lys689 of the CRL where neddylation occurs.

As the collective variable for TMD, we used the residue-based RMSD of the last ~70 C*α* atoms of the Cullin C-terminus (where neddylation and subsequent activation occur) of Cullin1 from the 6TTU structure^61^ as the reference state and modeled Cullin2 from its inactive form in the 5N4W structure to this reference state. In addition, the C*α* atoms of the entire NEDD8 protein from the 6TTU structure was also used as a reference structure during TMD. Residues 135 to 425 from Cullin2 and corresponding residues from Cullin1 were used for alignment during TMD. The force constant for TMD was set to 30 kJ/mol/nm^2^. The system in a rectangular simulation box with a total number of ~500K atoms and an ionic concentration of 0.120 M using KCl. Hydrogen mass repartitioning (HMR) was used to enable 4 fs timestep simulations using the the AMBER ff14SB force field parameters. The TMD structure was then used to build the entire complex for CRL-VHL-Degrader-SMARCA2*^BD^*. The system also included E2 and ubiquitin from the 6TTU structure. This system was solvated in a truncated octahedral box to avoid protein rotation during simulation and it was equilibrated for about 30 ns before subsequent meta-eABF simulations for identifying the ubiquitination zone.

### 4.14 Meta-eABF simulations on full Cullin-RING E3 ubiquitin ligases (CRL) complex

We employ an advanced path-based simulation method that combines metadynamics with extended adaptive biasing force (meta-eABF) to study the dynamic nature of the full CRL-VHL-degrader-SMARCA2*^BD^* complex and generate a diverse set of putative closed conformations that place the E2-loaded ubiquitin close to lysine residues on SMARCA2*^BD^*. The results from the meta-eABF simulation are used to seed additional simulations for unbiased ensemble-scale sampling.

Detailed description of the meta-eABF algorithm and its variants can be found elsewhere,^92–95^ but for clarity we present a brief account here. Similar to adaptive biasing force (ABF) methods, meta-eABF simulations also utilize adaptive free energy biasing force to enhance sampling along one or more collective variables (CVs), but the practical implementation is different. Meta-eABF evokes the extended Lagrangian formalism of ABF whereby an auxiliary simulation is introduced with a small number of degrees of freedom equal to the number of CVs, and each real CV is associated with its so-called fictitious counterpart in the low-dimensional auxiliary simulation. The real CV is tethered to its fictitious CV via a stiff spring with a large force constant and the adaptive biasing force is equal to the running average of the negative of the spring force. The biasing force is only applied to the fictitious CV, which in turn “drags” the real simulation along the real CV via the spring by periodically injecting the instantaneous spring force back into the real simulation. Moreover, the main tenet of the meta-eABF method is employing metadynamics (MtD) or well-tempered metadynamics (WTM) to enhance sampling of the fictitious CV itself. The combined approach provides advantages from both MtD/WTM and eABF.

For CRL-VHL closure we chose a single CV, the center-of-mass (COM) distance between SMARCA2*^BD^* and E2 ligase-ubiquitin (E2-Ub) complex. The initial COM distance after relaxation was ~65 Å, and we ran 40 ns of meta-eABF simulation biasing the COM distance between 25-75 Å. During this simulation we saw multiple ring closing-opening events with the last frame representing a slightly open conformation with COM distance ~36 Å. We then continued the meta-eABF simulation for another 80 ns but narrowing the bias range on the COM distance to 25-40 Å in order to focus the sampling on closed or nearly closed conformations. The simulations were run using OpenMM 7.5^96^ interfaced with PLUMED 2.7.^97^

### 4.15 Mass spectrometry-based proteomics and ubiquitin analysis

Hela cells were cultured at a seeding density of 6E6 cells per 150 cm dish the day before in IMDM + 10% FCS. Next day, the cells were treated for 1 h with either i) 300 nM of ACBI1, ii) 300 nM of ACBI1 + 10 µM MG132 or, iii) vehicle (DMSO) alone. Three plates of cells were treated for triplicate measurement in each condition. The cell pellets were collected after 1 h and lysed in 50 mM TEAB (pH 7.5) buffer containing 5% (w/v) SDS. Protein amounts were quantified using a BCA assay (Thermo Fisher Scientific) according to manufacturers’ instructions. A total of 5 mg of each sample was processed and digested overnight using the S-trap (midi S-trap)-based approach according to a published protocol.^98^ Enrichment of ubiquitinated peptides (GG-remnants) was performed using an anti-diGly remnant antibody (CST, PTMScan® Ubiquitin Remnant Motif (K-*ɛ*-GG) Kit) following a previously reported protocol.^99^ We used 10 µL of slurry beads (corresponding to 62.5 µg antibody) for each ubiquitin pull-down. Each sample was desalted and separated into four fractions using basic reversed-phase tip columns as previously described.^100^ Fractions were dried down and stored at −20°C until further analysis.

### LC–MS/MS analysis

Peptides were dissolved in 0.1% formic acid (FA) and analyzed on a Q-Exactive Plus mass spectrometer (Thermo Scientific) coupled to an Ultimate 3000 RSLCnano ultra HPLC system (Thermo Scientific). The samples were separated in a 120 min gradient (from 4% solvent B to 32% solvent B over 100 min; Solvent A 0.1% FA, 5% DMSO in water; solvent B 0.% FA, 5% DMSO in acetonitrile. The loading buffer was 0.1% formic acid in water. The mass spectrometer was operated in a data-dependent acquisition (DDA) mode with an MS1 scan from 360–1300 m/z, acquired at 70,000 resolution. The MS1 scan was followed by 20 m/z dependent MS2 scans. The precursor ions were fragmented by higher energy collision dissociation (HCD) and acquired at a resolution of 17,500. The automatic gain control (AGC) targets for MS1 and MS2 were set at 3 × 10E6 ions and 1 × 10E5 ions, respectively. The maximum ion injection time for MS1 was set to 25 ms for MS1 and 50 ms for MS2 acquisition, with a dynamic exclusion of 35 sec. The normalized collision energy was set at 28%. Peptide and protein identification. MaxQuant software v.2.0.1.0 was used for protein identification and label-free quantification (LFQ). The raw mass spectrometry data files were searched against the Human UniProt database using trypsin as the digestion enzyme with up to two missed cleavages allowed. Carbamidomethylated cysteine was set as a static modification. Oxidation of methionine, protein N-terminal acetylation, and GlyGly on lysine were set as variable modifications. The match-between-run option in MaxQuant was switched on. To control for false positives, a 1% false discovery rate was used on the PSM and the protein level.

### Data analysis

For the ubiquitination profiling data, the distribution of ubiquitination sites/protein intensity ratios between each sample and the vehicle samples were computed. A constant scaling factor per sample was determined so that the median of this distribution becomes 1, assuming the intensities of most ubi-site proteins do not change. All intensities in the samples were multiplied with this scaling factor for normalization. Statistical testing was performed using the limma R package^101^ on normalized log2-transformed intensities. Missing values were imputed as long as no replicate of the same treatment had an intensity larger than the median intensity of the treatment. Proteome-corrected ubiquitination values were calculated using imputed intensities. Statistical significance was determined using pairwise t-test with Benjamini-Hochberg correction for multiple testing.

### Data availability

The mass spectrometry proteomics data have been deposited to the ProteomeXchange Consortium via the PRIDE partner repository^102^ with the dataset identifier PXD033763.

## Supporting information

Supplementary Ubiquitination Sites Proteomics

## Acknowledgement

This research used resources of the Oak Ridge Leadership Computing Facility at the Oak Ridge National Laboratory, which is supported by the Office of Science of the U.S. Department of Energy under Contract No. DE-AC05-00OR22725.

We thank the University of Massachusetts Institute of Applied Life Sciences Mass Spectrometry Core (RRID:SCR 019063) and Stephen J. Eyles for their support and mentorship during the collection, and processing of all Hydrogen Deuterium Exchange Data. We thank Helix Biostructures LLC for their assistance with X-Ray data collection and raw data reduction. SAXS measurements were based upon research conducted at the Structural Biology Platform of the Université de Montréal, which is supported by the Canadian Foundation for Innovation award #30574.We would also like to thank OmicScouts GmbH for their support on Proteomics data generation.

We are grateful to all the citizen scientists who contributed their compute power to make parts of this work possible, and members of the Folding@home community who volunteered to help with technical support to run these simulations.

## 5 Author contributions

TDi and BM ran and analyzed WE-HDX simulations. DMa performed crystallography and HDX-MS experiments. TDa ran docking simulations. SL wrote software to support simulations and analysis. DMc ran MD simulations and analyses. SSh and RP performed homology modeling and analyzed data. URS ran HREMD simulations and compared to SAXS. RW and ZAM ran FAH simulations and performed conformational landscape analyses. FP analyzed HDX data. JVR assisted with visualization and analyses. TW and VS helped scale WESTPA on Summit and run all simulations efficiently in our HPC cluster. NG and SJ performed protein production. SSp performed SAXS analyses. YL and AV performed SPR experiments. XZ oversaw synthesis of degrader molecules. AMR and IK performed CRL simulations and ubiquitination analyses. JA and BR designed and analyzed proteomics experiments JIm, AE, and LB helped edit the paper. AD, HX, WS and JAI directed the research presented in this paper. All authors wrote the paper.

## Supporting Information Available

We make all experimental data used in this study available, including HDX-MS and a crystal structure of SMARCA2*^BD^*:ACBI1:VHL-Elongin C-Elongin B (PDB ID: 7S4E). We also make available trajectory data for the conformational sampling of the crystal structures and the ternary complex formation simulations at https://console.cloud.google.com/storage/browser/paperdata.

We have created a repository information about the format of the WE-HDX trajectory data, and source code needed to run WE-HDX at https://github.com/stxinsite/degrader-ternary-complex-prediction.

## References

(1) Wu, T.; Yoon, H.; Xiong, Y.; Dixon-Clarke, S. E.; Nowak, R. P.; Fischer, E. S. Targeted protein degradation as a powerful research tool in basic biology and drug target discovery. NAT STRUCT MOL BIOL 2020, 27, 605–614.

(2) Schneider, M.; Radoux, C. J.; Hercules, A.; Ochoa, D.; Dunham, I.; Zalmas, L.-P.; Hessler, G.; Ruf, S.; Shanmugasundaram, V.; Hann, M. M.; Thomas, P. J.; Queisser, M. A.; Benowitz, A. B.; Brown, K.; Leach, A. R. The PROTACtable genome. NAT REV DRUG DISCOV 2021, 1–9.

(3) Schapira, M.; Calabrese, M. F.; Bullock, A. N.; Crews, C. M. Targeted protein degradation: expanding the toolbox. NAT REV DRUG DISCOV 2019, 18, 949– 963.

(4) Coleman, K. G.; Crews, C. M. Proteolysis–Targeting Chimeras: Harnessing the Ubiquitin–Proteasome System to Induce Degradation of Specific Target Proteins. Annual Review of Cancer Biology 2017, 2, 1–18.

(5) Matyskiela, M. E. et al. A Cereblon Modulator (CC-220) with Improved Degradation of Ikaros and Aiolos. Journal of Medicinal Chemistry 2018, 61, 535–542.

(6) Chamberlain, P. P. et al. Structure of the human Cereblon–DDB1–lenalidomide complex reveals basis for responsiveness to thalidomide analogs. Nature Structural & Molecular Biology 2014, 21, 803–809.

(7) Krönke, J. et al. Lenalidomide Causes Selective Degradation of IKZF1 and IKZF3 in Multiple Myeloma Cells. Science 343, 301–305.

(8) Ohoka, N. et al. In Vivo Knockdown of Pathogenic Proteins via Specific and Nongenetic Inhibitor of Apoptosis Protein (IAP)-dependent Protein Erasers (SNIPERs)*. Journal of Biological Chemistry 2017, 292, 4556–4570.

(9) Wei, J. et al. Harnessing the E3 Ligase KEAP1 for Targeted Protein Degradation. Journal of the American Chemical Society 2021, 143, 15073–15083.

(10) Rodriguez-Gonzalez, A.; Cyrus, K.; Salcius, M.; Kim, K.; Crews, C. M.; Deshaies, R. J.; Sakamoto, K. M. Targeting steroid hormone receptors for ubiquitination and degradation in breast and prostate cancer. Oncogene 2008, 27, 7201–7211.

(11) Hon, W.-C.; Wilson, M. I.; Harlos, K.; Claridge, T. D.; Schofield, C. J.; Pugh, C. W.; Maxwell, P. H.; Ratcliffe, P. J.; Stuart, D. I.; Jones, E. Y. Structural basis for the recognition of hydroxyproline in HIF-1*α* by pVHL. Nature 2002, 417, 975–978.

(12) Sakamoto, K. M.; Kim, K. B.; Kumagai, A.; Mercurio, F.; Crews, C. M.; Deshaies, R. J. Protacs: Chimeric molecules that target proteins to the Skp1–Cullin–F box complex for ubiquitination and degradation. Proceedings of the National Academy of Sciences 2001, 98, 8554–8559.

(13) Schapira, M.; Calabrese, M. F.; Bullock, A. N.; Crews, C. M. Targeted protein degradation: expanding the toolbox. Nature reviews Drug discovery 2019, 18, 949–963.

(14) Imaide, S.; Riching, K. M.; Makukhin, N.; Vetma, V.; Whitworth, C.; Hughes, S. J.; Trainor, N.; Mahan, S. D.; Murphy, N.; Cowan, A. D., et al. Trivalent PROTACs enhance protein degradation via combined avidity and co-operativity. Nature chemical biology 2021, 17, 1157–1167.

(15) Cowan, A. D.; Ciulli, A. Driving E3 Ligase Substrate Specificity for Targeted Protein Degradation: Lessons from Nature and the Laboratory. Annual Review of Biochemistry 2022, 91.

(16) Roy, M. J.; Winkler, S.; Hughes, S. J.; Whitworth, C.; Galant, M.; Farnaby, W. l.; Rumpel, K.; Ciulli, A. SPR-Measured Dissociation Kinetics of PROTAC Ternary Complexes Influence Target Degradation Rate. ACS CHEM BIOL 2019, 14, 361–368.

(17) Casement, R.; Bond, A.; Craigon, C.; Ciulli, A. Targeted Protein Degradation; Springer, 2021; pp 79–113.

(18) Rodriguez-Rivera, F. P.; Levi, S. M. Unifying catalysis framework to dissect proteasomal degradation paradigms. ACS Central Science 2021, 7, 1117–1125.

(19) Li, W.; Zhang, J.; Guo, L.; Wang, Q. Importance of Three-Body Problems and Protein–Protein Interactions in Proteolysis-Targeting Chimera Modeling: Insights from Molecular Dynamics Simulations. Journal of Chemical Information and Modeling 2022,

(20) Hughes, S.; Ciulli, A. Molecular recognition of ternary complexes: a new dimension in the structure-guided design of chemical degraders. ESSAYS BIOCHEM 2017, 61, 505–516.

(21) Zorba, A. et al. Delineating the role of cooperativity in the design of potent PROTACs for BTK. Proceedings of the National Academy of Sciences 2018, 115, 201803662.

(22) Schiemer, J. et al. Snapshots and ensembles of BTK and cIAP1 protein degrader ternary complexes. NAT CHEM BIOL 2021, 17, 152–160.

(23) Huang, H.-T.; Dobrovolsky, D.; Paulk, J.; Yang, G.; Weisberg, E. L.; Doctor, Z. M.; Buckley, D. L.; Cho, J.-H.; Ko, E.; Jang, J., et al. A chemoproteomic approach to query the degradable kinome using a multi-kinase degrader. Cell chemical biology 2018, 25, 88–99.

(24) Bondeson, D. P.; Smith, B. E.; Burslem, G. M.; Buhimschi, A. D.; Hines, J.; Jaime-Figueroa, S.; Wang, J.; Hamman, B. D.; Ishchenko, A.; Crews, C. M. Lessons in PROTAC design from selective degradation with a promiscuous warhead. Cell chemical biology 2018, 25, 78–87.

(25) Ward, C. C.; Kleinman, J. I.; Brittain, S. M.; Lee, P. S.; Chung, C. Y. S.; Kim, K.; Petri, Y.; Thomas, J. R.; Tallarico, J. A.; McKenna, J. M., et al. Covalent ligand screening uncovers a RNF4 E3 ligase recruiter for targeted protein degradation applications. ACS chemical biology 2019, 14, 2430–2440.

(26) Zengerle, M.; Chan, K.-H.; Ciulli, A. Selective Small Molecule Induced Degradation of the BET Bromodomain Protein BRD4. ACS CHEM BIOL 2015, 10, 1770–1777.

(27) Gadd, M. S.; Testa, A.; Lucas, X.; Chan, K.-H.; Chen, W.; Lamont, D. J.; Zẽngerle, M.; Ciulli, A. Structural basis of PROTAC cooperative recognition for selective protein degradation. NAT CHEM BIOL 2017, 13, 514–521.

(28) Farnaby, W. et al. BAF complex vulnerabilities in cancer demonstrated via structure-based PROTAC design. NAT CHEM BIOL 2019, 15, 672–680.

(29) Testa, A.; Hughes, S. J.; Lucas, X.; Wright, J. E.; Ciulli, A. Structure-Based Design of a Macrocyclic PROTAC. Angewandte Chemie International Edition 2020, 59, 1727–1734.

(30) Zaidman, D.; Prilusky, J.; London, N. PRosettaC: Rosetta Based Modeling of PROTAC Mediated Ternary Complexes. J CHEM INF MODEL 2020, 60, 4894–4903.

(31) Bai, N.; Kirubakaran, P.; Karanicolas, J. Rationalizing PROTAC-mediated ternary complex formation using Rosetta. J. Chem. Inf. Model. 2021, 61, 1368–1382.

(32) Drummond, M. L.; Henry, A.; Li, H.; Williams, C. I. Improved Accuracy for Modeling PROTAC-Mediated Ternary Complex Formation and Targeted Protein Degradation via New In Silico Methodologies. J CHEM INF MODEL 2020, 60, 5234–5254.

(33) Shaheer, M.; Singh, R.; Sobhia, M. E. Protein degradation: a novel computational approach to design protein degrader probes for main protease of SARS-CoV-2. J BIOMOL STRUCT DYN 2021, 1–13.

(34) Drummond, M. L.; Henry, A.; Li, H.; Williams, C. I. Improved Accuracy for Modeling PROTAC-Mediated Ternary Complex Formation and Targeted Protein Degradation via New In Silico Methodologies. J CHEM INF MODEL 2020, 60, 5234–5254.

(35) Eron, S. J.; Huang, H.; Agafonov, R. V.; Fitzgerald, M. E.; Patel, J.; Michael, R. E.; Lee, T. D.; Hart, A. A.; Shaulsky, J.; Nasveschuk, C. G.; Phillips, A. J.; Fisher, S. L.; Good, A. Structural Characterization of Degrader-Induced Ternary Complexes Using Hydrogen–Deuterium Exchange Mass Spectrometry and Computational Modeling: Implications for Structure-Based Design. ACS Chemical Biology 2021,

(36) Farnaby, W.; Koegl, M.; Roy, M. J.; Whitworth, C.; Diers, E.; Trainor, N.; Zollman, D.; Steurer, S.; Karolyi-Oezguer, J.; Riedmueller, C., et al. BAF complex vulnerabilities in cancer demonstrated via structure-based PROTAC design. Nature chemical biology 2019, 15, 672–680.

(37) Liu, X.; Zhang, X.; Lv, D.; Yuan, Y.; Zheng, G.; Zhou, D. Assays and technologies for developing proteolysis targeting chimera degraders. Future Medicinal Chemistry 2020, 12, 1155–1179.

(38) Jubb, H. C.; Higueruelo, A. P.; Ochoa-Montaño, B.; Pitt, W. R.; Ascher, D. B.; Blundell, T. L. Arpeggio: A Web Server for Calculating and Visualising Interatomic Interactions in Protein Structures. Journal of Molecular Biology 2017, 429, 365–371.

(39) Nowak, R. P.; DeAngelo, S. L.; Buckley, D.; He, Z.; Donovan, K. A.; An, J.; Safaee, N.; Jedrychowski, M. P.; Ponthier, C. M.; Ishoey, M.; Zhang, T.; Mancias, J. D.; Gray, N. S.; Bradner, E. S., J. E. Fischer Plasticity in binding confers selectivity in ligand-induced protein degradation. Nature Chemical Biology 2018, 14, 706–714.

(40) Deller, M. C.; Kong, L.; Rupp, B. Protein stability: A crystallographer’s perspective. Acta Crystallogr F Struct Biol Commun 2016, 72, 72–95.

(41) Skinner, S. P.; Radou, G.; Tuma, R.; Houwing-Duistermaat, J. J.; Paci, E. Estimating Constraints for Protection Factors from HDX-MS Data. Biophysical Journal 2019, 116, 1194–1203.

(42) Devaurs, D.; Antunes, D. A.; Borysik, A. J. Computational Modeling of Molecular Structures Guided by Hydrogen-Exchange Data. Journal of the American Society for Mass Spectrometry 2022, 33, 215–237, PMID: 35077179.

(43) Wales, T. E.; Engen, J. R. Hydrogen exchange mass spectrometry for the analysis of protein dynamics. MASS SPECTROM REV 2006, 25, 158–170.

(44) Gallagher, E. S.; Hudgens, J. W. Mapping Protein-Ligand Interactions with Proteolytic Fragmentation, Hydrogen/Deuterium Exchange-Mass Spectrometry. Methods in Enzymology 2016, 566.

(45) Huber, G. A.; Kim, S. Weighted-ensemble Brownian dynamics simulations for protein association reactions. BIOPHYS J 1996, 70, 97–110.

(46) Zuckerman, D. M.; Chong, L. T. Weighted Ensemble Simulation: Review of Methodology, Applications, and Software. ANN REV BIOPHYS 2017, 46, 43–57.

(47) Saglam, A. S.; Chong, L. T. Protein–protein binding pathways and calculations of rate constants using fully-continuous, explicit-solvent simulations. Chemical Science 2018, 10, 2360–2372.

(48) Dickson, A. Mapping the Ligand Binding Landscape. Biophysical Journal 2018, 115, 1707–1719.

(49) Méndez, R.; Leplae, R.; De Maria, L.; Wodak, S. J. Assessment of blind predictions of protein–protein interactions: Current status of docking methods. PROTEINS 2003, 52, 51–67.

(50) Huang, L.; So, P.-K.; Yao, Z.-P. Protein Dynamics Revealed by Hydrogen Deuterium Exchange Mass Spectrometry: Correlation between Experiments and Simulation. Rapid communications in mass spectrometry: RCM 2018, 33, 83– 89.

(51) Lotz, S. D.; Dickson, A. Wepy: A Flexible Software Framework for Simulating Rare Events with Weighted Ensemble Resampling. ACS Omega 2020, 5, 31608– 31623.

(52) Dixon, T.; Uyar, A.; Ferguson-Miller, S.; Dickson, A. Membrane-Mediated Ligand Unbinding of the PK-11195 Ligand from TSPO. Biophysical Journal 2021, 120, 158–167.

(53) Copperman, J.; Zuckerman, D. M. Accelerated Estimation of Long-Timescale Kinetics from Weighted Ensemble Simulation via Non-Markovian “Microbin” Analysis. Journal of Chemical Theory and Computation 2020, 16, 6763–6775.

(54) DeGrave, A. J.; Bogetti, A. T.; Chong, L. T. The RED scheme: Rate-constant estimation from pre-steady state weighted ensemble simulations. The Journal of Chemical Physics 2021, 154, 114111.

(55) Zhang, M. M.; Beno, B. R.; Huang, R. Y.-C.; Adhikari, J.; Deyanova, E. G.; Li, J.; Chen, G.; Gross, M. L. An Integrated Approach for Determining a Protein–Protein Binding Interface in Solution and an Evaluation of Hydrogen–Deuterium Exchange Kinetics for Adjudicating Candidate Docking Models. Anal. Chem. 2019, 91, 15709–15717.

(56) Scherer, M. K.; Trendelkamp-Schroer, B.; Paul, F.; Pérez-Hernández, G.; Hoffmann, M.; Plattner, N.; Wehmeyer, C.; Prinz, J.-H.; Noé, F. PyEMMA 2: A Software Package for Estimation, Validation, and Analysis of Markov Models. Journal of Chemical Theory and Computation 2015, 11, 5525–5542.

(57) Husic, B. E.; Pande, V. S. Markov state models: From an art to a science. Journal of the American Chemical Society 2018, 140, 2386–2396.

(58) Molgedey, L.; Schuster, H. G. Separation of a mixture of independent signals using time delayed correlations. Phys. Rev. Lett. 1994, 72, 3634–3637.

(59) Husic, B. E.; McGibbon, R. T.; Sultan, M. M.; Pande, V. S. Optimized parameter selection reveals trends in Markov state models for protein folding. The Journal of chemical physics 2016, 145, 194103.

(60) Buhimschi, A. D.; Crews, C. M. Evolving rules for protein degradation? Insights from the zinc finger degrome. Biochemistry 2019, 58, 861–864.

(61) Baek, K.; Krist, D. T.; Prabu, J. R.; Hill, S.; Klügel, M.; Neumaier, L.-M.; von Gronau, S.; Kleiger, G.; Schulman, B. A. NEDD8 nucleates a multivalent cullin–RING–UBE2D ubiquitin ligation assembly. Nature 2020, 578, 461–466.

(62) Yauch, R.; Cantley, J.; Ye, X.; Rousseau, E.; Januario, T.; Hamman, B.; Rose, C.; Cheung, T.; Hickle, T.; Soto, L., et al. Selective PROTAC-mediated degradation of SMARCA2 is efficacious in SMARCA4 mutant cancers. 2022,

(63) Dagbay, K. B.; Bolik-Coulon, N.; Savinov, S. N.; Hardy, J. A. Caspase-6 Undergoes a Distinct Helix-Strand Interconversion upon Substrate Binding*. J BIOL CHEM 2017, 292, 4885–4897.

(64) Dagbay, K. B.; Hardy, J. A. Multiple proteolytic events in caspase-6 self-activation impact conformations of discrete structural regions. P NATL ACAD SCI USA 2017, 114, E7977–E7986.

(65) MacPherson, D. J.; Mills, C. L.; Ondrechen, M. J.; Hardy, J. A. Tri-arginine exosite patch of caspase-6 recruits substrates for hydrolysis. J BIOL CHEM 2019, 294, 71–88.

(66) Kochert, B. A.; Iacob, R. E.; Wales, T. E.; Makriyannis, A.; Engen, J. R. Hydrogen-Deuterium Exchange Mass Spectrometry to Study Protein Complexes. Methods in Molecular Biology 2018, 1764, 153–171.

(67) Wales, T. E.; Fadgen, K. E.; Gerhardt, G. C.; Engen, J. R. High-Speed and High-Resolution UPLC Separation at Zero Degrees Celsius. AANAL BIOANAL CHEM 2008, 80, 6815–6820.

(68) Hopkins, J. B.; Gillilan, R. E.; Skou, S. *BioXTAS RAW*: improvements to a free open-source program for small-angle X-ray scattering data reduction and analysis. J APPL CRYSTALLOGR 2017, 50, 1545–1553.

(69) Maier, J. A.; Martinez, C.; Kasavajhala, K.; Wickstrom, L.; Hauser, K. E.; Simmerling, C. ff14SB: Improving the Accuracy of Protein Side Chain and Backbone Parameters from ff99SB. J CHEM THEORY COMPUT 2015, 11, 3696–3713, PMID: 26574453.

(70) Zwier, M. C.; Adelman, J. L.; Kaus, J. W.; Pratt, A. J.; Wong, K. F.; Rego, N. B.; Suarez, E.; Lettieri, S.; Wang, D. W.; Grabe, M.; Zuckerman, D. M.; Chong, L. T. WESTPA: An Interoperable, Highly Scalable Software Package for Weighted Ensemble Simulation and Analysis. J CHEM THEORY COMPUT 2015, 11, 800–809.

(71) Russo, J. D. et al. WESTPA 2.0: High-Performance Upgrades for Weighted Ensemble Simulations and Analysis of Longer-Timescale Applications. Journal of Chemical Theory and Computation 2022, 18, 638–649.

(72) Zhang, B. W.; Jasnow, D.; Zuckerman, D. M. The “weighted ensemble” path sampling method is statistically exact for a broad class of stochastic processes and binning procedures. J CHEM PHYS 2010, 132, 054107.

(73) Pearlman, D. A.; Case, D. A.; Caldwell, J. W.; Ross, W. S.; Cheatham III, T. E.; DeBolt, S.; Ferguson, D.; Seibel, G.; Kollman, P. AMBER, a package of computer programs for applying molecular mechanics, normal mode analysis, molecular dynamics and free energy calculations to simulate the structural and energetic properties of molecules. COMPUT PHYS COMMUN 1995, 91, 1–41.

(74) Pracht, P.; Bohle, F.; Grimme, S. Automated exploration of the low-energy chemical space with fast quantum chemical methods. Phys. Chem. Chem. Phys. 2020, 22, 7169–7192.

(75) Grimme, S. Exploration of Chemical Compound, Conformer, and Reaction Space with Meta-Dynamics Simulations Based on Tight-Binding Quantum Chemical Calculations. J. Chem. Theory Comput. 2019, 15, 2847–2862.

(76) Bannwarth, C.; Ehlert, S.; Grimme, S. GFN2-xTB—An Accurate and Broadly Parametrized Self-Consistent Tight-Binding Quantum Chemical Method with Multipole Electrostatics and Density-Dependent Dispersion Contributions. J. Chem. Theory Comput. 2019, 15, 1652–1671.

(77) Gray, J. J.; Moughon, S.; Wang, C.; Schueler-Furman, O.; Kuhlman, B.; Rohl, C. A.; Baker, D. Protein–Protein Docking with Simultaneous Optimization of Rigid-body Displacement and Side-chain Conformations. J. Mol. Biol. 2003, 331, 281–299.

(78) Marze, N. A.; Roy Burman, S. S.; Sheffler, W.; Gray, J. J. Efficient flexible backbone protein–protein docking for challenging targets. Bioinformatics 2018, 34, 3461–3469.

(79) Maier, J. A.; Martinez, C.; Kasavajhala, K.; Wickstrom, L.; Hauser, K. E.; Simmerling, C. ff14SB: Improving the Accuracy of Protein Side Chain and Backbone Parameters from ff99SB. J CHEM THEORY COMPUT 2015, 11, 3696–3713.

(80) Jorgensen, W. L.; Chandrasekhar, J.; Madura, J. D.; Impey, R. W.; Klein, M. L. Comparison of simple potential functions for simulating liquid water. J CHEM PHYS 1983, 79, 926–935.

(81) Hess, B.; Bekker, H.; Berendsen, H. J. C.; Fraaije, J. G. E. M. LINCS: A linear constraint solver for molecular simulations. J COMPUT CHEM 1997, 18, 1463– 1472.

(82) Gunsteren, W. F. V.; Berendsen, H. J. C. A Leap-frog Algorithm for Stochastic Dynamics. MOL SIMULAT 1988, 1, 173–185.

(83) Darden, T.; York, D.; Pedersen, L. Particle mesh Ewald: An N*·*log(N) method for Ewald sums in large systems. jcp 1993, 98, 10089–10092.

(84) Bussi, G.; Donadio, D.; Parrinello, M. Canonical sampling through velocity rescaling. J CHEM PHYS 2007, 126, 014101.

(85) Parrinello, M.; Rahman, A. Polymorphic transitions in single crystals: A new molecular dynamics method. J APPL PHYS 1981, 52, 7182–7190.

(86) Cao, S.; Montoya-Castillo, A.; Wang, W.; Markland, T. E.; Huang, X. On the advantages of exploiting memory in Markov state models for biomolecular dynamics. The Journal of Chemical Physics 2020, 153, 014105.

(87) Svergun, D.; Barberato, C.; Koch, M. H. J. *CRYSOL* – a Program to Evaluate X-ray Solution Scattering of Biological Macromolecules from Atomic Coordinates. J APPL CRYSTALLOGR 1995, 28, 768–773.

(88) Manalastas-Cantos, K.; Konarev, P. V.; Hajizadeh, N. R.; Kikhney, A. G.; Petoukhov, M. V.; Molodenskiy, D. S.; Panjkovich, A.; Mertens, H. D. T.; Gruzinov, A.; Borges, C.; Jeffries, C. M.; Svergun, D. I.; Franke, D. *ATSAS 3.0*: expanded functionality and new tools for small-angle scattering data analysis. J APPL CRYSTALLOGR 2021, 54, 343–355.

(89) Borreguero, J. M.; Islam, F. F.; Shrestha, U. R.; Petridis, L. idpflex: Analysis of Intrinsically Disordered Proteins by Comparing Simulations to Small Angle Scattering Experiments. Journal of Open Source Software 2018, 3.

(90) Cheng, X.; Wang, H.; Grant, B.; Sine, S. M.; McCammon, J. A. Targeted molecular dynamics study of C-loop closure and channel gating in nicotinic receptors. PLoS computational biology 2006, 2, e134.

(91) Edmondson, S. D.; Yang, B.; Fallan, C. Proteolysis Targeting Chimeras (PROTACs) in ‘Beyond Rule-of-Five’ Chemical Space: Recent Progress and Future Challenges. BIOORG MED CHEM LETT 2019, 29, 1555–1564.

(92) Comer, J.; Gumbart, J. C.; Hénin, J.; Lelièvre, T.; Pohorille, A.; Chipot, C. The adaptive biasing force method: Everything you always wanted to know but were afraid to ask. The Journal of Physical Chemistry B 2015, 119, 1129–1151.

(93) Lesage, A.; Lelievre, T.; Stoltz, G.; Hénin, J. Smoothed biasing forces yield unbiased free energies with the extended-system adaptive biasing force method. The Journal of Physical Chemistry B 2017, 121, 3676–3685.

(94) Fu, H.; Zhang, H.; Chen, H.; Shao, X.; Chipot, C.; Cai, W. Zooming across the free-energy landscape: shaving barriers, and flooding valleys. The journal of physical chemistry letters 2018, 9, 4738–4745.

(95) Fu, H.; Shao, X.; Cai, W.; Chipot, C. Taming rugged free energy landscapes using an average force. Accounts of chemical research 2019, 52, 3254–3264.

(96) Eastman, P.; Swails, J.; Chodera, J. D.; McGibbon, R. T.; Zhao, Y.; Beauchamp, K. A.; Wang, L.-P.; Simmonett, A. C.; Harrigan, M. P.; Stern, C. D., et al. OpenMM 7: Rapid development of high performance algorithms for molecular dynamics. PLoS computational biology 2017, 13, e1005659.

(97) Bonomi, M. Promoting transparency and reproducibility in enhanced molecular simulations. Nature methods 2019, 16, 670–673.

(98) Zougman, A.; Selby, P. J.; Banks, R. E. Suspension trapping (STrap) sample preparation method for bottom-up proteomics analysis. PROTEOMICS 2014, 14, 1006–1000.

(99) Udeshi, N. D.; Mani, D. C.; Satpathy, S.; Fereshetian, S.; Gasser, J. A.; Svinkina, T.; Olive, M. E.; Ebert, B. L.; Mertins, P.; Carr, S. A. Rapid and deep-scale ubiquitylation profiling for biology and translational research. Nature Communications 2020, 11, 359.

(100) Ruprecht, B.; Zecha, J.; Zolg, D. P.; Kuster, B. Proteomics. Methods in Molecular Biology 2017, 1550, 83–98.

(101) Ritchie, M. E.; Phipson, B.; Wu, D.; Hu, Y.; Law, C. W.; Shi, W.; Smyth, G. K. limma powers differential expression analyses for RNA-sequencing and microarray studies. Nucleic acids research 2015, 43, e47.

(102) Perez-Riverol, Y.; Bai, J.; Bandla, C.; García-Seisdedos, D.; Hewapathirana, S.; Kamatchinathan, S.; Kundu, D. J.; Prakash, A.; Frericks-Zipper, A.; Eisenacher, M.; Walzer, M.; Wang, S.; Brazma, A.; Vizcaíno, J. A. The PRIDE database resources in 2022: a hub for mass spectrometry-based proteomics evidences. Nucleic acids research 2022, 50, D543–D552.

