## Supplementary Information for "Atomic-Resolution Prediction of Degrader-mediated Ternary Complex Structures by Combining Molecular Simulations with Hydrogen Deuterium Exchange"

#### X-ray crystallography.

Supplemental Table 1: Crystallographic table for protein crystal structure 7S4E iso2-SMARCA2$^{BD}$:ACBI1:VHL. Asterisk (*) denotes values obtained for the highest-resolution shell.

| | | $SMARCA2^{BD}$: ACBI1 : VCB |
|---|---|---|
| ***Data collection*** | | |
| Space Group | | P 21 21 21 |
| ***Cell Dimension*** | | |
| a, b, c, (Å) | | 80.14, 116.57, 122.32 |
| α, β, ɣ (˚) | | 90, 90, 90 |
| Resolution (Å) | | 37.89-2.25 (2.31-2.25)* |
| $R_{merge}$ | | 0.15 (2.602)* |
| <I/σI> | | 8.0 (1.27)* |
| CC(1/2) | | 0.998 (0.317)* |
| Completeness (%) | | 99.9 (99.9)* |
| Redundancy | | 7.4 (7.4)* |
| ***Refinement*** | | |
| Resolution (Å) | | 2.25 |
| No. Reflections | | 52206 |
| $R_{work}/R_{merge}$ | | 21.9/25.9 |
| No. Atoms | | 7356 |
| | Protein | 7070 |
| | Ligand/Ions | 196 |
| | Water | 90 |
| ***B factors*** | | |
| | Protein | 61.19 |
| | Ligand/Ions | 57.66 |
| | Water | 53.56 |
| ***R.M.S deviations*** | | |
| Bond length (Å) | | 0.009 |
| Bond angles (˚) | | 1.519 |
| Ramachandran Favored (%) | | 96.73 |
| Ramachandran Outliers (%) | | 0.00 |
| Clashscore | | 3.64 |

** Denotes values obtained for the highest-resolution shell.*

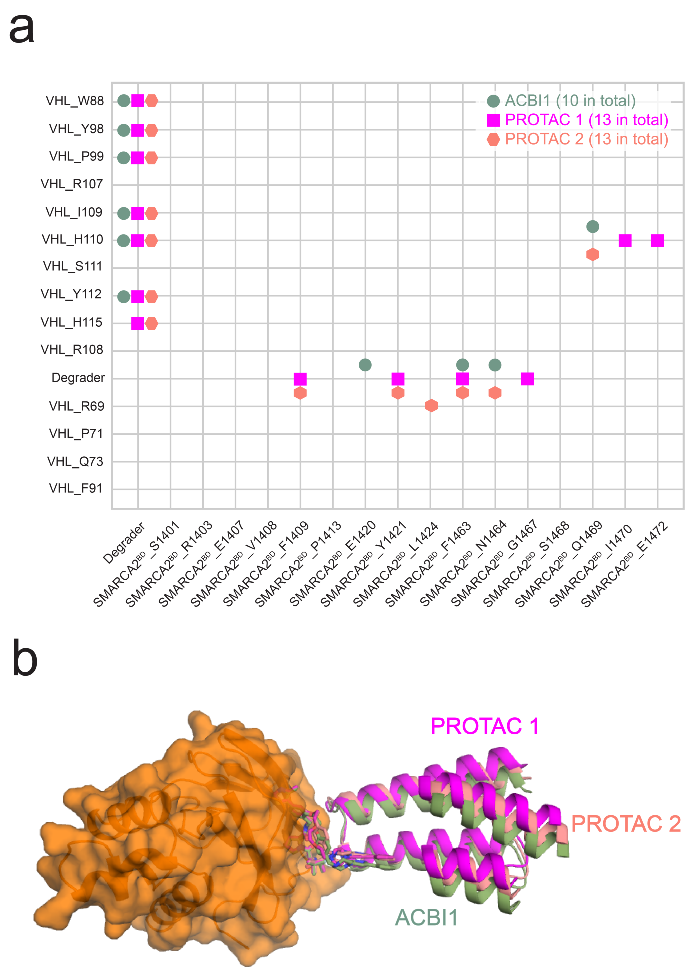


Supplemental Fig. 1: Comparison of the crystal structures of SMARCA2$^{BD}$:VHL complex with PROTAC 1 (PDB ID: 6HAY), PROTAC 2 (PDB ID: 6HAX), and ACBI1 (PDB ID: 7SE4) bound. (**a**) Protein-protein and protein-degrader contacts observed in the ternary complex crystal structures. The total number of contacts are indicated for each complex in the top right. (**b**) The alignment of the three crystal structures. VHL is colored in orange and shown in cartoon and surface representation. SMARCA2$^{BD}$ of the ternary complexes with PROTAC 1, PROTAC 2 and ACBI1 are shown in cartoon and colored in magenta, salmon, and green respectively.

#### WE-HDX simulations.

The weighted-ensemble (WE) strategy enhances the sampling of rare events by running in parallel multiple simulations with well-defined probabilities. The periodic pruning and replication of trajectories allows progress to be made along a collective variable in conformational space, either by simulating trajectories in pre-defined regions ("binned method") or by optimizing an objective function ("bin-less method"). For the formation of degrader ternary complexes, we have applied both WE simulation variants. Mainly, we run the bin-less REVO simulations[1] that maximize an objective function called the trajectory variation, defined as the sum of distances between individual trajectories. The distance metric itself is based on observables such as the number of atomic contacts or the warhead-RMSD (w-RMSD) with respect to the crystal structure of the target-warhead complex. Alternatively, in the binned WE method, we use these observables as collective variables to sample less-visited regions of conformational space.

Data from HDX-MS experiments are integrated with the WE simulations in a straightforward manner to facilitate the formation of contacts between the protected residues on SMARCA2$^{BD}$ and those on VHL as determined from HDX-MS (see Supplemental Table 2). This is achieved by replacing *any* SMARCA2$^{BD}$:VHL residue contact, in the (non-HDX) WE simulations, specifically with experimentally derived protected-residue contacts at the interface of the two binding partners, in the WE-HDX simulations. Detailed information on the WE-HDX simulations is provided in the Methods.

Supplemental Fig. 2 reveals that both WE variants yield similar results in that the bulk of simulated SMARCA2$^{BD}$:PROTAC 2:VHL ternary complexes, in particular when guided by the HDX-MS data, have (minimum) interface-RMSDs (I-RMSDs) $< 4$ Å. The inclusion of the HDX-MS data, i.e., in WE-HDX and Docking-HDX, yields distributions of "tighter bound" ternary complexes as they are shifted toward smaller minimum I-RMSD values – in particular for the WE-HDX simulations. Although the (non-HDX) WE simulations can sporadically produce SMARCA2$^{BD}$:PROTAC 2:VHL ternary complexes with a minimum I-RMSD that

Supplemental Table 2: Protected residues on VHL and SMARCA2$^{BD}$ used in the WE-HDX simulations.

| Protein | Protected residues |
|---|---|
| VHL | R60, V62, L63, R64, S65, V66, N67, S68, R69, E70, S72, Q73 |
| SMARCA2$^{BD}$ | F1409, I1410, Q1411, L1412, S1414, R1415, K1416, E1417, L1418, E1420, Y1421, Y1422, E1423, L1424, L1456, C1457, H1458, N1459, A1460, Q1461, T1462, F1463, N1464, L1465, E1466, G1467, S1468, Q1469, I1470 |

is even $< 0.5$ Å (see red profiles in Supplemental Figs. 2, 3), the notion of "tighter bound" ternary complexes is clearly confirmed by the discrepancy in the solvent-accessible surface area (SASA) measured for the HDX-derived protected residues, that is observed between WE and WE-HDX simulations (shown as a function of the minimum I-RMSD in Supplemental Fig. 3a). The reduced solvent-accessibility of protected residues in WE-HDX compared to the same residues in WE simulations suggests that the protein-protein interfaces of the corresponding structures are less solvent-exposed, i.e., the simulated SMARCA2$^{BD}$:PROTAC 2:VHL ternary complexes are bound tighter, in the WE-HDX simulations.

The enhanced ternary complex simulation by WE-HDX compared to WE is further emphasized in Supplemental Fig. 3b, showing that the HDX-augmented method converges toward structures with low minimum I-RMSD *and* low $C_\alpha$-RMSD values (with respect to a diverse set of ternary complex structures, as discussed below). These results demonstrate that the addition of HDX-MS information significantly improves the prediction accuracy of ternary complex WE simulations.

In previous work, the $C_\alpha$-RMSD of the entire ternary complex has been used as a metric to assess the accuracy of ternary complex predictions.[2] However, the enhanced simulations as

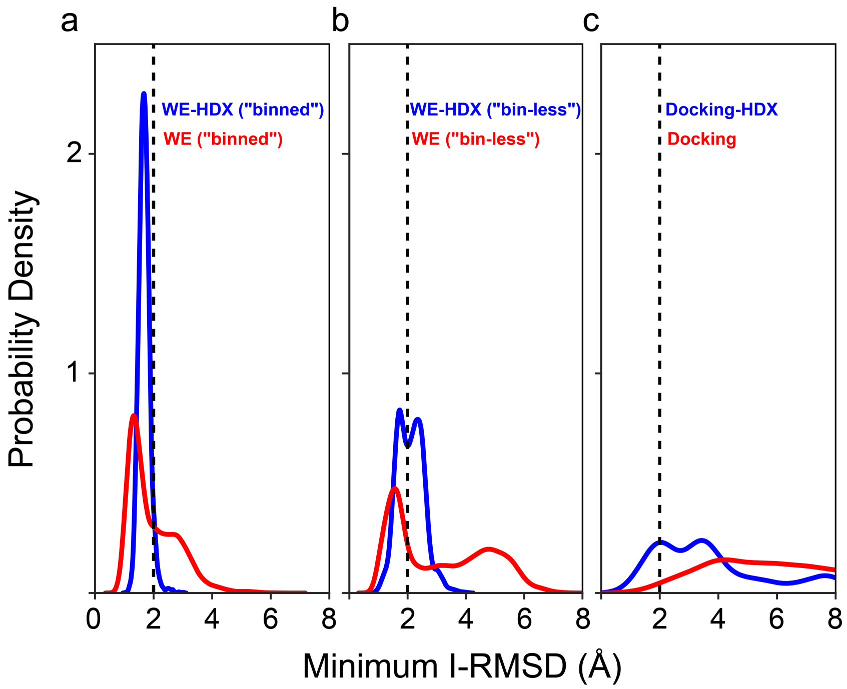


Supplemental Fig. 2: Comparison of WE simulations (**a**, "binned method", **b**, "bin-less method") of ternary complex formation and (**c**) ternary complex docking for SMARCA2$^{BD}$:PROTAC 2:VHL with (red) and without (blue) data from HDX-MS experiments. The vertical dashed lines indicate the threshold at 2 Å used to define a bound ternary complex (see discussion below).

well as the experiments presented in the main text reveal that the key structural determinants of differences in degradation efficiencies among the three degrader molecules studied are found at the SMARCA2$^{BD}$:VHL interface. Therefore, we focus on the interface-RMSD (I-RMSD) with respect to bound reference structures as the main parameter to evaluate the ternary complexes simulated with the WE-HDX method.

Importantly, in order to not only compare to a single reference structure, which, most likely, would be the experimentally obtained crystal structure, we apply an approach that compares the simulated ternary complex to a set of structurally diverse conformations. This procedure is crucial for a more accurate estimation of the validity of our simulated ternary conformations, which, as demonstrated in the main text, are highly flexible aggregates.

We derive the set of reference ternary structures from long ($> 1$ $\mu s$) brute-force MD simulations (see Methods). These simulations started from the corresponding ternary com-

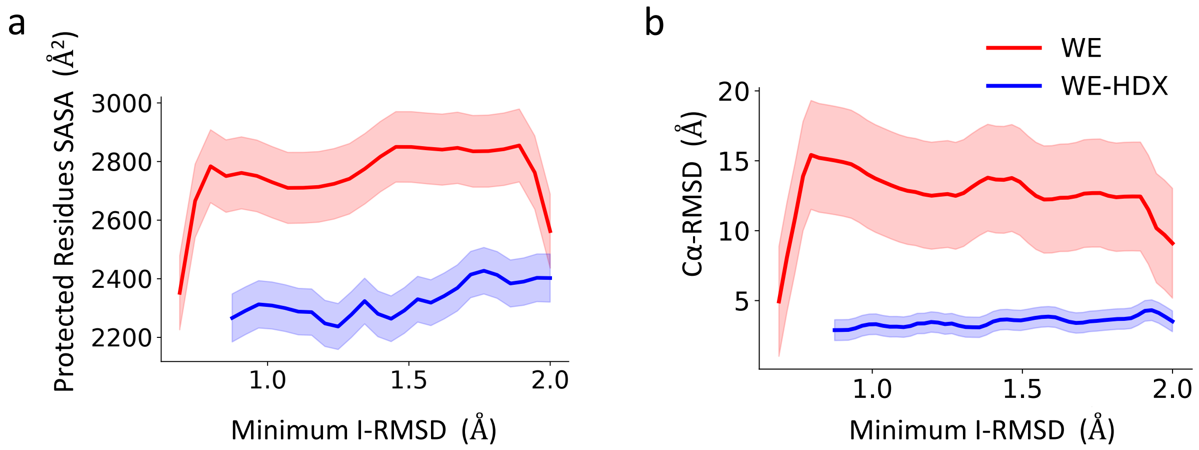


Supplemental Fig. 3: Minimum I-RMSD of SMARCA2$^{BD}$:PROTAC 2:VHL versus (**a**) the solvent-accessible surface area (SASA) of the HDX-protected residues (see Supplemental Table 2) and (**b**) the $C_\alpha$-RMSD during WE (red) and WE-HDX (blue) simulations of ternary complex formation. The minimum I-RMSD is with respect to a diverse set of reference structures (as described in the text) and the distribution of (**a**) SASAs or (**b**) $C_\alpha$-RMSDs is obtained for those structures with the minimum I-RMSDs. The solid lines and shaded regions are the arithmetic mean and standard error of (**a**) the SASAs or (**b**) the $C_\alpha$-RMSDs, respectively.

plex crystal structures with PROTAC 1, PROTAC 2, or ACBI1 and they sample, for each system, a variety of different binding poses. A $k$-means clustering based on interface residue distances, divides all sampled conformations into $k = 25$ subsets of relatively diverse ternary complexes. The 25 representative structures (or cluster centers) obtained, together with the experimental X-ray crystal structure, constitute the set of reference conformations used for the aforementioned I-RMSD calculations. Supplemental Fig. 4 presents superpositions of these reference sets for the three degraders connecting SMARCA2$^{BD}$ to VHL, highlighting the structural heterogeneity among them.

We use minimum I-RMSD with respect to this set of structures as the metric to assess the quality of predicted ternary complexes. As illustrated in Supplemental Fig. 3b, all sampled conformations with a minimum I-RMSD $< 2$ Å have a corresponding $C_\alpha$-RMSD $\leq \sim 5$ Å, which is clearly below the $C_\alpha$-RMSD threshold of 10 Å used by Drummond et al.,[3] thus suggesting that 2 Å is an appropriate I-RMSD threshold value to define bound ternary complexes in our study.

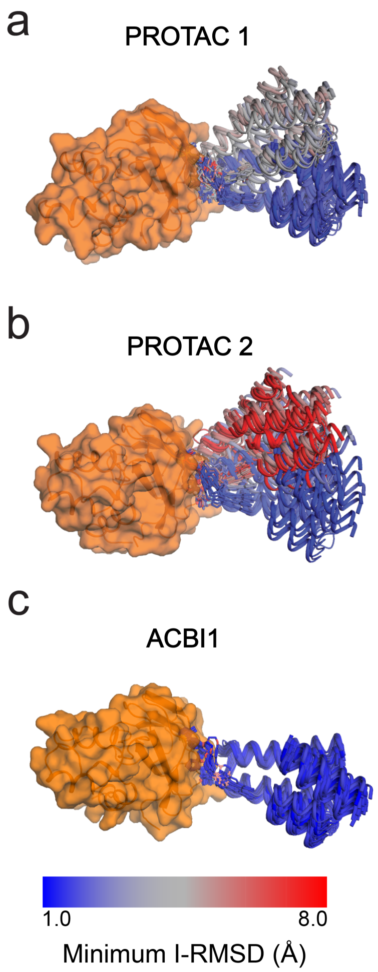


Supplemental Fig. 4: Superposition of the set of reference structures used for the analysis of SMARCA2$^{BD}$:degrader:VHL ternary complex predictions with a) PROTAC1, b) PROTAC 2, and c) ACBI1. The individual VHL molecules (orange, surface representation) are aligned, the degraders are shown in stick, and the SMARCA2$^{BD}$ in cartoon representations. SMARCA2$^{BD}$ is colored based on the I-RMSD value of the ternary complex with respect to the corresponding crystal structure (see color bar). Obviously, the ternary complex reference structures with PROTAC 2 are most diverse among the three degraders.

The synergy of WE simulations with information on protected residues obtained from HDX experiments is particularly useful for the formation of multi-component aggregates, such as the degrader ternary complexes, when a relatively large solvent-exposed surface is buried upon binding. More generally, the combination of experimental hydrogen-deuterium exchange data with computational modeling and simulation, which has surged over the last decade, can be divided into qualitative and quantitative approaches (as reviewed by Devaurs et al.[4]). In a vast number of recently published studies, HDX protection data are converted into distance or binding restraints that aid in the selection of docked protein complexes,[5–8] such as antibody-antigen[9] or enzyme-inhibitor[10] pairs, and have ultimately led to the development of integrated strategies.[11–13] Notably, Eron et al. have applied this approach for the prediction of degrader ternary complexes,[14] as described in the main text. These methods have recently also been augmented by molecular simulations, which were either informed by restraints[15,16] or were used to provide atomistic detail on experimentally derived interactions of important membrane and cytosolic proteins, thus establishing a qualitative connection between HDX experiments and simulation.[17–24]

On the other hand, all quantitative methods have in common that a "protection factor" is estimated (usually from molecular simulations) and correlated with results from HDX experiments.[25] This estimate has previously been computed in various ways, such as through the solvent-exposure of amide groups,[26,27] their hydrogen bond propensity,[28–30] or even the protein backbone acidity/reactivity[31,32] and flexibility,[33] in combination with packing densities.[34,35] Since the correlation to experimental protection factors is better when predictions stem from an ensemble of structures rather than from a single structure,[36,37] recent studies have relied on docking[38] and coarse-grained models[39] or enhanced and accelerated simulation techniques[40,41] with reweighting protocols.[42]

Clearly, the WE-HDX method, as implemented in our study, is a qualitative approach that correlates protection data with molecular simulations, as no protection factors or intensities are computed based on the simulated structures. Nevertheless, WE-HDX is distinctly different from most qualitative HDX-modeling protocols in that the WE simulations, which themselves are often referred to as "unbiased" due to the omission of steering forces or biasing potentials, are *guided* by information from HDX experiments. The conversion of protection data into interface residue contacts used as a collective variable in the WE-HDX simulations is, to our knowledge, a novel and, based on the results of degrader ternary complex formation, a most intriguing strategy to experimentally augment the simulation of rare events with minimal bias.

### Docking protocol.

The docking protocol relies on the core assumption that high-fidelity structures are available for both the SMARCA2$^{BD}$:warhead and the VHL:E3-ligand binary complexes. The RosettaDock keeps relative poses of the degrader moieties fixed with respect to their bound protein partners. Thus, if $A$ is the chain ID of SMARCA2$^{BD}$, $X$ is the chain ID of the warhead attached to SMARCA2$^{BD}$, $B$ is the chain ID of VHL and $Y$ is the chain ID of the E3-ligand attached to VHL, the command-line to run RosettaDock is the following:

```
docking_protocol.linuxgccrelease -database database/
-s input_0001.pdb -nstruct $NUM_STR -in:file:extra_res_fa
warhead.params  ligand.params -use_input_sc -docking
-dock_pert 2.7 15 -partners AX_BY -ex1 -ex2aro
-constraints:cst_file constraints.txt
-constraints:cst_fa_weight 10 -out:file:scorefile score.sc
```

This will keep the chains $A$ and $X$ fixed and dock chains $B$ and $Y$ as a single docking partner with respect to $AX$.

The input PDB file input_0001.pdb is a structure with pre-packed side-chains obtained by running:

```
docking_prepack_protocol.linuxgccrelease -database database/
-s input.pdb -use_input_sc -extra_res_fa warhead.params ligand.params
```

Alignment of the linker conformers is performed with the tool presented in:[43]

```
python ternary_model_prediction.py -da decoy_atom_list.txt
-la linker_atom_list.txt -wd decoy_atom_list_delete.txt
-ld linker_atom_list_delete.txt -dl listDecoysPDB.txt
-ll listConformersPDB.txt -c 0.3 -r rmsd_0.3A.txt -t specify
```

Generation of conformers for the linker is performed with a set of in-house developed Python programs that wrap the CREST software.[44] Pre-processing, post-processing and re-ranking is performed with a set of in-house Python scripts. Calculation of CAPRI parameters is performed by RosettaDock with respect to the input co-crystallized structures.

Supplemental Fig. 5 shows distributions of the top-N docking predictions into the four conventional CAPRI quality categories:[45] Incorrect, Acceptable, Medium and High. Obviously, the prediction accuracy is higher when HDX-derived restraints are incorporated (green bars) compared to the conventional docking protocol (orange bars) for Acceptable and Medium and High quality categories at all values of top-N. At the same time, the count of Incorrect predictions is lower for docking "with HDX" data, indicating the advantage of using the restraints derived from the HDX data. This advantage is especially striking for the High-quality predictions: without the HDX restraints, there are no High-quality structures within top-10 and top-50 predictions (see Table 3).

Supplemental Fig. 6 (**a, b, c**) shows the mean values for the fraction of native contacts, $fNat$, which are evaluated for docking predictions with respect to the crystal structure (PDB ID: 6HAX). The observation that HDX-derived restraints lead to higher values of $fNat$ (green versus orange bars) is a direct consequence of the nature of information determined in the experiments: sets of most protected residues that allegedly form the interface. We see this as an evidence for complementarity of employed methods and consistency of our research. Supplemental Fig. 6 (**d, e, f**) and (**g, h, i**) show the distributions of I-RMSD and C$\alpha$-RMSD values. Following ,[3] the C$\alpha$-RMSD is defined as the root-mean-square deviation of C$\alpha$ coordinates after a rigid-body superposition of the whole predicted structure onto the corresponding co-crystallized one. As seen from Supplemental Fig. 6, most of the predictions have C$\alpha$-RMSD values less than 10 Å and fall into the "crystal-like" category as defined by Drummond et al.[3] Also, the values of I-RMSD and C$\alpha$-RMSD are lower when the HDX-derived restraints are used. Similar results are obtained for the ternary complexes with PROTAC 1 and ACBI1 (see Supplemental Figs. 10 – 13).

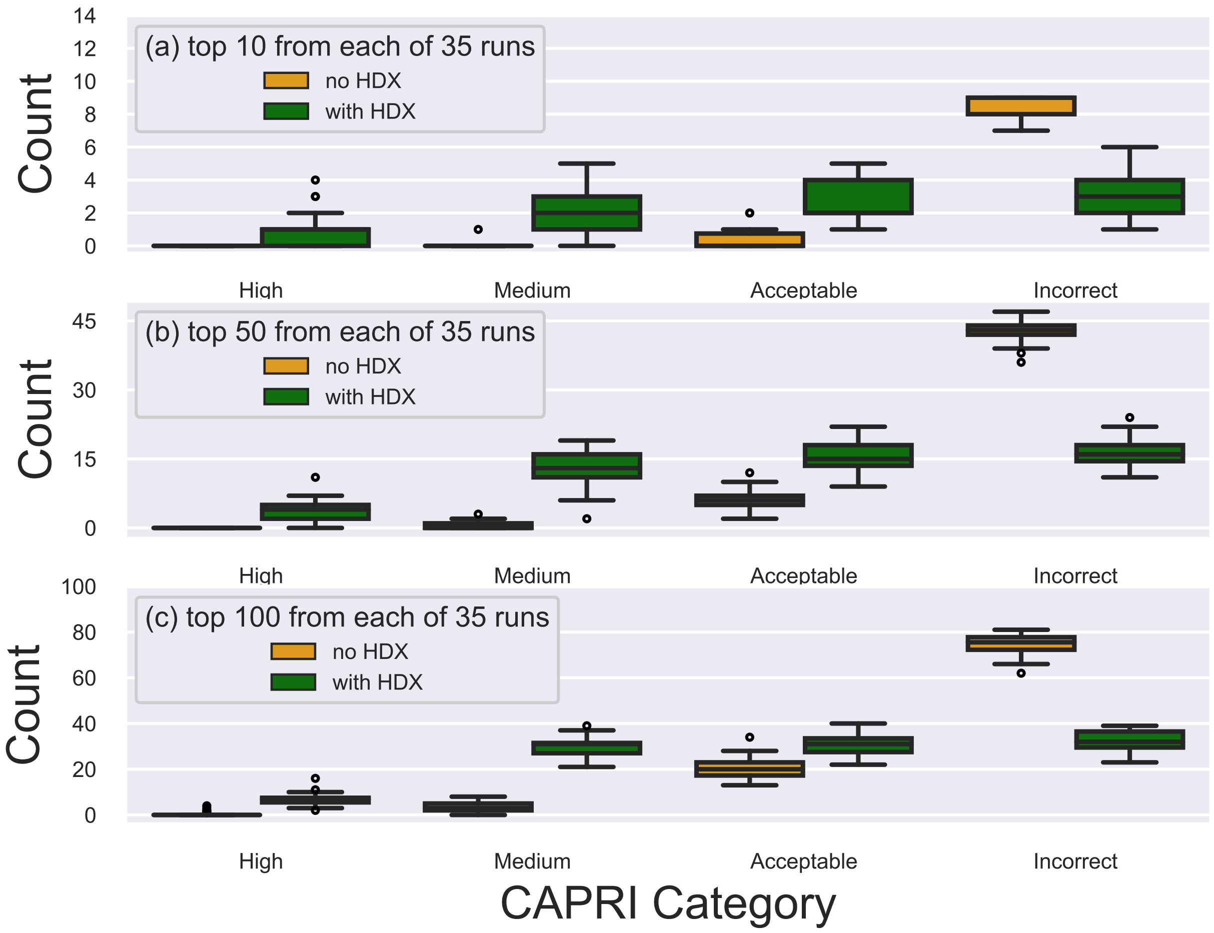


Supplemental Fig. 5: Distributions of the top-N docking predictions into the CAPRI quality categories High, Medium, Acceptable, and Incorrect. 35 independent docking runs have been performed and mean values have been calculated for (**a**) the top-10, (**b**) the top-50, and (**c**) the top-100 docking predictions with (green) and without (orange) HDX-derived restraints. Individual data points are shown as circles. The median, first and third quartiles are shown as horizontal lines. Results presented for SMARCA2$^{BD}$:PROTAC 2:VHL (PDB ID: 6HAX). See also Supplemental Table 3.

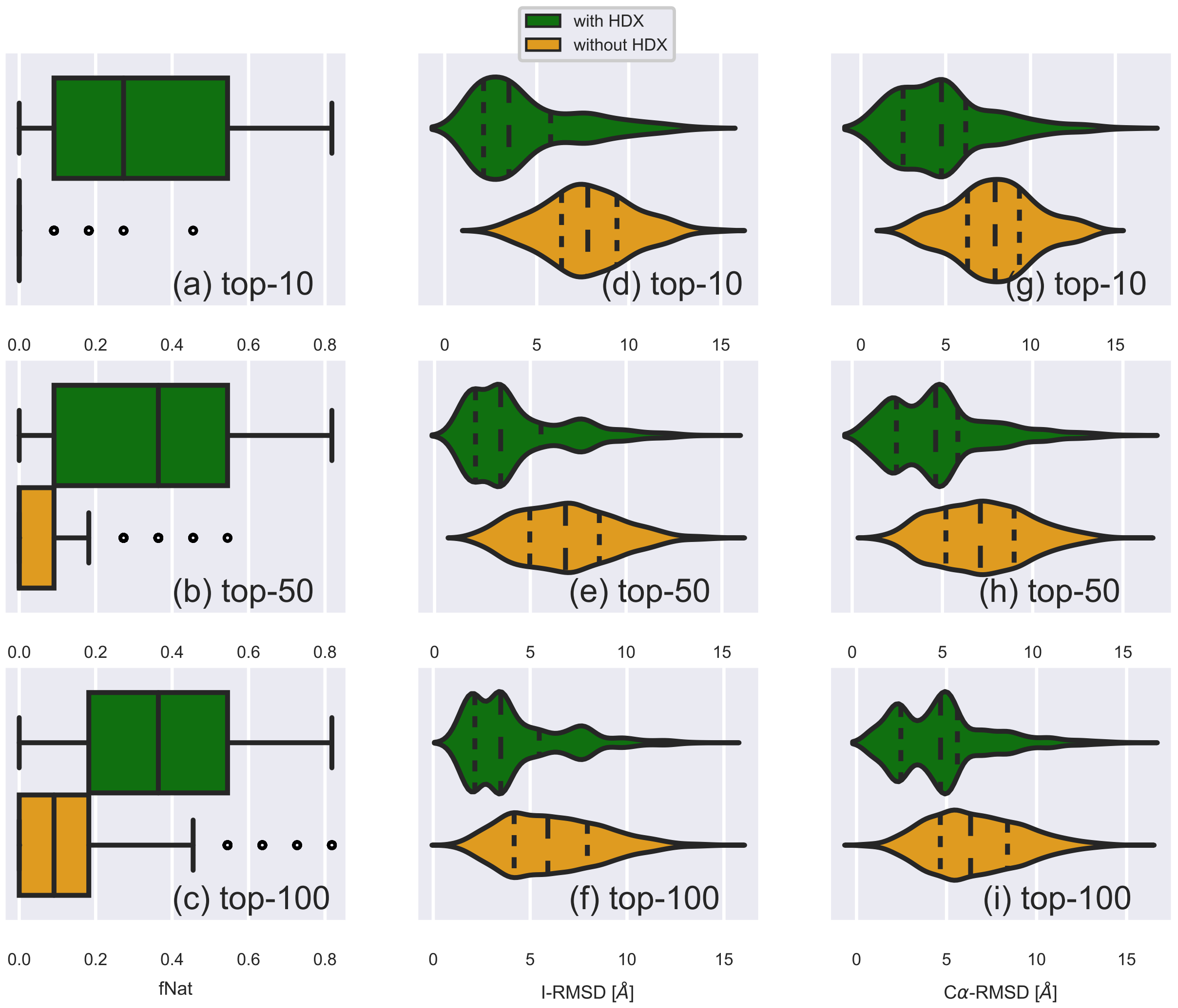


Supplemental Fig. 6: Values of the fNat (**a, b, c**) and distributions of the I-RMSDs (**d, e, f**) and $C\alpha$-RMSDs (**g, h, i**) calculated over 35 independent runs for the top-10 (**a, d, g**), the top-50 (**b, e, h**), and the top-100 (**c, f, i**) docking predictions with (green) and without (orange) HDX-derived restraints. Individual data points are shown as circles. The median and first and third quartiles are shown as vertical solid lines in (**a-c**) and as long and short-dashed lines in (**d-i**). Results presented for SMARCA2$^{BD}$:PROTAC 2:VHL (PDB ID: 6HAX)

Supplemental Fig. 7 shows examples of the docking structures that fall within different categories of quality of predictions. The structures of SMARCA2$^{BD}$ are aligned with the corresponding crystal structure (PDB ID: 6HAX) (salmon) and different predicted poses of the VHL are depicted in purple. The reference pose of the VHL from the same crystal structure is depicted in orange.

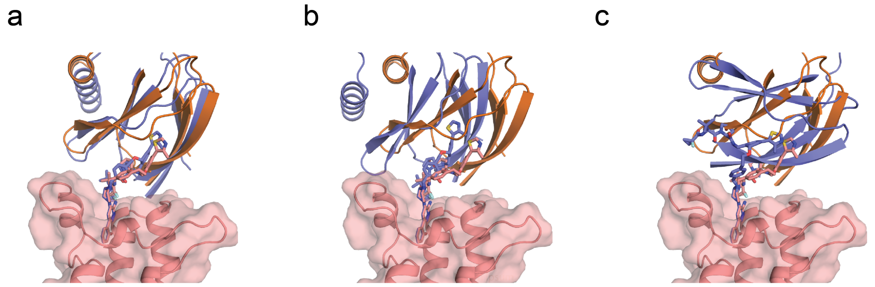


Supplemental Fig. 7: Illustration of the docking-based predicted ternary structures superimposed onto the corresponding co-crystallized structure (PDB ID: 6HAX). (**a**) High quality prediction; (**b**) Medium quality prediction; (**c**) Incorrect prediction. SMARCA2$^{BD}$ is shown in salmon cartoon and surface representation, the VHL pose from the crystal structure is shown in orange cartoon representation, and the predicted docking pose of VHL is shown in purple cartoon representation.

Supplemental Table 3: Distributions of predicted docking structures over the CAPRI quality categories (PDB ID: 6HAX). The 35 docking runs have been performed both with and without HDX-derived restraints to assure reproducibility of the results. The classification of predictions into the quality categories was done with the conventional CAPRI criteria. In the columns: *mean* is the mean number of models in a given category estimated over the 35 runs; *CI* is the Confidence Interval, i.e. the range within which the *mean* value will be found with the probability 0.95.

| | | High | | Medium | | Acceptable | | Incorrect | |
|---|---|---|---|---|---|---|---|---|---|
| | | mean | CI | mean | CI | mean | CI | mean | CI |
| Top-10 | no HDX | 0.0 | [0.0, 0.0] | 0.0 | [0.0, 0.1] | 0.3 | [0.1, 0.5] | 8.6 | [8.4, 8.8] |
| | with HDX | 0.9 | [0.6, 1.3] | 2.2 | [1.8, 2.6] | 2.7 | [2.3, 3.1] | 3.2 | [2.7, 3.7] |
| Top-50 | no HDX | 0.0 | [0.0, 0.0] | 0.4 | [0.2, 0.6] | 6.0 | [5.4, 6.8] | 43 | [42, 43] |
| | with HDX | 3.8 | [3.1, 4.6] | 13 | [12, 14] | 16 | [15, 17] | 16 | [15, 17] |
| Top-100 | no HDX | 0.4 | [0.1, 0.8] | 3.5 | [2.8, 4.2] | 21 | [19, 22] | 74 | [73, 76] |
| | with HDX | 6.8 | [6.0, 7.7] | 30 | [28, 31] | 30 | [29, 32] | 32 | [31, 34] |

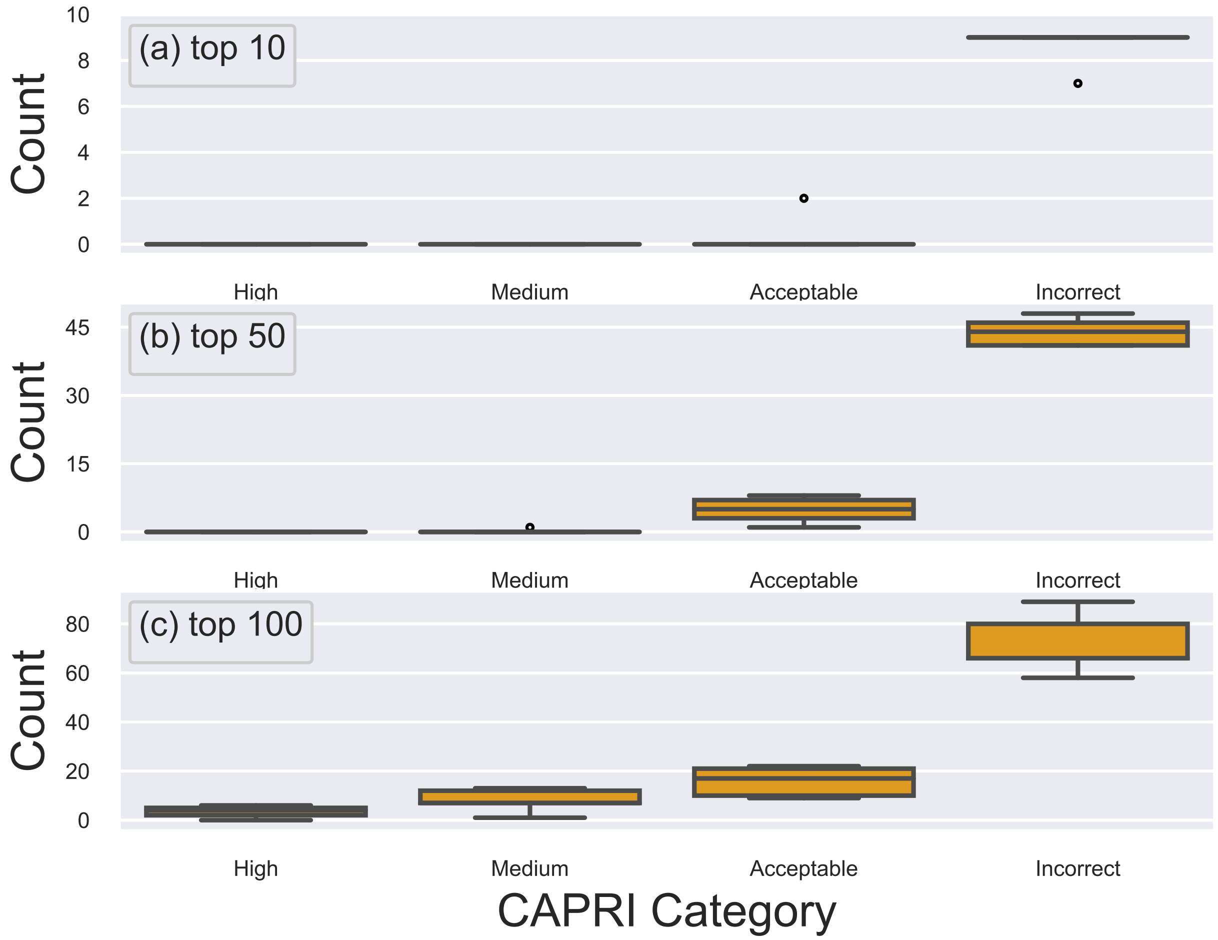


Supplemental Fig. 8: Distributions of the Top-N docking predictions into the CAPRI quality categories High, Medium, Acceptable and Incorrect. 5 independent docking runs have been performed and mean values have been calculated for (**a**) the top-10, (**b**) the top-50, and (**c**) the top-100 docking predictions. Individual data points are shown as circles. The median, first and third quartiles are shown as horizontal lines. Results presented for SMARCA2$^{BD}$:PROTAC 1:VHL (PDB ID: 6HAY) *no HDX-MS-derived restraints were used in docking*

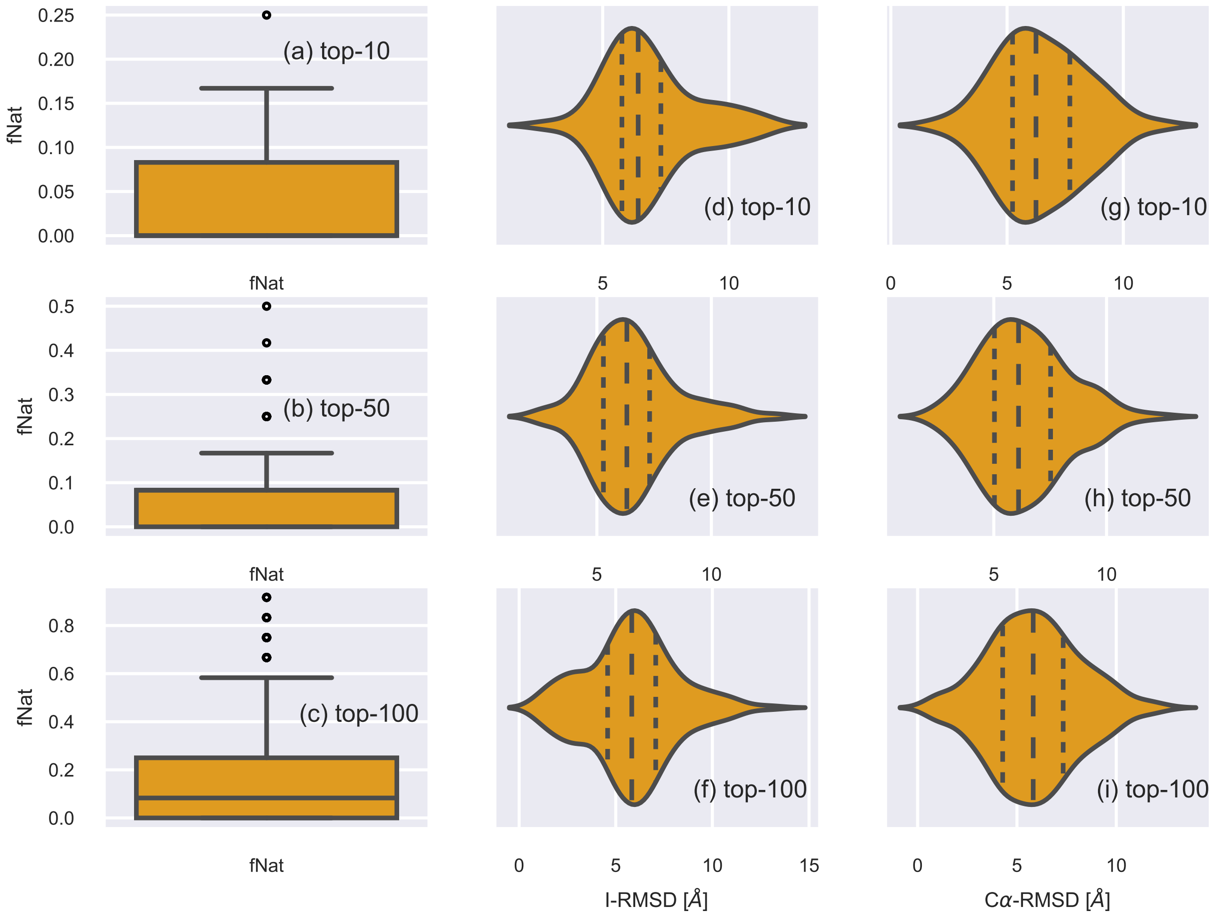


Supplemental Fig. 9: Values of the fNat (**a,b,c**) and distributions of the I-RMSD (**d, e, f**) and $C\alpha - RMSD$ (**g, h, i**) calculated over 5 independent runs for the top-10 (**a, d, g**), the top-50 (**b, e, h**), and the top-100 (**c, f, i**) docking predictions. The median and first and third quartiles are shown as vertical solid lines in (**a - c**) and as long and short-dashed lines in (**d - i**). Results presented for SMARCA2$^{BD}$:PROTAC 1:VHL (PDB ID: 6HAY) *no HDX-MS-derived restraints were used in docking*

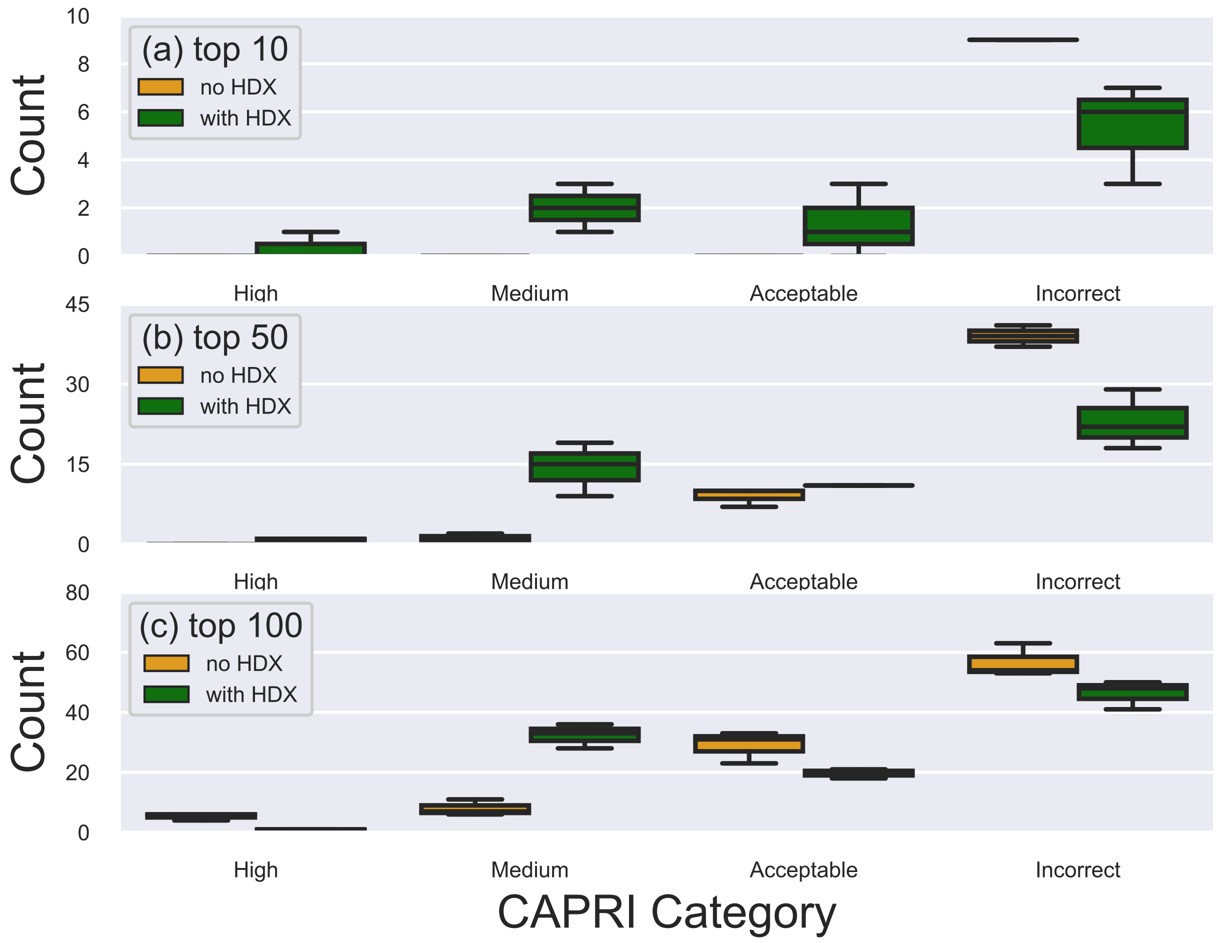


Supplemental Fig. 10: Distributions of the Top-N docking predictions into the CAPRI quality categories High, Medium, Acceptable, and Incorrect. 3 independent docking runs have been performed and mean values have been calculated for (**a**) the top-10, (**b**) the top-50, and (**c**) the top-100 docking predictions with (green) and without (orange) HDX-derived restraints. Individual data points are shown as circles. The median, first and third quartiles are shown as horizontal lines. Results presented for SMARCA2$^{BD}$-iso1:ACBI1:VHL.

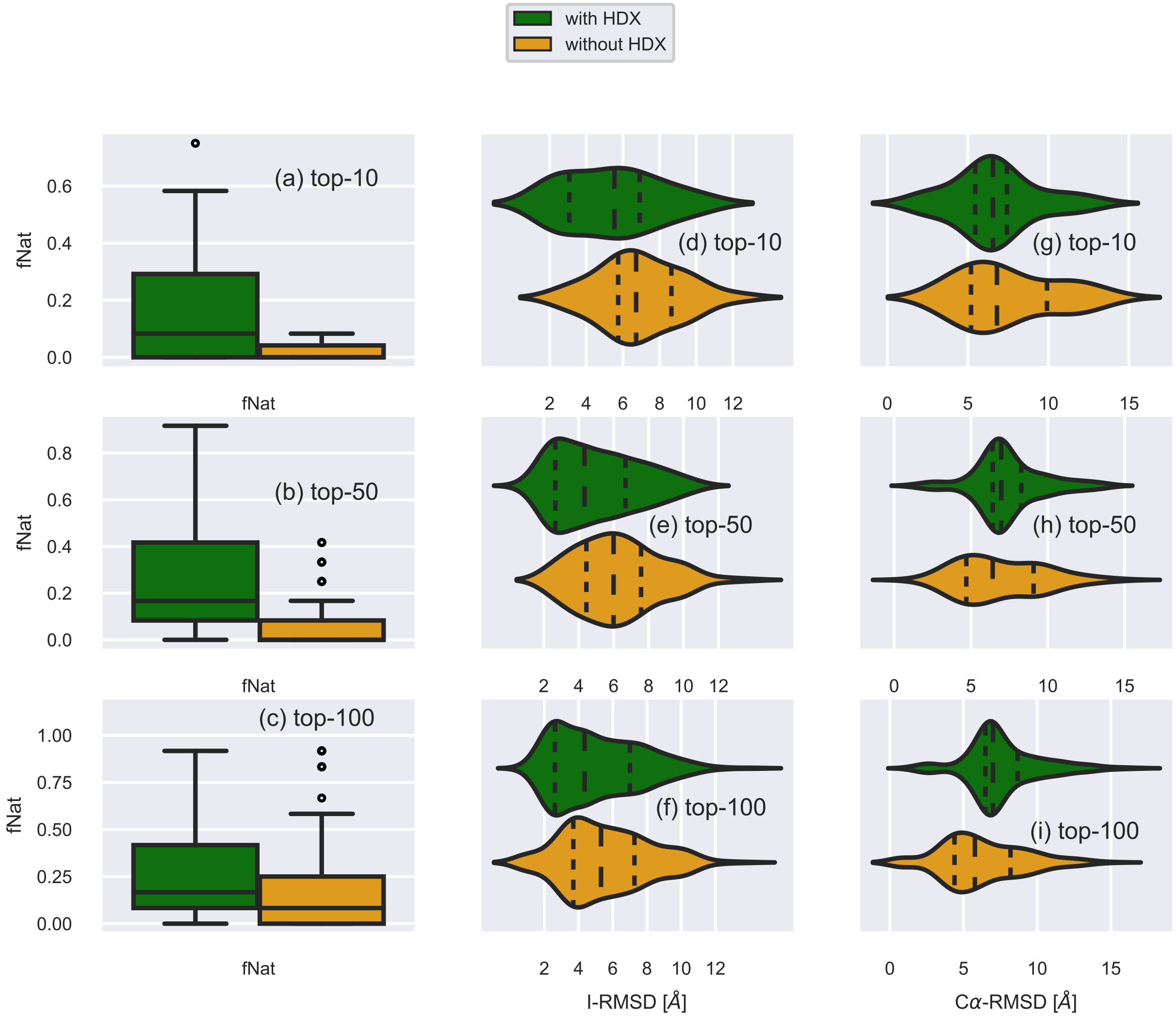


Supplemental Fig. 11: Values of the fNat (**a,b,c**) and distributions of the I-RMSDs (**d, e, f**) and $C\alpha - RMSDs$ (**g, h, i**) calculated over 3 independent runs for the top-10 (**a, d, g**), the top-50 (**b, e, h**) and the top-100 (**c, f, i**) docking predictions with (green) and without (orange) HDX-derived restraints. Individual data points are shown as circles. The median and first and third quartiles are shown as vertical solid lines in (**a - c**) and as long and short-dashed lines in (**d - i**). Results presented for SMARCA2$^{BD}$-iso1:ACBI1:VHL

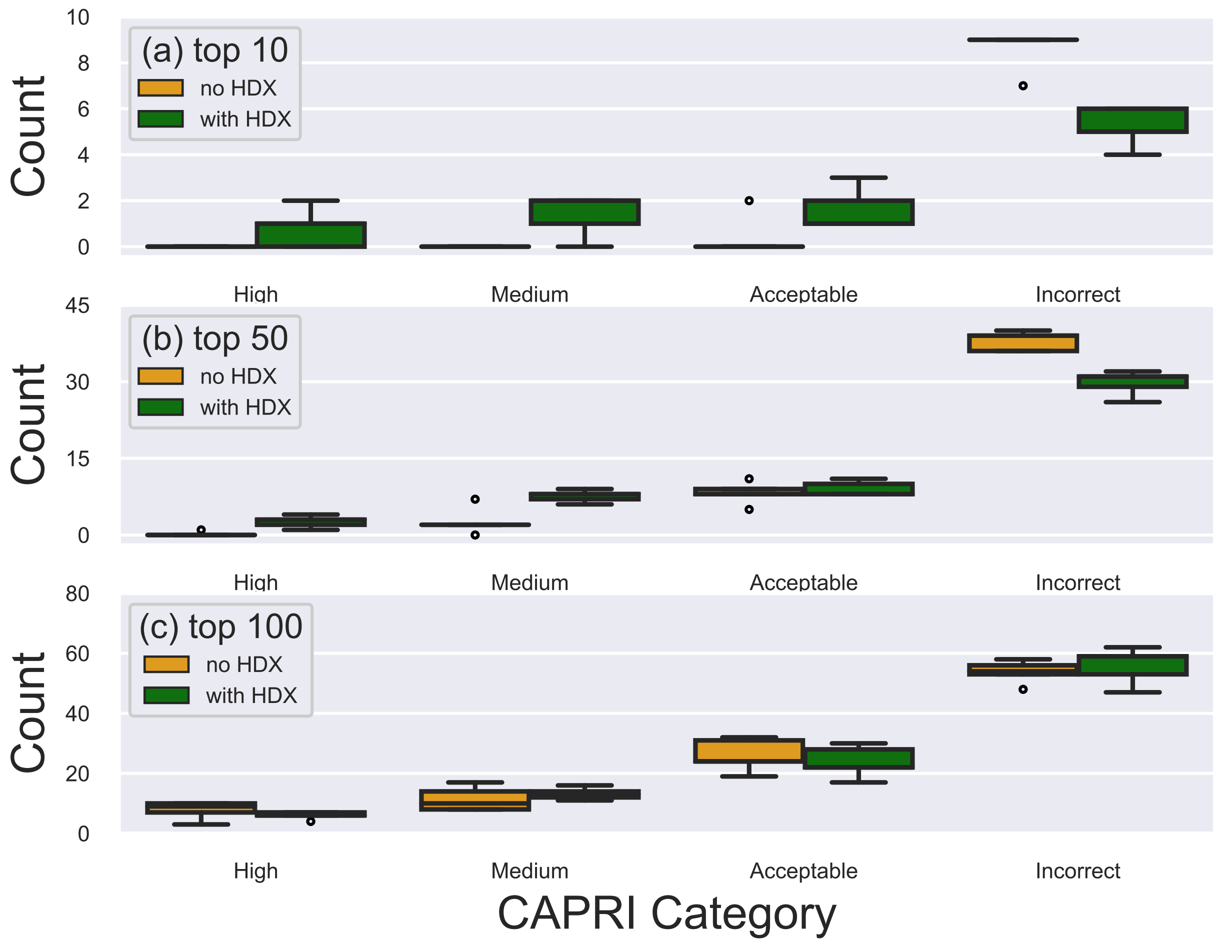


Supplemental Fig. 12: Distributions of the Top-N docking predictions into the CAPRI quality categories High, Medium, Acceptable and Incorrect. 5 Independent docking runs have been performed and mean values have been calculated for (**a**) the top-10, (**b**) the top-50, and (**c**) the top-100 docking predictions with (green) and without (orange) HDX-derived restraints. Individual data points are shown as circles. The median, first and third quartiles are shown as horizontal lines. Results presented for SMARCA2$^{BD}$-iso2:ACBI1:VHL

.

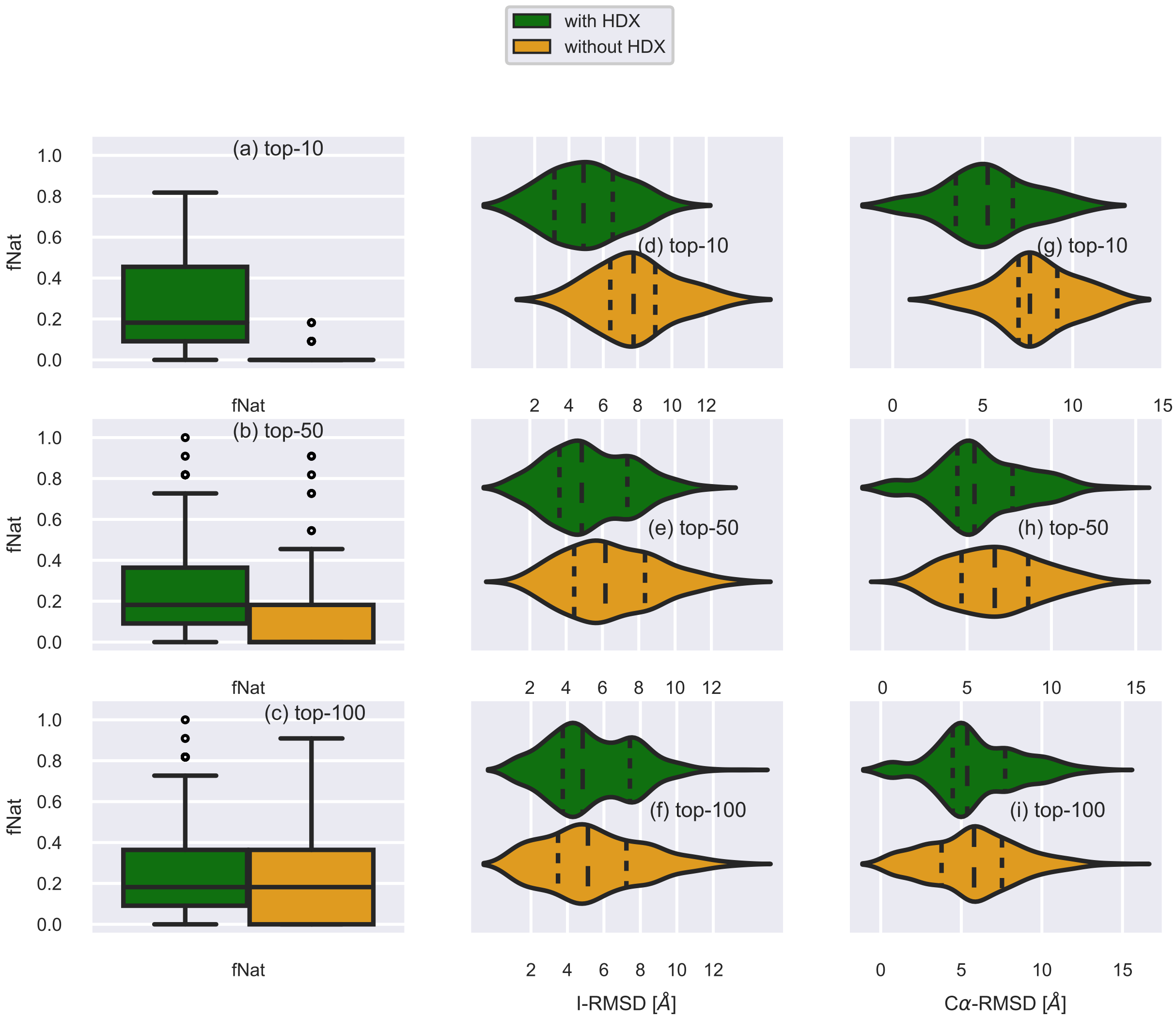


Supplemental Fig. 13: Values of the fNat (**a,b,c**) and distributions of the I-RMSDs (**d, e, f**) and $C\alpha - RMSDs$ (**g, h, i**) calculated over 5 independent runs for the top-10 (**a, d, g**), the top-50 (**b, e, h**) and the top-100 (**c, f, i**) docking predictions with (green) and without (orange) HDX-derived restraints. Individual data points are shown as circles. The median and first and third quartiles are shown as vertical solid lines in (**a - c**) and a long and short-dashed lines in (**d - i**). Results presented for SMARCA2$^{BD}$-iso2:ACBI1:VHL

161 **SAXS experiments.**

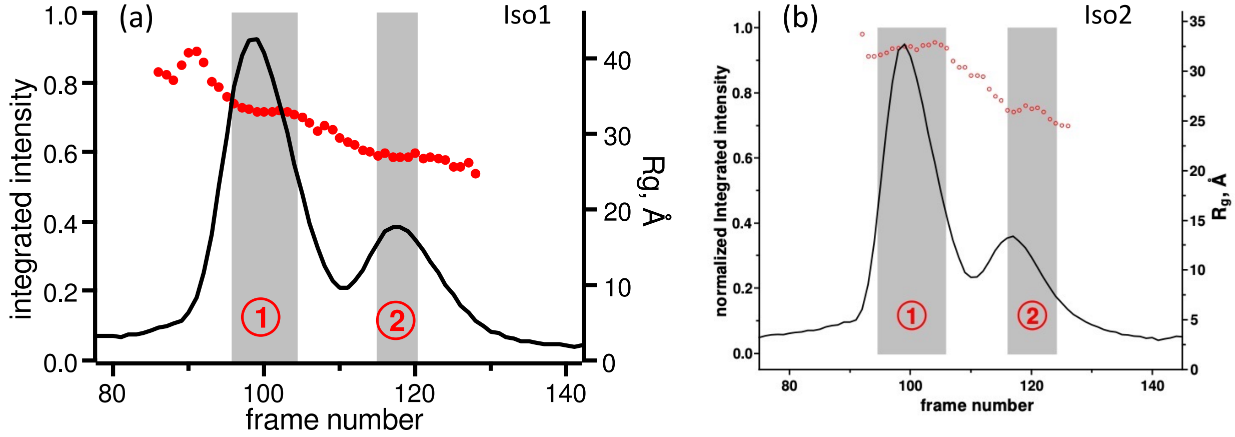


Supplemental Fig. 14: The normalized integrated intensity for each SAXS frame recorded during the elution peak 1 (full complex) and peak 2 (uncomplexed or binary) of (a) iso1-SMARCA2$^{BD}$:ACBI1:VCB and (b) iso2-SMARCA2$^{BD}$:ACBI1:VCB.

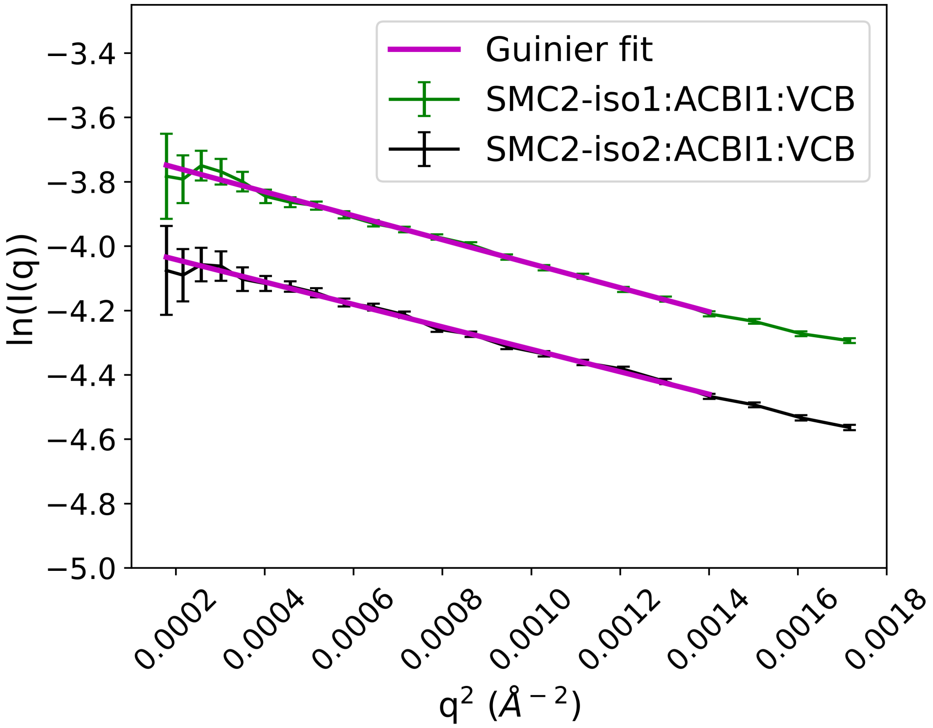


Supplemental Fig. 15: Guinier analysis. $R_g$ of complex is determined using Guinier approximation at low $q$-values (Eq. 1 in the main text). The magenta solid lines are the Guinier fits to the SAXS data.

#### HREMD simulations.

The Hamiltonian replica-exchange molecular dynamics (HREMD) simulation is a computationally efficient method to enhance the configurational sampling of a biomolecular system.[46,47] Recent studies revealed that HREMD-generated structures of intrinsically disordered proteins[48,49] and G protein-coupled receptors[50,51] achieved an excellent agreement with experiments. Here we implemented HREMD (specifically replica-exchange with solute tempering, REST2)[46,47] method using the software package GROMACS (v2018.8)[52–54] patched with PLUMED (v2.5.4)[55–59] to explore the conformational free energy landscape of the SMARCA2$^{BD}$:VHL degrader ternary complexes. Using this approach, we ran several parallel simulations (replicas) with scaled Hamiltonians by dividing the system into two regions as "hot" and "cold". The Hamiltonian of the "hot" region (viz., SMARCA2$^{BD}$, VHL, and degrader atoms) was scaled by a factor $\lambda$ (Supplemental Eq. 1) in higher rank replicas. Whereas the Hamiltonian of the "cold" region (solvent) was unaltered for all the replicas. Specifically, the Lennard-Jones parameter ($\epsilon$), the dihedral term, and the charge of the atoms in the "hot" region of the $i^{th}$ replica are scaled by $\lambda_i$, $\lambda_i$, and $\sqrt{\lambda_i}$ respectively given by Eq. 1,

$$\lambda_i = \frac{T_0}{T_i} = exp\Bigg( - \frac{i}{(n-1)} ln\Big(\frac{T_{max}}{T_0}\Big)\Bigg) \tag{1}$$

where $n$ is the total number of replicas, and $T_0$, $T_i$ and $T_{max}$ are the effective temperatures of the lowest (unscaled), the $i^{th}$, and the highest rank replicas, respectively. Therefore, only the force field terms that contribute to the energy barriers were scaled.[46] The exchange of coordinates was allowed after every 500 MD steps (1 ps) between the neighboring replicas if the Monte Carlo metropolis criterion was satisfied.[46] In principle, this scheme helps to rapidly sample the conformational substates of a "hot" region.

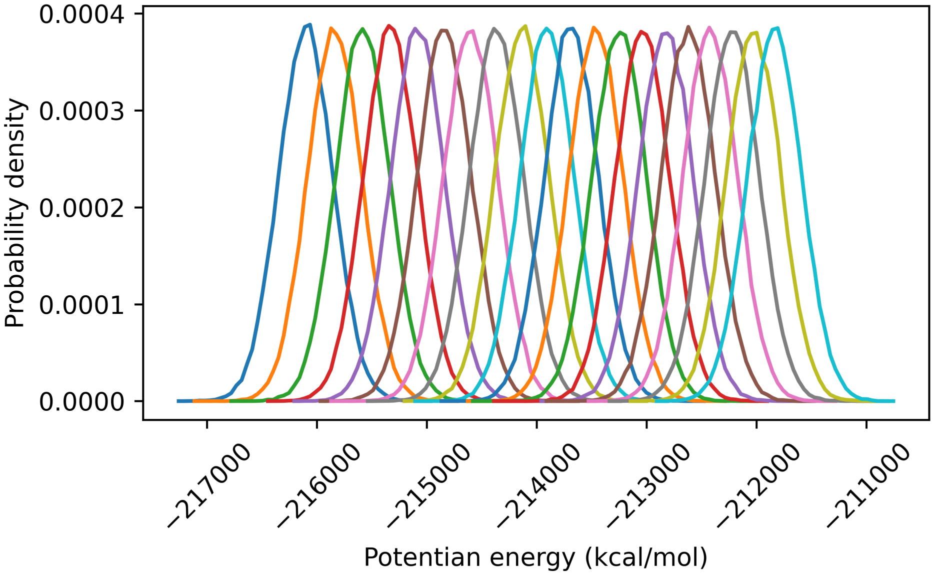


Supplemental Fig. 16: Potential energy of all replicas from HREMD simulation of **Sys7** (see Supplemental Table **??**). Left to right: rank 0 to rank 19. A good overlap between adjacent replicas suggests a sufficient number of replicas were employed and also confirms that no phase transition took place during the HREMD simulation.

Supplemental Table 4: Details of HREMD simulations. Protein complexes, number of atoms in a simulation box, number of replicas used and the aggregate length of the simulations are listed.

| **ID** | **Complex** | **# of atoms** | **# of replicas** | **Aggregate length ($\mu$s)** |
|---|---|---|---|---|
| Sys1 | iso1-SMARCA2$^{BD}$:ACBI1:VHL | 116,254 | 20 | 10 |
| Sys2 | iso1-SMARCA2$^{BD}$:ACBI1:VCB | 220,573 | 24 | 12 |
| Sys3 | iso2-SMARCA2$^{BD}$:ACBI1:VHL | 117,256 | 20 | 10 |
| Sys4 | iso2-SMARCA2$^{BD}$:ACBI1:VCB | 234,724 | 24 | 12 |
| Sys5 | iso2-SMARCA2$^{BD}$:PROTAC 1:VHL | 137,347 | 20 | 10 |
| Sys6 | iso1-SMARCA2$^{BD}$:PROTAC 2:VHL | 69,696 | 20 | 10 |
| Sys7 | iso2-SMARCA2$^{BD}$:PROTAC 2:VHL | 68,820 | 20 | 10 |
| Sys8 | iso2-SMARCA2$^{BD}$:PROTAC 2:VCB | 119,082 | 24 | 12 |

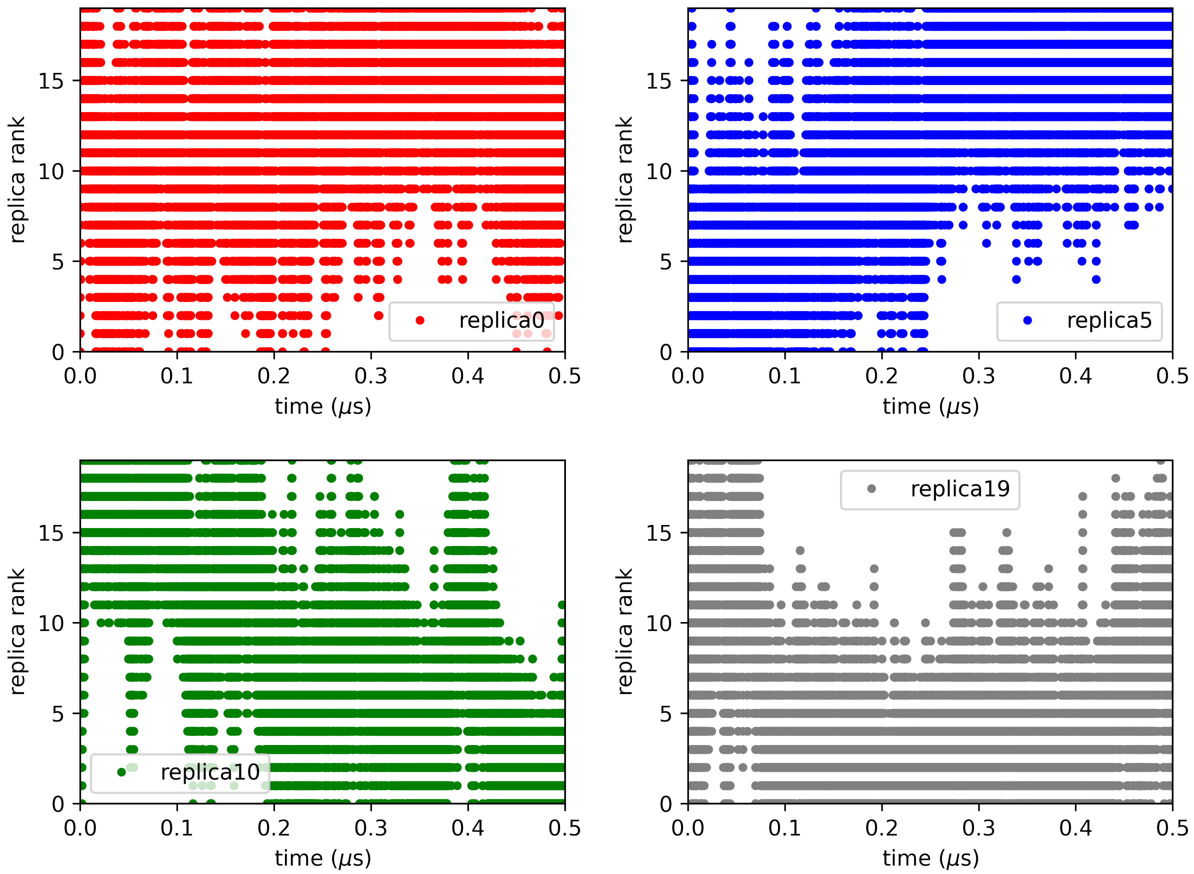


Supplemental Fig. 17: Effective temperature trajectories of replicas ranked 0 (red), 5 (blue), 10 (green) and 19 (grey) from HREMD simulation of **Sys7** (see Supplemental Table **??**).

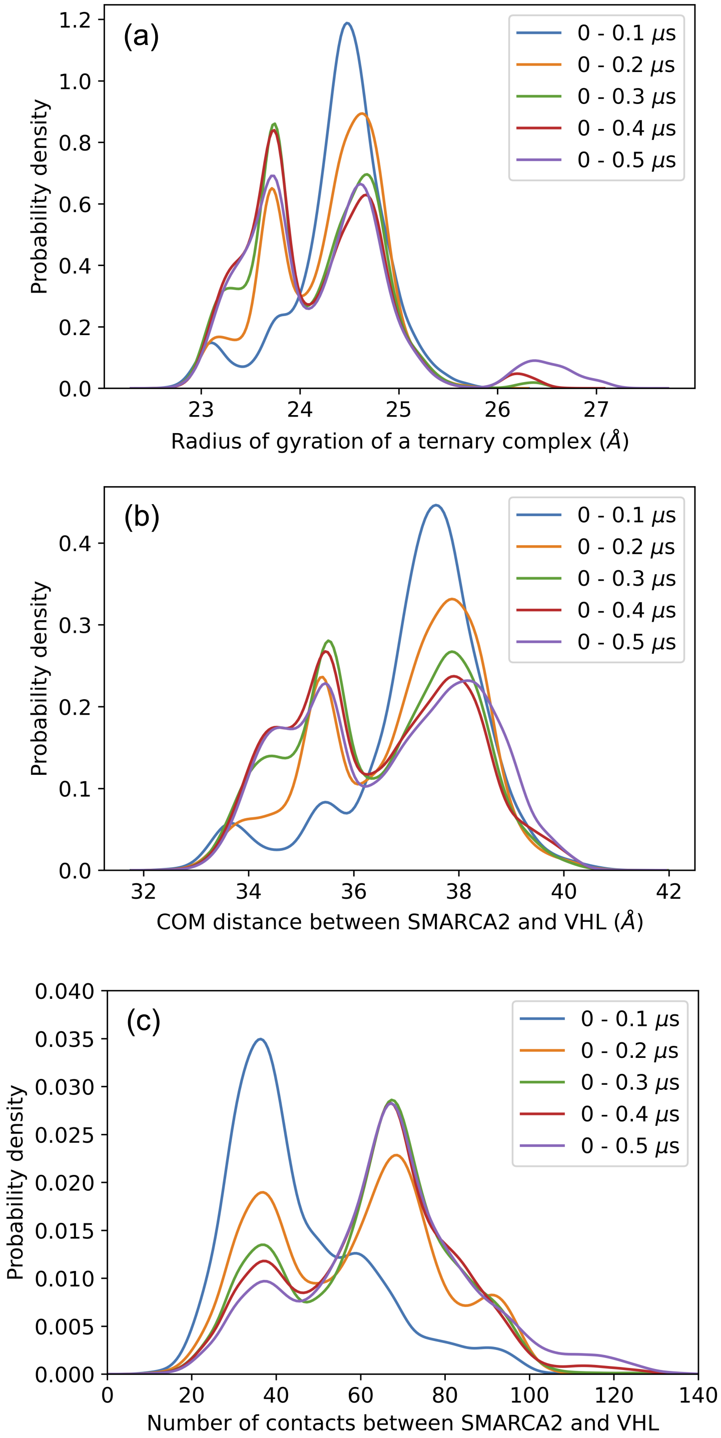


Supplemental Fig. 18: HREMD convergence test. The distribution of (a) radius of gyration of a ternary complex, (b) center of mass (COM) distance between SMARCA2$^{BD}$ and VHL, and (c) heavy atom contacts within 5 Å between SMARCA2$^{BD}$ and VHL are plotted with cumulative length of HREMD simulation for **Sys7** (see Supplemental Table **??**).

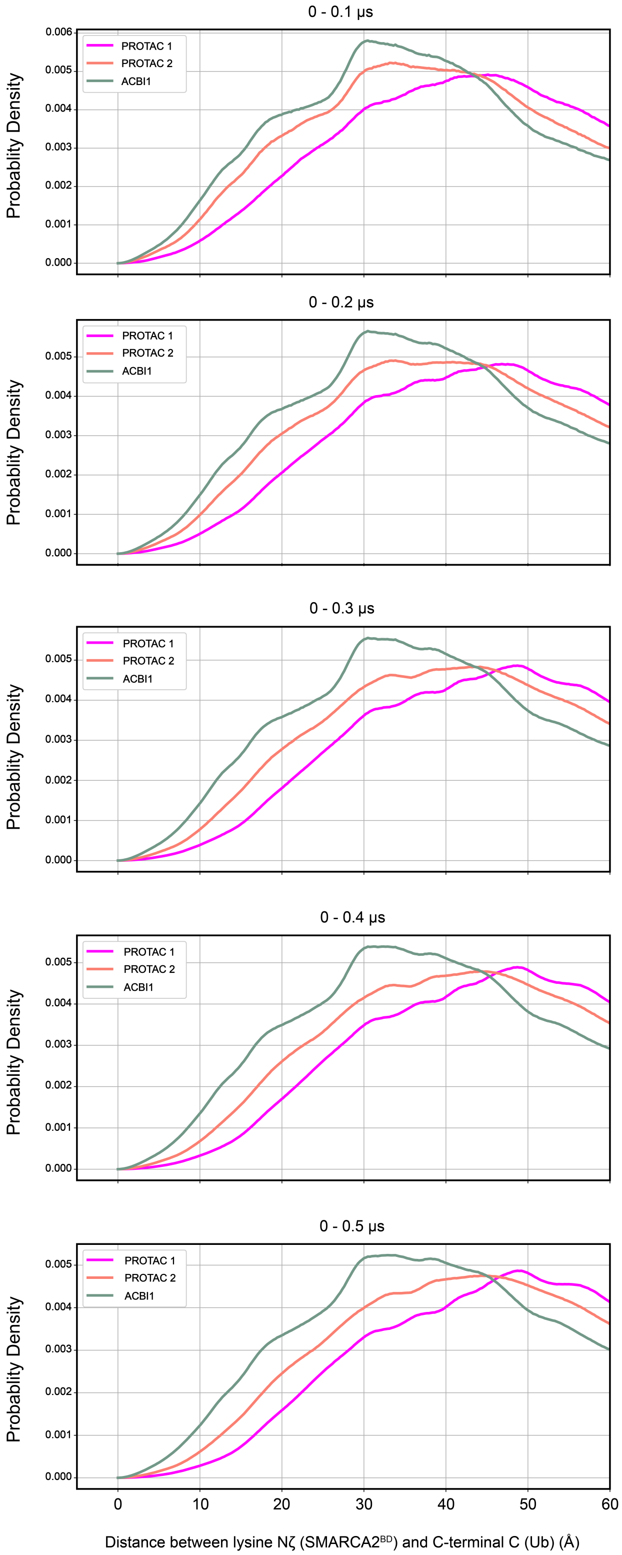


Supplemental Fig. 19: HREMD convergence test. Distance of Lys residues (side-chain nitrogen atom) from SMARCA2 to the C-terminus glycine C atom of ubiquitin for three different degraders (PROTAC 1, PROTAC 2 and ACBI1) plotted with simulation lengths to confirm the convergence of the result.

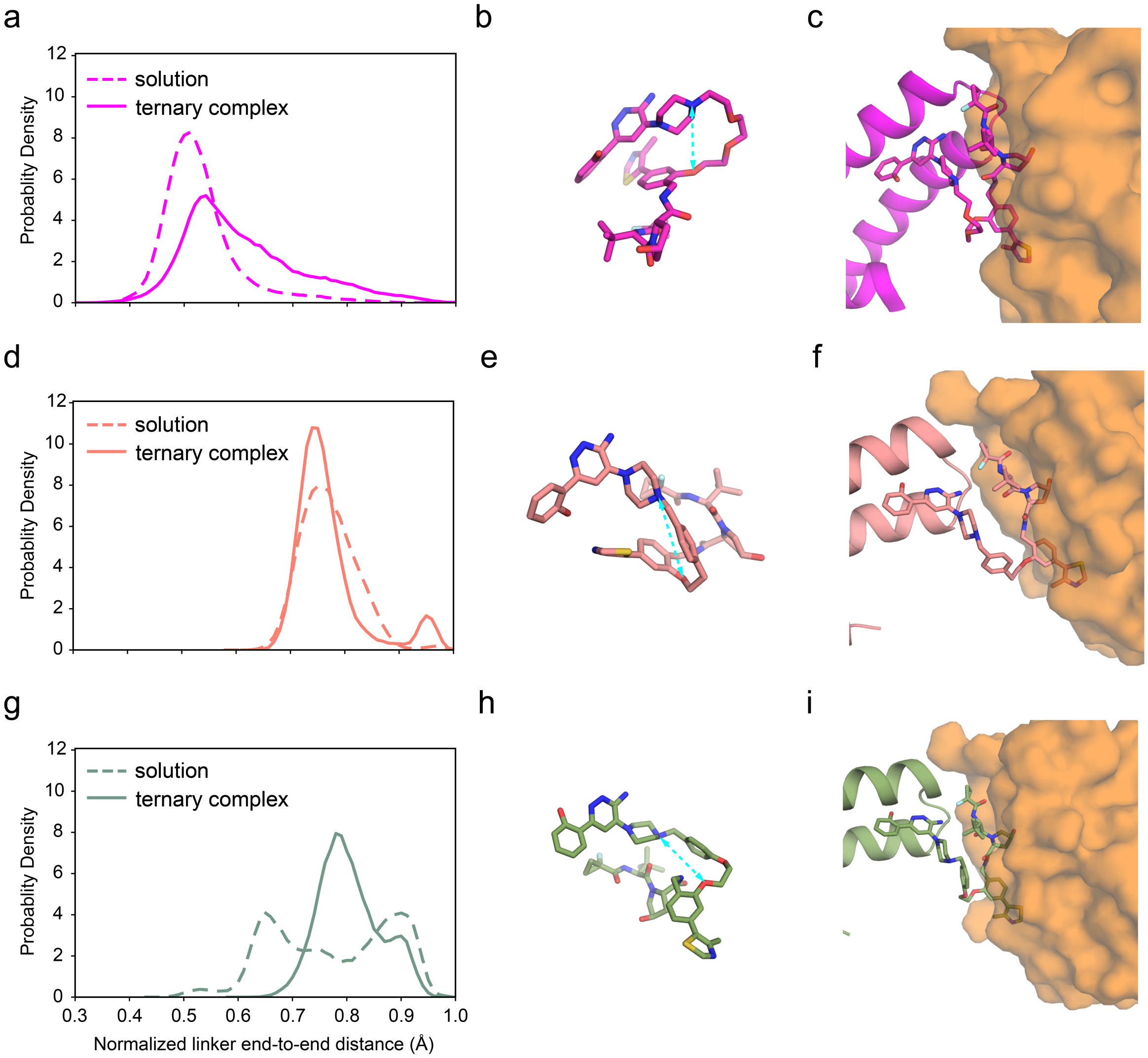


Supplemental Fig. 20: Degrader compatibility in a ternary complex. Comparison of linker end-to-end distance normalized by the number of linker backbone atoms in solution (dashed line) vs. in ternary complex (solid line) for (a) PROTAC 1 (d) PROTAC 2, and (g) ACBI1. Snapshots of conformation of PROTAC 1 (magenta, panels b and c), PROTAC 2 (salmon, panels e and f) and ACBI1 (green, panels h and i) in solution and in ternary complex are shown by stick representation. The cyan dashed line with arrows indicates the linker end-to-end distance.

184 **Long-timescale conformational sampling.**

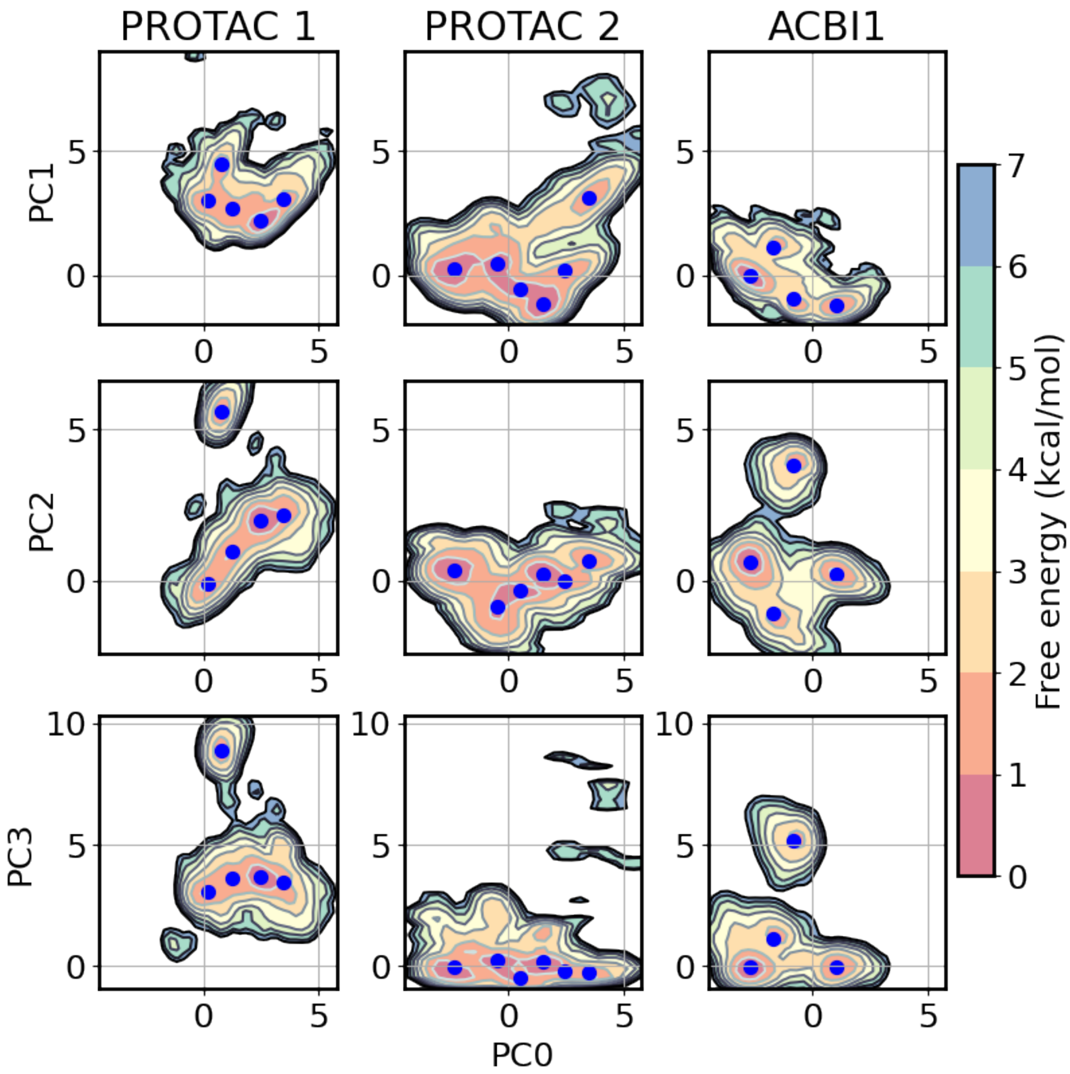


Supplemental Fig. 21: Free energy landscapes determined from PCA projections of iso2-SMARCA2$^{BD}$ bound to VHL via PROTAC 1 (left), PROTAC 2 (middle), and ACBI1 (right). Blue points indicate $k$-means centroids, which show an approximate correspondence with local minima of the free energy surface. To facilitate comparison, the landscapes are here shown projected onto the same PCA space, determined from interface distances of the PROTAC 2 system.

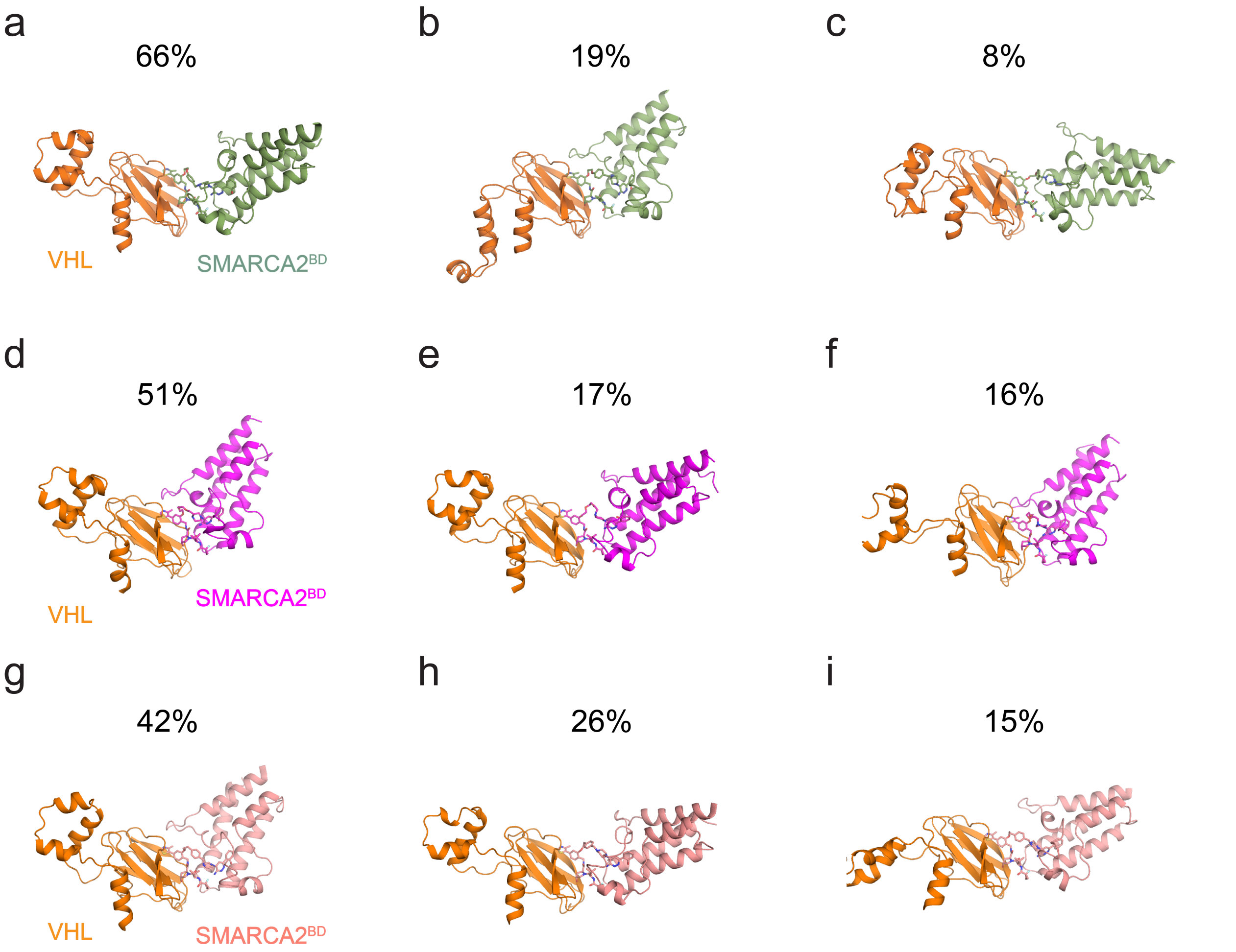


Supplemental Fig. 22: Cluster centroids from the three highest populated structures of iso2-SMARCA2$^{BD}$ bound to VHL via (a-c) ACBI1, (d-f) PROTAC 1, and (g-i) PROTAC 2, along with their populations. Less populated structures are omitted.

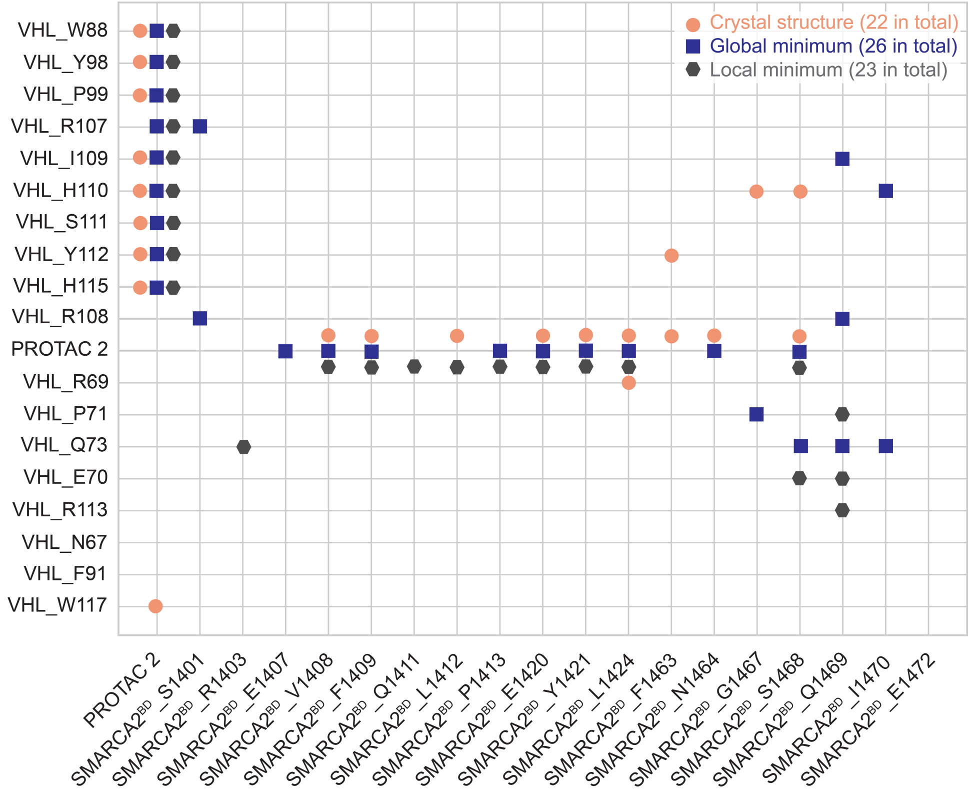


Supplemental Fig. 23: Contact maps for the iso2-SMARCA2$^{BD}$:PROTAC 2:VHL ternary complex crystal structure (PDB ID: 6HAX; orange circles) and its global minimum (blue squares) and metastable (gray hexagons) states identified by our MSM.

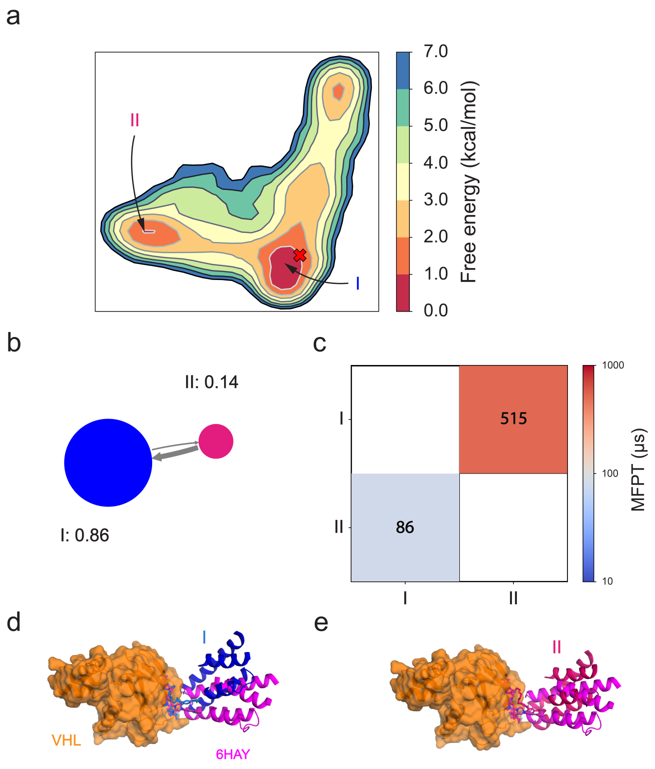


Supplemental Fig. 24: Markov-state Model for the iso2-SMARCA2$^{BD}$:PROTAC 1:VHL system. **a** Conformational free energy landscape as a function of the first two tICA features. The crystal structure (PDB ID: 6HAY) is shown as a red X. **b** Network diagram of the coarse-grained MSM calculated with a lag time of 100 ns, with the stationary probabilities associated with each state indicated. **c** Mean-first passage times (MFPTs) to transition from one state to another in the coarse-grained MSM. Numbers indicate predicted MFPTs in $\mu s$. **d-e** Comparison of the crystal structure (magenta) with the lowest free energy state (skyblue) and the metastable state (cyan) predicted by the MSM.

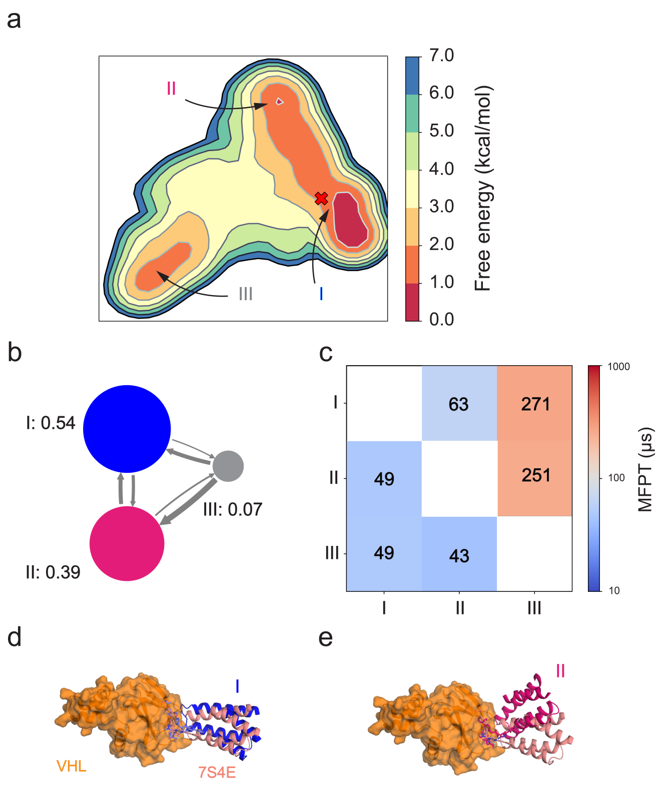


Supplemental Fig. 25: Markov-state Model for the iso2-SMARCA2$^{BD}$:ACBI1:VHL system. **a** Conformational free energy landscape as a function of the first two tICA features. The crystal structure (PDB ID: 7S4E) is shown as a red X. **b** Network diagram of the coarse-grained MSM calculated with a lag time of 100 ns, with the stationary probabilities associated with each state indicated. **c** Mean-first passage times (MFPTs) to transition from one state to another in the coarse-grained MSM. Numbers indicate predicted MFPTs in $\mu s$. **d-e** Comparison of the crystal structure (salmon) with the lowest free energy state (cyan) and the metastable state (skyblue) predicted by the MSM.

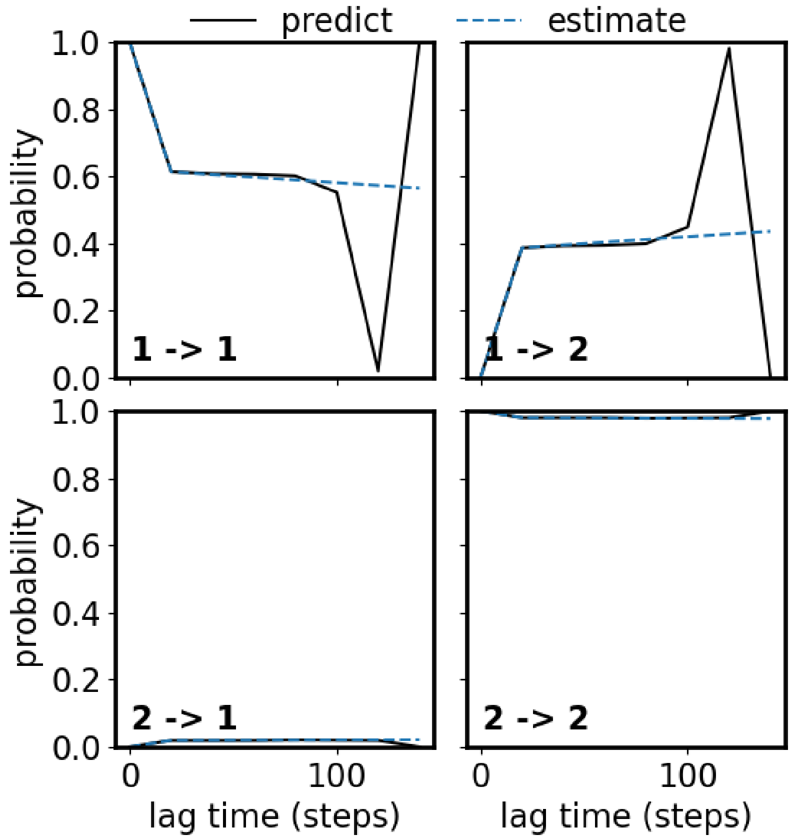


Supplemental Fig. 26: Chapman-Kolmogorov tests for PROTAC 1 MSM.

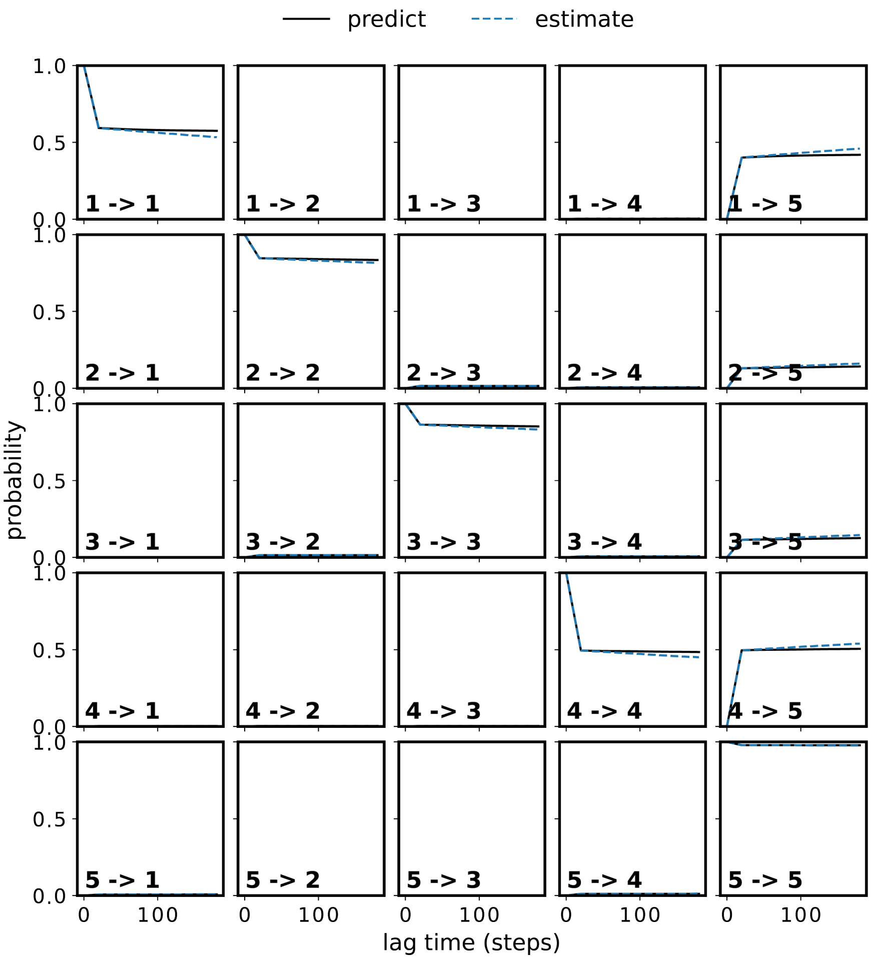


Supplemental Fig. 27: Chapman-Kolmogorov tests for PROTAC 2 MSM.

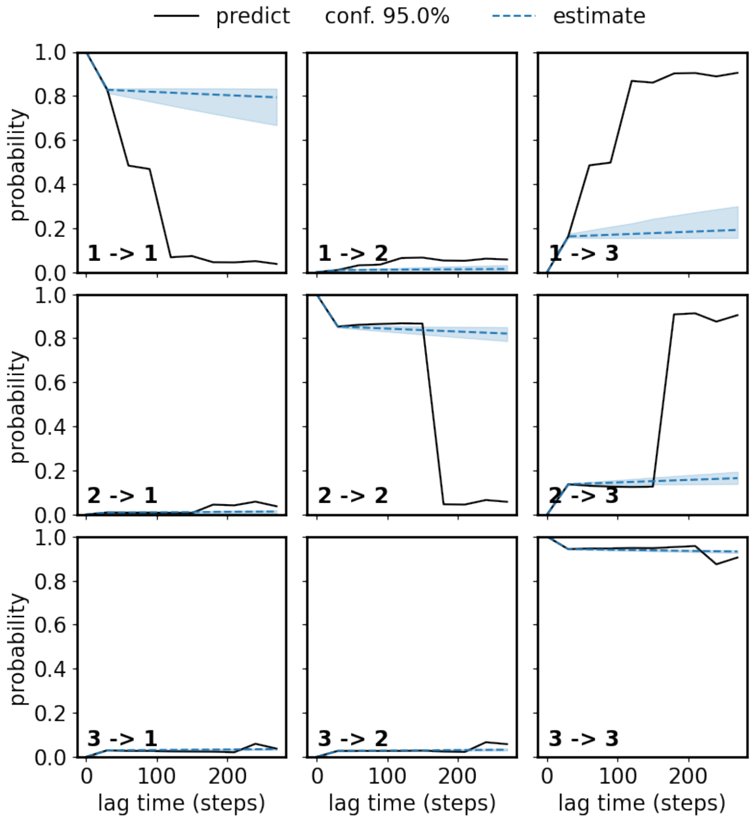


Supplemental Fig. 28: Chapman-Kolmogorov tests for ACBI1 MSM.

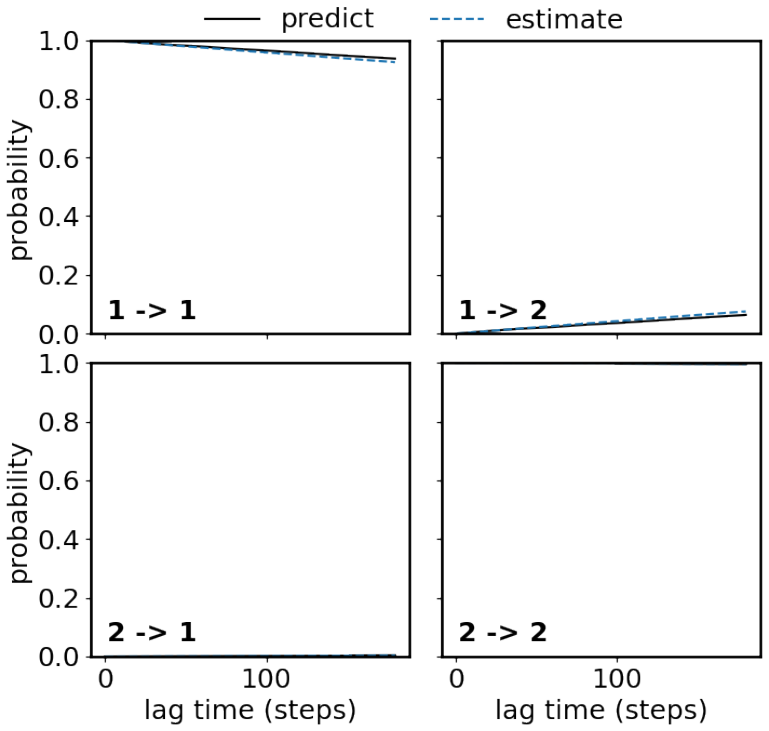


Supplemental Fig. 29: Chapman-Kolmogorov tests for PROTAC 1 HMM.

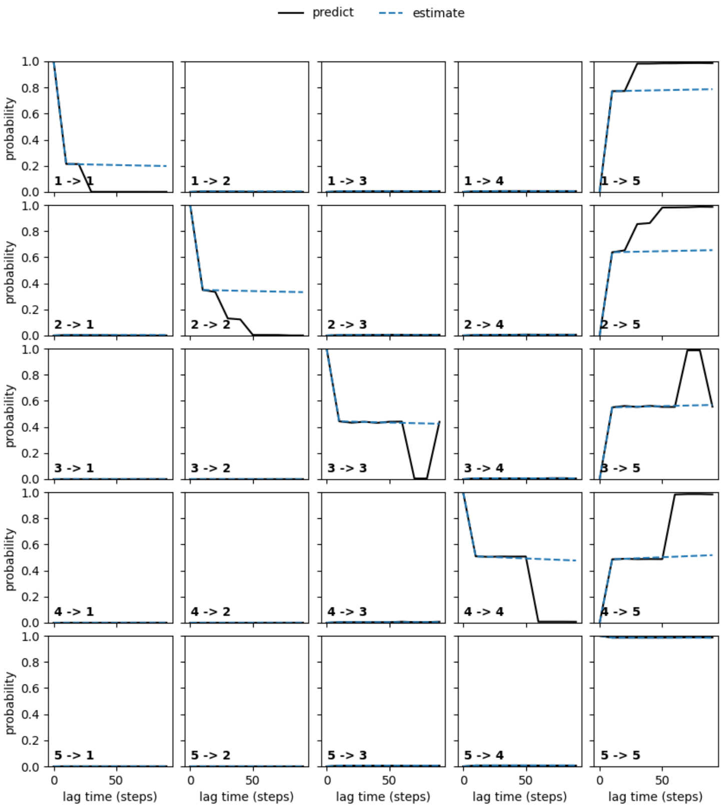


Supplemental Fig. 30: Chapman-Kolmogorov tests for PROTAC 2 HMM.

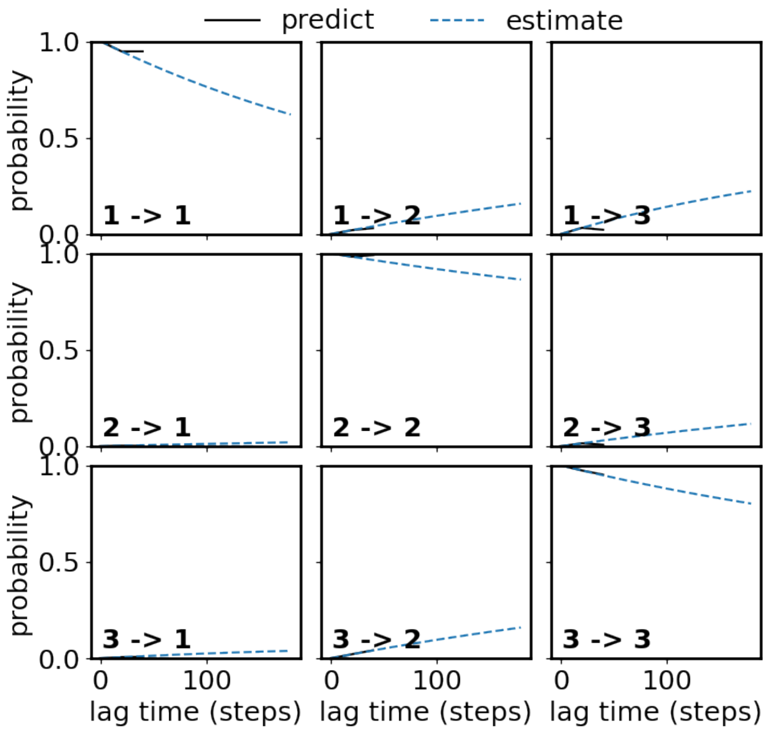


Supplemental Fig. 31: Chapman-Kolmogorov tests for ACBI1 HMM.

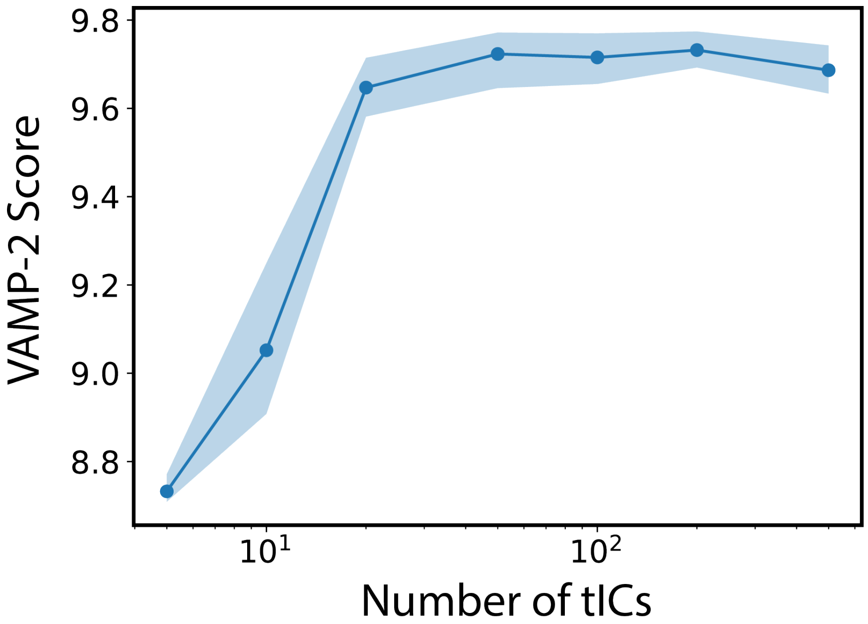


Supplemental Fig. 32: VAMP-2 score as a function of number of tICs used during fitting of our MSM for PROTAC 2. Beyond $\tilde{5}0$ tICs, the VAMP-2 score saturates. Shaded region indicates 90% confidence interval, determined from 5 independently fitted MSMs.

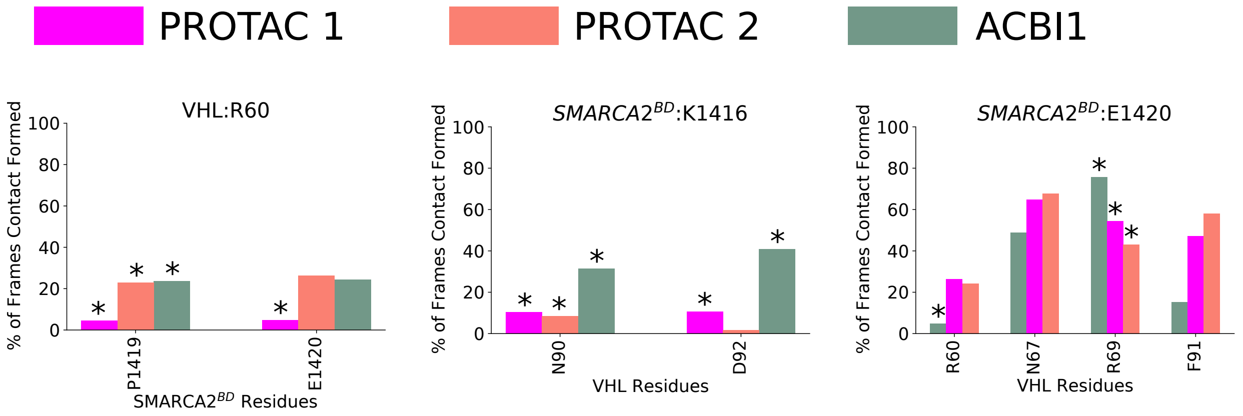


Supplemental Fig. 33: Comparison of the contacts formed between protected charged interface residues iso2-SMARCA2$^{BD}$ or VHL indicated in Section 2.1 (main text) and any residue from the opposite protein. A contact is defined as a minimum distance between heavy atoms of 5 Å Results are shown for contacts that were formed 20% of the time or higher and when the contact was formed for at least one of the systems (PROTAC 1 (magenta), PROTAC 2 (salmon), or ACBI1 (green)). This leaves three distinct charged interface residues, i.e., VHL:R60 (left), SMARCA2$^{BD}$:K1416 (middle), and SMARCA2$^{BD}$:E1420 (right). The asterisks indicate that the specific residue pairs are not observed to form contacts in the corresponding crystal structures.

### Ubiquitination predictions.

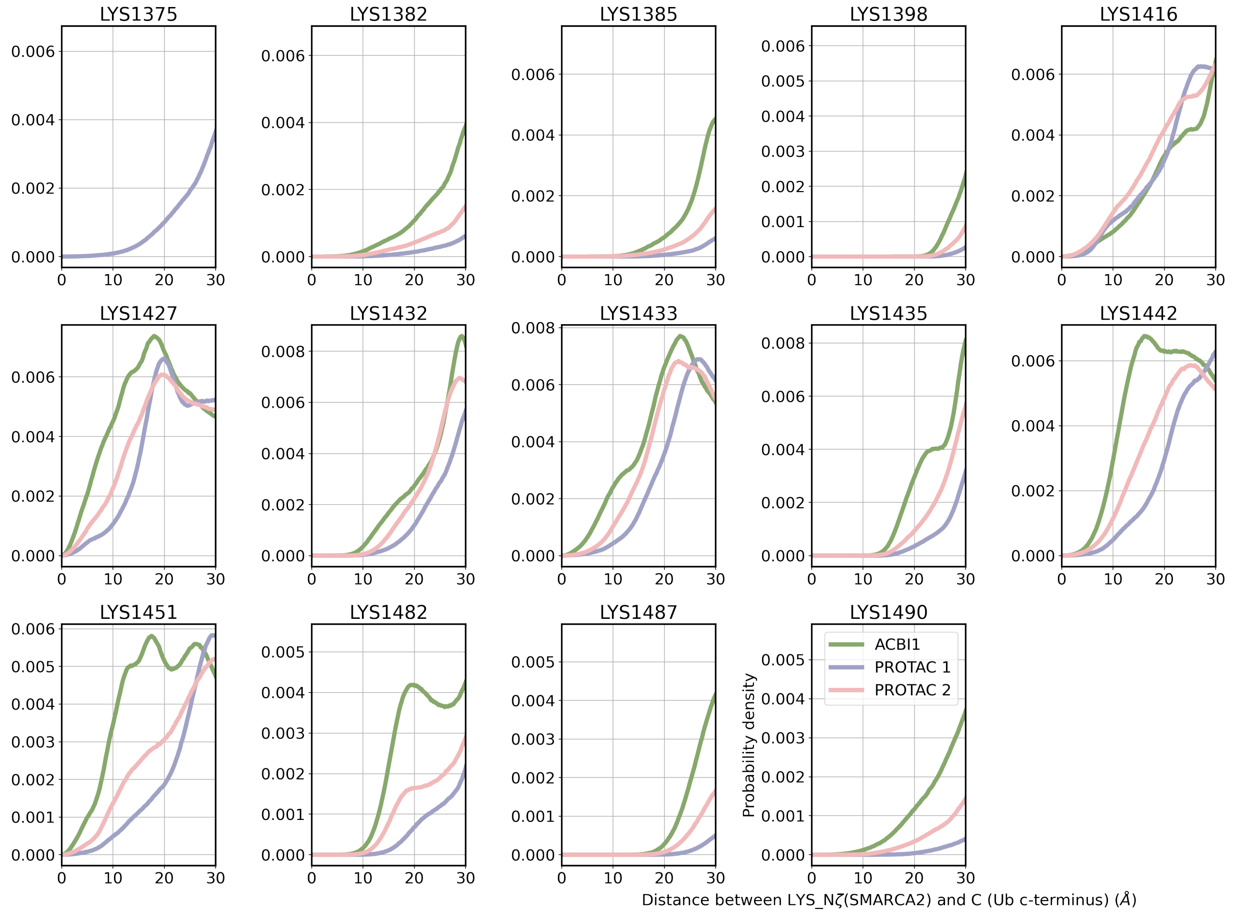


Supplemental Fig. 34: Probability distribution of individual lysine residue distances from ubiquitin based on HREMD simulations of ternary complexes with PROTAC 1, PROTAC 2, or ACBI1. The crystal structures with PROTAC 2 and ACBI1 lack Lys1375 on SMARCA2$^{BD}$, thus there is only a corresponding distribution for PROTAC 1 (top left panel).

### Movies.

Supplemental Table 5: Movies rendered from MD simulation trajectories.

| Movie filename | Download link |
| --- | --- |
| wes_6hax_ternary_complex_formation.mp4 | Click here |
| hremd_6HAX.mp4 | Click here |
| hremd_6HAY.mp4 | Click here |
| hremd_7S4E.mp4 | Click here |
| hremd_iso1_smrc2_acbi1_vhl.mp4 | Click here |
| meta-eabf_CRL_SMARCA2$^{BD}$_ACBI1_VHL_25to40Angs.mp4 | Click here |
| meta-eabf_CRL_SMARCA2$^{BD}$_ACBI1_VHL_25to75Angs.mp4 | Click here |

**HDX-MS experiments.**

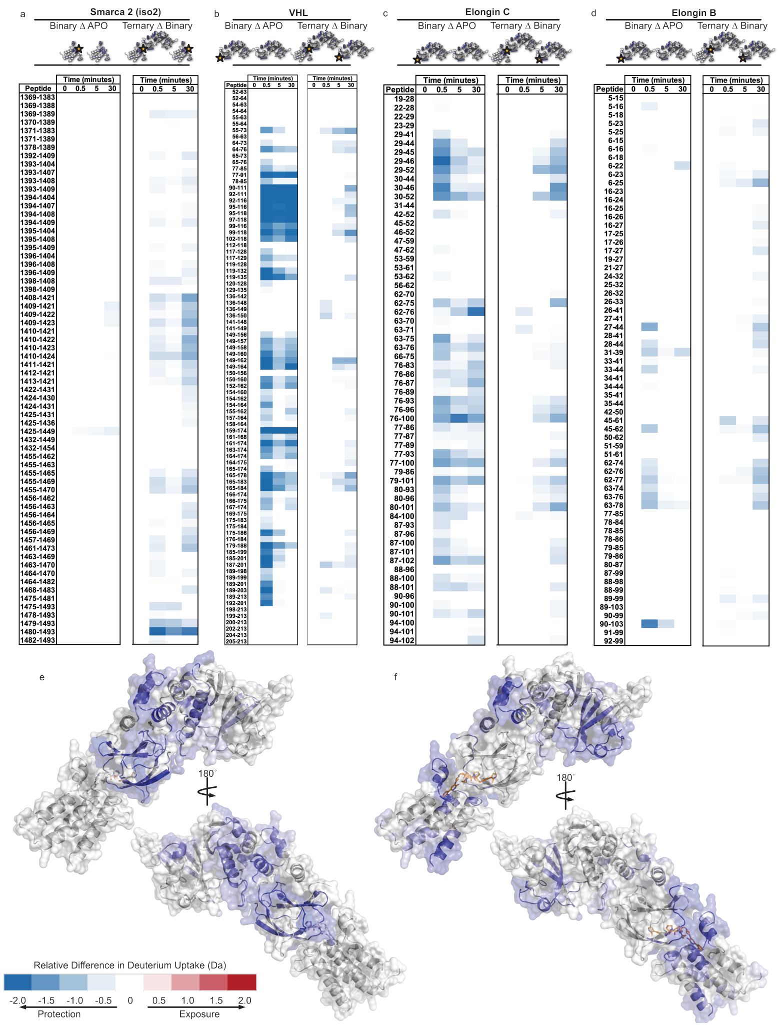

Supplemental Fig. 35: ACBI1-induced ternary complex formation of SMARCA2$^{BD}$:VCB leads to protection of specific sites: (a) iso2-SMARCA2$^{BD}$, (b) VHL, (c) Elongin C, and (d) Elongin B monitored for hydrogen-deuterium exchange over time. The difference plots of each protein are generated by subtracting the deuterium exchange of like peptides of the APO or binary from the binary or ternary states (defined as Binary$\Delta$APO and Ternary$\Delta$Binary), respectively. Regions that exchange significantly less than the comparative state are depicted in blue (negative), whereas regions that exchange significantly more appear in red (positive). The resultant difference plots of the binary (e), or ternary complex (f) were mapped onto the structure crystal structure (PDB ID: 7S4E). The experiments were repeated on 2 separate days. All raw relative uptake plots of the deuterium exchange for each state and experiment can be found in Supplemental Figs. 53– 103.

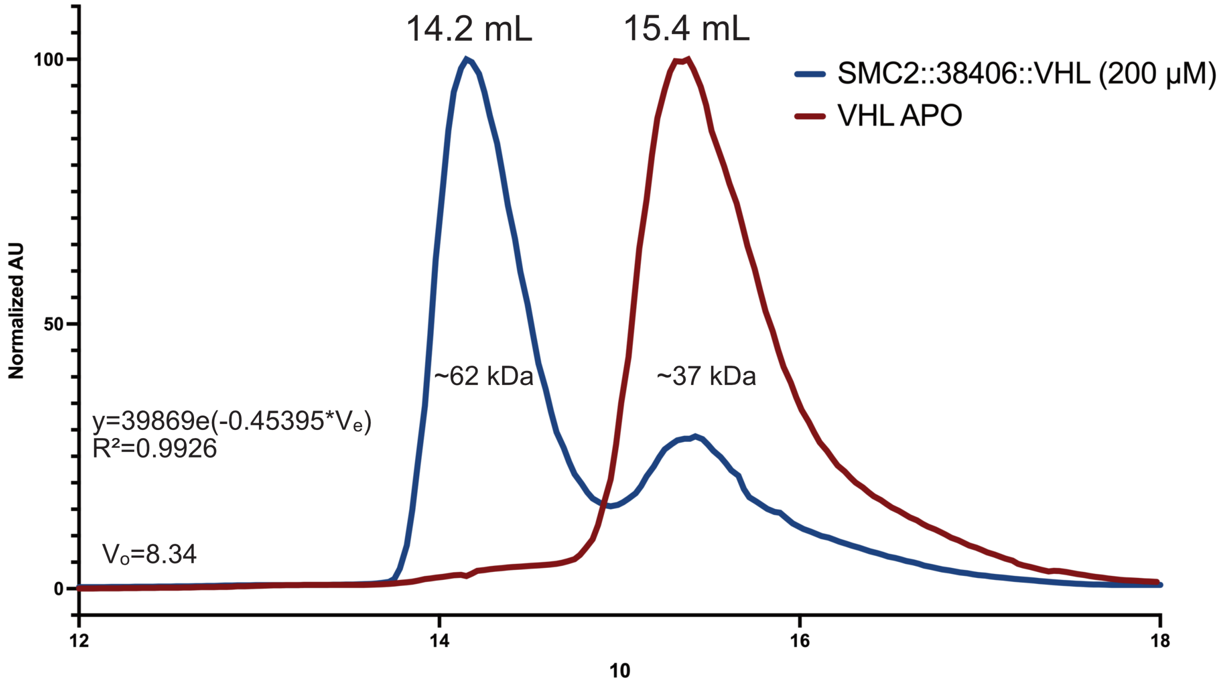


Supplemental Fig. 36: Analytical Size Exclusion with Superdex 200 GL-10/300 increase (Cytiva), ternary complex of iso2-SMARCA2$^{BD}$:ACBI1:VHL/ElonginC/ElonginB, separation from binary or APO state proteins.

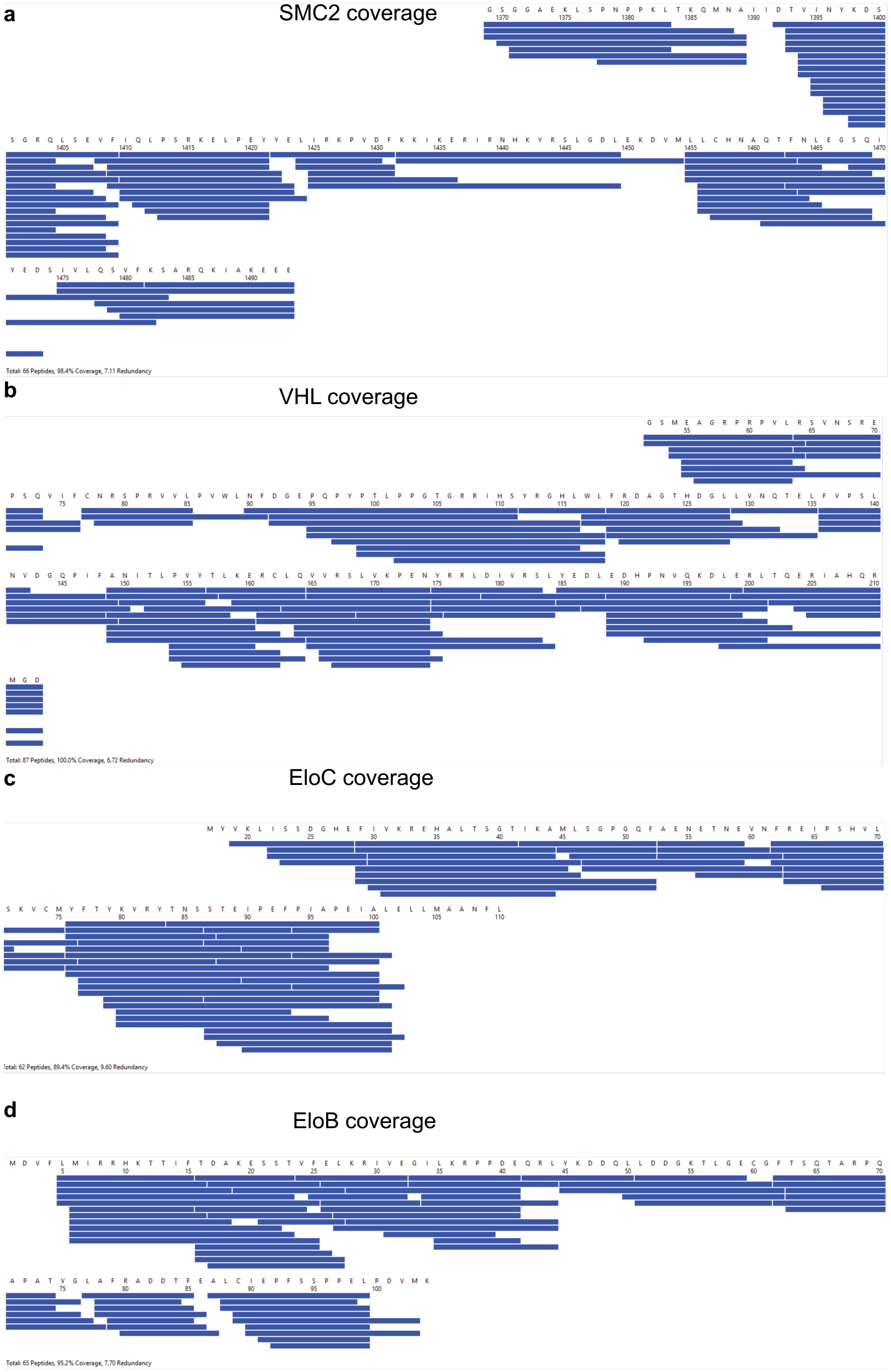


Supplemental Fig. 37: Peptic coverage map of proteolyzed proteins SMARCA2$^{BD}$, VHL, Elongin C and Elongin B.

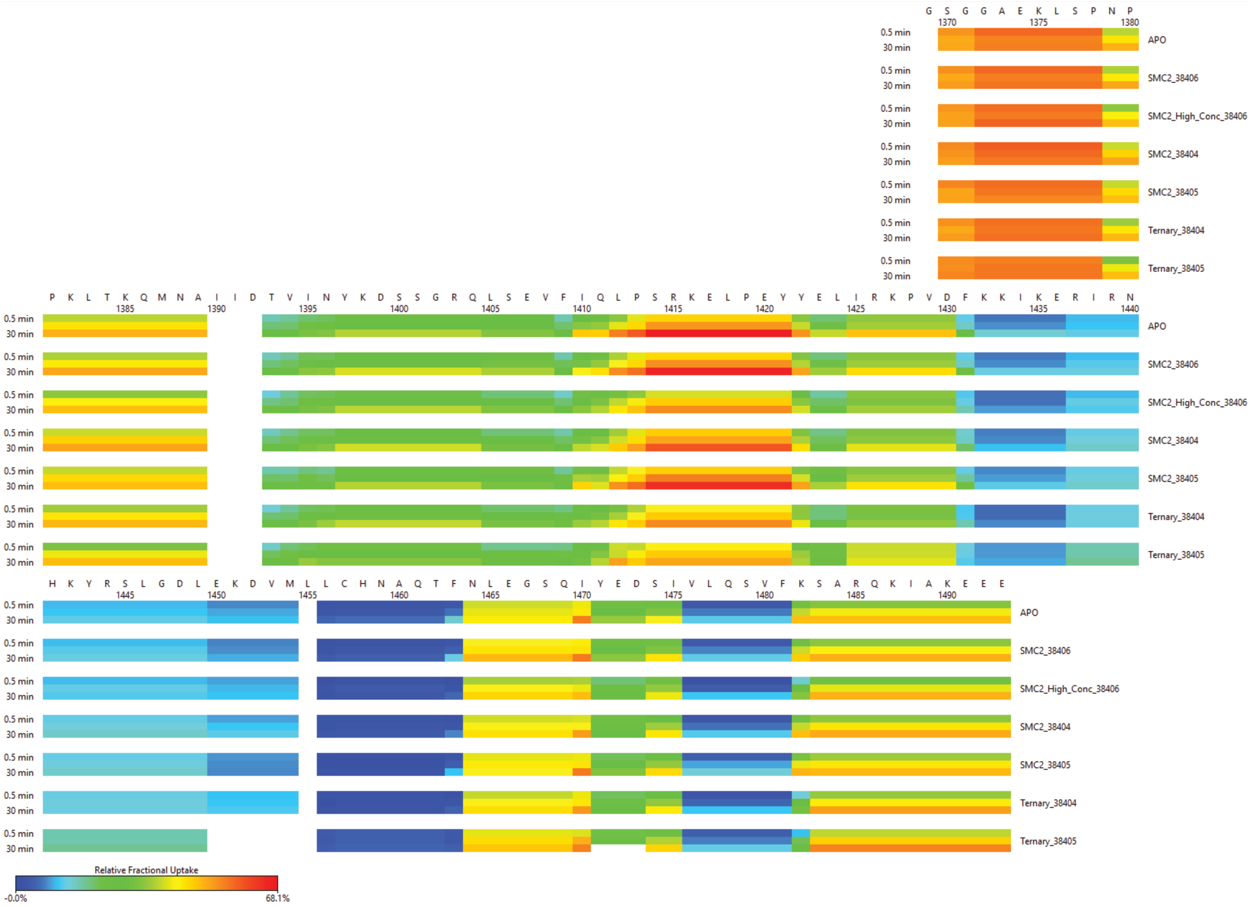


Supplemental Fig. 38: Relative uptake heat map of HDX-MS exchange data of all PROTAC 1, PROTAC 2, and ACBI1 degraders bound to binary and ternary state iso2-SMARCA2$^{BD}$.

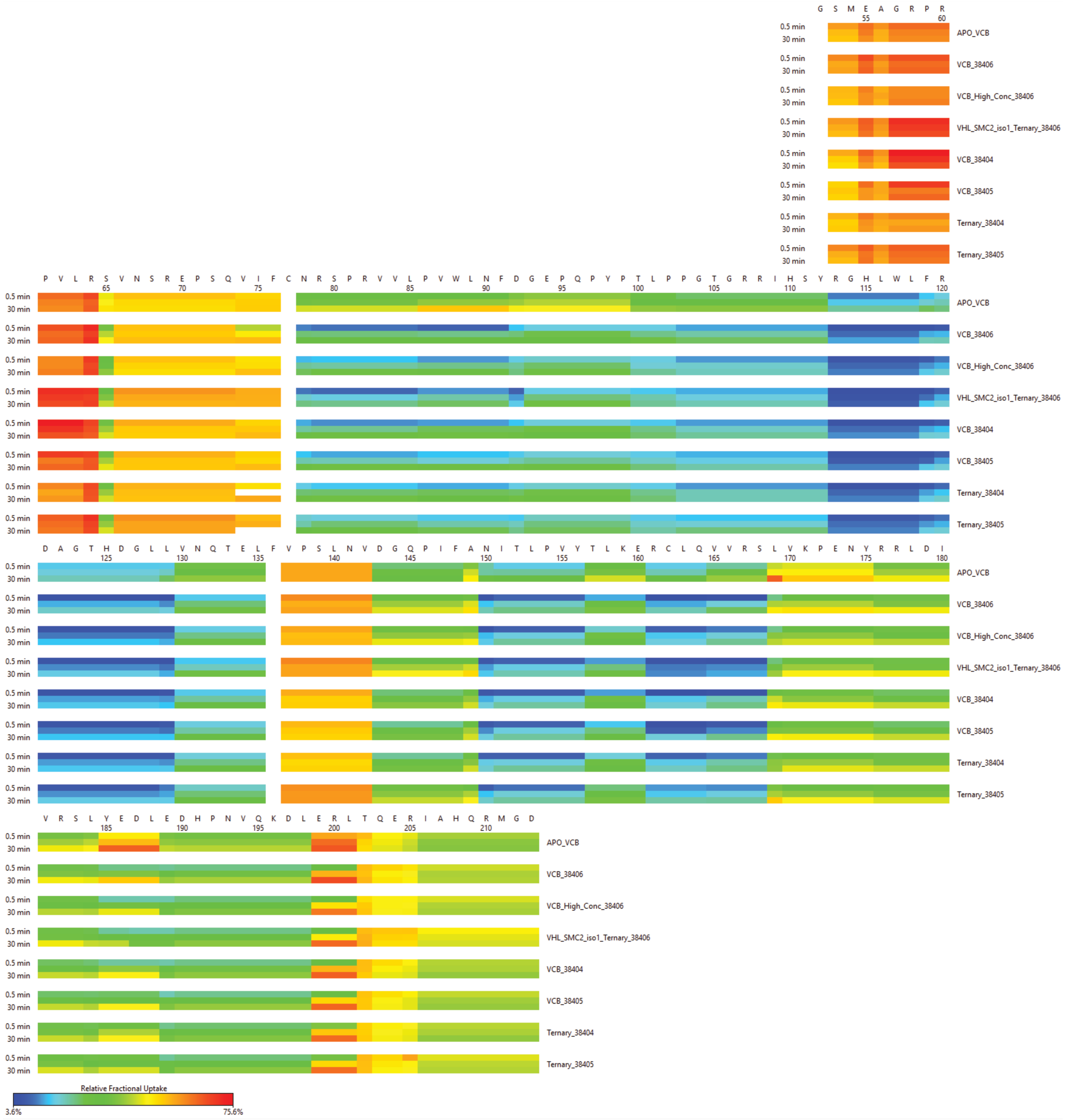

Supplemental Fig. 39: Relative uptake heat map of HDX-MS exchange data of all PROTAC 1, PROTAC 2, and ACBI1 degraders bound to binary and ternary state VHL.

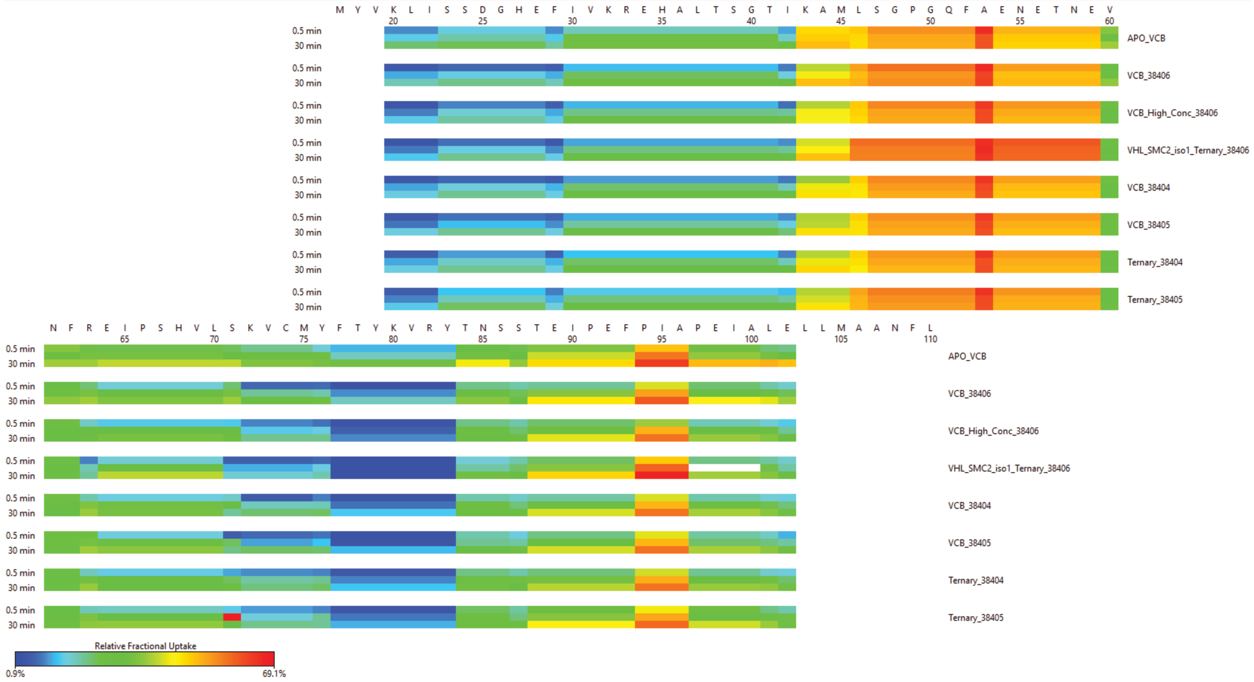


Supplemental Fig. 40: Relative uptake heat map of HDX-MS exchange data of all PROTAC 1, PROTAC 2, and ACBI1 degraders bound to binary and ternary state Elongin C.

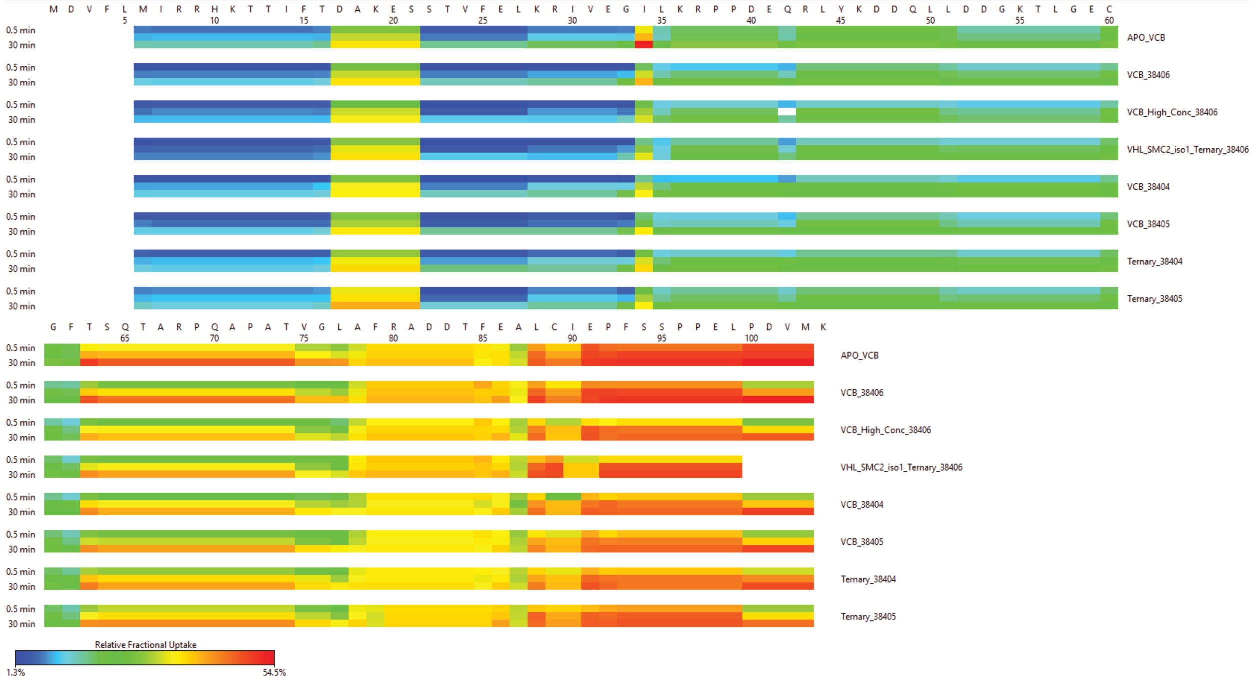


Supplemental Fig. 41: Relative uptake heat map of HDX-MS exchange data of all PROTAC 1, PROTAC 2, and ACBI1 degraders bound to binary and ternary state Elongin B.

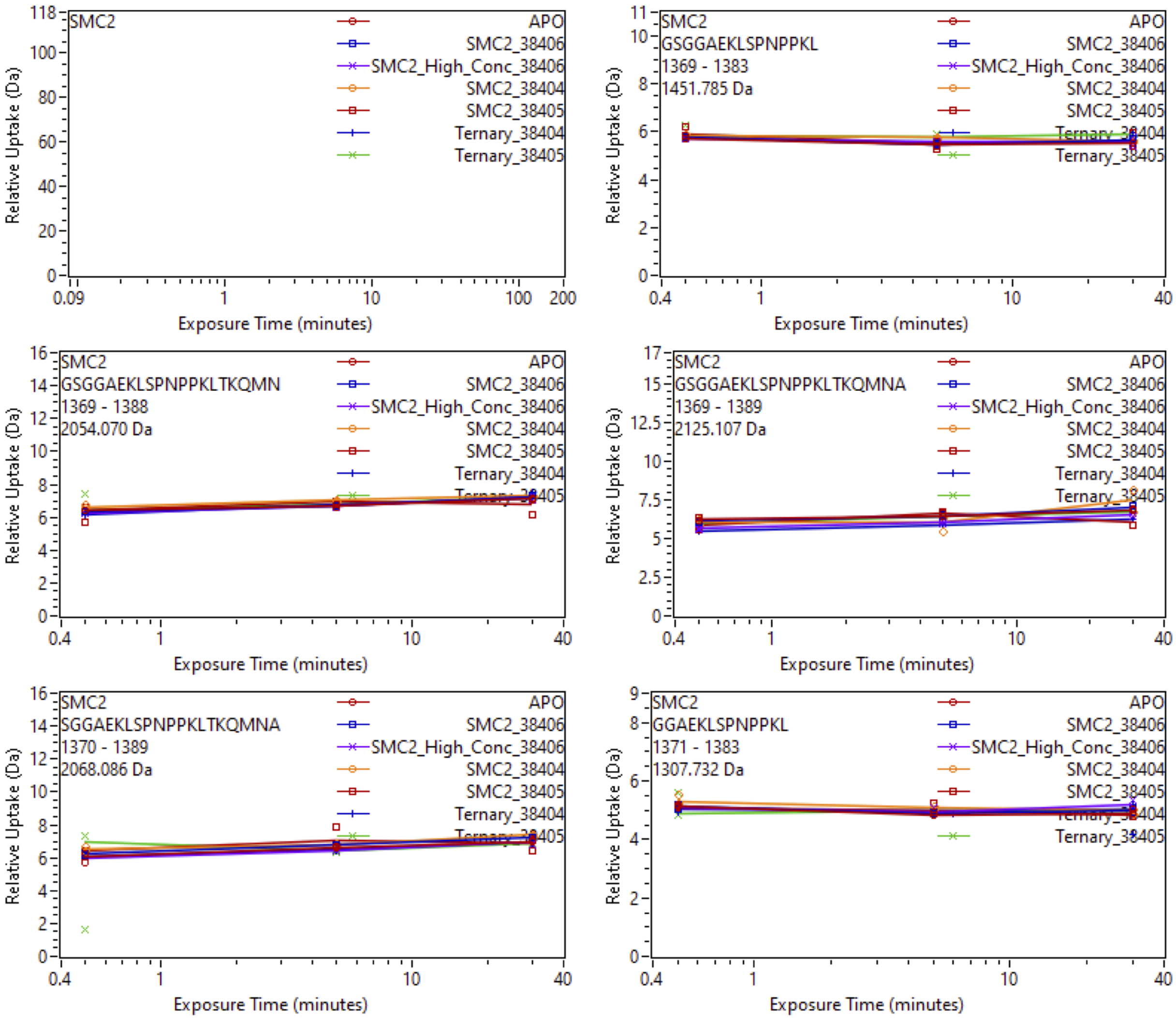


Supplemental Fig. 42: Relative deuterium uptake plots of peptic peptides of iso2-SMARCA2$^{BD}$ in the APO, Binary with SiTX-0038404 (PROTAC 1), SiTX-0038405(PROTAC 2), SiTX-0038406(ACBI1) or Ternary complex with 404, 405, 406 + VCB.

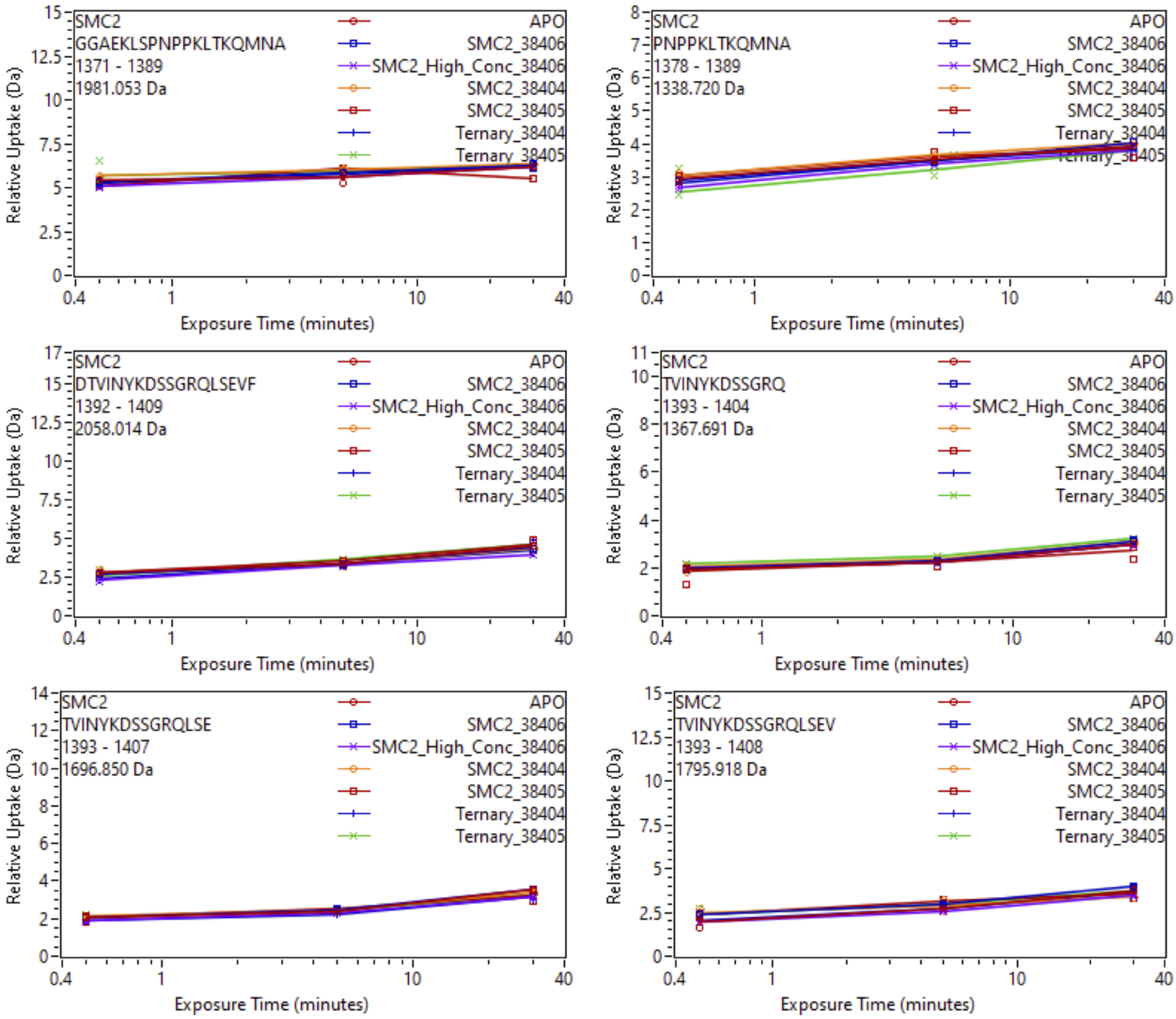


Supplemental Fig. 43: Relative deuterium uptake plots of peptic peptides of iso2-SMARCA2$^{BD}$ in the APO, Binary with SiTX-0038404 (PROTAC 1), SiTX-0038405(PROTAC 2), SiTX-0038406(ACBI1) or Ternary complex with 404, 405, 406 + VCB.

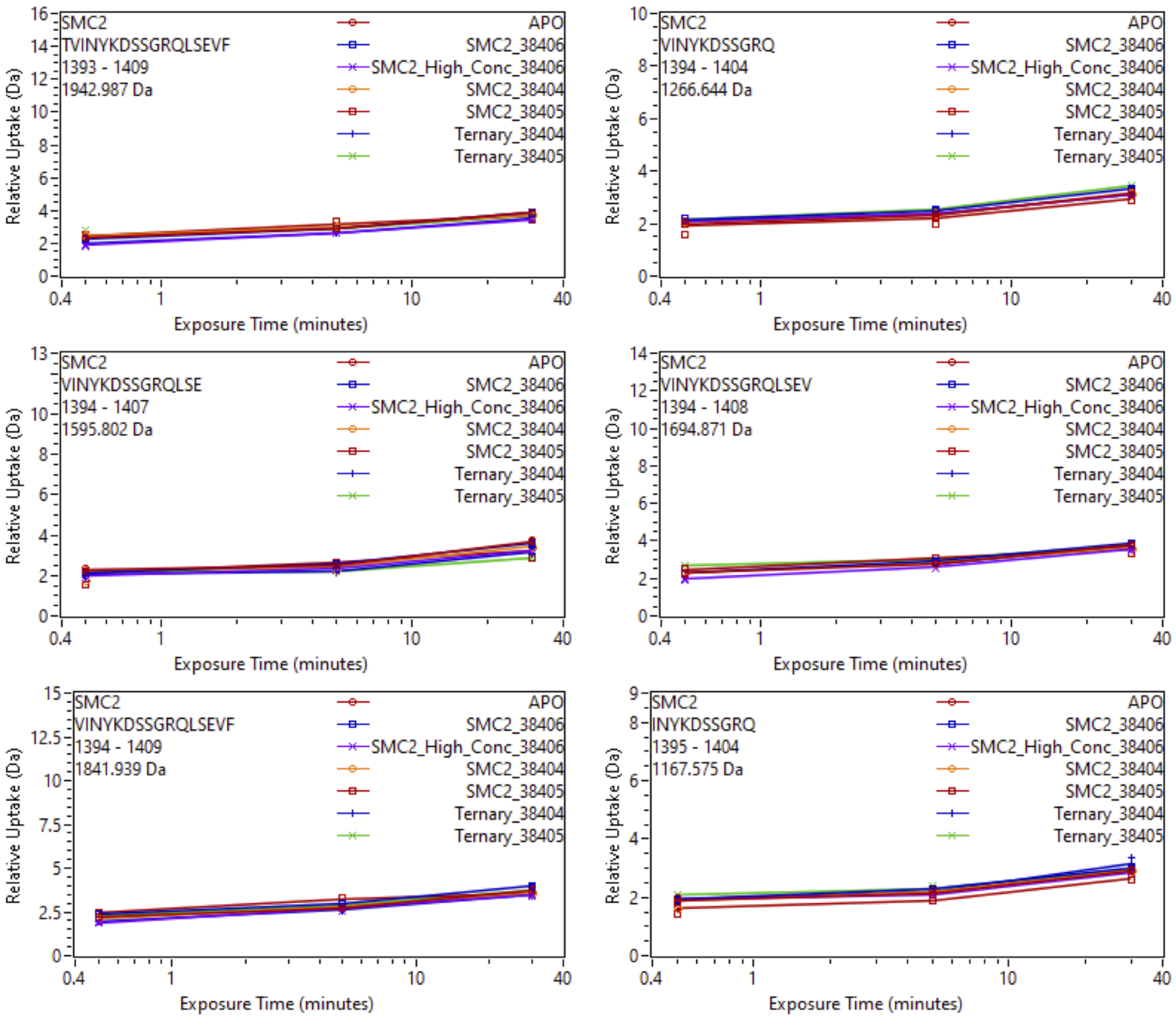


Supplemental Fig. 44: Relative deuterium uptake plots of peptic peptides of iso2-SMARCA2$^{BD}$ in the APO, Binary with SiTX-0038404 (PROTAC 1), SiTX-0038405(PROTAC 2), SiTX-0038406(ACBI1) or Ternary complex with 404, 405, 406 + VCB.

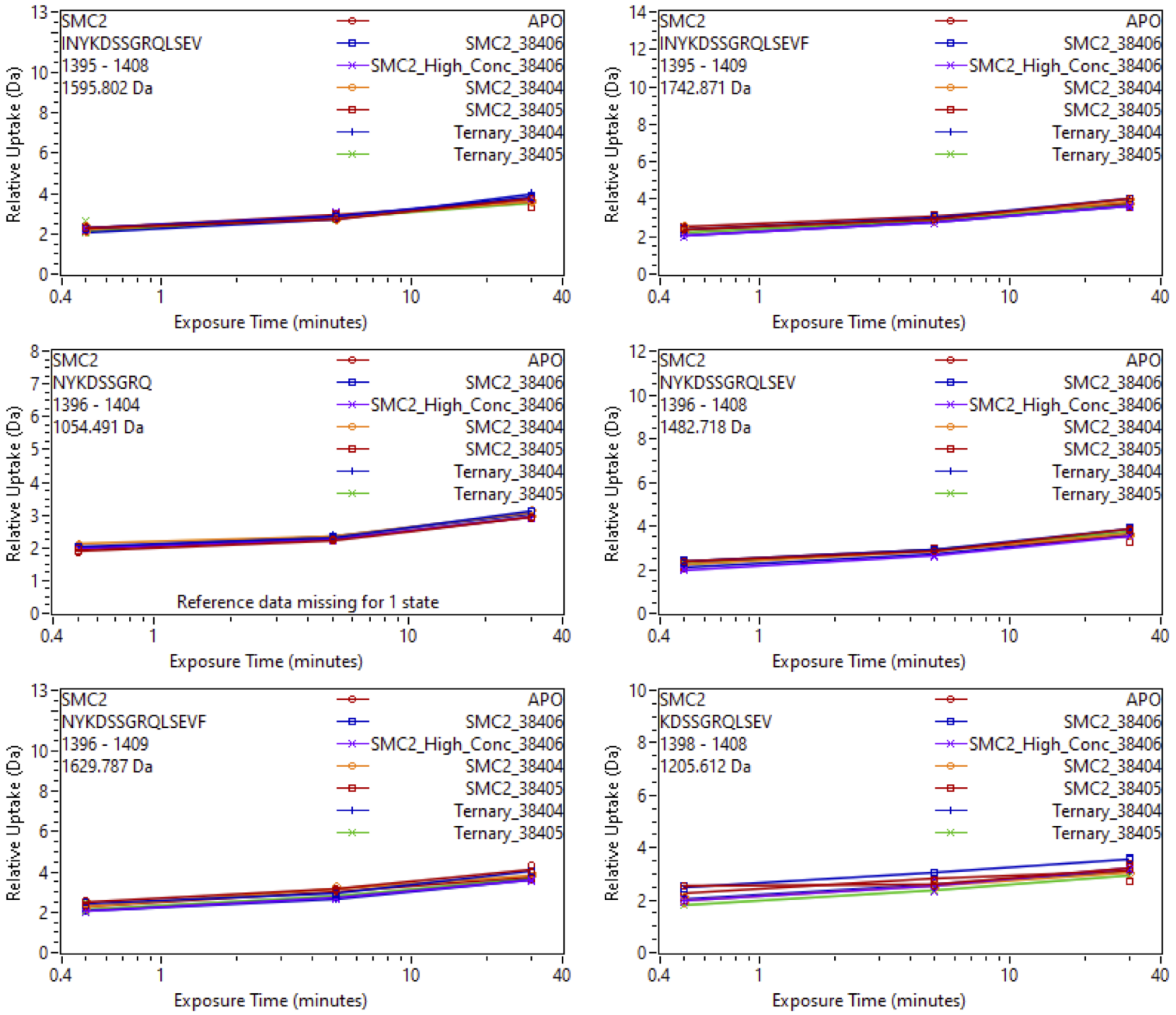


Supplemental Fig. 45: Relative deuterium uptake plots of peptic peptides of iso2-SMARCA2$^{BD}$ in the APO, Binary with SiTX-0038404 (PROTAC 1), SiTX-0038405(PROTAC 2), SiTX-0038406(ACBI1) or Ternary complex with 404, 405, 406 + VCB.

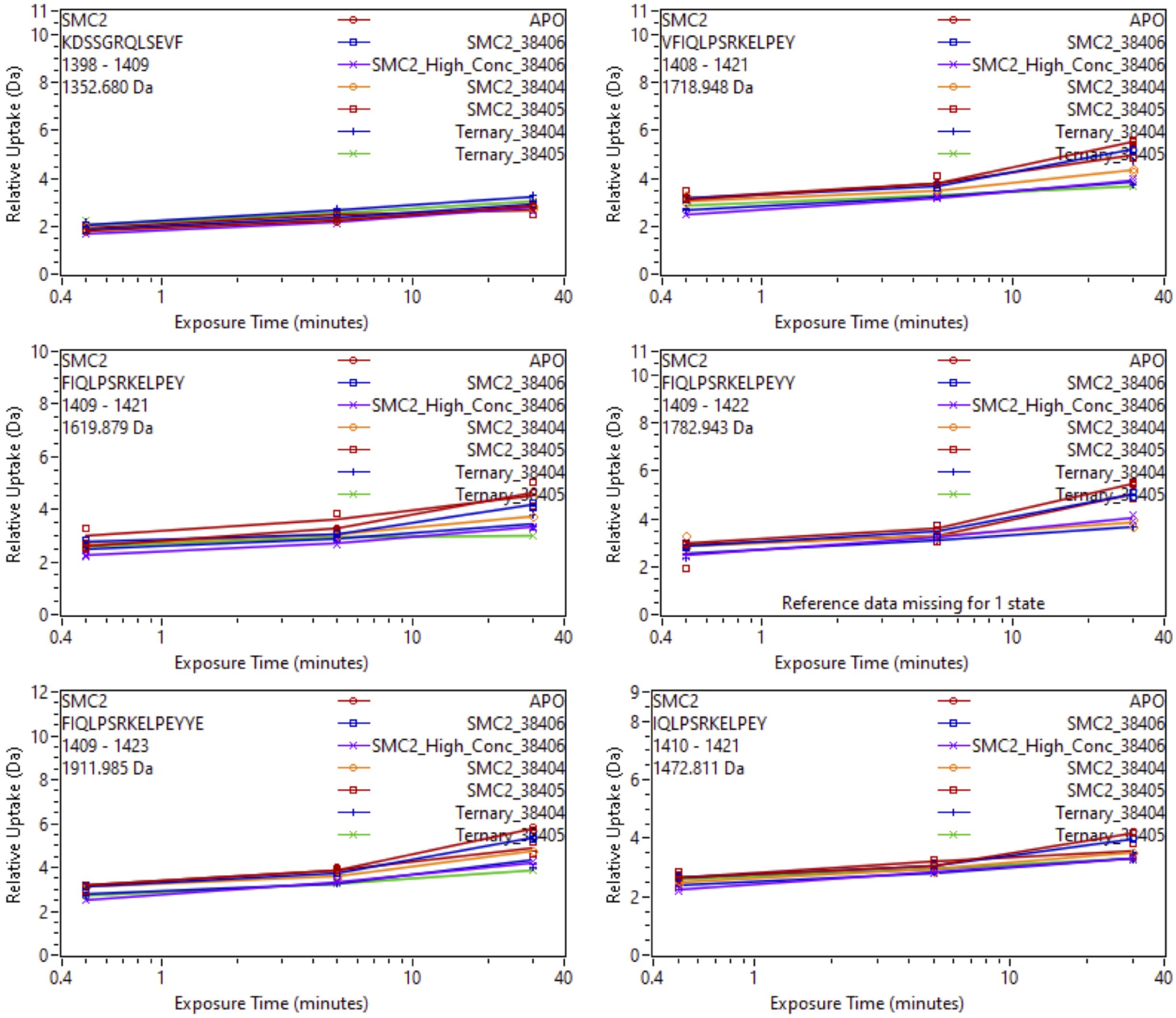


Supplemental Fig. 46: Relative deuterium uptake plots of peptic peptides of iso2-SMARCA2$^{BD}$ in the APO, Binary with SiTX-0038404 (PROTAC 1), SiTX-0038405(PROTAC 2), SiTX-0038406(ACBI1) or Ternary complex with 404, 405, 406 + VCB.

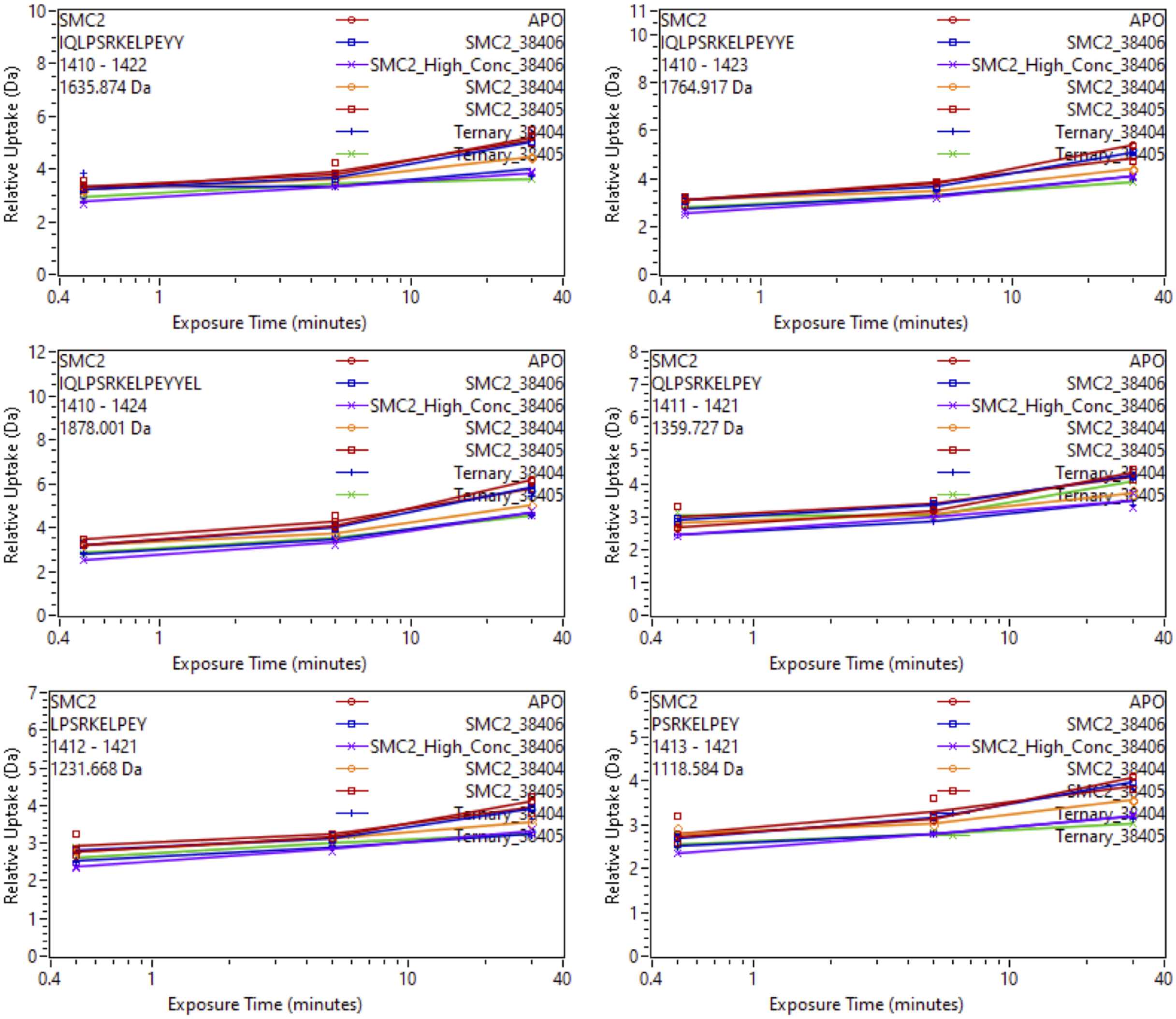


Supplemental Fig. 47: Relative deuterium uptake plots of peptic peptides of iso2-SMARCA2$^{BD}$ in the APO, Binary with SiTX-0038404 (PROTAC 1), SiTX-0038405(PROTAC 2), SiTX-0038406(ACBI1) or Ternary complex with 404, 405, 406 + VCB.

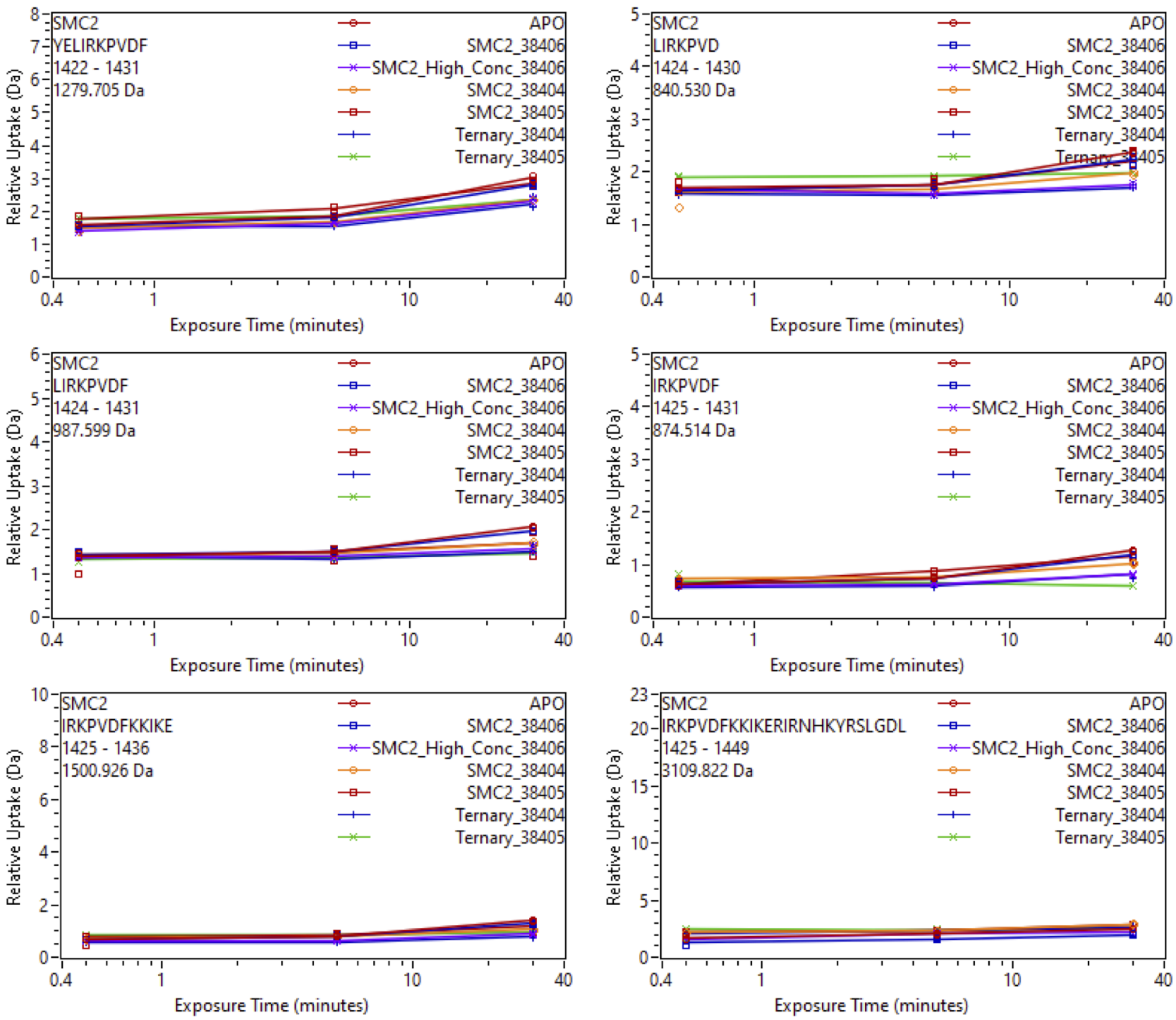


Supplemental Fig. 48: Relative deuterium uptake plots of peptic peptides of iso2-SMARCA2$^{BD}$ in the APO, Binary with SiTX-0038404 (PROTAC 1), SiTX-0038405(PROTAC 2), SiTX-0038406(ACBI1) or Ternary complex with 404, 405, 406 + VCB.

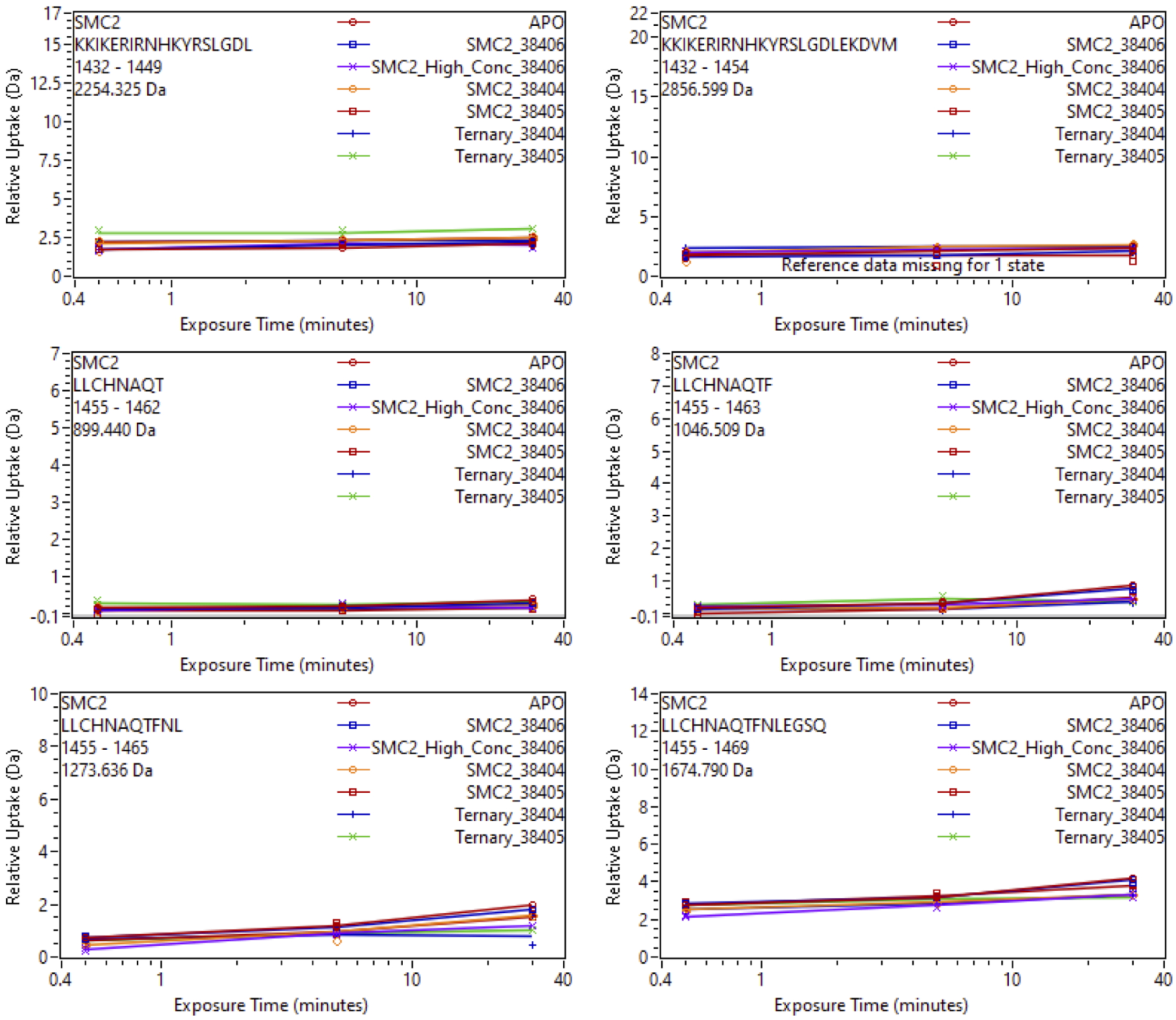


Supplemental Fig. 49: Relative deuterium uptake plots of peptic peptides of iso2-SMARCA2$^{BD}$ in the APO, Binary with SiTX-0038404 (PROTAC 1), SiTX-0038405(PROTAC 2), SiTX-0038406(ACBI1) or Ternary complex with 404, 405, 406 + VCB.

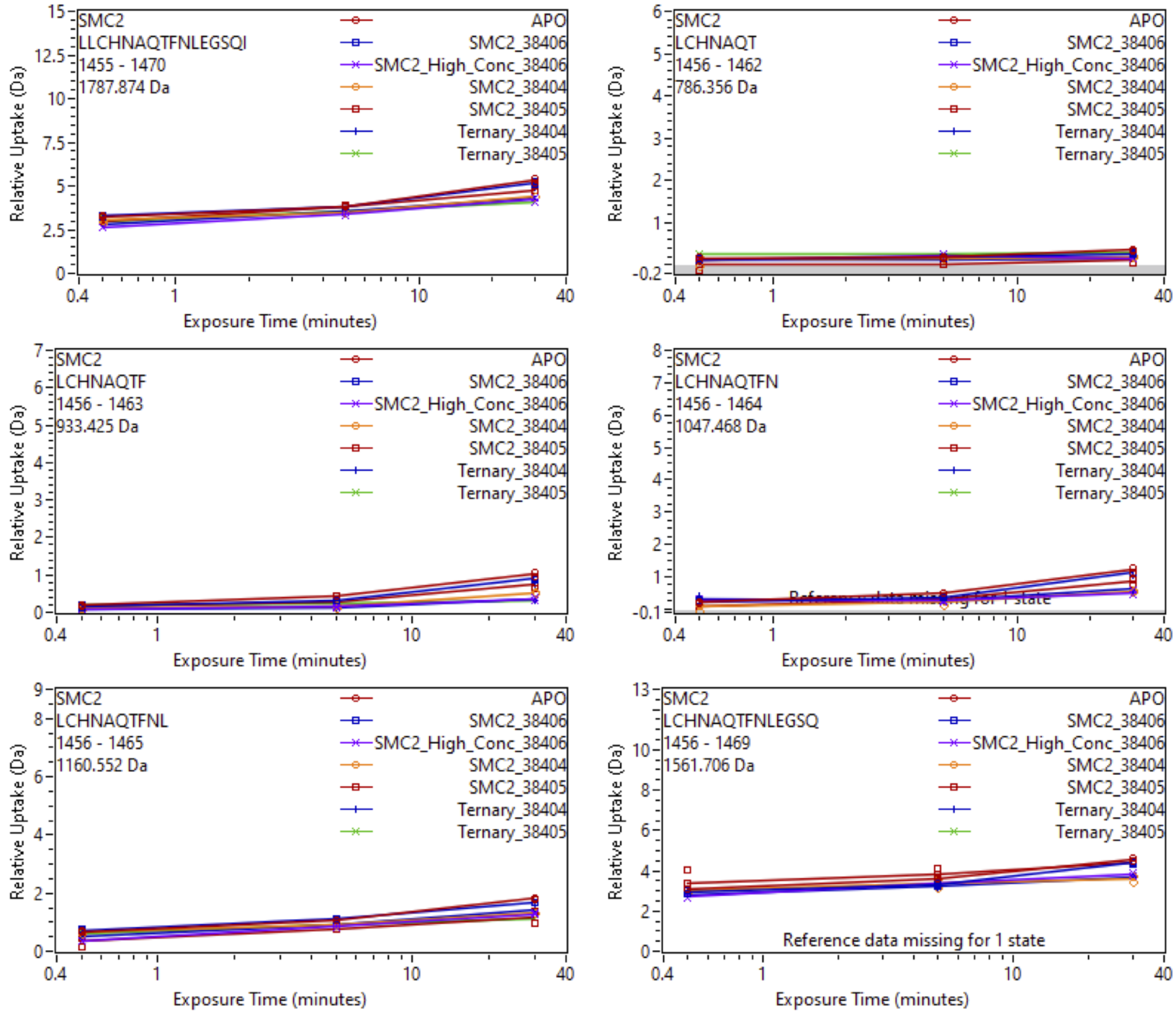


Supplemental Fig. 50: Relative deuterium uptake plots of peptic peptides of iso2-SMARCA2$^{BD}$ in the APO, Binary with SiTX-0038404 (PROTAC 1), SiTX-0038405(PROTAC 2), SiTX-0038406(ACBI1) or Ternary complex with 404, 405, 406 + VCB.

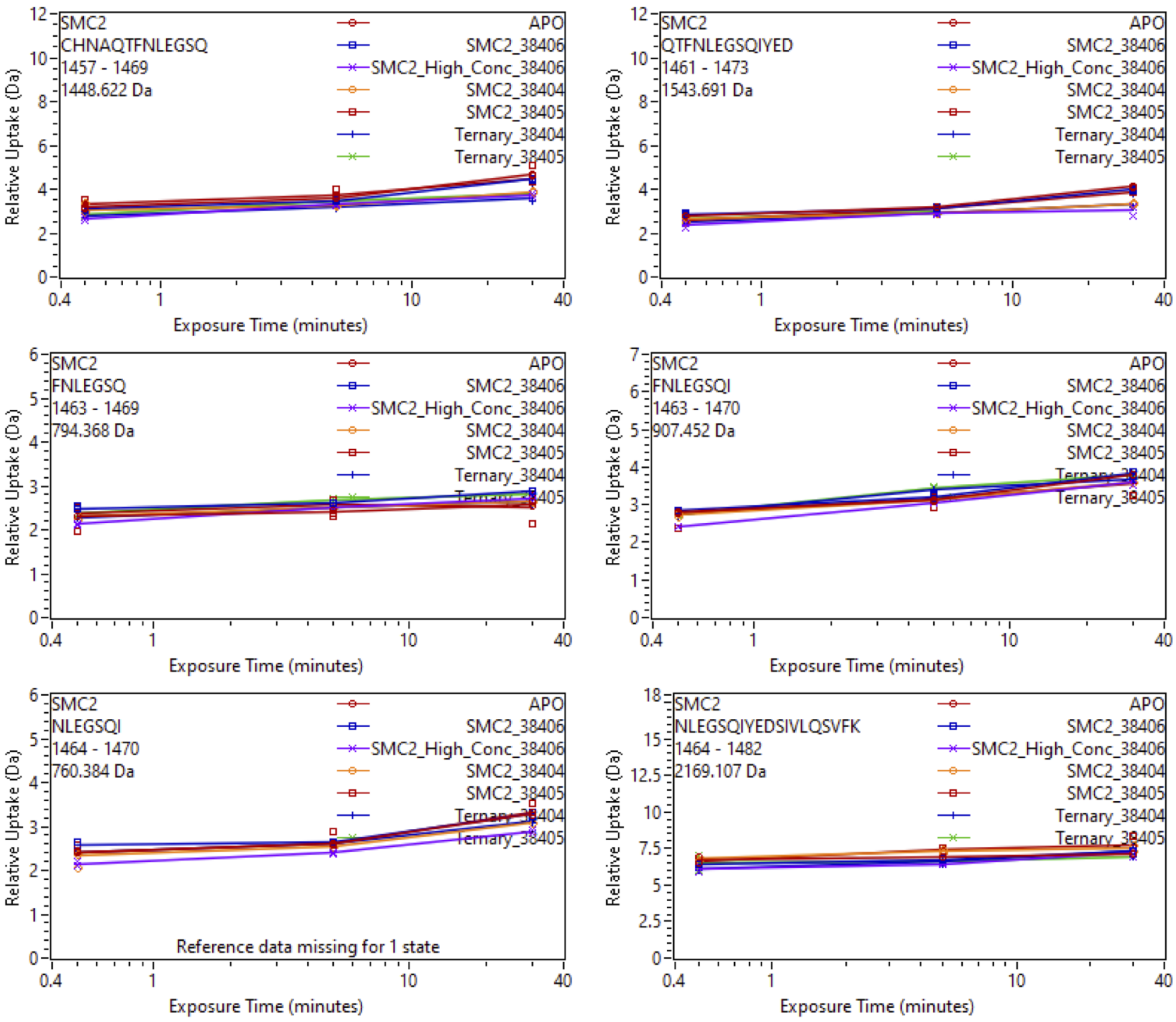


Supplemental Fig. 51: Relative deuterium uptake plots of peptic peptides of iso2-SMARCA2$^{BD}$ in the APO, Binary with SiTX-0038404 (PROTAC 1), SiTX-0038405(PROTAC 2), SiTX-0038406(ACBI1) or Ternary complex with 404, 405, 406 + VCB.

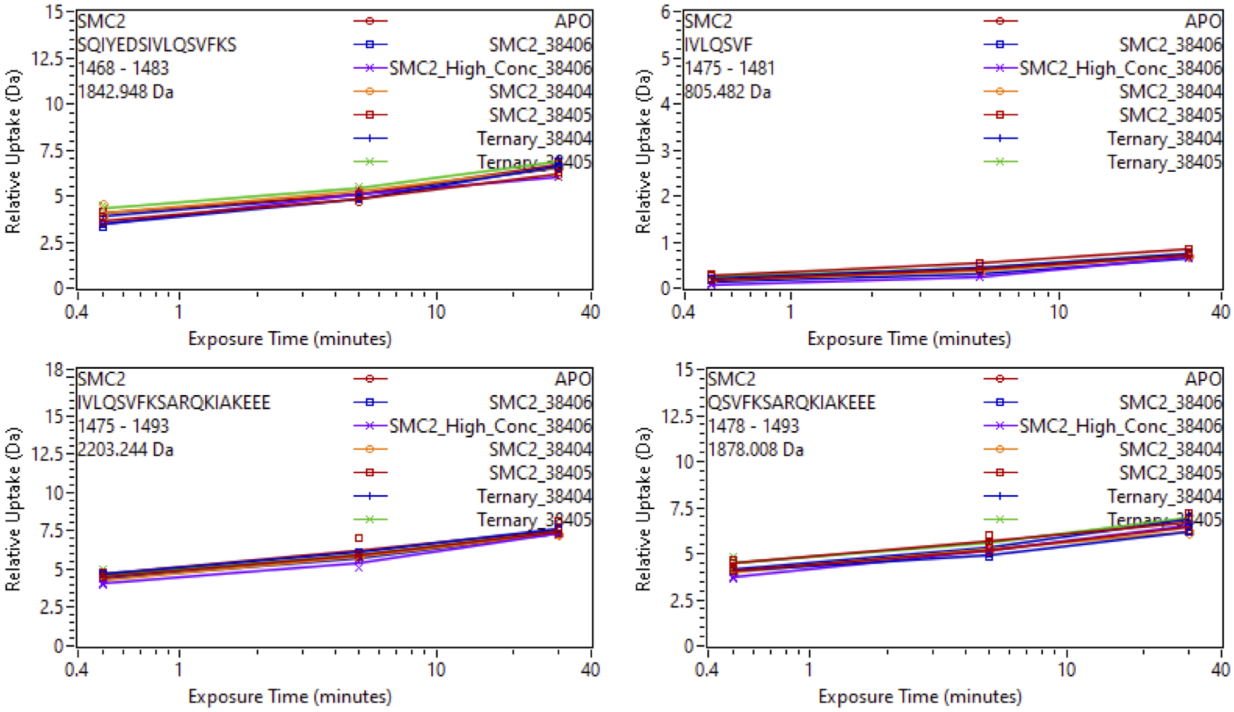


Supplemental Fig. 52: Relative deuterium uptake plots of peptic peptides of iso2-SMARCA2$^{BD}$ in the APO, Binary with SiTX-0038404 (PROTAC 1), SiTX-0038405(PROTAC 2), SiTX-0038406(ACBI1) or Ternary complex with 404, 405, 406 + VCB.

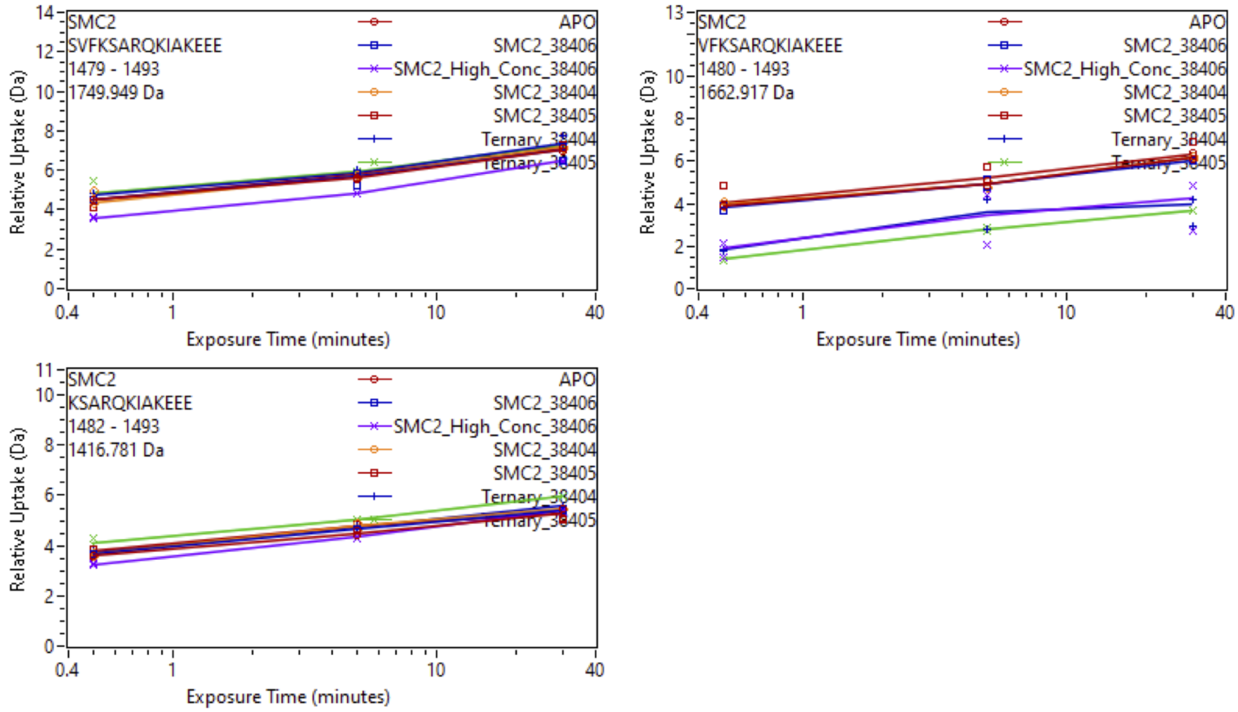


Supplemental Fig. 53: Relative deuterium uptake plots of peptic peptides of iso2-SMARCA2$^{BD}$ in the APO, Binary with SiTX-0038404 (PROTAC 1), SiTX-0038405(PROTAC 2), SiTX-0038406(ACBI1) or Ternary complex with 404, 405, 406 + VCB.

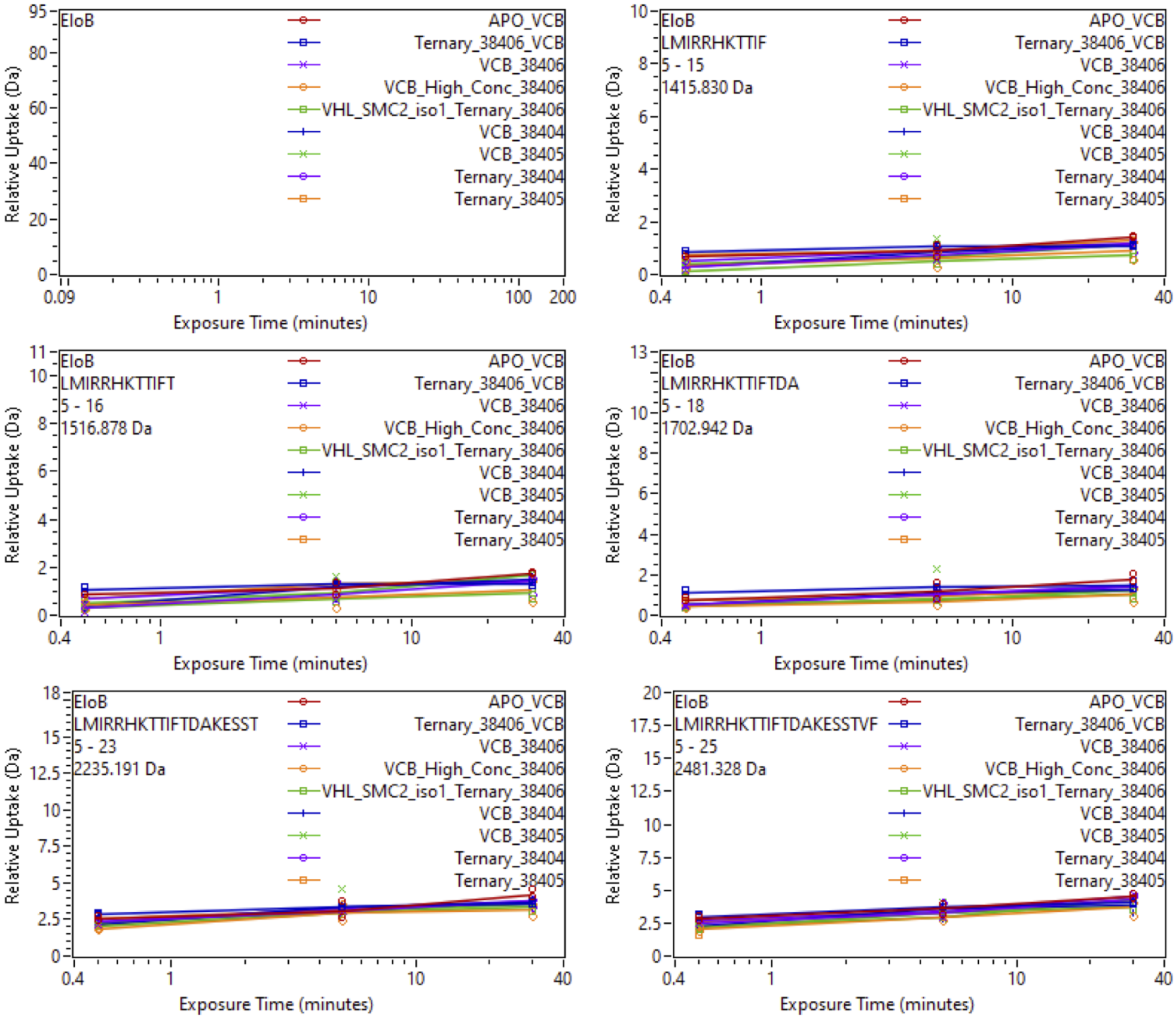


Supplemental Fig. 54: Relative deuterium uptake plots of peptic peptides Elongin B of the VCB complex in the APO, Binary with SiTX-0038404 (PROTAC 1), SiTX-0038405(PROTAC 2), SiTX-0038406(ACBI1) or Ternary complex with 404, 405, 406 + SMARCA2$^{BD}$.

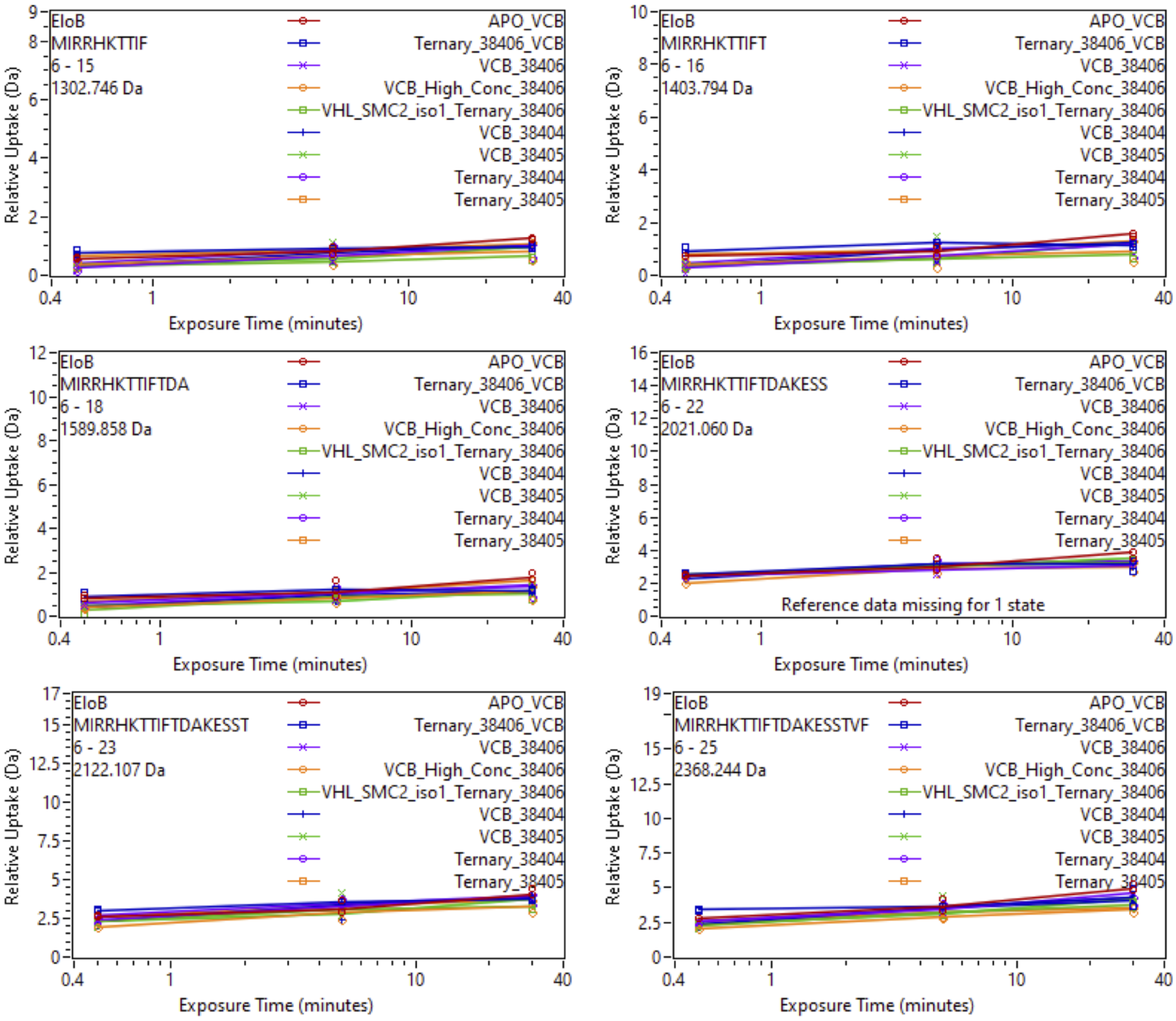


Supplemental Fig. 55: Relative deuterium uptake plots of peptic peptides Elongin B of the VCB complex in the APO, Binary with SiTX-0038404 (PROTAC 1), SiTX-0038405(PROTAC 2), SiTX-0038406(ACBI1) or Ternary complex with 404, 405, 406 + SMARCA2$^{BD}$.

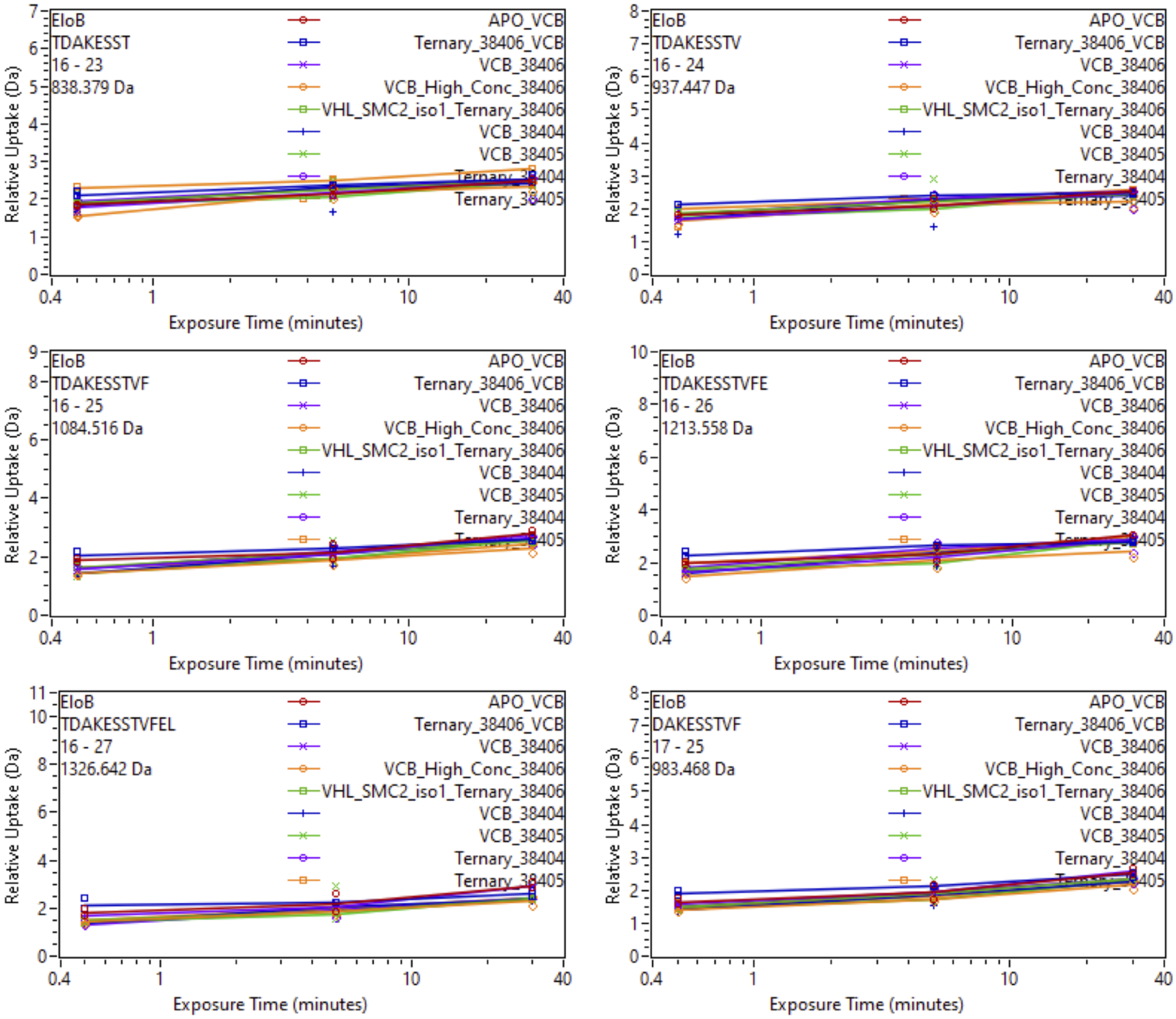


Supplemental Fig. 56: Relative deuterium uptake plots of peptic peptides Elongin B of the VCB complex in the APO, Binary with SiTX-0038404 (PROTAC 1), SiTX-0038405(PROTAC 2), SiTX-0038406(ACBI1) or Ternary complex with 404, 405, 406 + SMARCA2$^{BD}$.

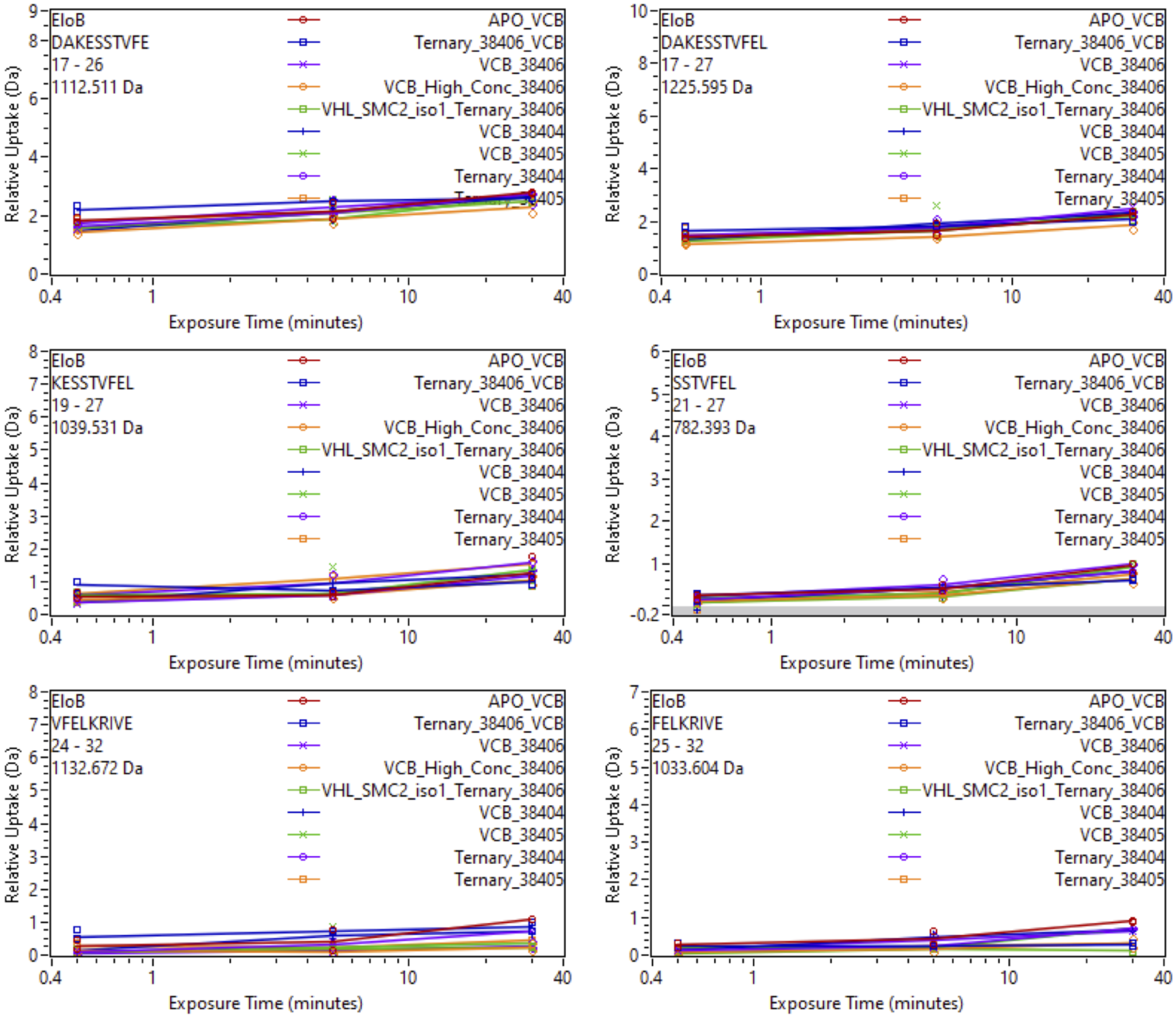


Supplemental Fig. 57: Relative deuterium uptake plots of peptic peptides Elongin B of the VCB complex in the APO, Binary with SiTX-0038404 (PROTAC 1), SiTX-0038405(PROTAC 2), SiTX-0038406(ACBI1) or Ternary complex with 404, 405, 406 + SMARCA2$^{BD}$.

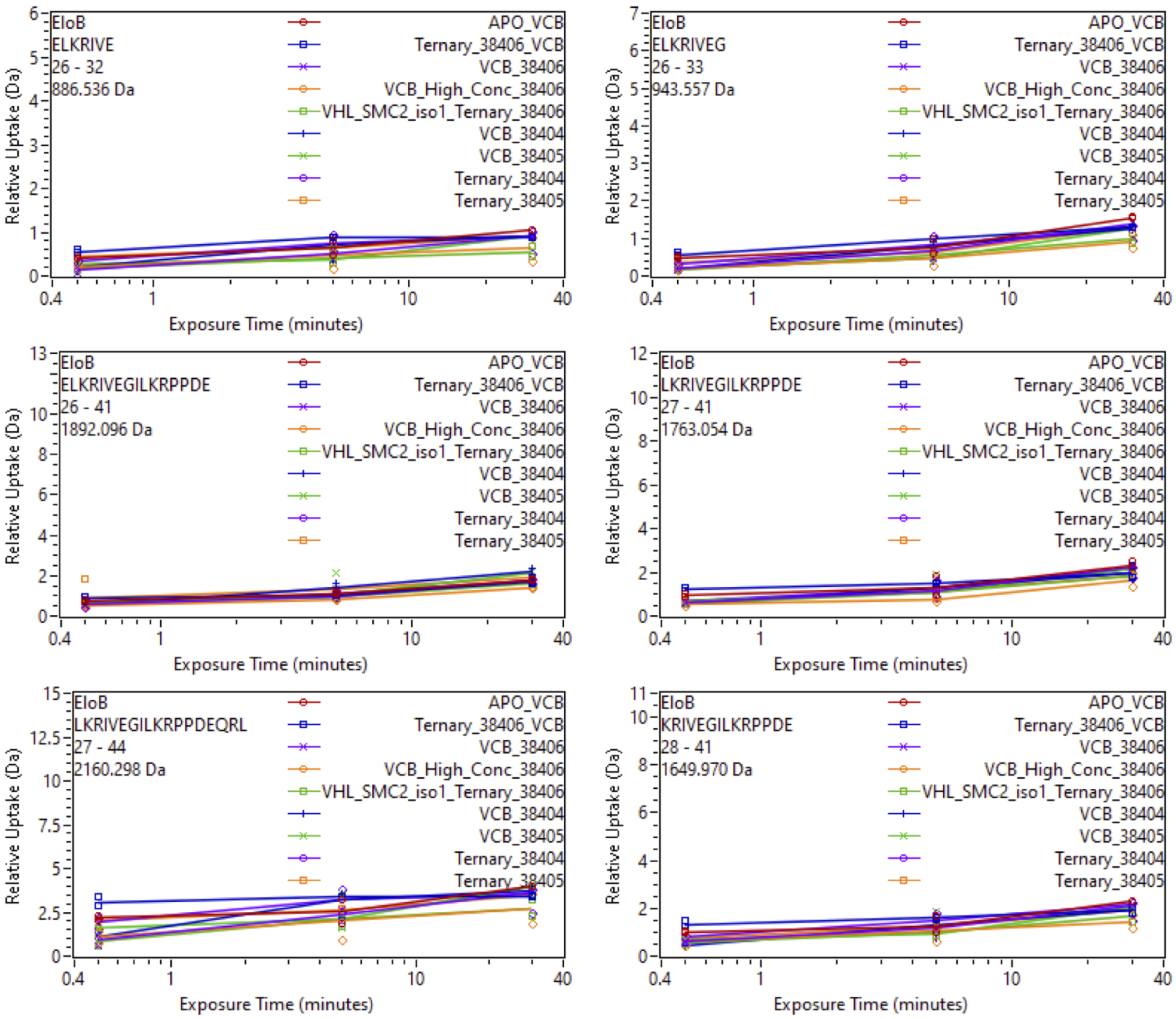


Supplemental Fig. 58: Relative deuterium uptake plots of peptic peptides Elongin B of the VCB complex in the APO, Binary with SiTX-0038404 (PROTAC 1), SiTX-0038405(PROTAC 2), SiTX-0038406(ACBI1) or Ternary complex with 404, 405, 406 + SMARCA2$^{BD}$.

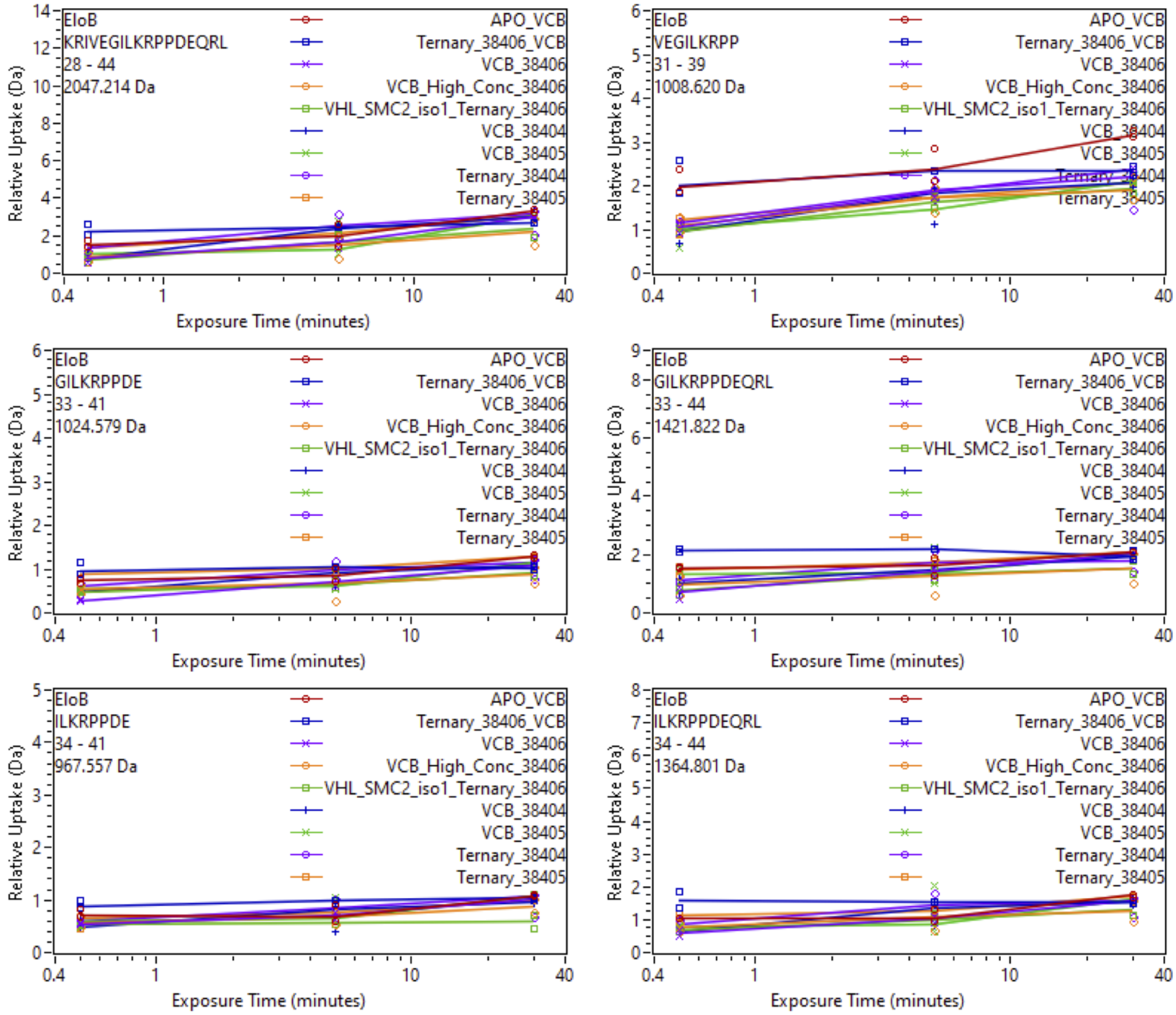


Supplemental Fig. 59: Relative deuterium uptake plots of peptic peptides Elongin B of the VCB complex in the APO, Binary with SiTX-0038404 (PROTAC 1), SiTX-0038405(PROTAC 2), SiTX-0038406(ACBI1) or Ternary complex with 404, 405, 406 + SMARCA2$^{BD}$.

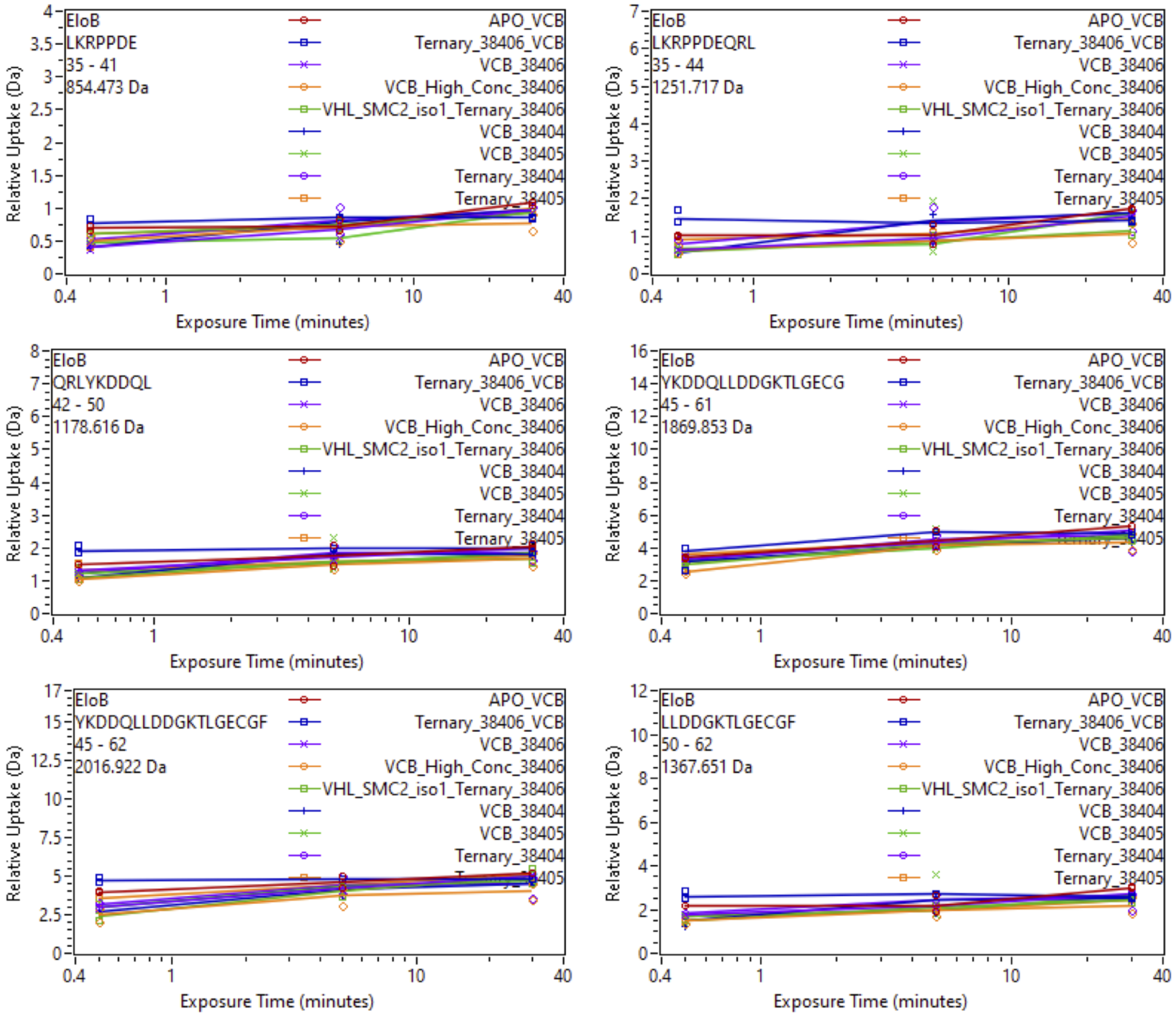


Supplemental Fig. 60: Relative deuterium uptake plots of peptic peptides Elongin B of the VCB complex in the APO, Binary with SiTX-0038404 (PROTAC 1), SiTX-0038405(PROTAC 2), SiTX-0038406(ACBI1) or Ternary complex with 404, 405, 406 + SMARCA2$^{BD}$.

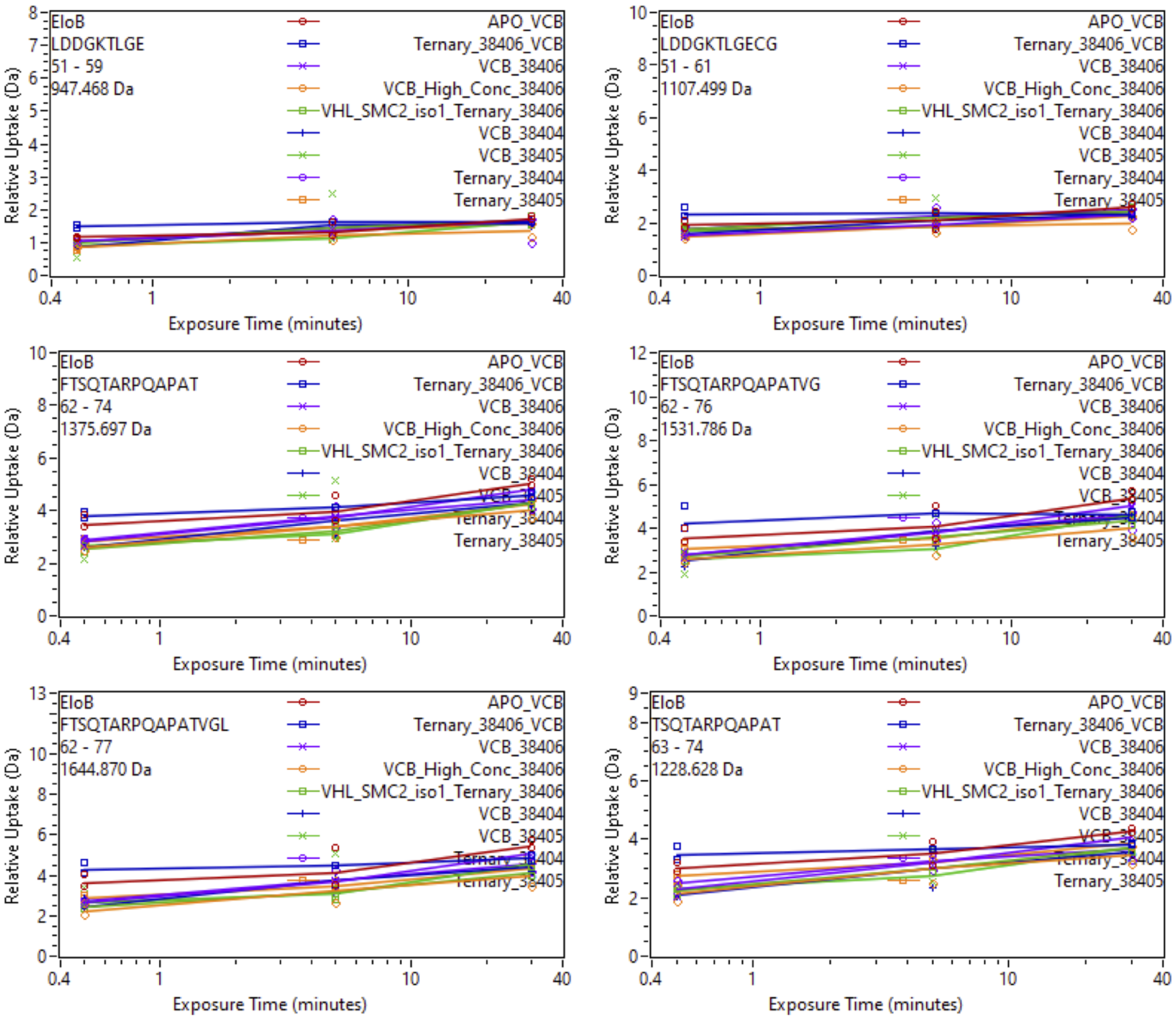


Supplemental Fig. 61: Relative deuterium uptake plots of peptic peptides Elongin B of the VCB complex in the APO, Binary with SiTX-0038404 (PROTAC 1), SiTX-0038405(PROTAC 2), SiTX-0038406(ACBI1) or Ternary complex with 404, 405, 406 + SMARCA2$^{BD}$.

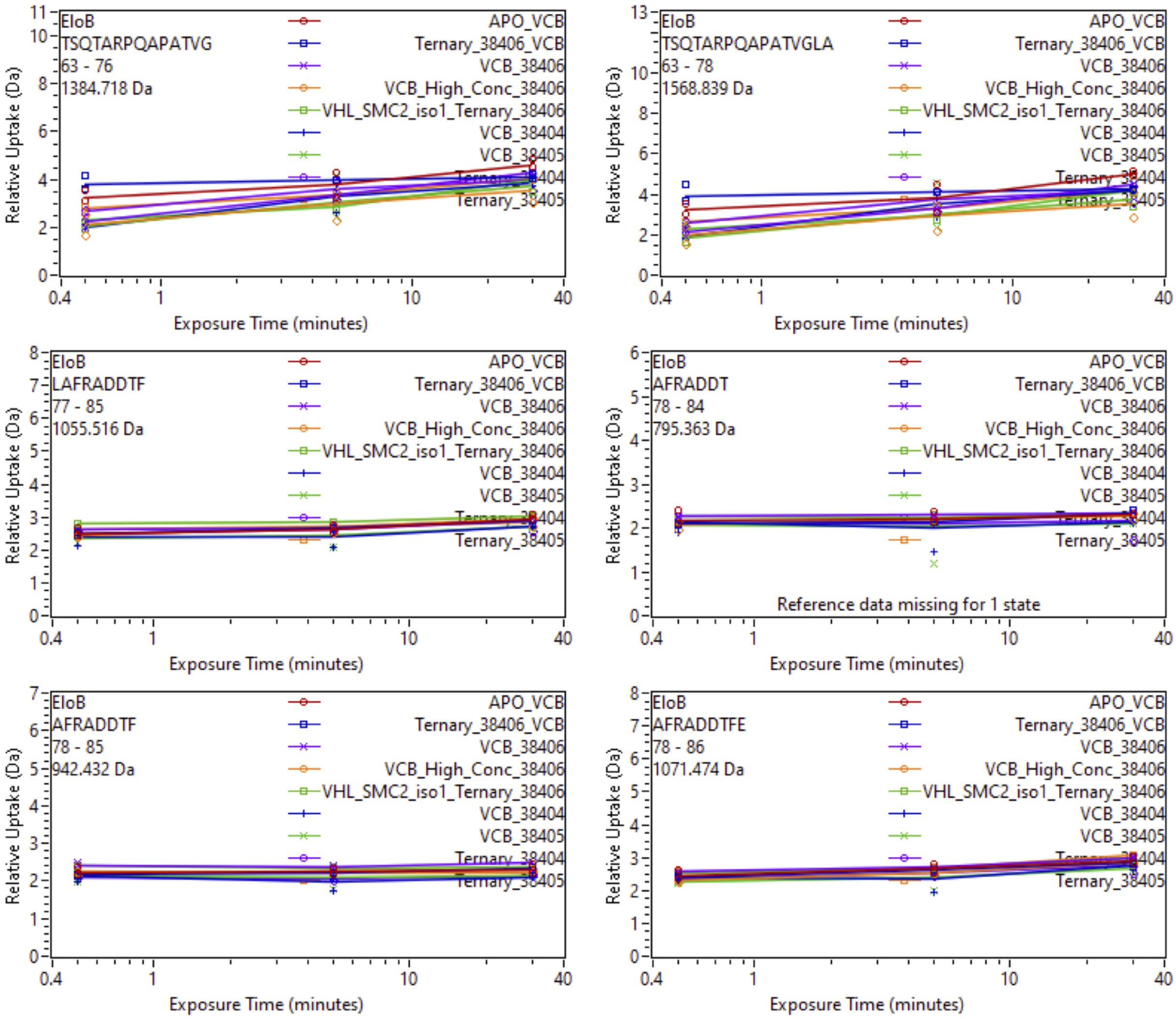


Supplemental Fig. 62: Relative deuterium uptake plots of peptic peptides Elongin B of the VCB complex in the APO, Binary with SiTX-0038404 (PROTAC 1), SiTX-0038405(PROTAC 2), SiTX-0038406(ACBI1) or Ternary complex with 404, 405, 406 + SMARCA2$^{BD}$.

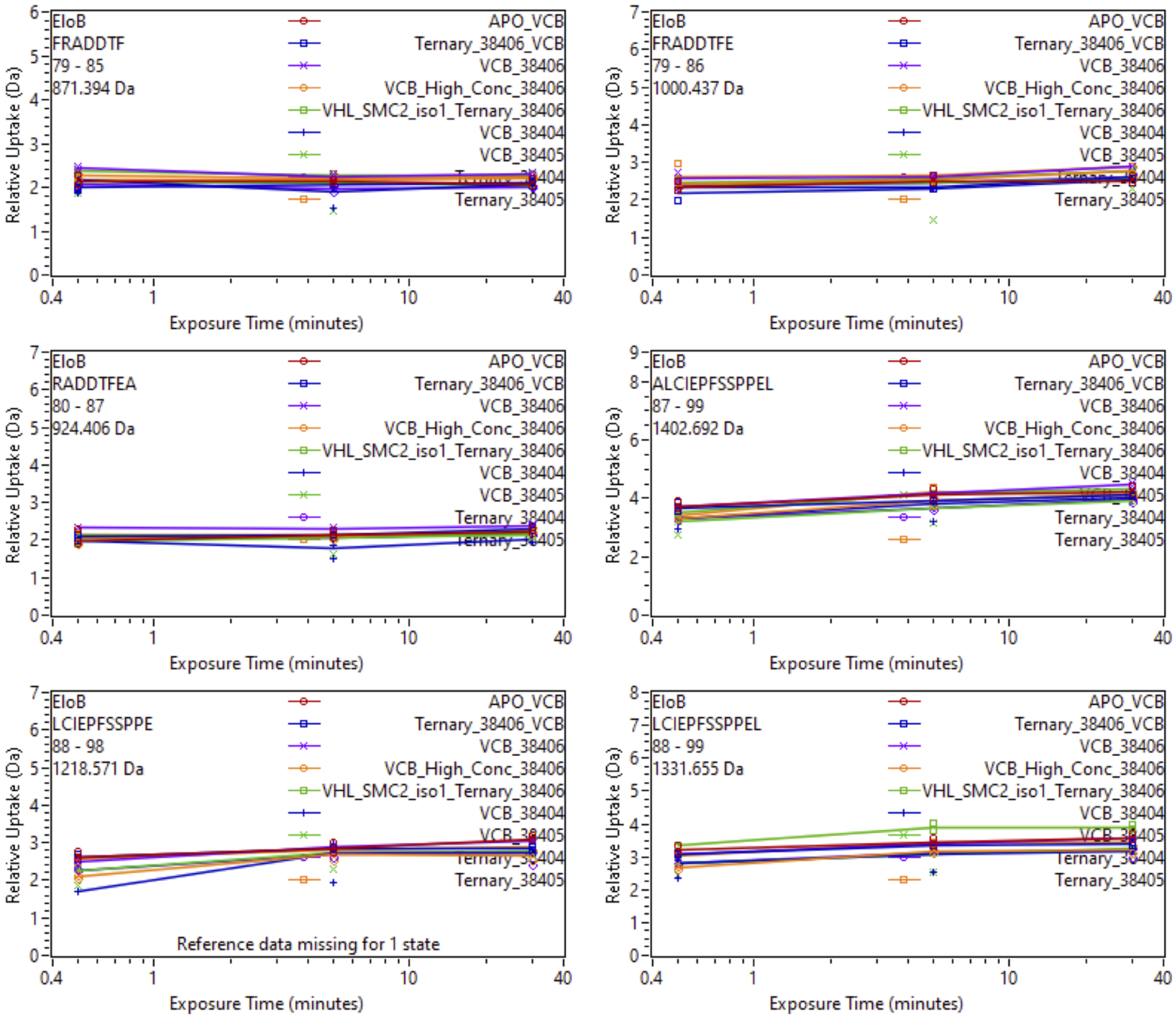


Supplemental Fig. 63: Relative deuterium uptake plots of peptic peptides Elongin B of the VCB complex in the APO, Binary with SiTX-0038404 (PROTAC 1), SiTX-0038405(PROTAC 2), SiTX-0038406(ACBI1) or Ternary complex with 404, 405, 406 + SMARCA2$^{BD}$.

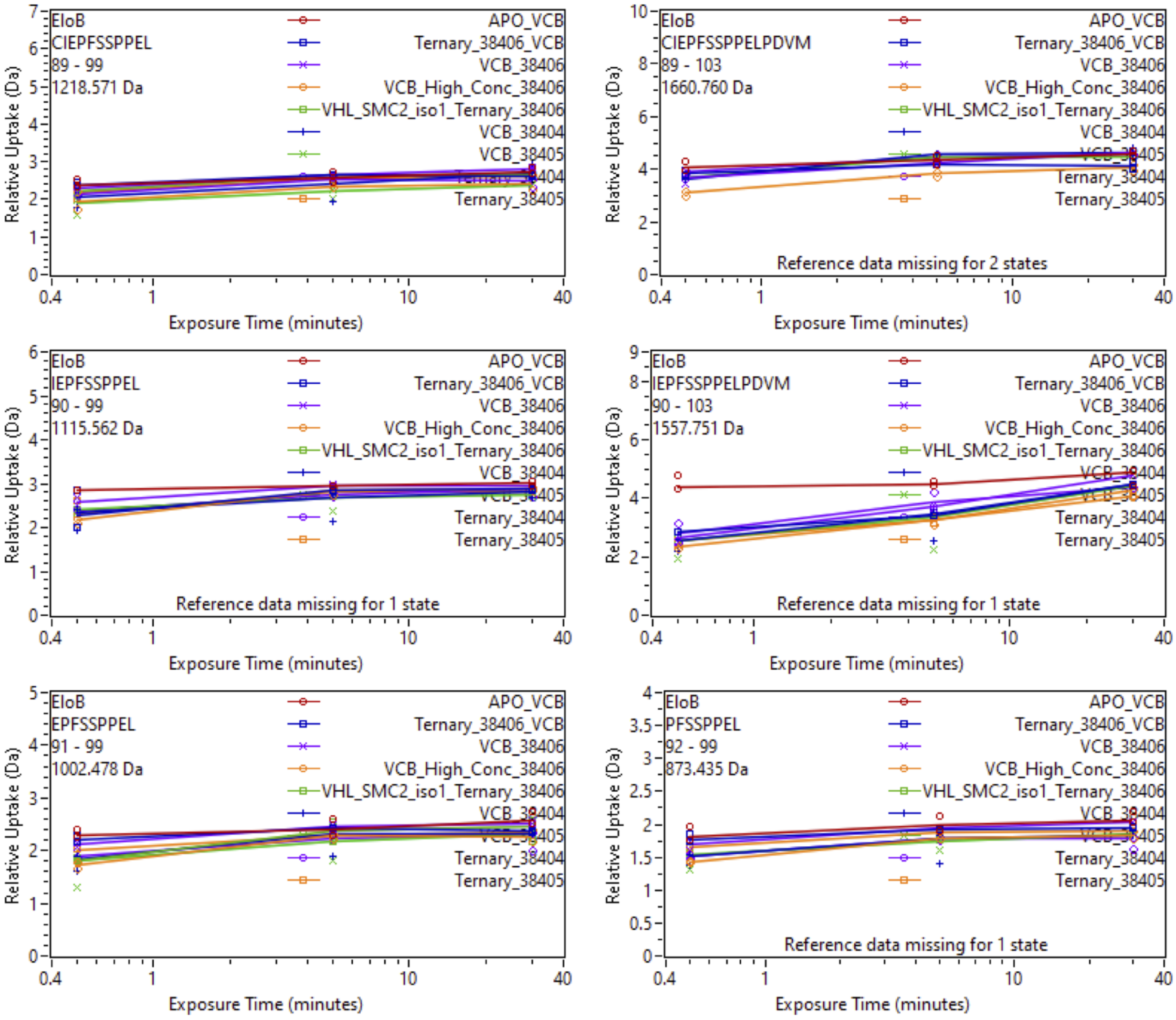


Supplemental Fig. 64: Relative deuterium uptake plots of peptic peptides Elongin B of the VCB complex in the APO, Binary with SiTX-0038404 (PROTAC 1), SiTX-0038405(PROTAC 2), SiTX-0038406(ACBI1) or Ternary complex with 404, 405, 406 + SMARCA2$^{BD}$.

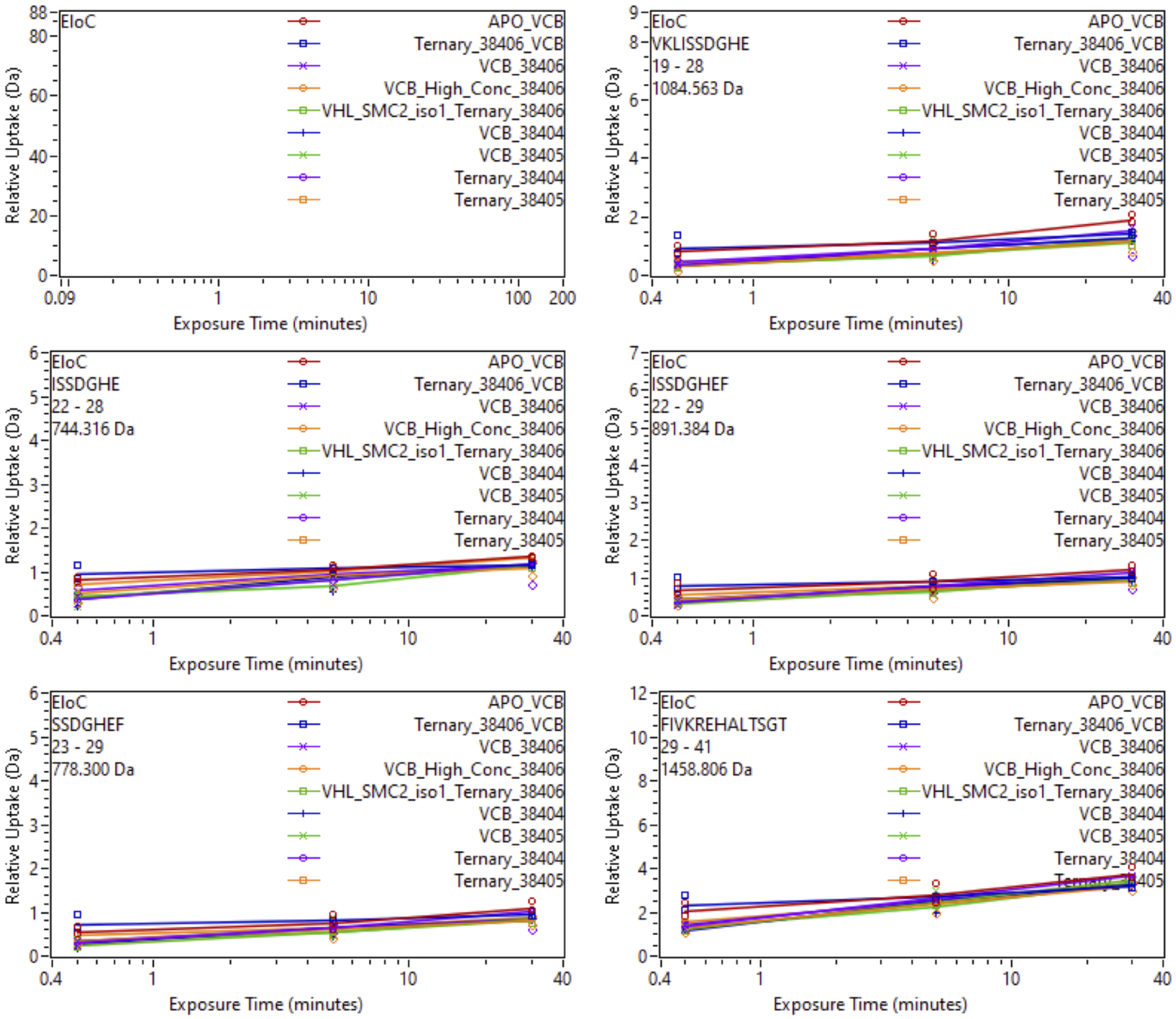


Supplemental Fig. 65: Relative deuterium uptake plots of peptic peptides of Elongin C of the VCB complex in the APO, Binary with SiTX-0038404 (PROTAC 1), SiTX-0038405(PROTAC 2), SiTX-0038406(ACBI1) or Ternary complex with 404, 405, 406 + SMARCA2$^{BD}$.

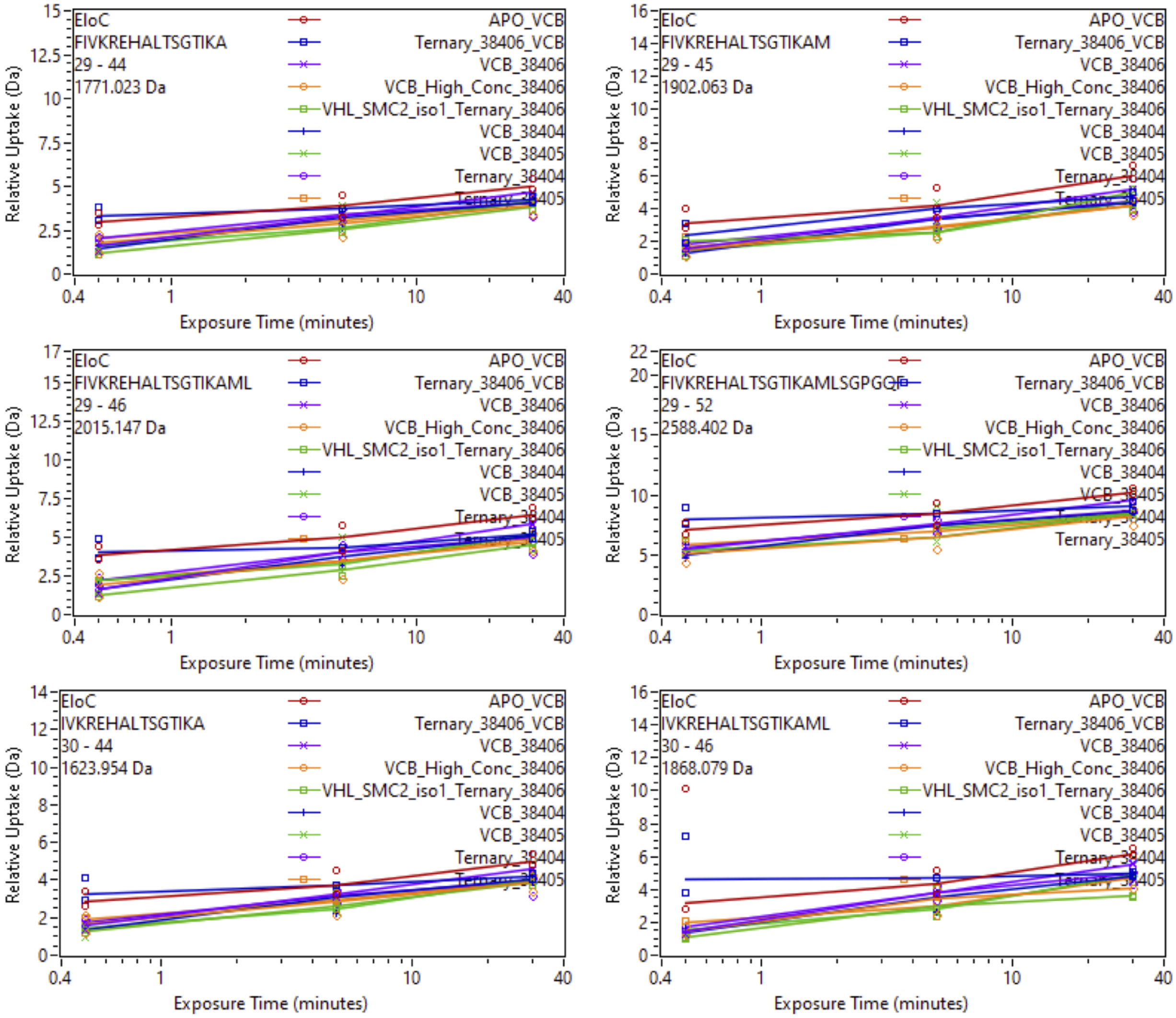


Supplemental Fig. 66: Relative deuterium uptake plots of peptic peptides of Elongin C of the VCB complex in the APO, Binary with SiTX-0038404 (PROTAC 1), SiTX-0038405(PROTAC 2), SiTX-0038406(ACBI1) or Ternary complex with 404, 405, 406 + SMARCA2$^{BD}$.

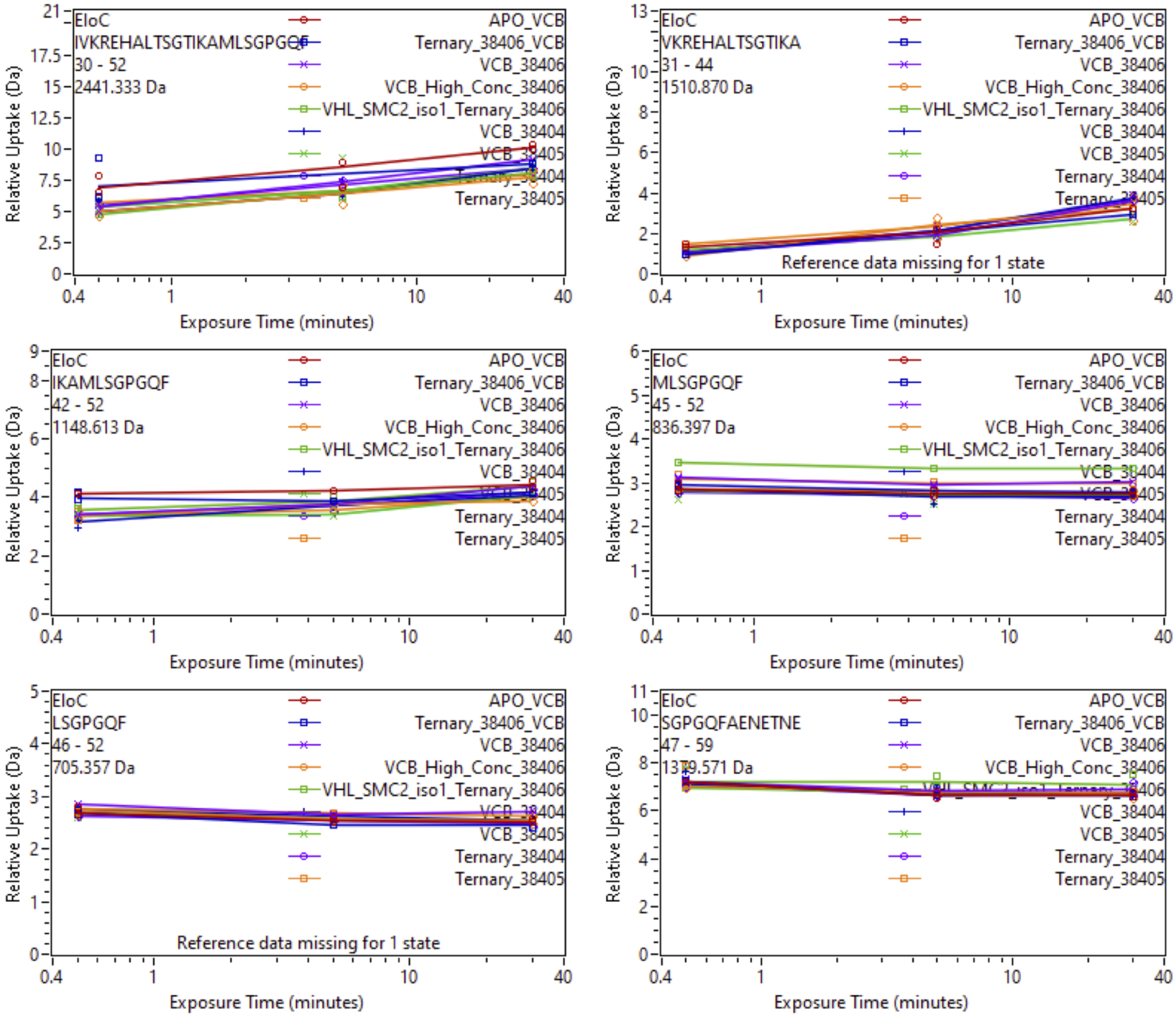


Supplemental Fig. 67: Relative deuterium uptake plots of peptic peptides of Elongin C of the VCB complex in the APO, Binary with SiTX-0038404 (PROTAC 1), SiTX-0038405(PROTAC 2), SiTX-0038406(ACBI1) or Ternary complex with 404, 405, 406 + SMARCA2$^{BD}$.

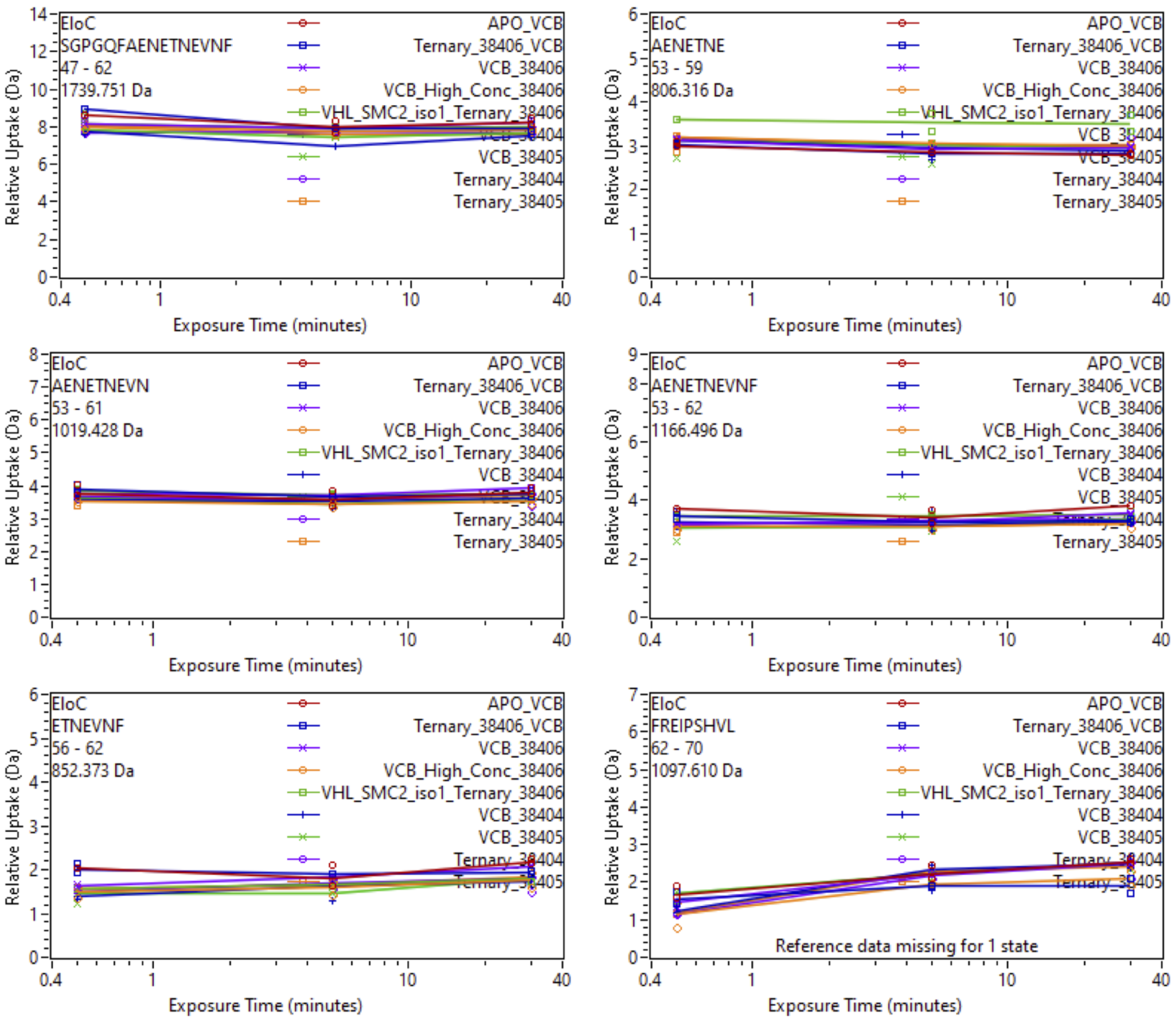


Supplemental Fig. 68: Relative deuterium uptake plots of peptic peptides of Elongin C of the VCB complex in the APO, Binary with SiTX-0038404 (PROTAC 1), SiTX-0038405(PROTAC 2), SiTX-0038406(ACBI1) or Ternary complex with 404, 405, 406 + SMARCA2$^{BD}$.

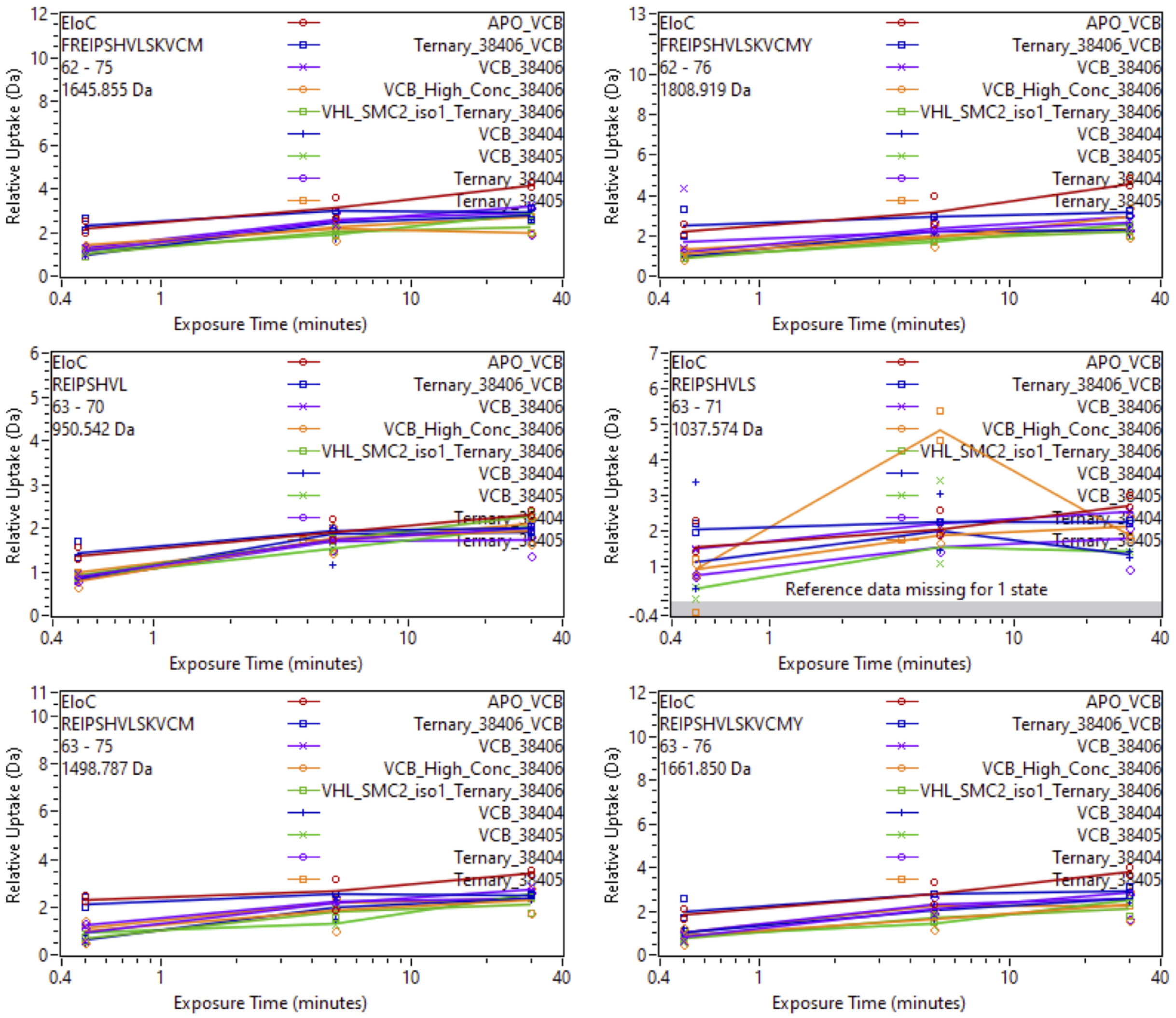


Supplemental Fig. 69: Relative deuterium uptake plots of peptic peptides of Elongin C of the VCB complex in the APO, Binary with SiTX-0038404 (PROTAC 1), SiTX-0038405(PROTAC 2), SiTX-0038406(ACBI1) or Ternary complex with 404, 405, 406 + SMARCA2$^{BD}$.

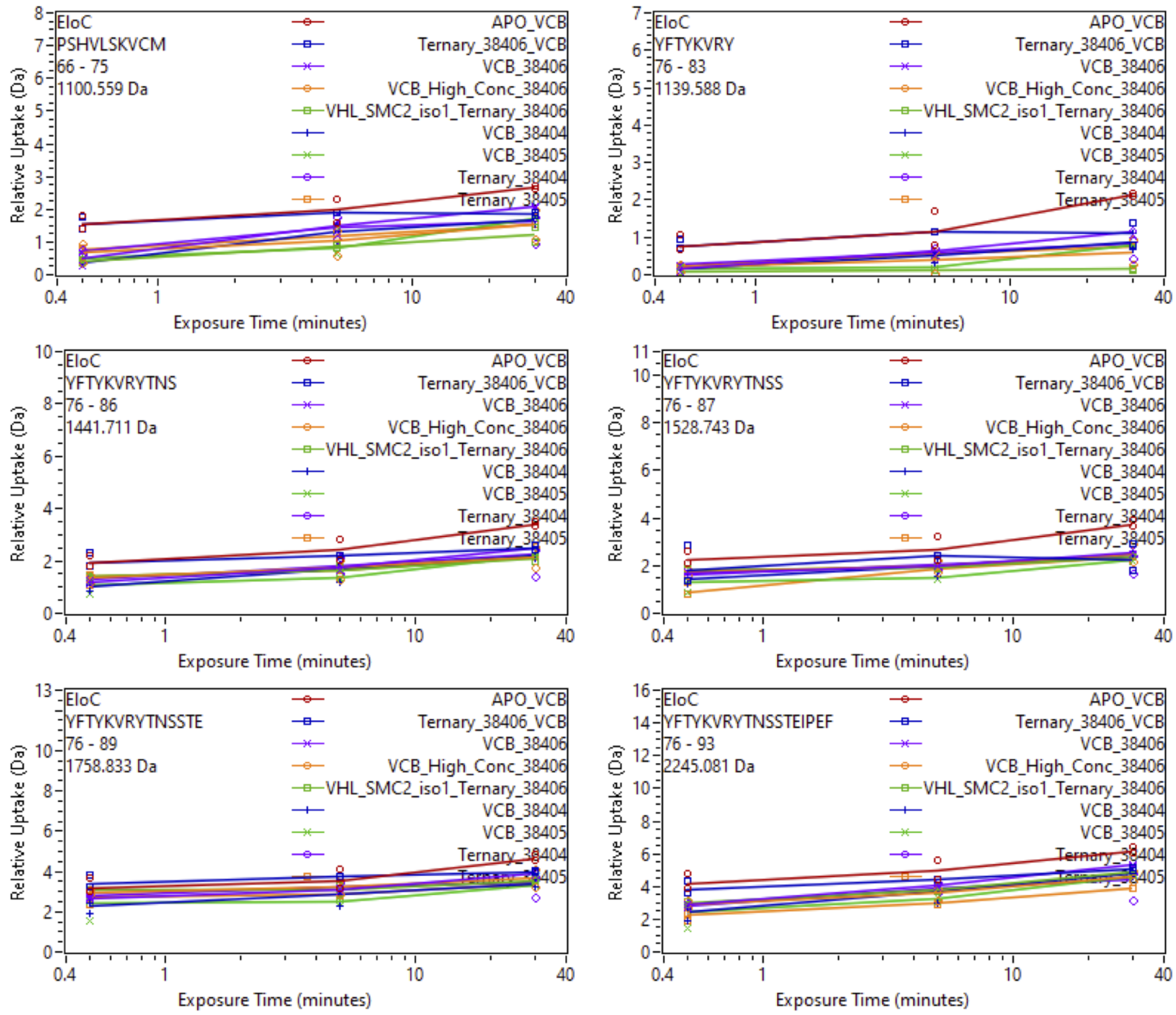


Supplemental Fig. 70: Relative deuterium uptake plots of peptic peptides of Elongin C of the VCB complex in the APO, Binary with SiTX-0038404 (PROTAC 1), SiTX-0038405(PROTAC 2), SiTX-0038406(ACBI1) or Ternary complex with 404, 405, 406 + SMARCA2$^{BD}$.

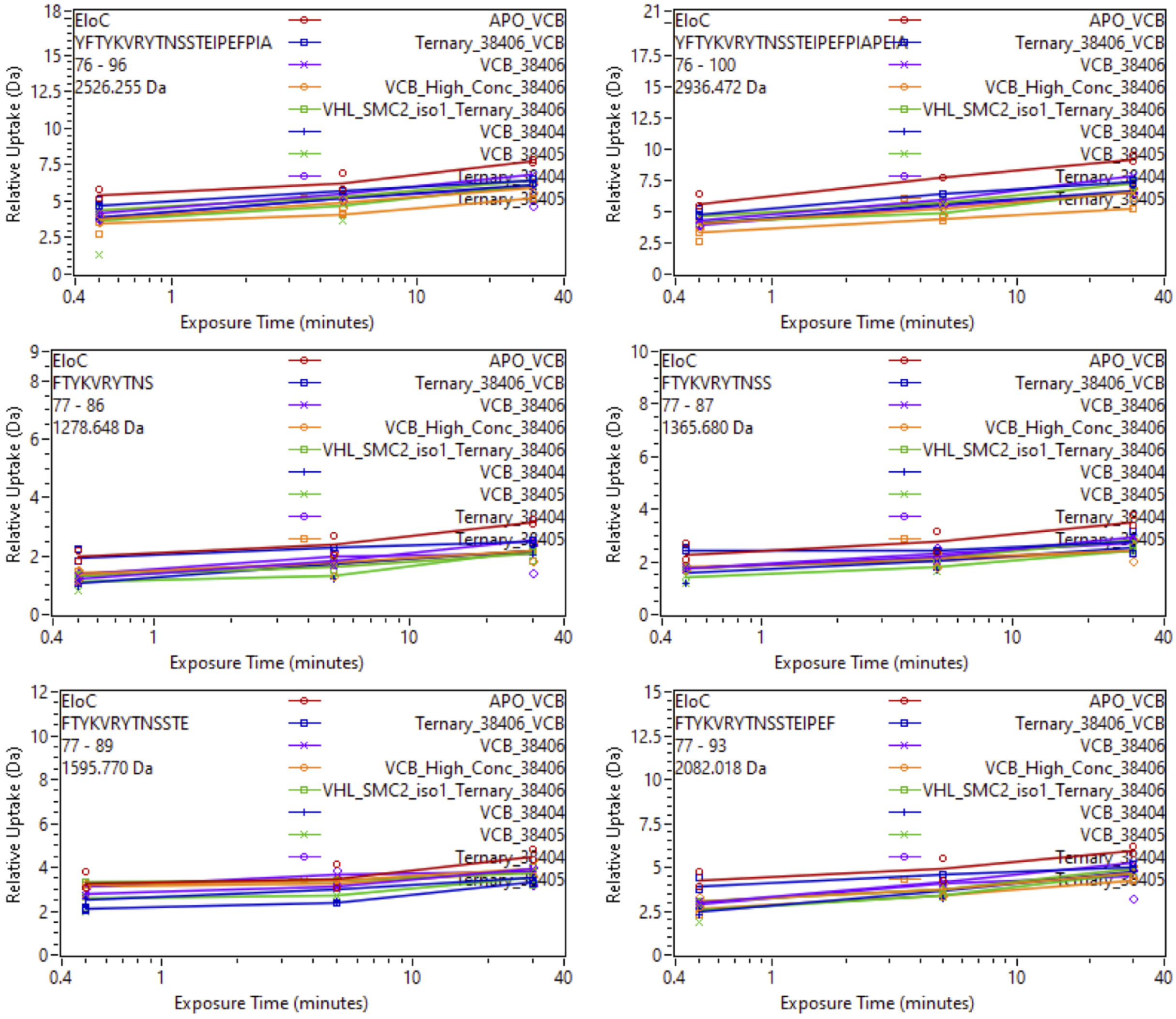


Supplemental Fig. 71: Relative deuterium uptake plots of peptic peptides of Elongin C of the VCB complex in the APO, Binary with SiTX-0038404 (PROTAC 1), SiTX-0038405(PROTAC 2), SiTX-0038406(ACBI1) or Ternary complex with 404, 405, 406 + SMARCA2$^{BD}$.

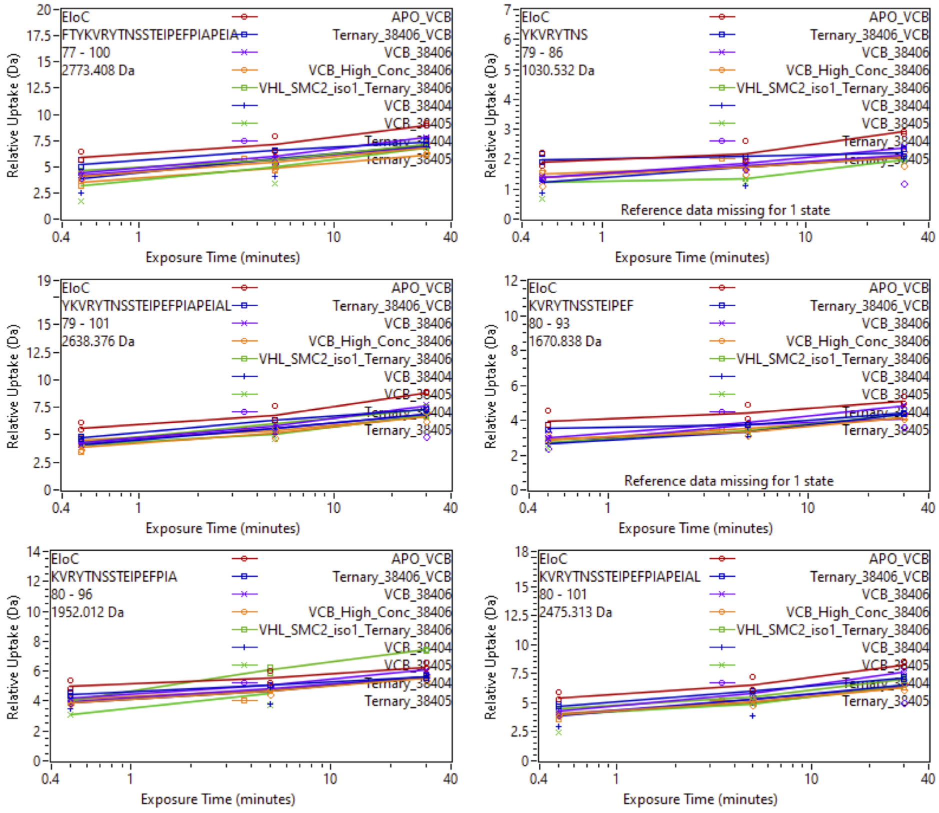




Supplemental Fig. 72: Relative deuterium uptake plots of peptic peptides of Elongin C of the VCB complex in the APO, Binary with SiTX-0038404 (PROTAC 1), SiTX-0038405(PROTAC 2), SiTX-0038406(ACBI1) or Ternary complex with 404, 405, 406 + SMARCA2$^{BD}$.

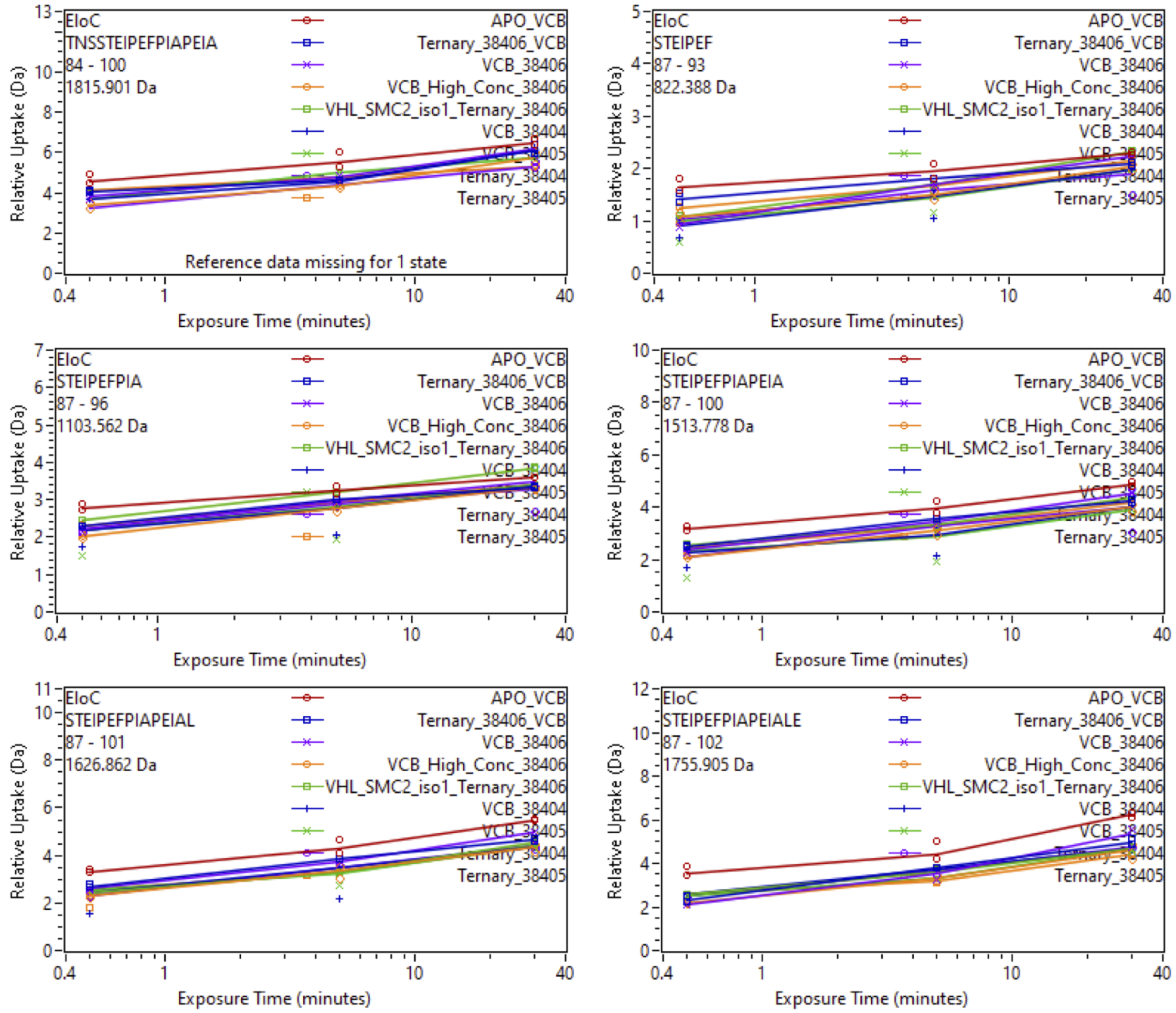


Supplemental Fig. 73: Relative deuterium uptake plots of peptic peptides of Elongin C of the VCB complex in the APO, Binary with SiTX-0038404 (PROTAC 1), SiTX-0038405(PROTAC 2), SiTX-0038406(ACBI1) or Ternary complex with 404, 405, 406 + SMARCA2$^{BD}$.

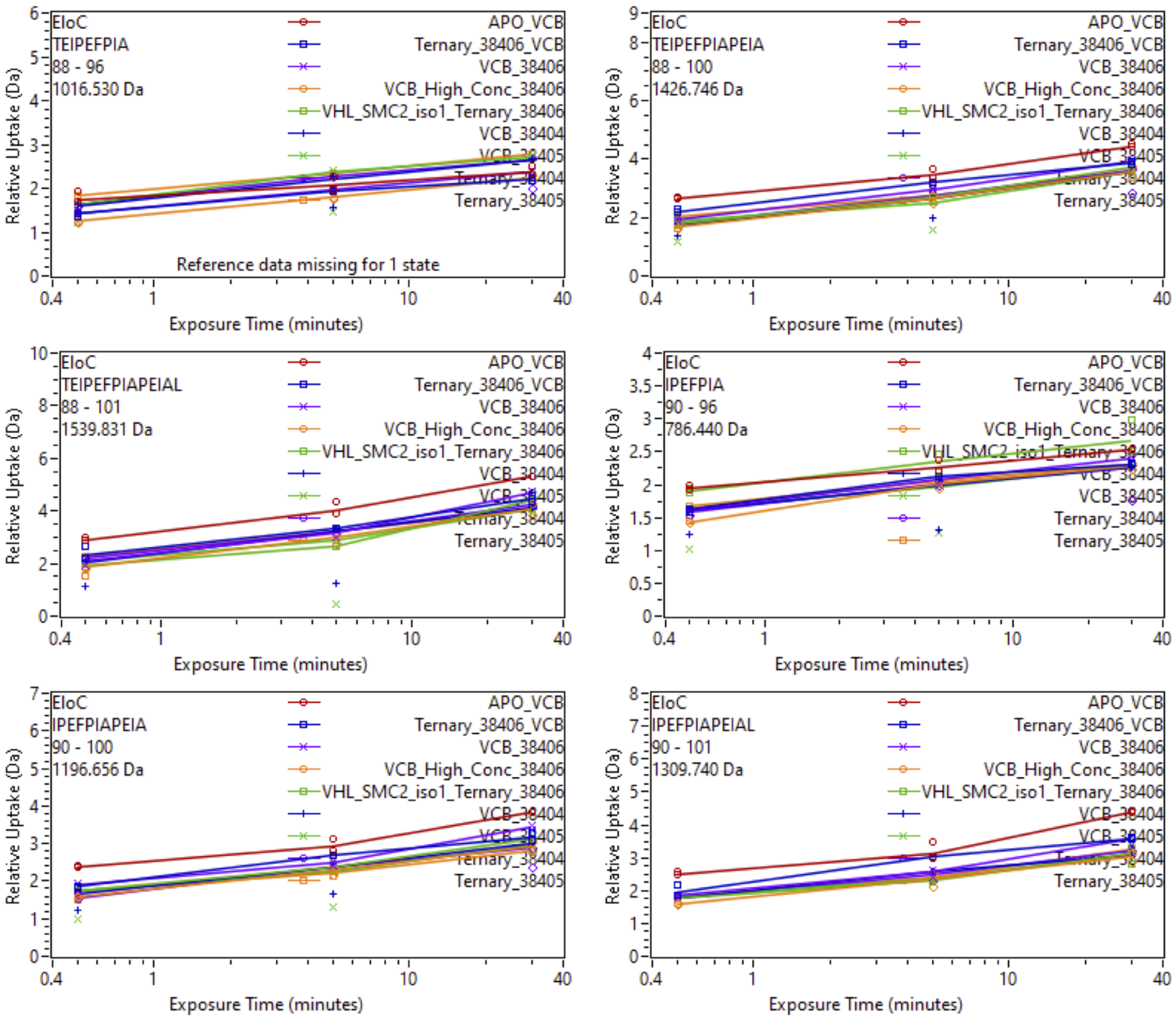


Supplemental Fig. 74: Relative deuterium uptake plots of peptic peptides of Elongin C of the VCB complex in the APO, Binary with SiTX-0038404 (PROTAC 1), SiTX-0038405(PROTAC 2), SiTX-0038406(ACBI1) or Ternary complex with 404, 405, 406 + SMARCA2$^{BD}$.

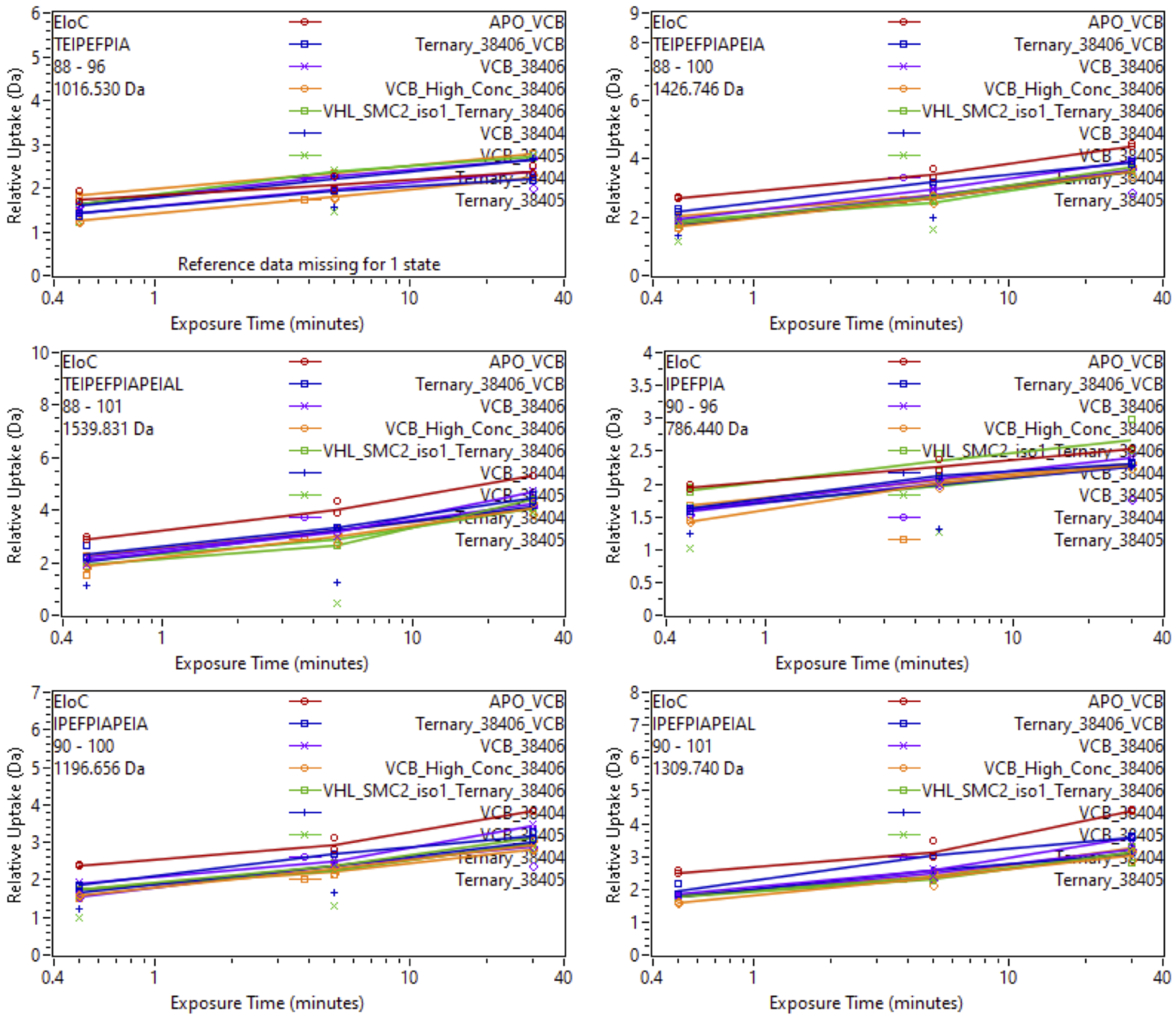


Supplemental Fig. 75: Relative deuterium uptake plots of peptic peptides of VHL of the VCB complex in the APO, Binary with SiTX-0038404 (PROTAC 1), SiTX-0038405(PROTAC 2), SiTX-0038406(ACBI1) or Ternary complex with 404, 405, 406 + SMARCA2$^{BD}$.

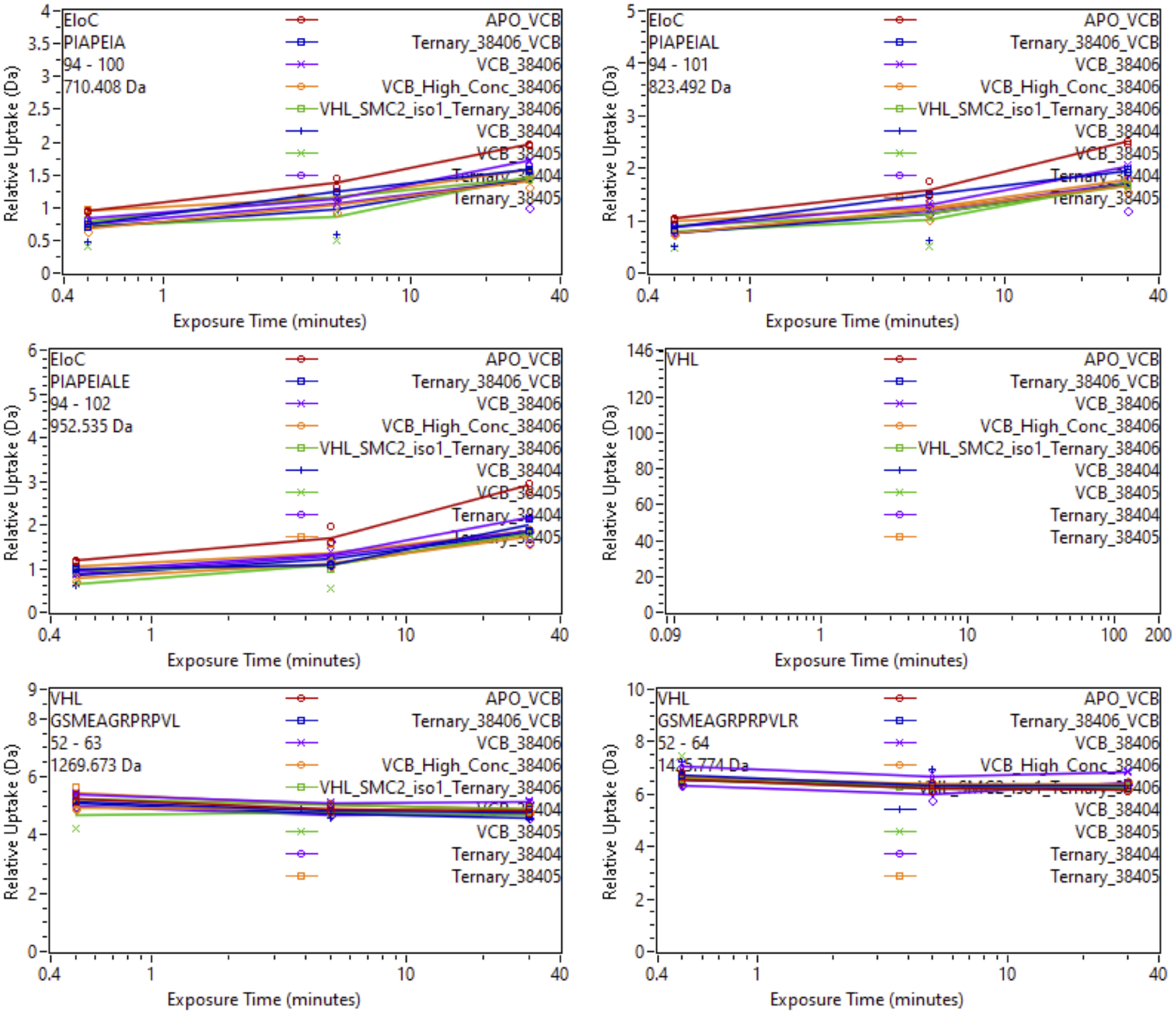


Supplemental Fig. 76: Relative deuterium uptake plots of peptic peptides of VHL of the VCB complex in the APO, Binary with SiTX-0038404 (PROTAC 1), SiTX-0038405(PROTAC 2), SiTX-0038406(ACBI1) or Ternary complex with 404, 405, 406 + SMARCA2$^{BD}$.

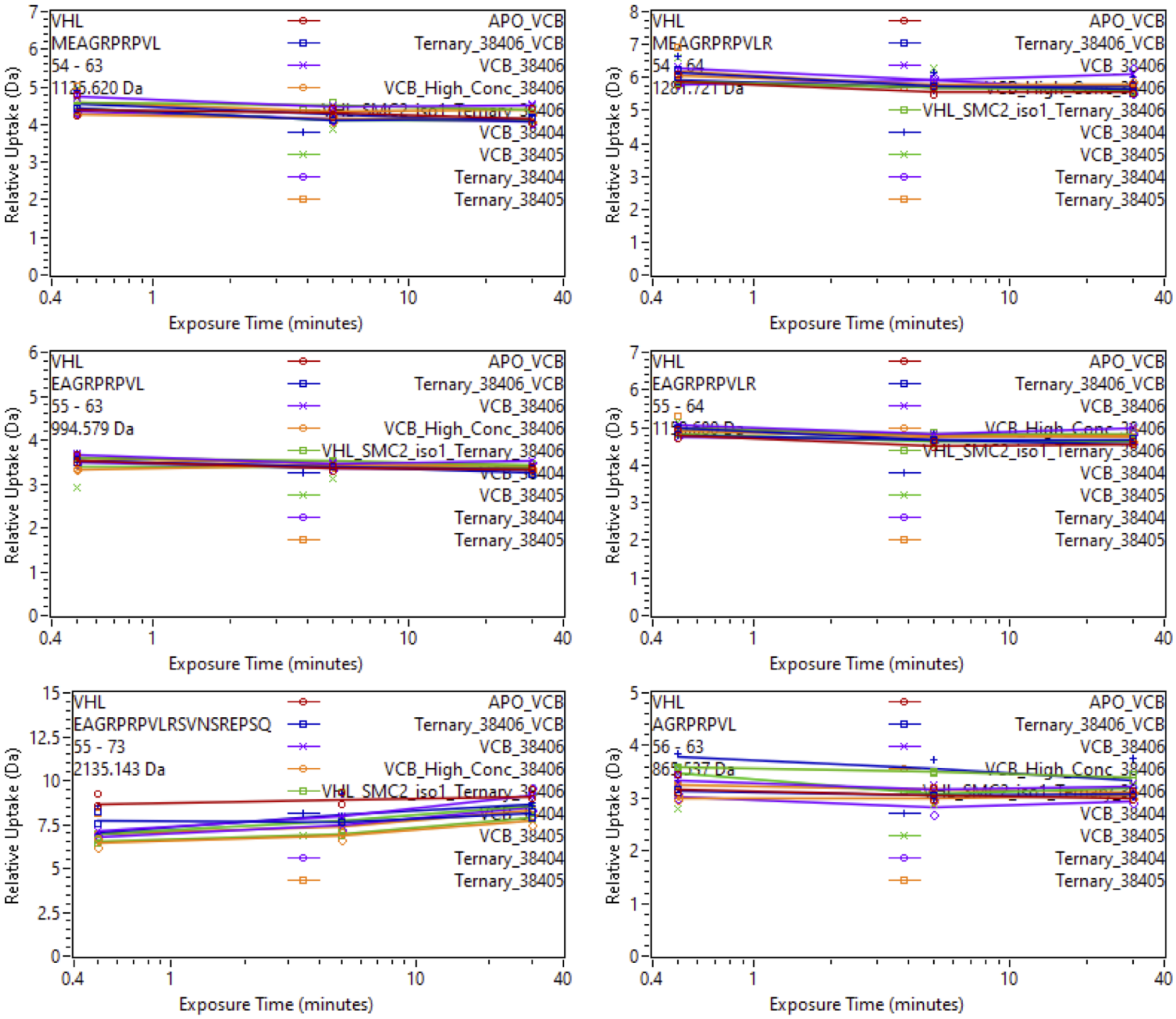


Supplemental Fig. 77: Relative deuterium uptake plots of peptic peptides of VHL of the VCB complex in the APO, Binary with SiTX-0038404 (PROTAC 1), SiTX-0038405(PROTAC 2), SiTX-0038406(ACBI1) or Ternary complex with 404, 405, 406 + SMARCA2$^{BD}$.

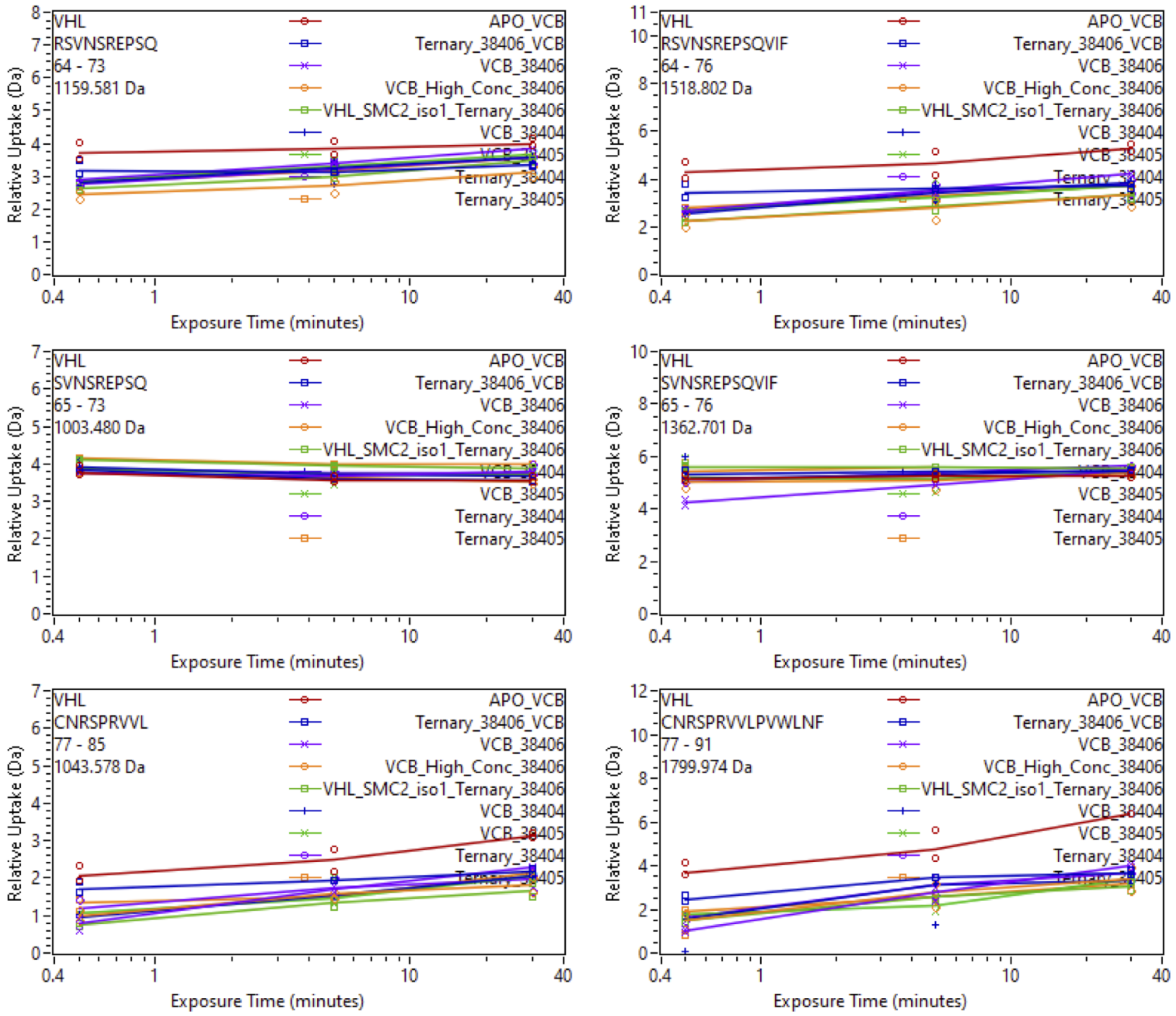


Supplemental Fig. 78: Relative deuterium uptake plots of peptic peptides of VHL of the VCB complex in the APO, Binary with SiTX-0038404 (PROTAC 1), SiTX-0038405(PROTAC 2), SiTX-0038406(ACBI1) or Ternary complex with 404, 405, 406 + SMARCA2$^{BD}$.

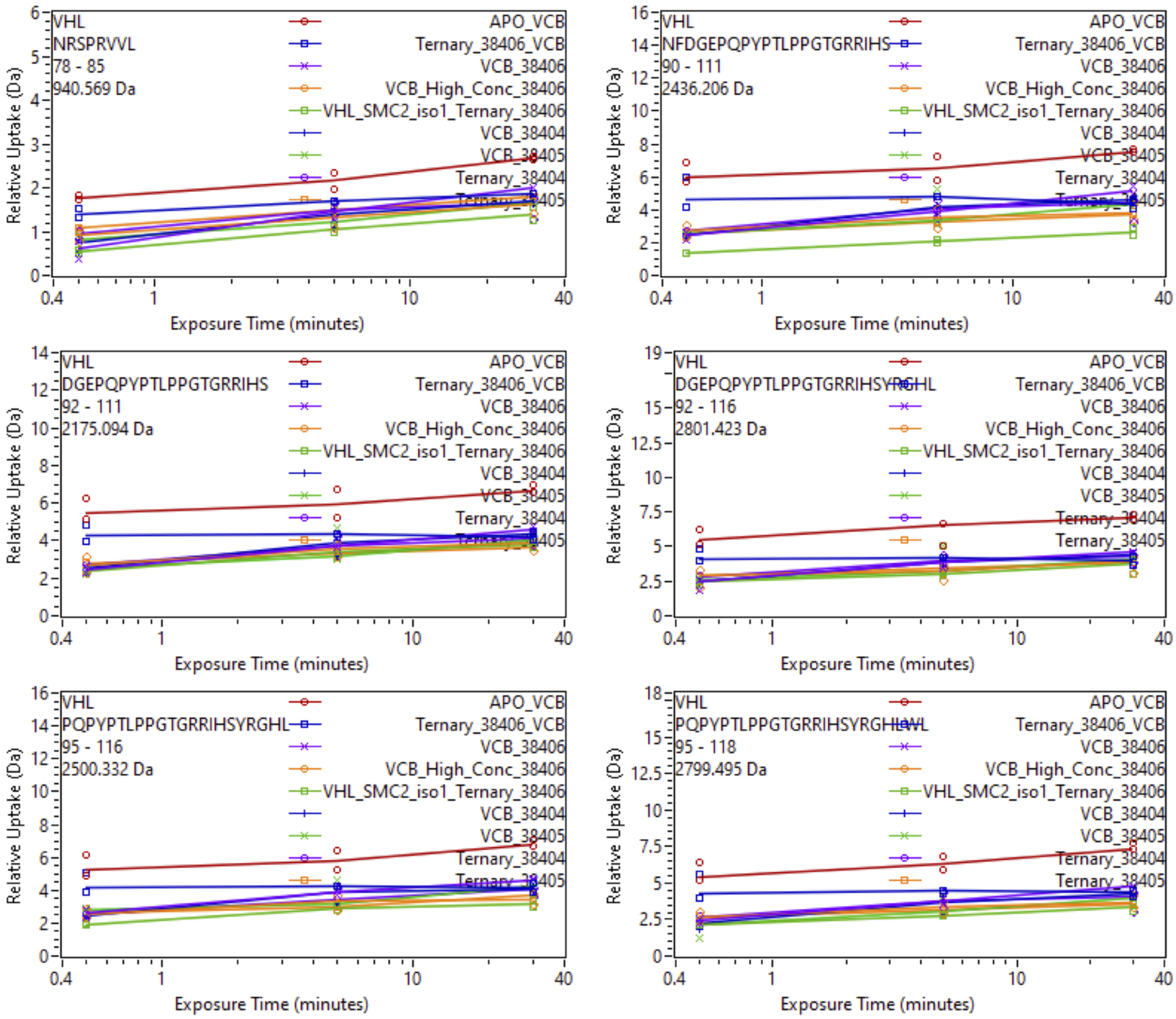


Supplemental Fig. 79: Relative deuterium uptake plots of peptic peptides of VHL of the VCB complex in the APO, Binary with SiTX-0038404 (PROTAC 1), SiTX-0038405(PROTAC 2), SiTX-0038406(ACBI1) or Ternary complex with 404, 405, 406 + SMARCA2$^{BD}$.

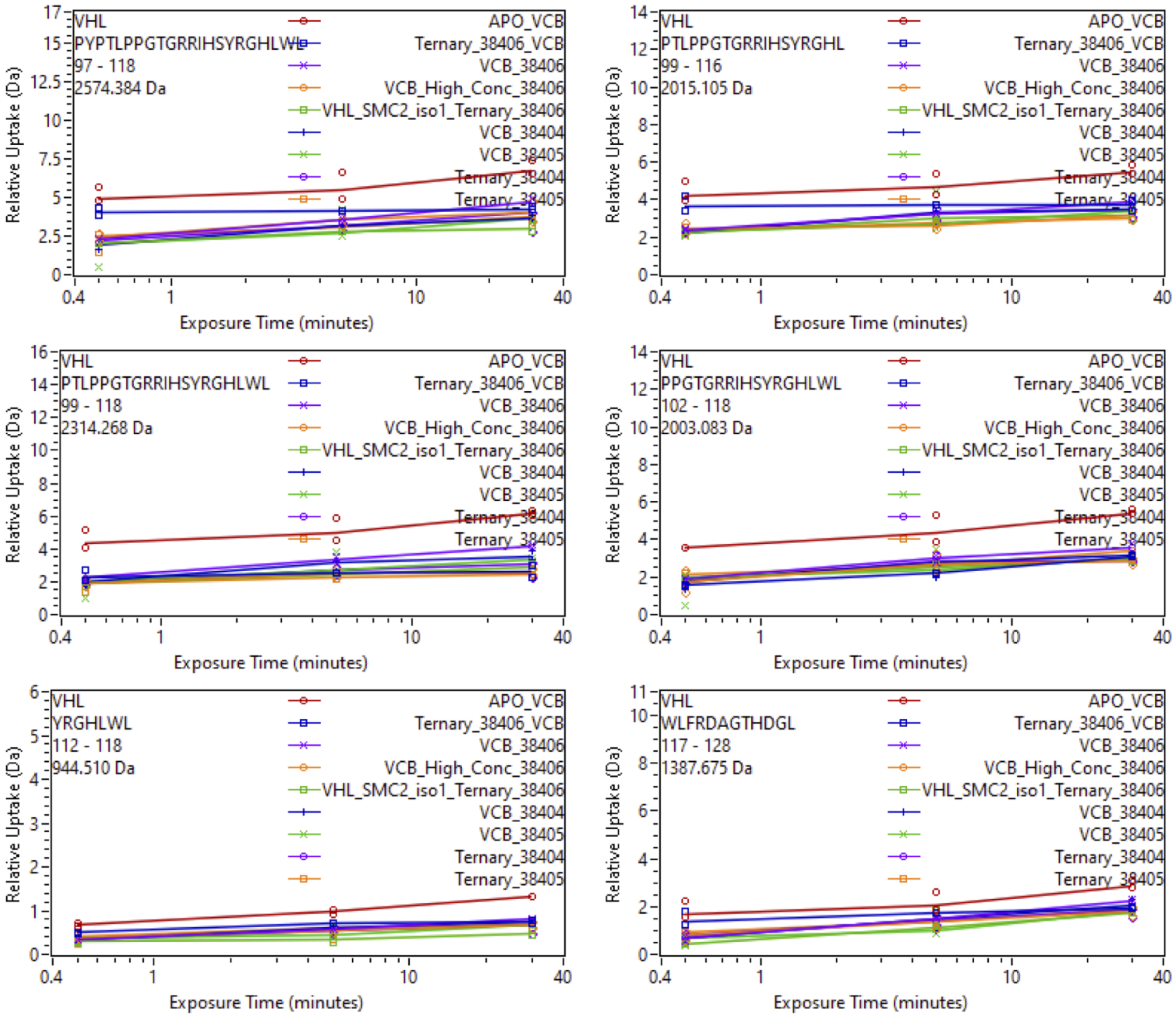


Supplemental Fig. 80: Relative deuterium uptake plots of peptic peptides of VHL of the VCB complex in the APO, Binary with SiTX-0038404 (PROTAC 1), SiTX-0038405(PROTAC 2), SiTX-0038406(ACBI1) or Ternary complex with 404, 405, 406 + SMARCA2$^{BD}$.

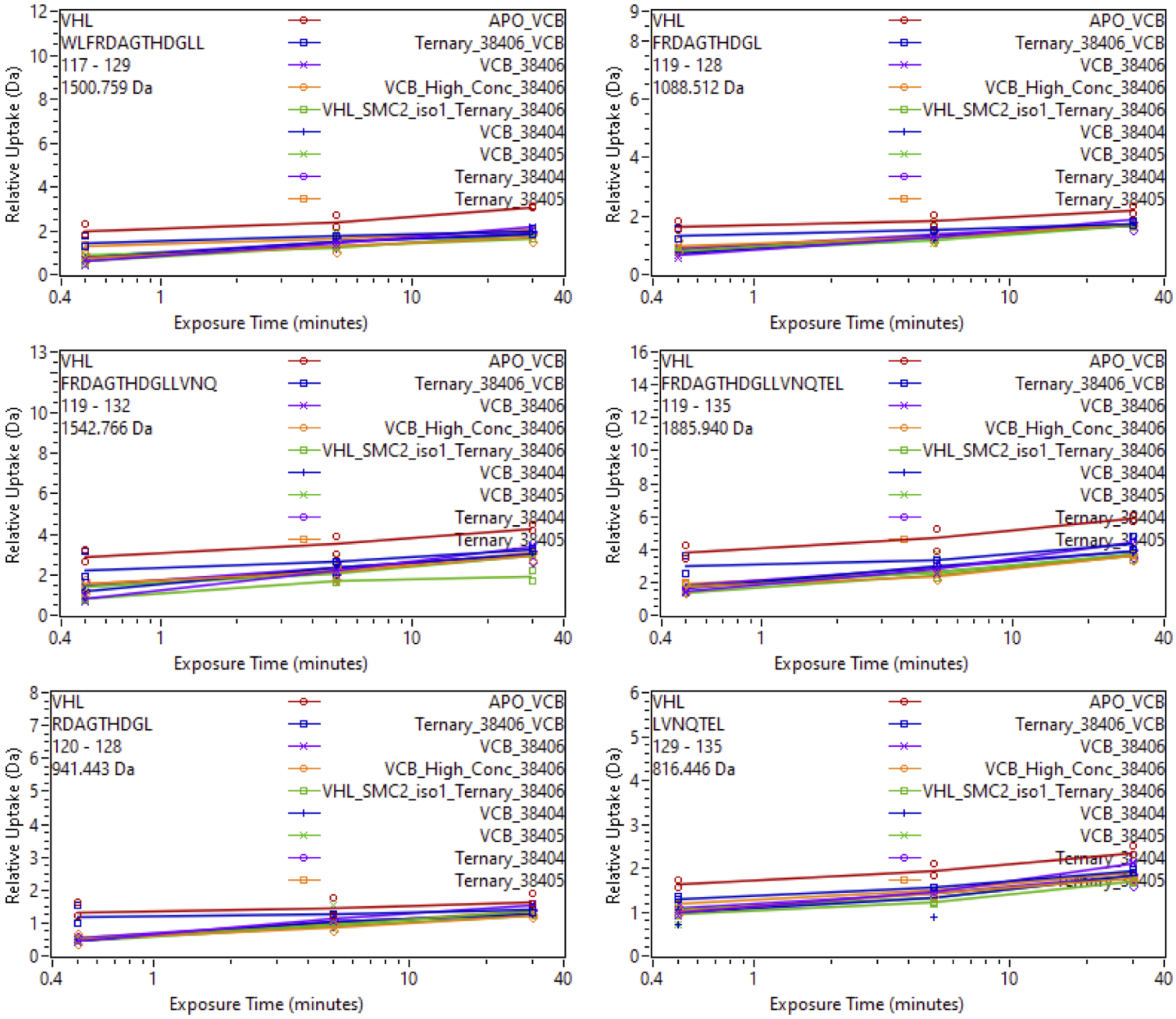


Supplemental Fig. 81: Relative deuterium uptake plots of peptic peptides of VHL of the VCB complex in the APO, Binary with SiTX-0038404 (PROTAC 1), SiTX-0038405(PROTAC 2), SiTX-0038406(ACBI1) or Ternary complex with 404, 405, 406 + SMARCA2$^{BD}$.

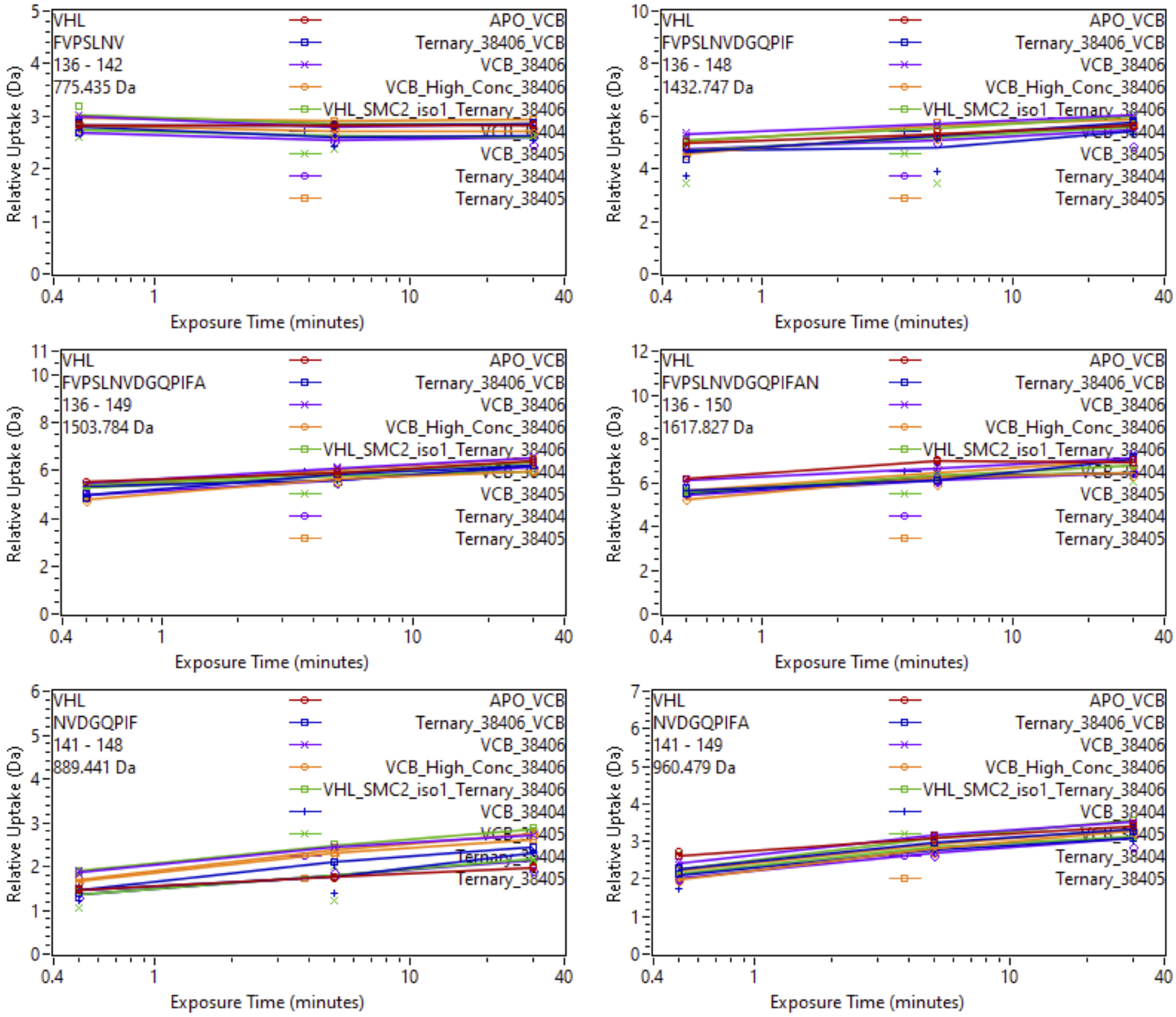


Supplemental Fig. 82: Relative deuterium uptake plots of peptic peptides of VHL of the VCB complex in the APO, Binary with SiTX-0038404 (PROTAC 1), SiTX-0038405(PROTAC 2), SiTX-0038406(ACBI1) or Ternary complex with 404, 405, 406 + SMARCA2$^{BD}$.

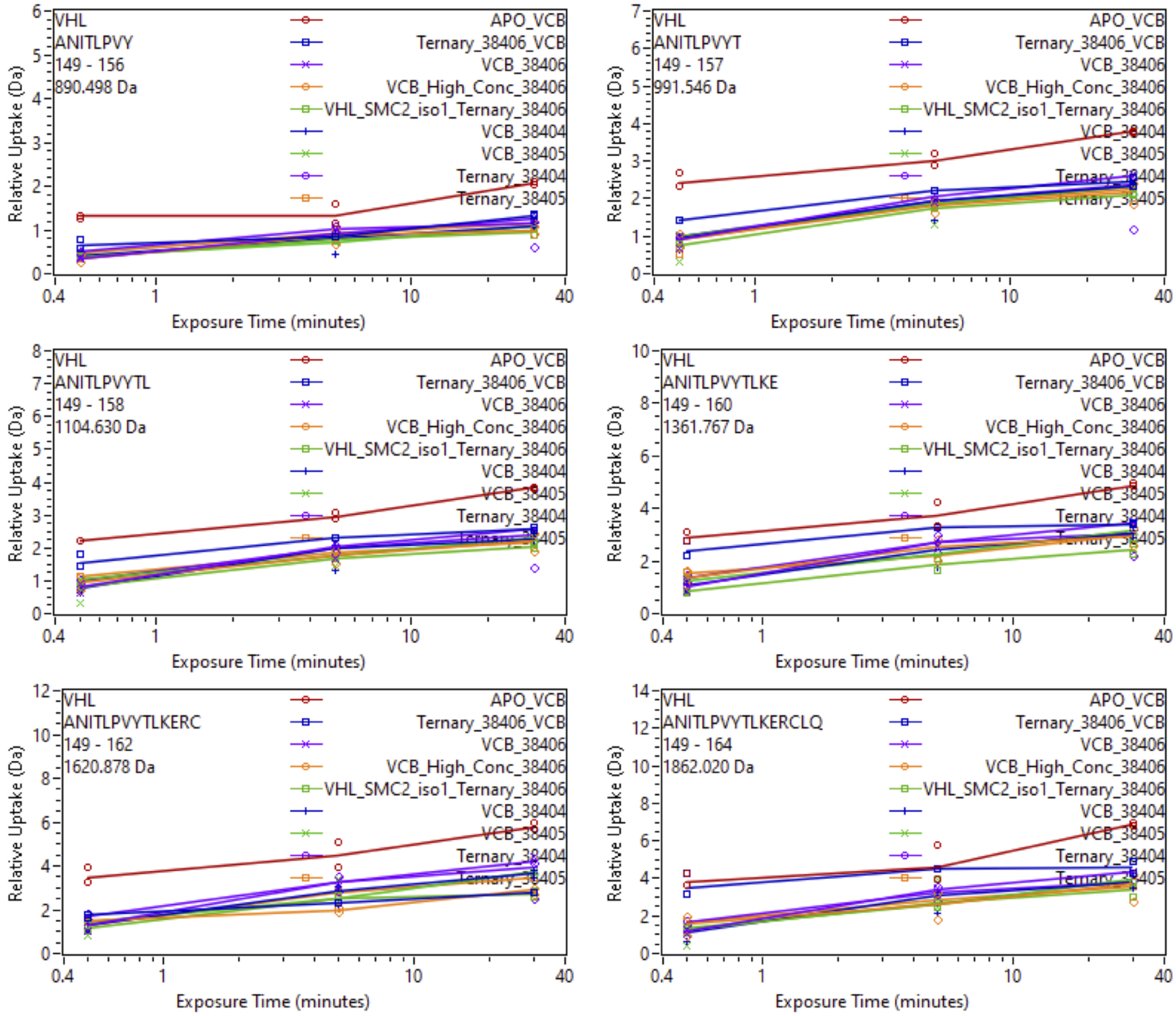


Supplemental Fig. 83: Relative deuterium uptake plots of peptic peptides of VHL of the VCB complex in the APO, Binary with SiTX-0038404 (PROTAC 1), SiTX-0038405(PROTAC 2), SiTX-0038406(ACBI1) or Ternary complex with 404, 405, 406 + SMARCA2$^{BD}$.

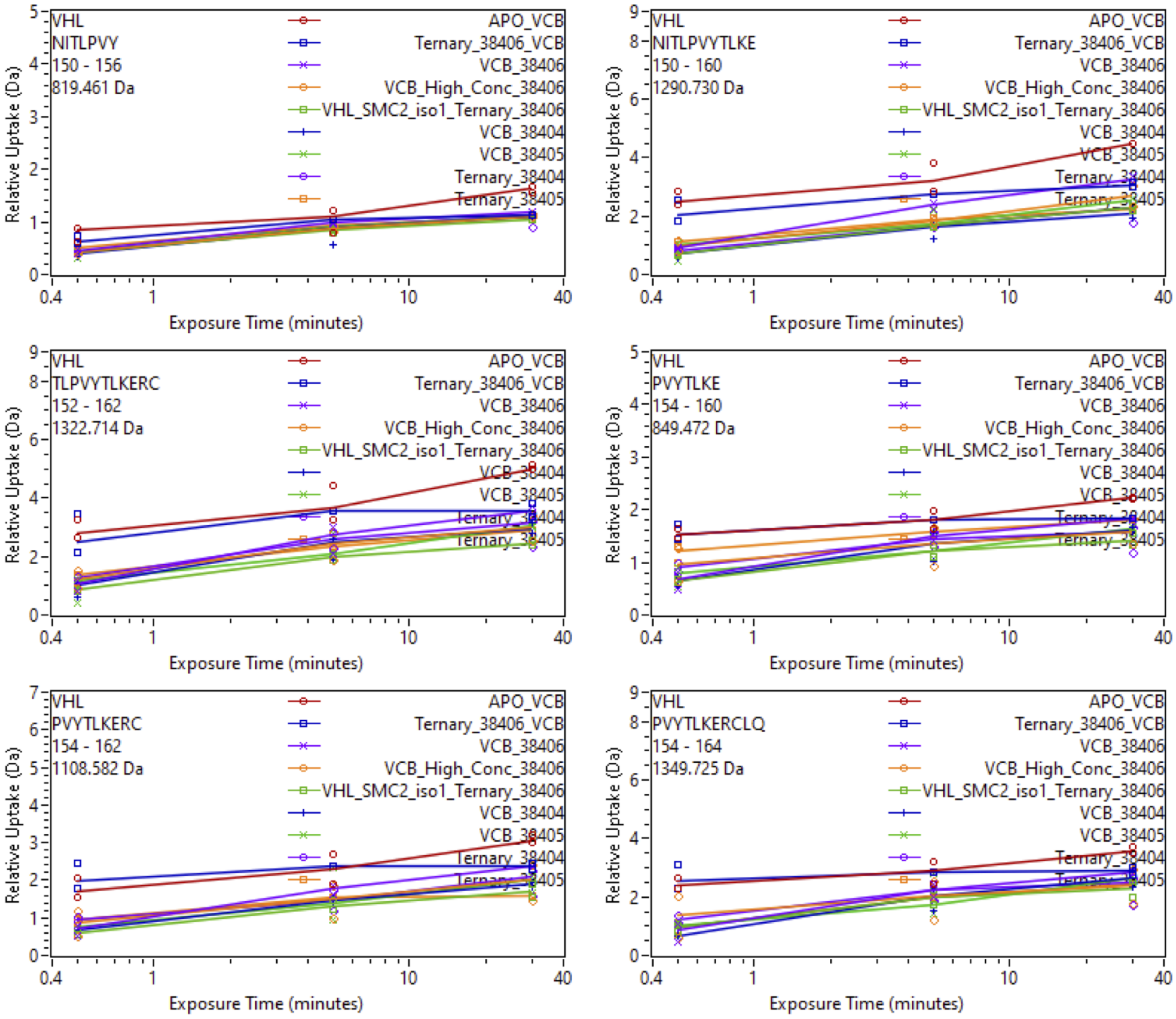


Supplemental Fig. 84: Relative deuterium uptake plots of peptic peptides of VHL of the VCB complex in the APO, Binary with SiTX-0038404 (PROTAC 1), SiTX-0038405(PROTAC 2), SiTX-0038406(ACBI1) or Ternary complex with 404, 405, 406 + SMARCA2$^{BD}$.

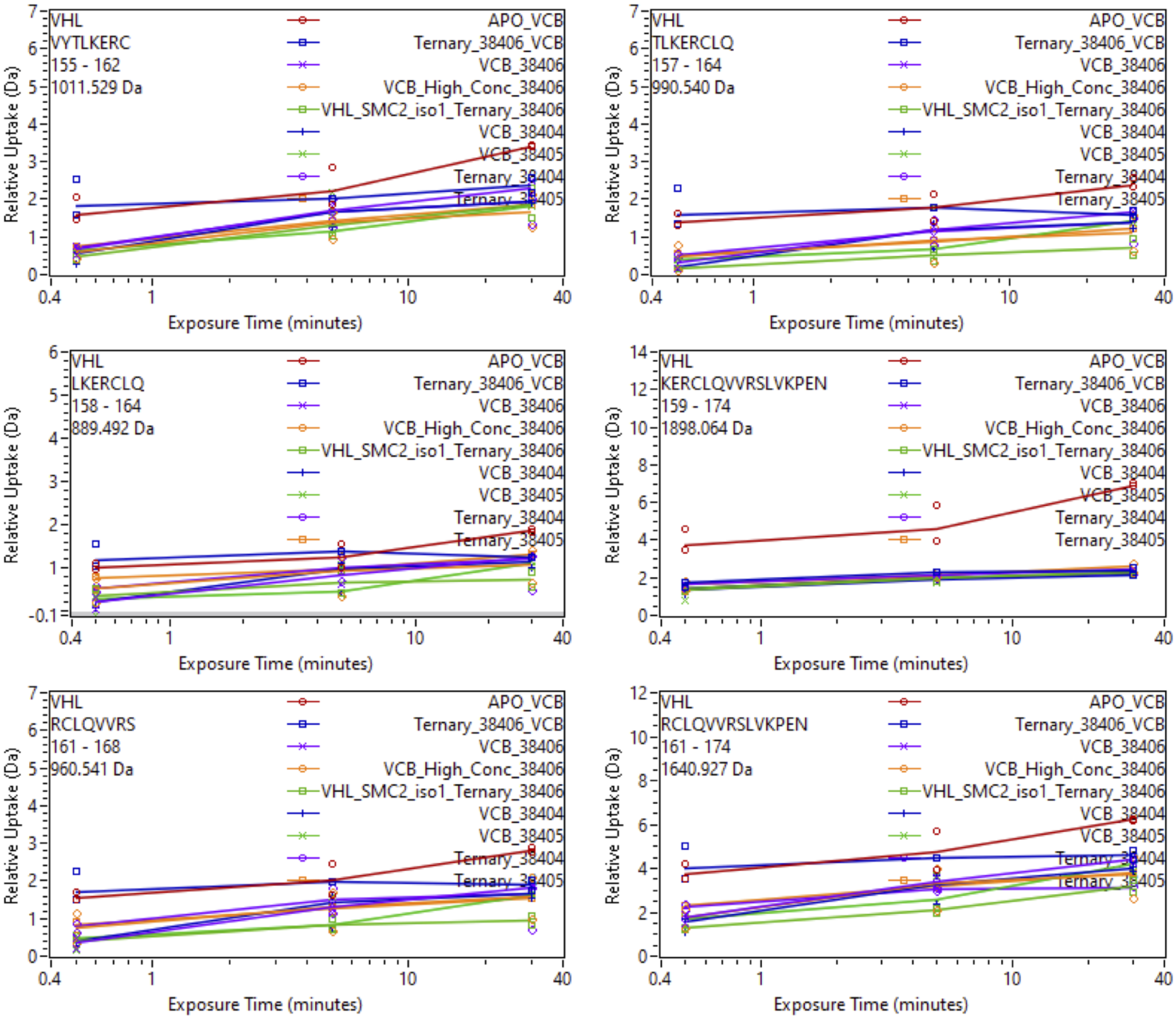


Supplemental Fig. 85: Relative deuterium uptake plots of peptic peptides of VHL of the VCB complex in the APO, Binary with SiTX-0038404 (PROTAC 1), SiTX-0038405(PROTAC 2), SiTX-0038406(ACBI1) or Ternary complex with 404, 405, 406 + SMARCA2$^{BD}$.

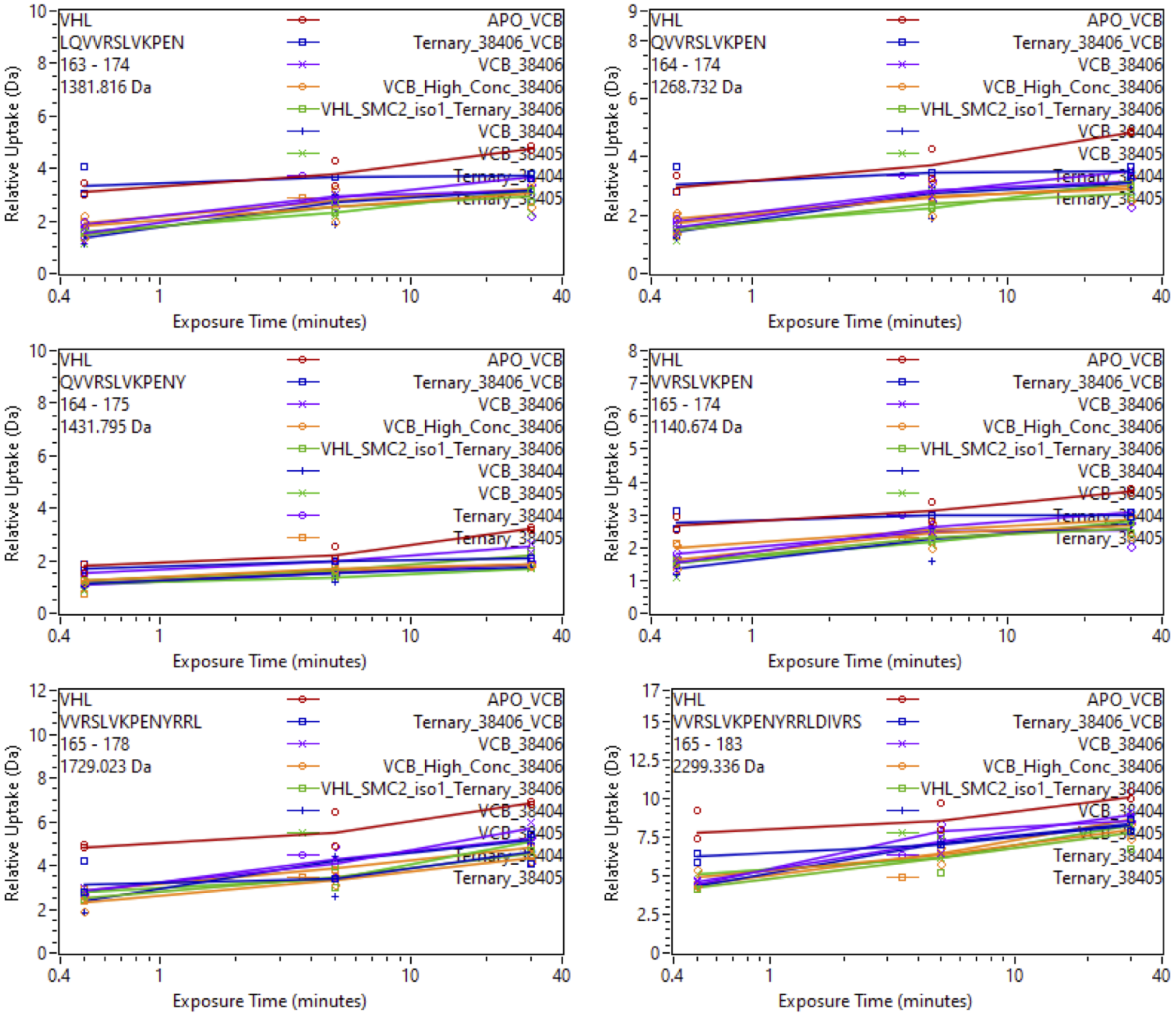


Supplemental Fig. 86: Relative deuterium uptake plots of peptic peptides of VHL of the VCB complex in the APO, Binary with SiTX-0038404 (PROTAC 1), SiTX-0038405(PROTAC 2), SiTX-0038406(ACBI1) or Ternary complex with 404, 405, 406 + SMARCA2$^{BD}$.

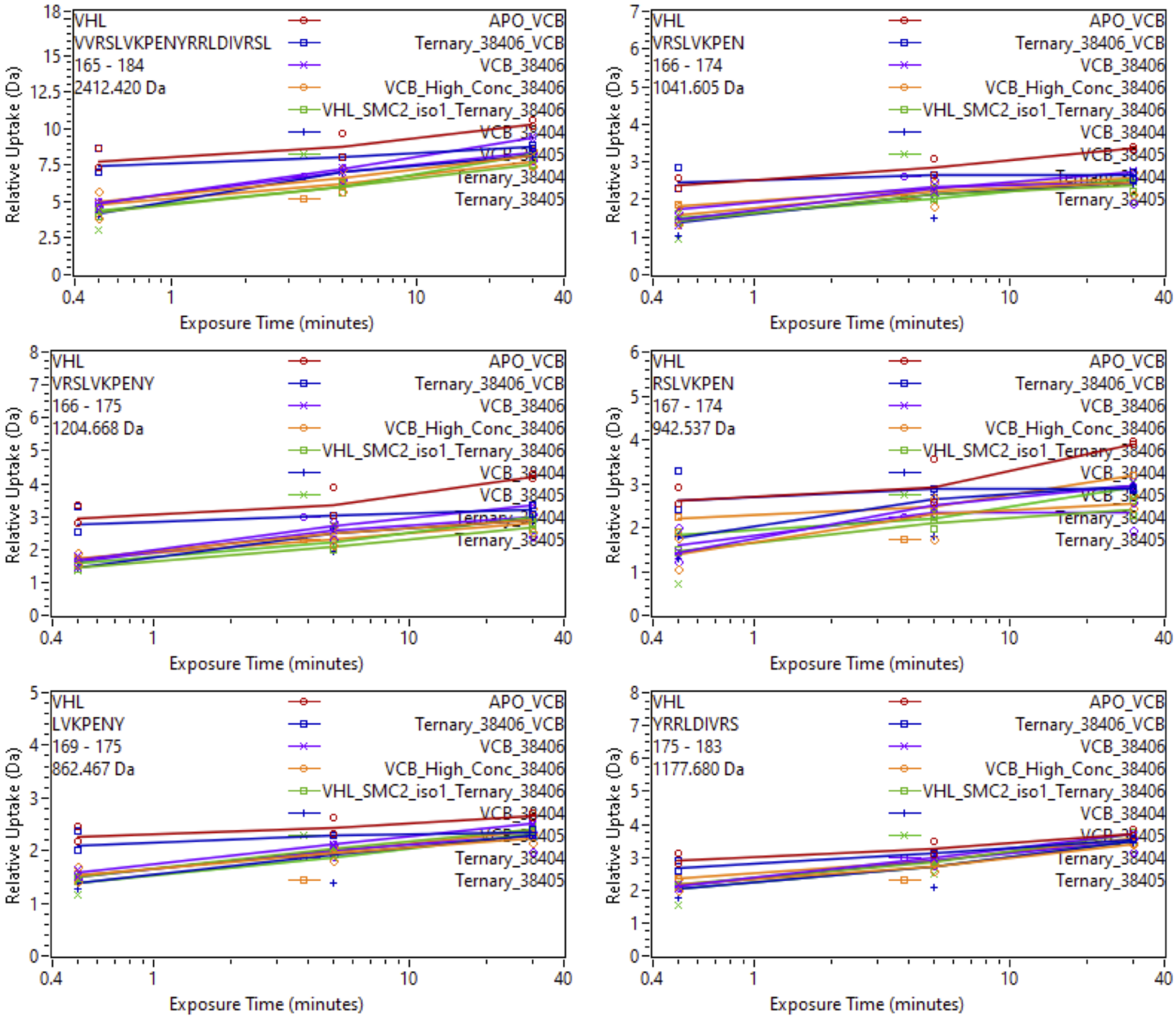


Supplemental Fig. 87: Relative deuterium uptake plots of peptic peptides of VHL of the VCB complex in the APO, Binary with SiTX-0038404 (PROTAC 1), SiTX-0038405(PROTAC 2), SiTX-0038406(ACBI1) or Ternary complex with 404, 405, 406 + SMARCA2$^{BD}$.

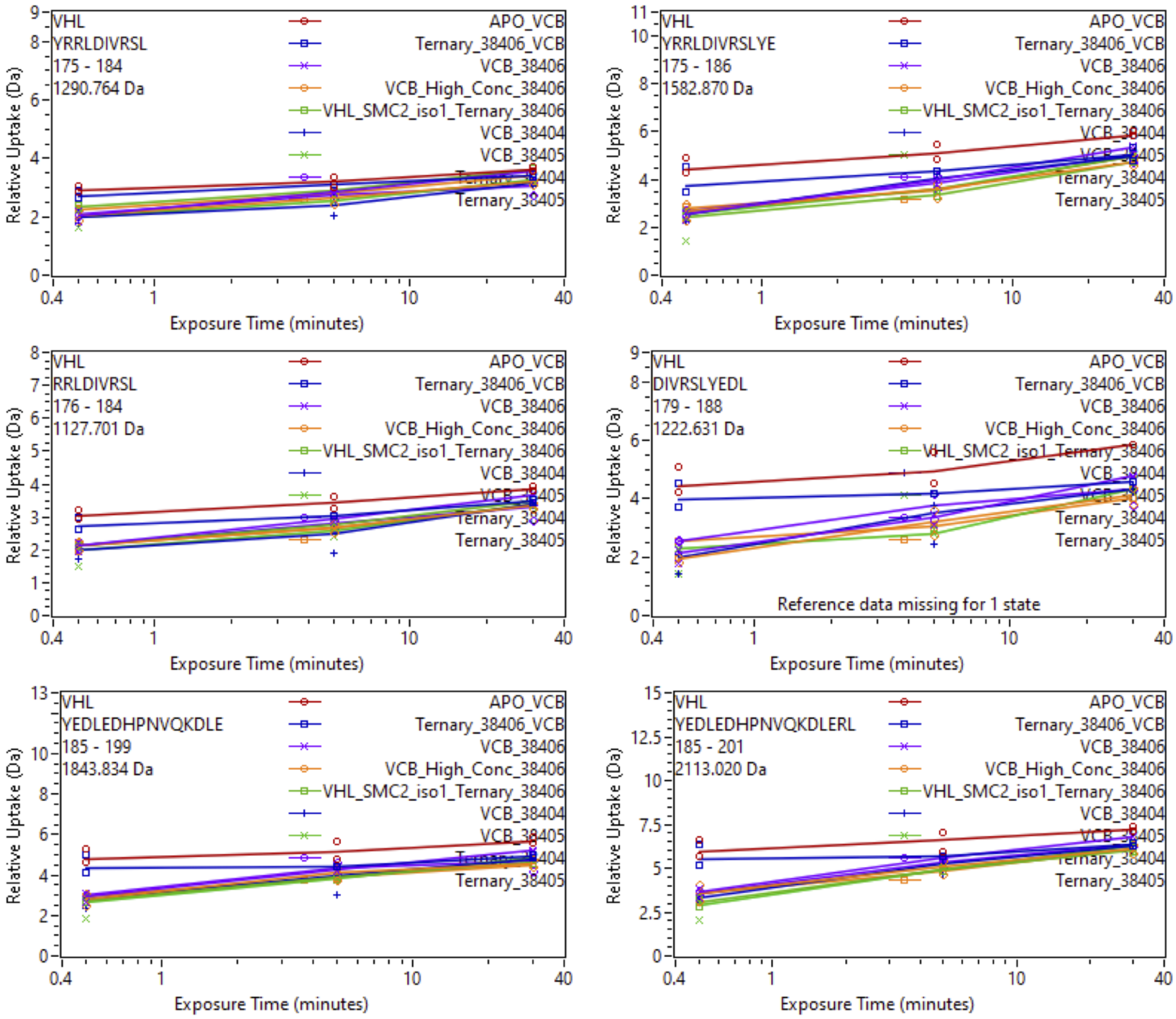


Supplemental Fig. 88: Relative deuterium uptake plots of peptic peptides of VHL of the VCB complex in the APO, Binary with SiTX-0038404 (PROTAC 1), SiTX-0038405(PROTAC 2), SiTX-0038406(ACBI1) or Ternary complex with 404, 405, 406 + SMARCA2$^{BD}$.

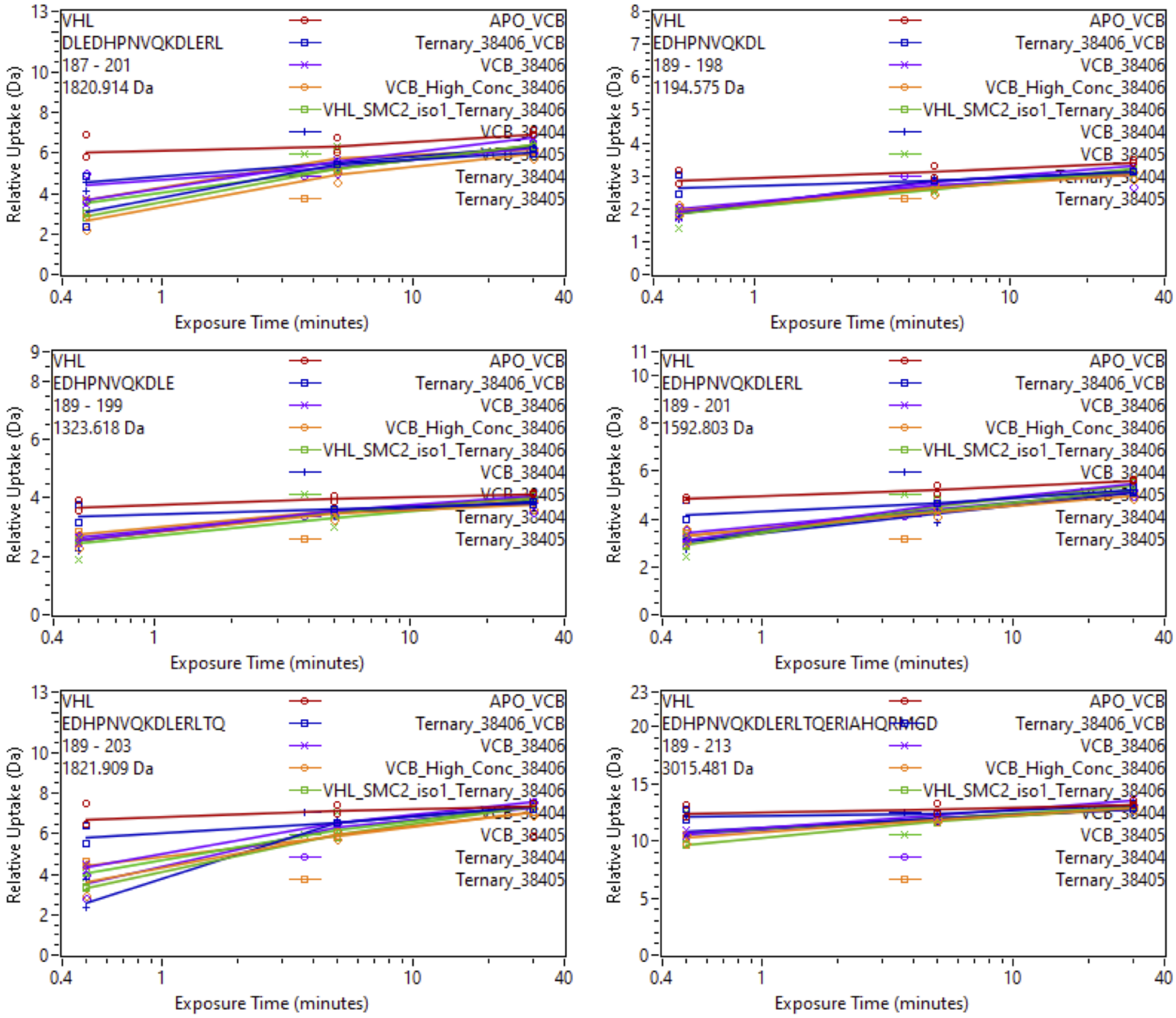


Supplemental Fig. 89: Relative deuterium uptake plots of peptic peptides of VHL of the VCB complex in the APO, Binary with SiTX-0038404 (PROTAC 1), SiTX-0038405(PROTAC 2), SiTX-0038406(ACBI1) or Ternary complex with 404, 405, 406 + SMARCA2$^{BD}$.

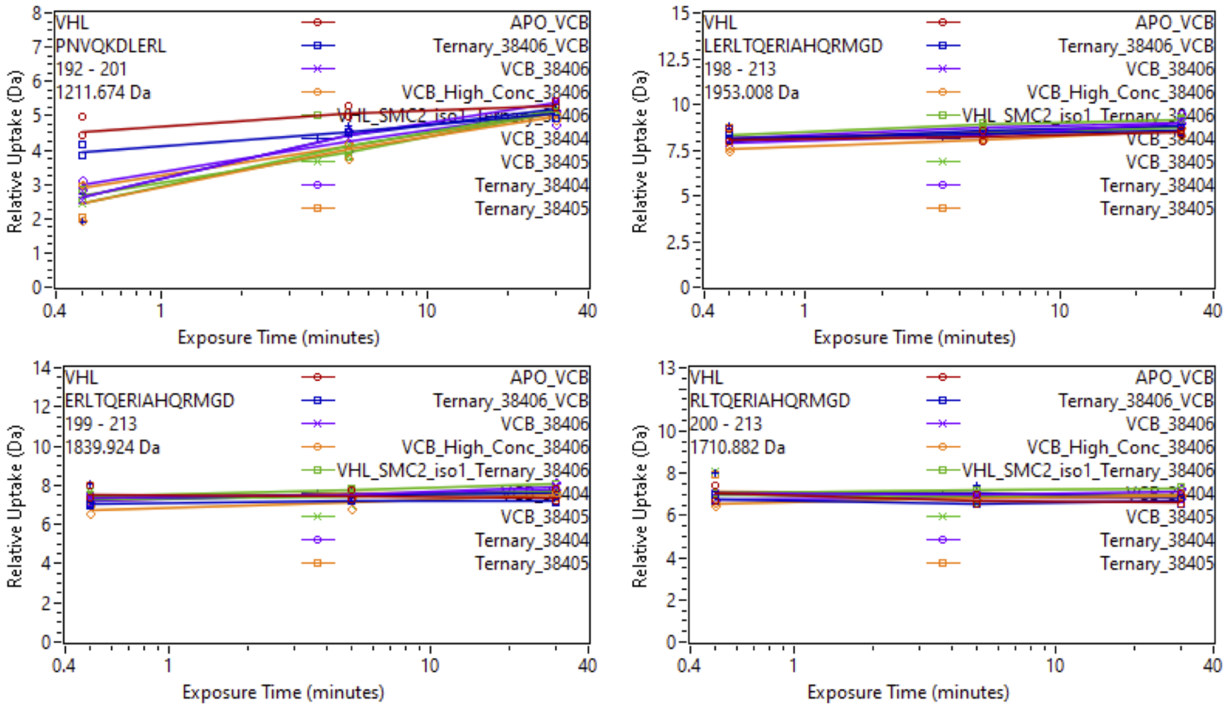


Supplemental Fig. 90: Relative deuterium uptake plots of peptic peptides of VHL of the VCB complex in the APO, Binary with SiTX-0038404 (PROTAC 1), SiTX-0038405(PROTAC 2), SiTX-0038406(ACBI1) or Ternary complex with 404, 405, 406 + SMARCA2$^{BD}$.

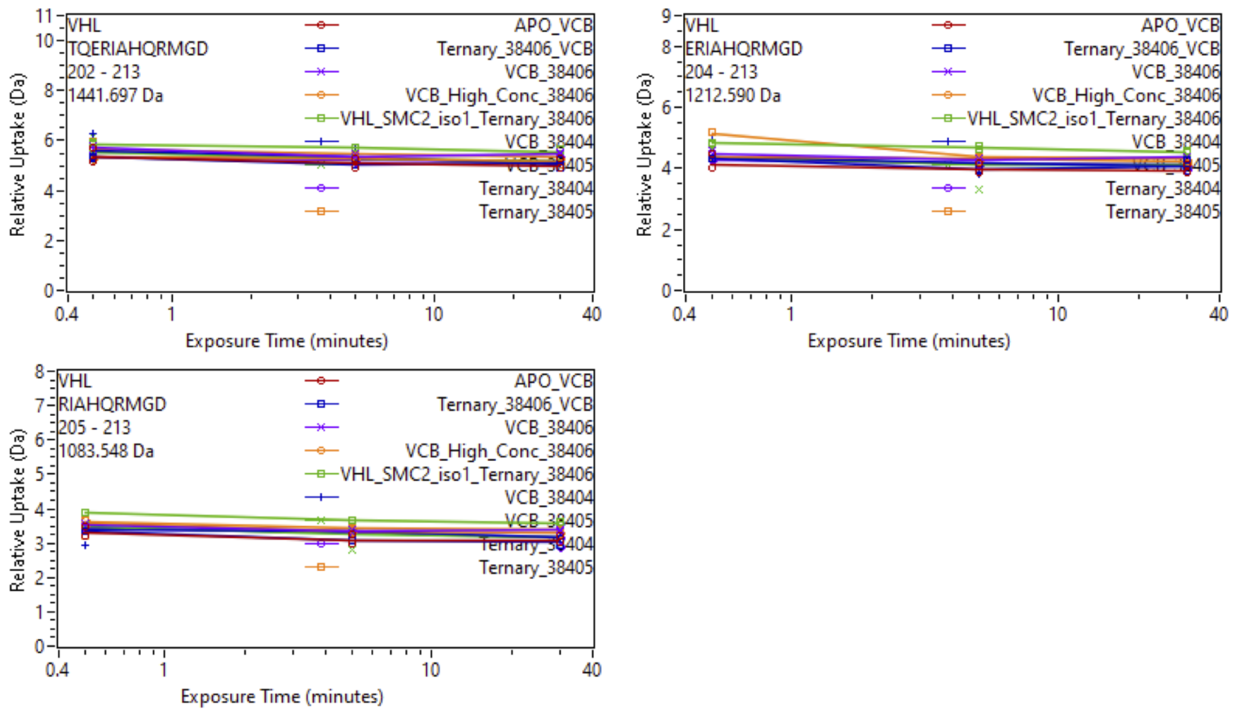


Supplemental Fig. 91: Relative deuterium uptake plots of peptic peptides of VHL of the VCB complex in the APO, Binary with SiTX-0038404 (PROTAC 1), SiTX-0038405(PROTAC 2), SiTX-0038406(ACBI1) or Ternary complex with 404, 405, 406 + SMARCA2$^{BD}$.

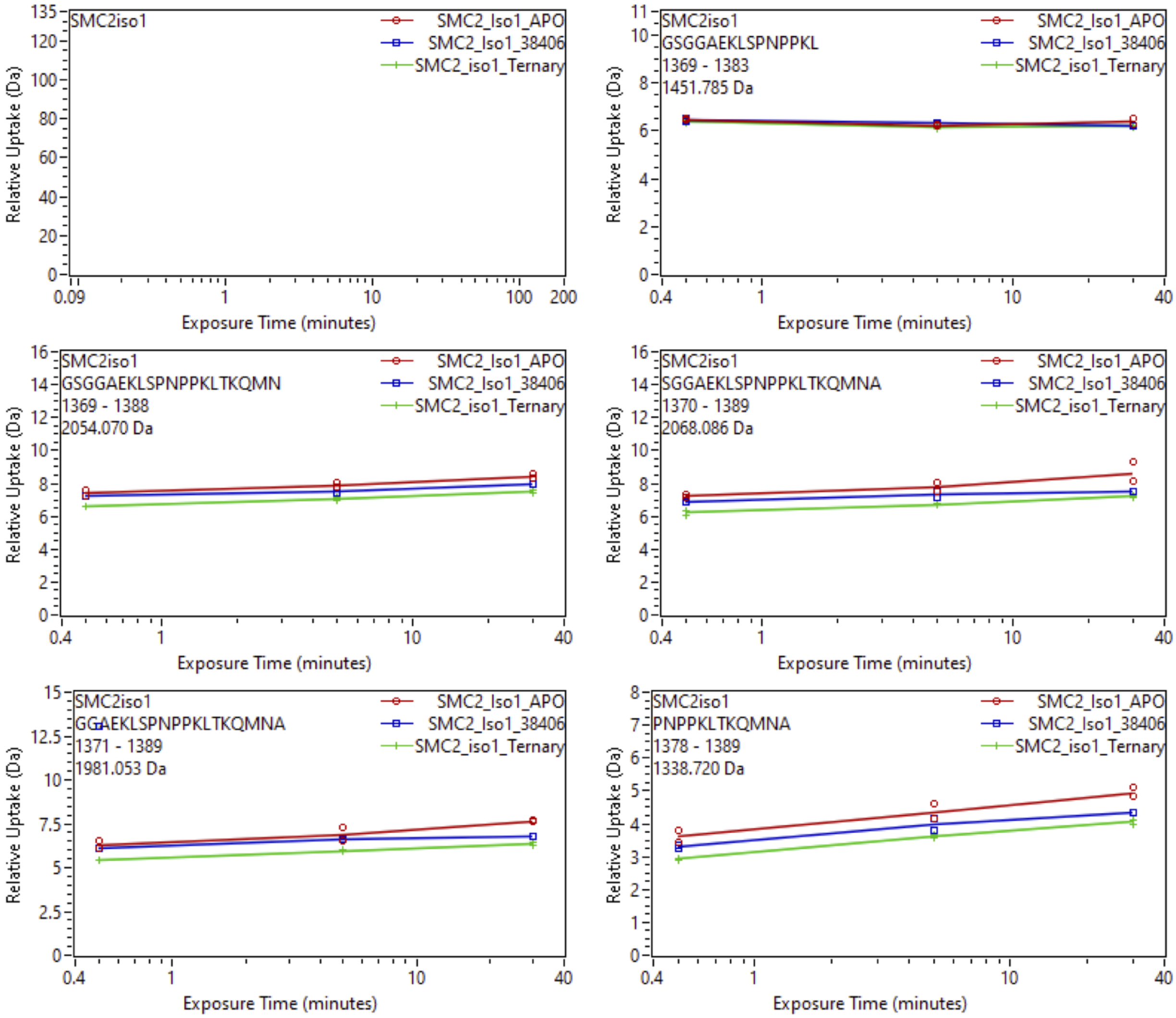


Supplemental Fig. 92: Relative deuterium uptake plots of peptic peptides of iso1-SMARCA2$^{BD}$ in the APO, Binary with SiTX-0038404 (PROTAC 1), SiTX-0038405(PROTAC 2), SiTX-0038406(ACBI1) or Ternary complex with 404, 405, 406 + VCB.

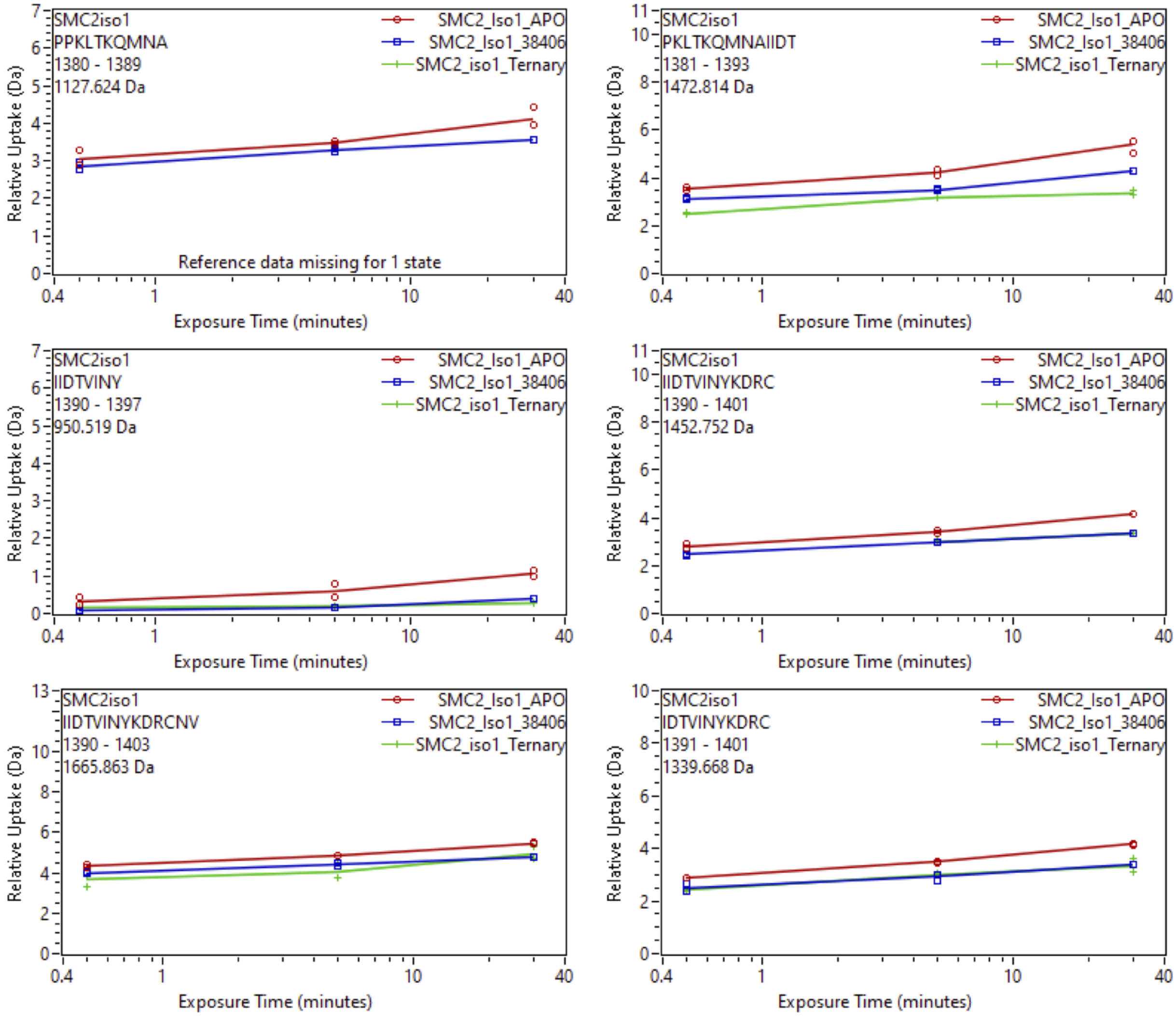


Supplemental Fig. 93: Relative deuterium uptake plots of peptic peptides of iso1-SMARCA2$^{BD}$ in the APO, Binary with SiTX-0038404 (PROTAC 1), SiTX-0038405(PROTAC 2), SiTX-0038406(ACBI1) or Ternary complex with 404, 405, 406 + VCB.

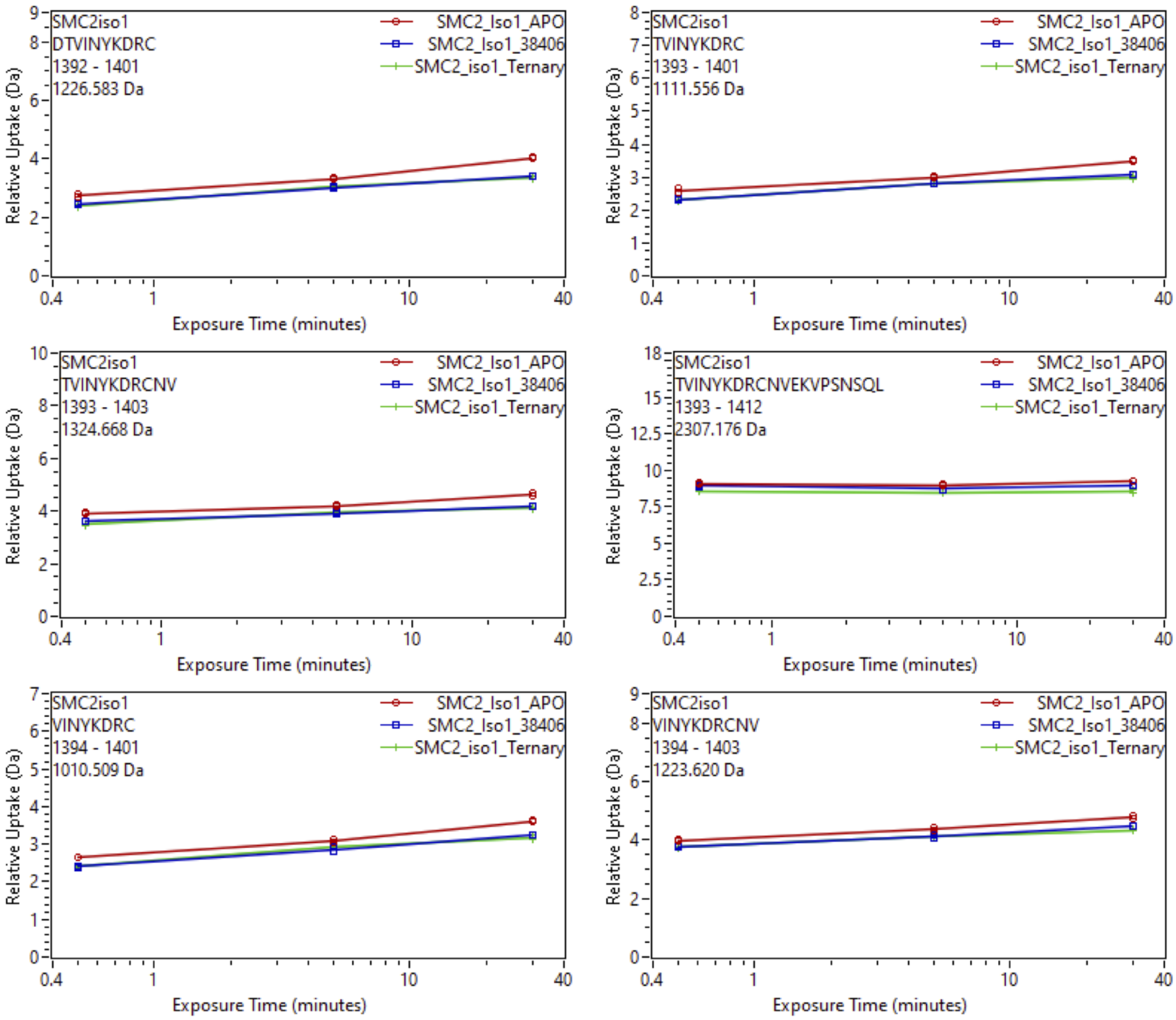


Supplemental Fig. 94: Relative deuterium uptake plots of peptic peptides of iso1-SMARCA2$^{BD}$ in the APO, Binary with SiTX-0038404 (PROTAC 1), SiTX-0038405(PROTAC 2), SiTX-0038406(ACBI1) or Ternary complex with 404, 405, 406 + VCB.

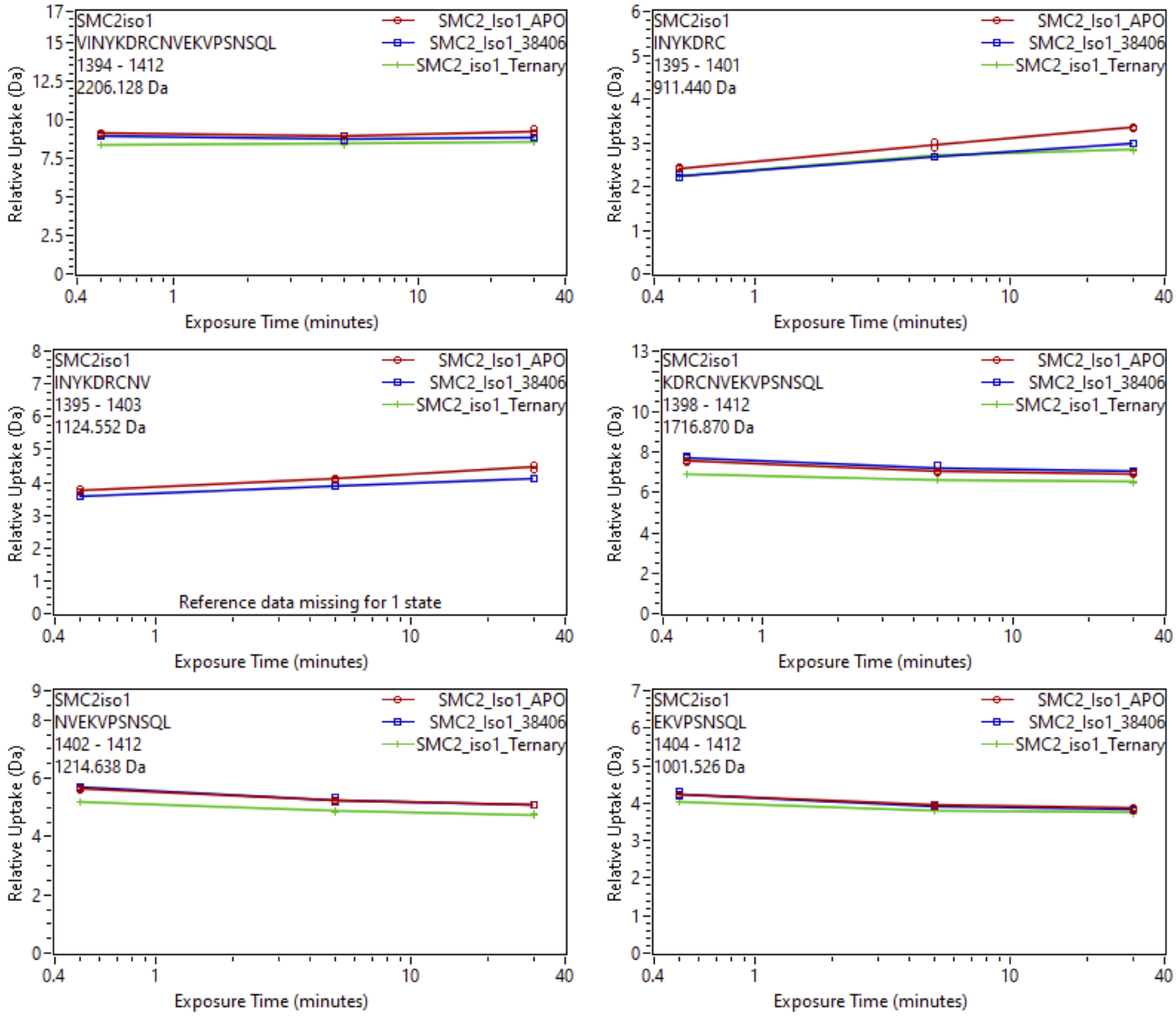


Supplemental Fig. 95: Relative deuterium uptake plots of peptic peptides of iso1-SMARCA2$^{BD}$ in the APO, Binary with SiTX-0038404 (PROTAC 1), SiTX-0038405(PROTAC 2), SiTX-0038406(ACBI1) or Ternary complex with 404, 405, 406 + VCB.

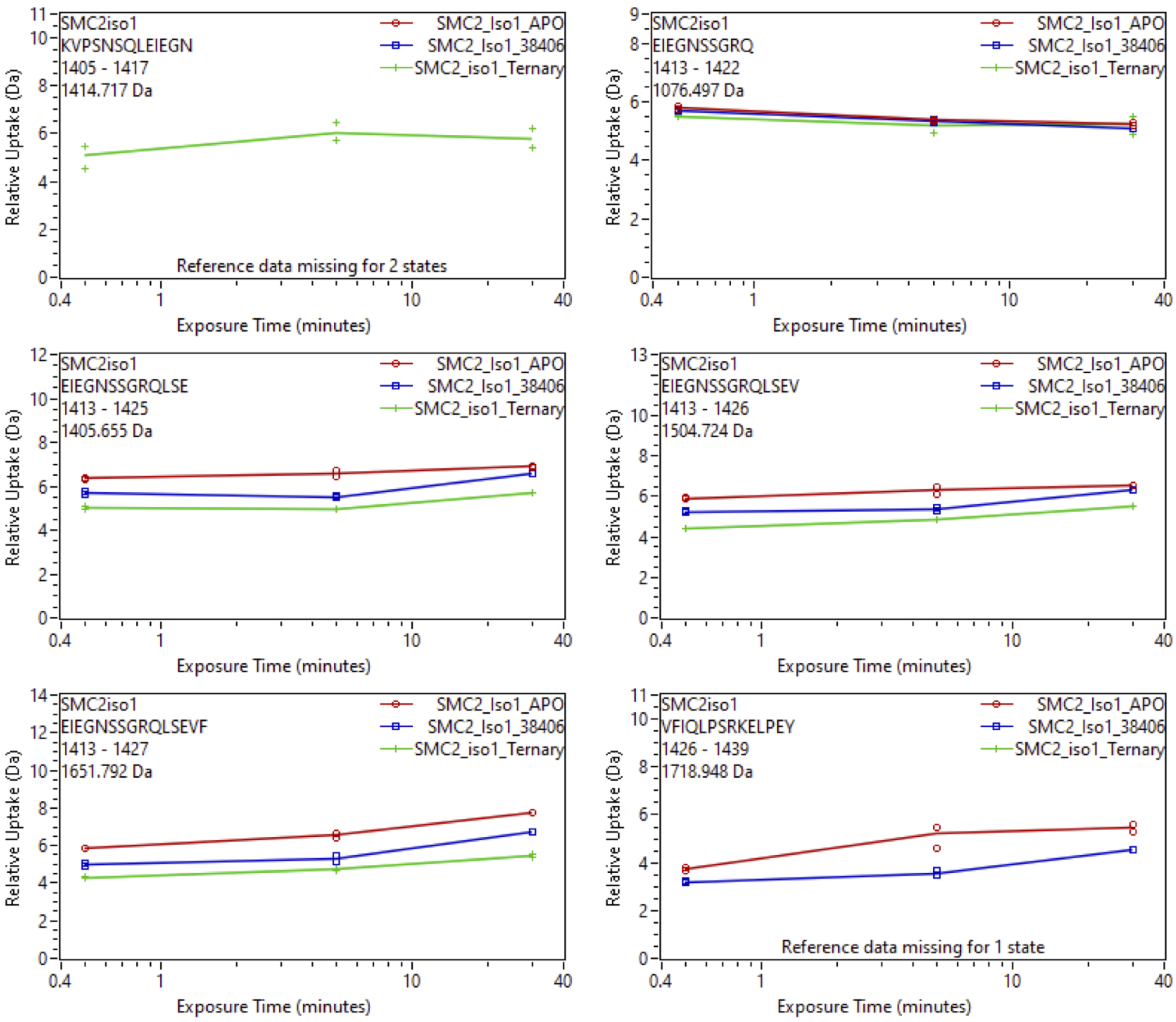


Supplemental Fig. 96: Relative deuterium uptake plots of peptic peptides of iso1-SMARCA2$^{BD}$ in the APO, Binary with SiTX-0038404 (PROTAC 1), SiTX-0038405(PROTAC 2), SiTX-0038406(ACBI1) or Ternary complex with 404, 405, 406 + VCB.

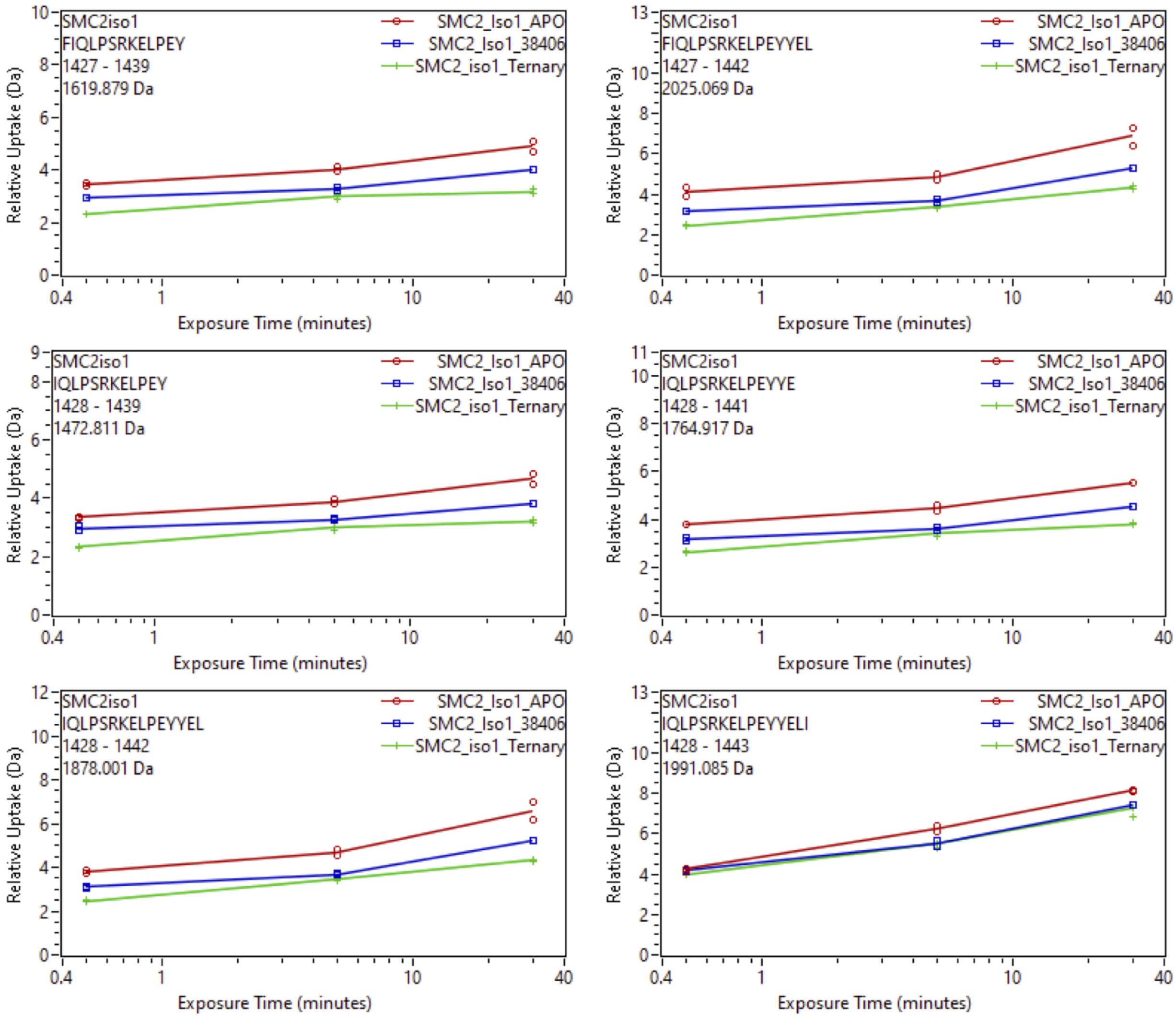


Supplemental Fig. 97: Relative deuterium uptake plots of peptic peptides of iso1-SMARCA2$^{BD}$ in the APO, Binary with SiTX-0038404 (PROTAC 1), SiTX-0038405(PROTAC 2), SiTX-0038406(ACBI1) or Ternary complex with 404, 405, 406 + VCB.

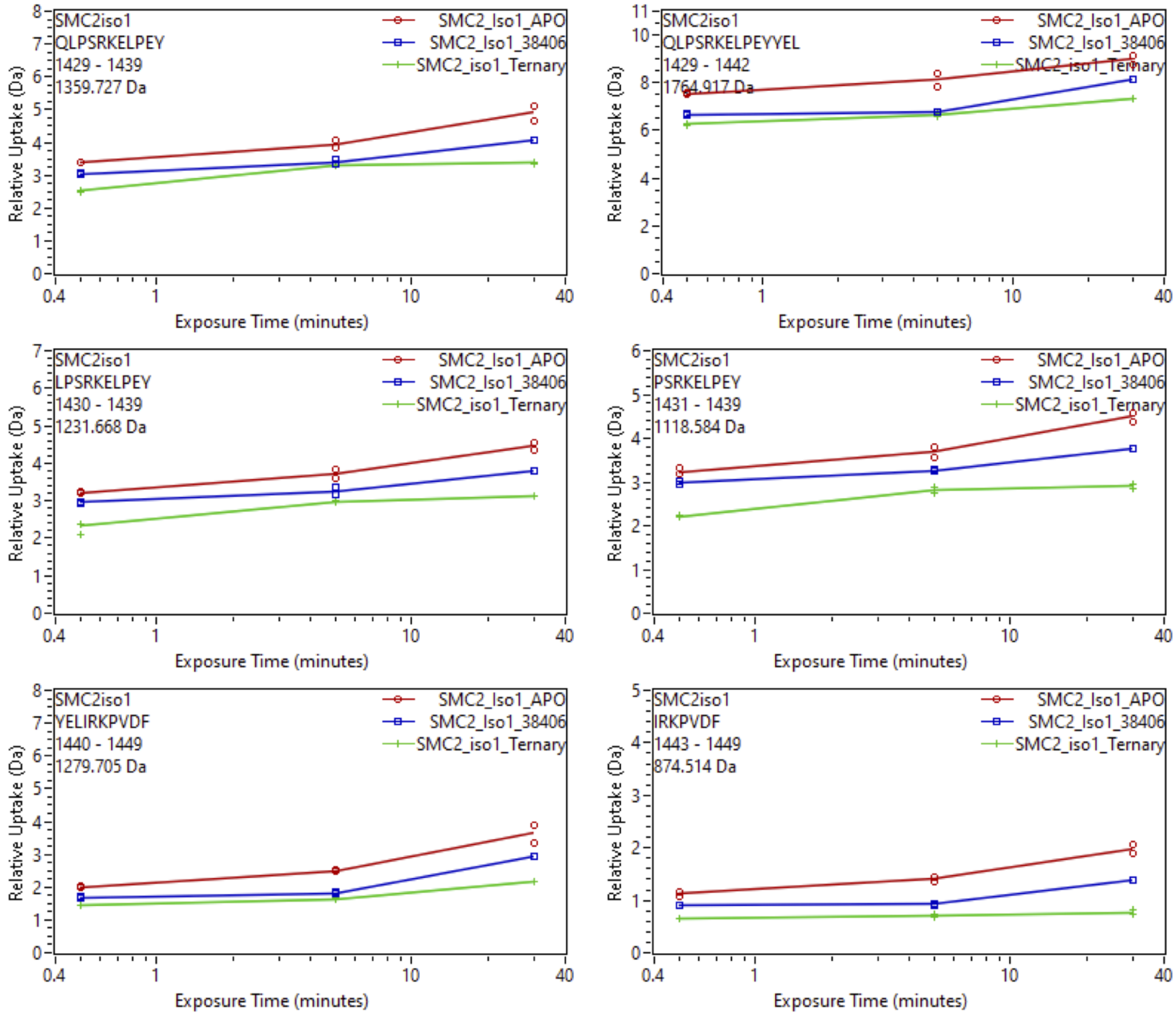


Supplemental Fig. 98: Relative deuterium uptake plots of peptic peptides of iso1-SMARCA2$^{BD}$ in the APO, Binary with SiTX-0038404 (PROTAC 1), SiTX-0038405(PROTAC 2), SiTX-0038406(ACBI1) or Ternary complex with 404, 405, 406 + VCB.

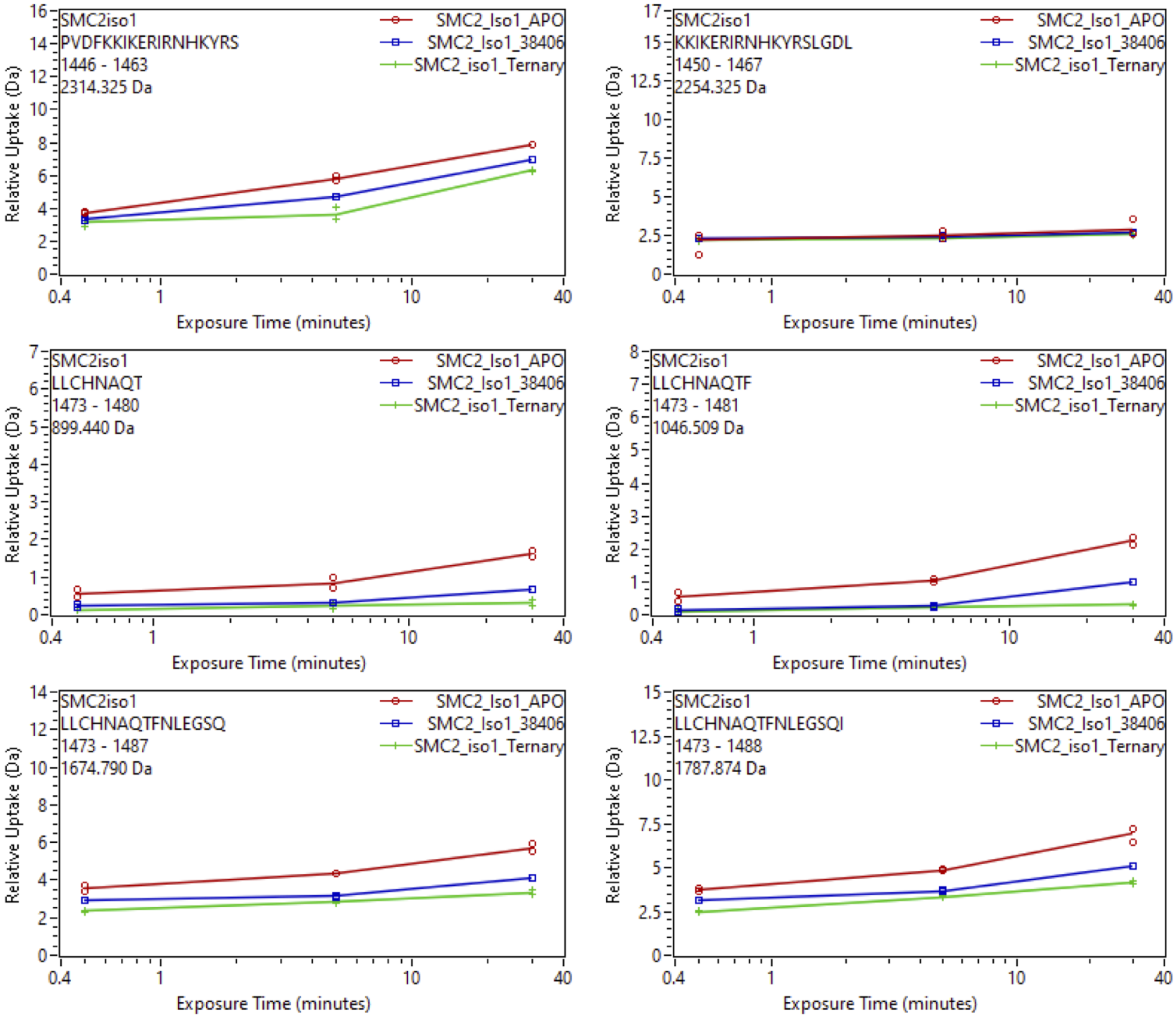


Supplemental Fig. 99: Relative deuterium uptake plots of peptic peptides of iso1-SMARCA2$^{BD}$ in the APO, Binary with SiTX-0038404 (PROTAC 1), SiTX-0038405(PROTAC 2), SiTX-0038406(ACBI1) or Ternary complex with 404, 405, 406 + VCB.

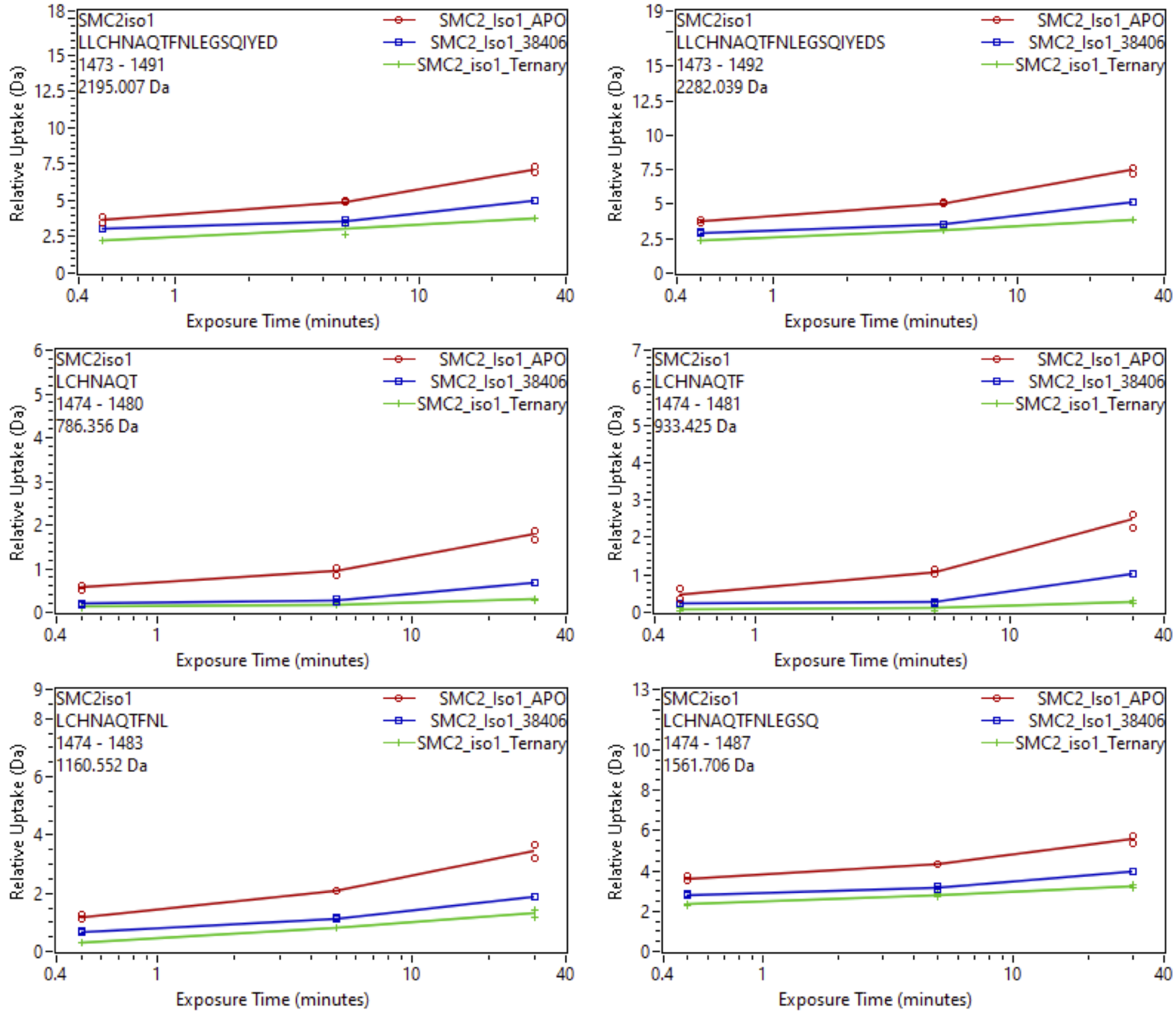


Supplemental Fig. 100: Relative deuterium uptake plots of peptic peptides of iso1-SMARCA2$^{BD}$ in the APO, Binary with SiTX-0038404 (PROTAC 1), SiTX-0038405(PROTAC 2), SiTX-0038406(ACBI1) or Ternary complex with 404, 405, 406 + VCB.

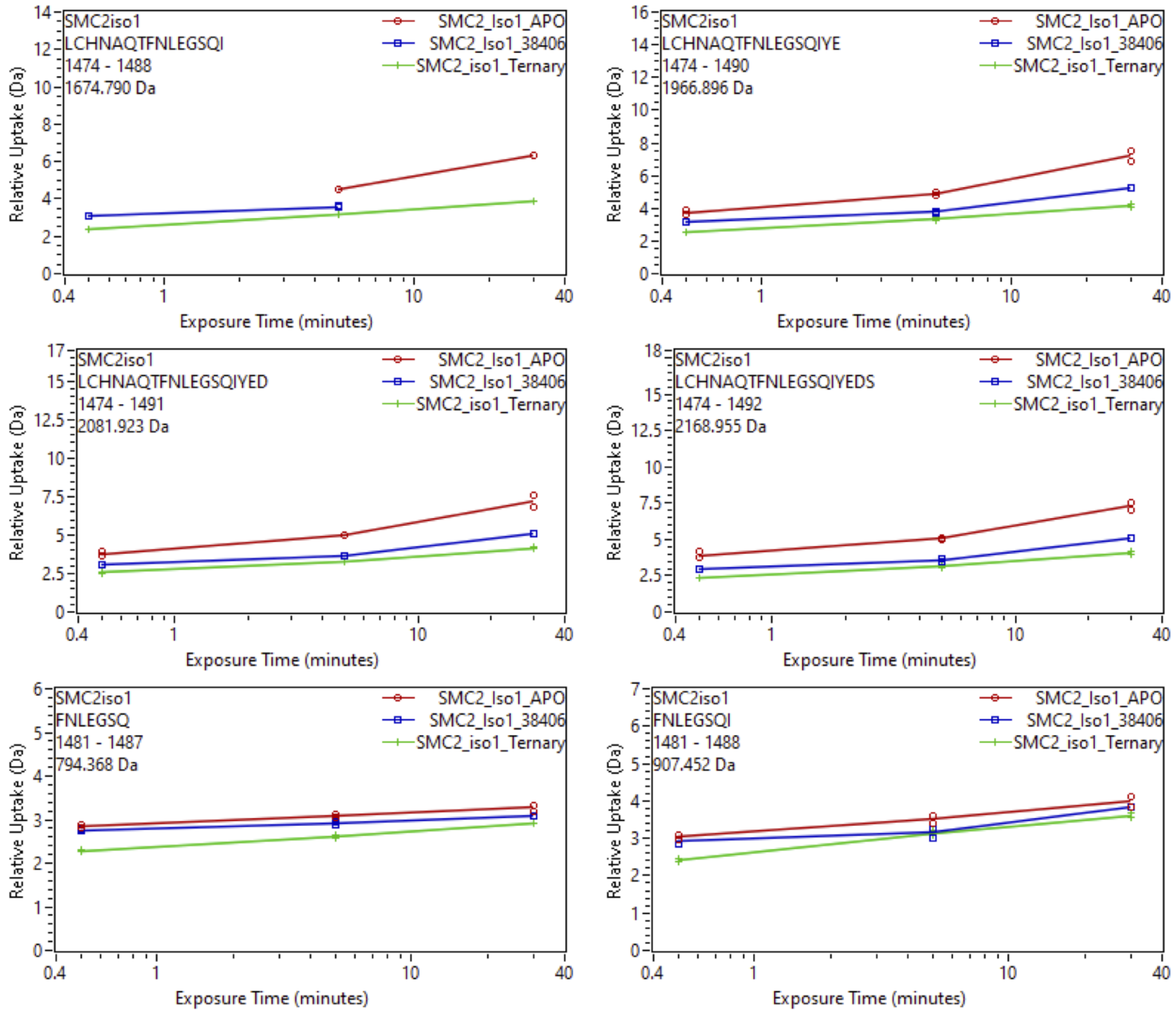


Supplemental Fig. 101: Relative deuterium uptake plots of peptic peptides of iso1-SMARCA2$^{BD}$ in the APO, Binary with SiTX-0038404 (PROTAC 1), SiTX-0038405(PROTAC 2), SiTX-0038406(ACBI1) or Ternary complex with 404, 405, 406 + VCB.

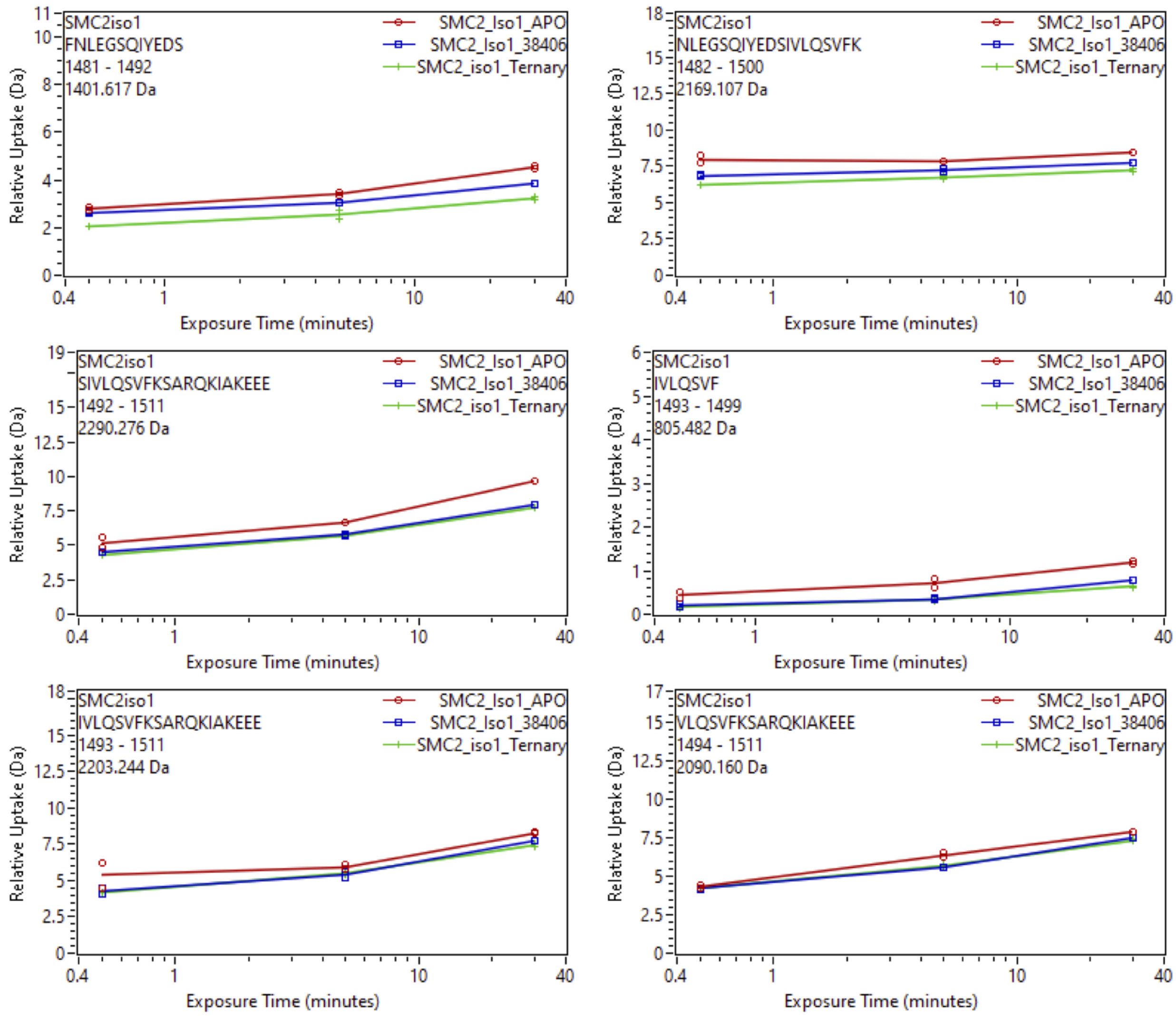


Supplemental Fig. 102: Relative deuterium uptake plots of peptic peptides of iso1-SMARCA2$^{BD}$ in the APO, Binary with SiTX-0038404 (PROTAC 1), SiTX-0038405(PROTAC 2), SiTX-0038406(ACBI1) or Ternary complex with 404, 405, 406 + VCB.

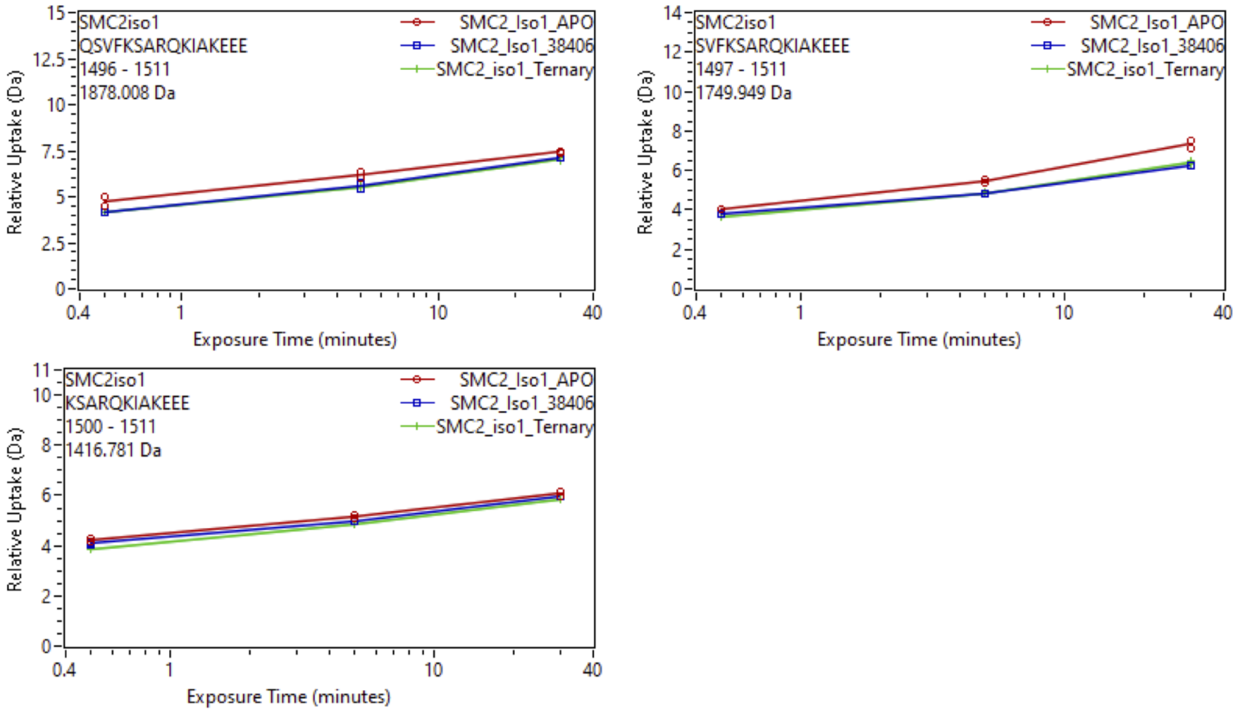


Supplemental Fig. 103: Relative deuterium uptake plots of peptic peptides of iso1-SMARCA2$^{BD}$ in the APO, Binary with SiTX-0038404 (PROTAC 1), SiTX-0038405(PROTAC 2), SiTX-0038406(ACBI1) or Ternary complex with 404, 405, 406 + VCB.

### References


(1) Donyapour, N.; Roussey, N. M.; Dickson, A. REVO: Resampling of ensembles by variation optimization. *J. Chem. Phys.* **2019**, *150*.

(2) Saglam, A. S.; Chong, L. T. Protein–protein binding pathways and calculations of rate constants using fully-continuous, explicit-solvent simulations. *Chemical Science* **2018**, *10*, 2360–2372.

(3) Drummond, M. L.; Henry, A.; Li, H.; Williams, C. I. Improved Accuracy for Modeling PROTAC-Mediated Ternary Complex Formation and Targeted Protein Degradation via New In Silico Methodologies. *J CHEM INF MODEL* **2020**, *60*, 5234–5254.

(4) Devaurs, D.; Antunes, D. A.; Borysik, A. J. Computational Modeling of Molecular Structures Guided by Hydrogen-Exchange Data. *Journal of the American Society for Mass Spectrometry* **2022**, *33*, 215–237, PMID: 35077179.

(5) Anand, G. S.; Law, D.; Mandell, J. G.; Snead, A. N.; Tsigelny, I.; Taylor, S. S.; Eyck, L. F. T.; Komives, E. A. Identification of the protein kinase A regulatory R I -catalytic subunit interface by amide H/ 2 H exchange and protein docking. *Proceedings of the National Academy of Sciences* **2003**, *100*, 13264–13269.

(6) Zhang, M. M.; Beno, B. R.; Huang, R. Y.-C.; Adhikari, J.; Deyanova, E. G.; Li, J.; Chen, G.; Gross, M. L. An Integrated Approach for Determining a Protein–Protein Binding Interface in Solution and an Evaluation of Hydrogen–Deuterium Exchange Kinetics for Adjudicating Candidate Docking Models. *Anal. Chem.* **2019**, *91*, 15709–15717.

(7) Rampler, E.; Stranzl, T.; Orban-Nemeth, Z.; Hollenstein, D. M.; Hudecz, O.; Schlögelhofer, P.; Mechtler, K. Comprehensive Cross-Linking Mass Spectrometry Reveals Parallel Orientation and Flexible Conformations of Plant HOP2-MND1. *Journal of proteome research* **2015**, *14*, 5048–62.

(8) Lin, S.-J.; Chen, Y.-F.; Hsu, K.-C.; Chen, Y.-L.; Ko, T.-P.; Lo, C.-F.; Wang, H.-C.; Wang, H.-C. Structural Insights to the Heterotetrameric Interaction between the Vibrio parahaemolyticus PirAvp and PirBvp Toxins and Activation of the Cry-Like Pore-Forming Domain. *Toxins* **2019**, *11*, 233.

(9) Pandit, D.; Tuske, S. J.; Coales, S. J.; E, S. Y.; Liu, A.; Lee, J. E.; Morrow, J. A.; Nemeth, J. F.; Hamuro, Y. Mapping of discontinuous conformational epitopes by amide hydrogen/deuterium exchange mass spectrometry and computational docking. *Journal of molecular recognition : JMR* **2012**, *25*, 114–24.

(10) Roberts, V. A.; Pique, M. E.; Hsu, S.; Li, S. Combining H/D Exchange Mass Spectrometry and Computational Docking To Derive the Structure of Protein–Protein Complexes. *Biochemistry* **2017**, *56*, 6329–6342.

(11) Rey, M.; Sarpe, V.; Burns, K. M.; Buse, J.; Baker, C. A. H.; van Dijk, M.; Wordeman, L.; Bonvin, A. M. J. J.; Schriemer, D. C. Mass Spec Studio for Integrative Structural Biology. *Structure* **2014**, *22*, 1538–1548.

(12) Merkle, P. S.; Irving, M.; Hongjian, S.; Ferber, M.; Jørgensen, T. J. D.; Scholten, K.; Luescher, I.; Coukos, G.; Zoete, V.; Cuendet, M. A.; Michielin, O.; Rand, K. D. The T-Cell Receptor Can Bind to the Peptide-Bound Major Histocompatibility Complex and Uncomplexed 2 -Microglobulin through Distinct Binding Sites. *Biochemistry* **2017**, *56*, 3945–3961.

(13) Komolov, K. E.; Du, Y.; Duc, N. M.; Betz, R. M.; Rodrigues, J. P. G. L. M.; Leib, R. D.; Patra, D.; Skiniotis, G.; Adams, C. M.; Dror, R. O.; Chung, K. Y.; Kobilka, B. K.; Benovic, J. L. Structural and Functional Analysis of a 2 -Adrenergic Receptor Complex with GRK5. *Cell* **2017**, *169*, 407–421.e16.

(14) Eron, S. J.; Huang, H.; Agafonov, R. V.; Fitzgerald, M. E.; Patel, J.; Michael, R. E.; Lee, T. D.; Hart, A. A.; Shaulsky, J.; Nasveschuk, C. G.; Phillips, A. J.; Fisher, S. L.;

Good, A. Structural Characterization of Degrader-Induced Ternary Complexes Using Hydrogen–Deuterium Exchange Mass Spectrometry and Computational Modeling: Implications for Structure-Based Design. *ACS Chemical Biology* **2021**,

(15) Brodie, N. I.; Popov, K. I.; Petrotchenko, E. V.; Dokholyan, N. V.; Borchers, C. H. Solving protein structures using short-distance cross-linking constraints as a guide for discrete molecular dynamics simulations. *Science advances* **2017**, *3*, e1700479.

(16) Marsh, J. A.; Forman-Kay, J. D. Structure and disorder in an unfolded state under nondenaturing conditions from ensemble models consistent with a large number of experimental restraints. *Journal of molecular biology* **2009**, *391*, 359–74.

(17) Martens, C.; Shekhar, M.; Lau, A. M.; Tajkhorshid, E.; Politis, A. Integrating hydrogen–deuterium exchange mass spectrometry with molecular dynamics simulations to probe lipid-modulated conformational changes in membrane proteins. *Nature Protocols* **2019**, *14*, 3183–3204.

(18) Jia, R.; Martens, C.; Shekhar, M.; Pant, S.; Pellowe, G. A.; Lau, A. M.; Findlay, H. E.; Harris, N. J.; Tajkhorshid, E.; Booth, P. J.; Politis, A. Hydrogen-deuterium exchange mass spectrometry captures distinct dynamics upon substrate and inhibitor binding to a transporter. *Nature Communications* **2020**, *11*, 6162.

(19) Zhang, H. et al. Structure of the full-length glucagon class B G-protein-coupled receptor. *Nature* **2017**, *546*, 259–264.

(20) Harrison, R. A.; Lu, J.; Carrasco, M.; Hunter, J.; Manandhar, A.; Gondi, S.; Westover, K. D.; Engen, J. R. Structural Dynamics in Ras and Related Proteins upon Nucleotide Switching. *Journal of Molecular Biology* **2016**, *428*, 4723–4735.

(21) Xiao, Y.; Shaw, G. S.; Konermann, L. Calcium-Mediated Control of S100 Proteins: Allosteric Communication via an Agitator/Signal Blocking Mechanism. *Journal of the American Chemical Society* **2017**, *139*, 11460–11470.

(22) Singh, J.; Udgaonkar, J. B. Unraveling the Molecular Mechanism of pH-Induced Misfolding and Oligomerization of the Prion Protein. *Journal of Molecular Biology* **2016**, *428*, 1345–1355.

(23) Petruk, A. A.; Defelipe, L. A.; Limardo, R. G. R.; Bucci, H.; Marti, M. A.; Turjanski, A. G. Molecular Dynamics Simulations Provide Atomistic Insight into Hydrogen Exchange Mass Spectrometry Experiments. *Journal of Chemical Theory and Computation* **2012**, *9*, 658–669.

(24) Shan, Y.; Arkhipov, A.; Kim, E. T.; Pan, A. C.; Shaw, D. E. Transitions to catalytically inactive conformations in EGFR kinase. *Proceedings of the National Academy of Sciences* **2013**, *110*, 7270–7275.

(25) Huang, L.; So, P.-K.; Yao, Z.-P. Protein Dynamics Revealed by Hydrogen Deuterium Exchange Mass Spectrometry: Correlation between Experiments and Simulation. *Rapid communications in mass spectrometry : RCM* **2018**, *33*, 83–89.

(26) Sheinerman, F. B.; Brooks, C. L. Molecular picture of folding of a small / protein. *Proceedings of the National Academy of Sciences* **1998**, *95*, 1562–1567.

(27) McAllister, R. G.; Konermann, L. Challenges in the Interpretation of Protein H/D Exchange Data: A Molecular Dynamics Simulation Perspective. *Biochemistry* **2015**, *54*, 2683–2692.

(28) Fazelinia, H.; Xu, M.; Cheng, H.; Roder, H. Ultrafast Hydrogen Exchange Reveals Specific Structural Events during the Initial Stages of Folding of Cytochrome c. *Journal of the American Chemical Society* **2013**, *136*, 733–740.

(29) Skinner, J. J.; Lim, W. K.; Bédard, S.; Black, B. E.; Englander, S. W. Protein dynamics viewed by hydrogen exchange: Protein Dynamics from Hydrogen Exchange. *Protein Science* **2012**, *21*, 996–1005.

(30) Ma, B.; Nussinov, R. Polymorphic triple beta-sheet structures contribute to amide hydrogen/deuterium (H/D) exchange protection in the Alzheimer amyloid beta42 peptide. *The Journal of biological chemistry* **2011**, *286*, 34244–53.

(31) Hernández, G.; Anderson, J. S.; LeMaster, D. M. Assessing the native state conformational distribution of ubiquitin by peptide acidity. *Biophysical Chemistry* **2010**, *153*, 70–82.

(32) Hernández, G.; Anderson, J. S.; LeMaster, D. M. Experimentally assessing molecular dynamics sampling of the protein native state conformational distribution. *Biophysical chemistry* **2012**, *163-164*, 21–34.

(33) Xu, J.; Lee, Y.; Beamer, L. J.; Doren, S. R. V. Phosphorylation in the catalytic cleft stabilizes and attracts domains of a phosphohexomutase. *Biophysical journal* **2015**, *108*, 325–37.

(34) Devaurs, D.; Antunes, D. A.; Papanastasiou, M.; Moll, M.; Ricklin, D.; Lambris, J. D.; Kavraki, L. E. Coarse-Grained Conformational Sampling of Protein Structure Improves the Fit to Experimental Hydrogen-Exchange Data. *Frontiers in Molecular Biosciences* **2017**, *4*, 13.

(35) Wan, H.; Ge, Y.; Razavi, A.; Voelz, V. A. Reconciling Simulated Ensembles of Apomyoglobin with Experimental Hydrogen/Deuterium Exchange Data Using Bayesian Inference and Multiensemble Markov State Models. *Journal of Chemical Theory and Computation* **2020**, *16*, 1333–1348.

(36) Radou, G.; Dreyer, F.; Tuma, R.; Paci, E. Functional Dynamics of Hexameric Helicase Probed by Hydrogen Exchange and Simulation. *Biophysical Journal* **2014**, *107*, 983–990.

(37) Adhikary, S.; Deredge, D. J.; Nagarajan, A.; Forrest, L. R.; Wintrode, P. L.; Singh, S. K. Conformational dynamics of a neurotransmitter:sodium symporter in a lipid bilayer.

*Proceedings of the National Academy of Sciences of the United States of America* **2017**, *114*, E1786–E1795.

(38) Borysik, A. J. Simulated Isotope Exchange Patterns Enable Protein Structure Determination. *Angewandte Chemie International Edition* **2017**, *56*, 9396–9399.

(39) Devaurs, D.; Papanastasiou, M.; Antunes, D. A.; Abella, J. R.; Moll, M.; Ricklin, D.; Lambris, J. D.; Kavraki, L. E. Native state of complement protein C3d analysed via hydrogen exchange and conformational sampling. *International Journal of Computational Biology and Drug Design* **2018**, *11*, 90.

(40) Markwick, P. R. L.; Peacock, R. B.; Komives, E. A. Accurate Prediction of Amide Exchange in the Fast Limit Reveals Thrombin Allostery. *Biophysical Journal* **2019**, *116*, 49–56.

(41) Aytenfisu, A. H.; Deredge, D.; Klontz, E. H.; Du, J.; Sundberg, E. J.; MacKerell, A. D. Insights into substrate recognition and specificity for IgG by Endoglycosidase S2. *PLoS computational biology* **2021**, *17*, e1009103.

(42) Kihn, K. C.; Wilson, T.; Smith, A. K.; Bradshaw, R. T.; Wintrode, P. L.; Forrest, L. R.; Wilks, A.; Deredge, D. J. Modeling the native ensemble of PhuS using enhanced sampling MD and HDX-ensemble reweighting. *Biophysical journal* **2021**, *120*, 5141–5157.

(43) Bai, N.; Kirubakaran, P.; Karanicolas, J. Rationalizing PROTAC-mediated ternary complex formation using Rosetta. *J. Chem. Inf. Model.* **2021**, *61*, 1368–1382.

(44) Pracht, P.; Bohle, F.; Grimme, S. Automated exploration of the low-energy chemical space with fast quantum chemical methods. *Phys. Chem. Chem. Phys.* **2020**, *22*, 7169–7192.

(45) Méndez, R.; Leplae, R.; De Maria, L.; Wodak, S. J. Assessment of blind predictions of

protein–protein interactions: Current status of docking methods. *PROTEINS* **2003**, *52*, 51–67.

(46) Bussi, G. Hamiltonian replica exchange in GROMACS: a flexible implementation. *MOL PHYS* **2014**, *112*, 379–384.

(47) Wang, L.; Friesner, R. A.; Berne, B. J. Replica exchange with solute scaling: a more efficient version of replica exchange with solute tempering (REST2). *J PHYS CHEM B* **2011**, *115*, 9431–9438.

(48) Shrestha, U. R.; Juneja, P.; Zhang, Q.; Gurumoorthy, V.; Borreguero, J. M.; Urban, V.; Cheng, X.; Pingali, S. V.; Smith, J. C.; O'Neill, H. M.; Petridis, L. Generation of the configurational ensemble of an intrinsically disordered protein from unbiased molecular dynamics simulation. *P NATL ACAD SCI USA* **2019**, *116*, 20446–20452.

(49) Shrestha, U. R.; Smith, J. C.; Petridis, L. Full structural ensembles of intrinsically disordered proteins from unbiased molecular dynamics simulations. *Communications Biology* **2021**, *4*, 243.

(50) Cong, X.; Golebiowski, J. Allosteric Na+-binding site modulates CXCR4 activation. *Phys. Chem. Chem. Phys.* **2018**, *20*, 24915–24920.

(51) Cong, X.; Chéron, J.-B.; Golebiowski, J.; Antonczak, S.; Fiorucci, S. Allosteric Modulation Mechanism of the mGluR5 Transmembrane Domain. *Journal of Chemical Information and Modeling* **2019**, *59*, 2871–2878, PMID: 31025859.

(52) Abraham, M. J.; Murtola, T.; Schulz, R.; Páll, S.; Smith, J. C.; Hess, B.; Lindahl, E. GROMACS: High performance molecular simulations through multi-level parallelism from laptops to supercomputers. *SoftwareX* **2015**, *1-2*, 19–25.

(53) Páll, S.; Abraham, M. J.; Kutzner, C.; Hess, B.; Lindahl, E. Tackling Exascale Soft-

ware Challenges in Molecular Dynamics Simulations with GROMACS. Solving Software Challenges for Exascale. Cham, 2015; pp 3–27.

(54) Hess, B.; Kutzner, C.; van der Spoel, D.; Lindahl, E. GROMACS 4: Algorithms for Highly Efficient, Load-Balanced, and Scalable Molecular Simulation. *J CHEM THEORY COMPUT* **2008**, *4*, 435–447.

(55) Bonomi, M.; Branduardi, D.; Bussi, G.; Camilloni, C.; Provasi, D.; Raiteri, P.; Donadio, D.; Marinelli, F.; Pietrucci, F.; Broglia, R. A.; Parrinello, M. PLUMED: A portable plugin for free-energy calculations with molecular dynamics. *COMPUT PHYS COMMUN* **2009**, *180*, 1961–1972.

(56) Bonomi, M. Promoting transparency and reproducibility in enhanced molecular simulations. *Nature methods* **2019**, *16*, 670–673.

(57) Tribello, G. A.; Bonomi, M.; Branduardi, D.; Camilloni, C.; Bussi, G. PLUMED 2: New feathers for an old bird. *COMPUT PHYS COMMUN* **2014**, *185*, 604–613.

(58) Sugita, Y.; Okamoto, Y. Replica-exchange molecular dynamics method for protein folding. *CHEM PHYS LETT* **1999**, *314*, 141–151.

(59) Okabe, T.; Kawata, M.; Okamoto, Y.; Mikami, M. Replica-exchange Monte Carlo method for the isobaric–isothermal ensemble. *CHEM PHYS LETT* **2001**, *335*, 435–439.
